# Click-Based Determination of Accumulation of Molecules in *Escherichia coli*

**DOI:** 10.1101/2023.06.20.545103

**Authors:** George M. Ongwae, Zichen Liu, Shasha Feng, Mahendra D. Chordia, Rachita Dash, Yuchen He, Mohammad Sharifian Gh, Brianna E. Dalesandro, Taijie Guo, Karl Barry Sharpless, Jiajia Dong, M. Sloan Siegrist, Wonpil Im, Marcos M. Pires

## Abstract

Gram-negative bacterial pathogens pose a significant challenge in drug development due to their outer membranes, which hinder the permeation of small molecules. The lack of widely adoptable methods to measure the cytosolic accumulation of compounds in bacterial cells has hindered drug discovery efforts. To address this challenge, we developed the CHloroalkane Azide Membrane Permeability (CHAMP) assay, specifically designed to assess molecule accumulation in the cytosol of Gram-negative bacteria. The CHAMP analysis utilizes biorthogonal epitopes anchored within HaloTag-expressing bacteria and measures the cytosolic arrival of azide-bearing test molecules through strain-promoted azide-alkyne cycloaddition. This workflow allows for robust and rapid accumulation measurements of thousands of azide-tagged small molecules. Our approach consistently produces comprehensive accumulation profiles, significantly surpassing the scale of previous measurements in *Escherichia coli* (*E. coli*). We have validated the CHAMP assay across various chemical and biological contexts, including hyperporinated cells, membrane-permeabilized cells, and *E. coli* strains with impaired TolC function, a key component of the efflux pump. The CHAMP platform provides a simple, high-throughput, and accessible method that enables the analysis of over 1,000 molecules within hours. This technique addresses a critical gap in antimicrobial research and has the potential to accelerate the development of effective agents against Gram-negative pathogens.

## MAIN

The increased prevalence of multidrug resistant Gram-negative bacterial infections is extremely alarming.^1^ The CDC issued a report in 2019 that categorized bacterial pathogens based on their threat level. The list of pathogens considered as “urgent threats” was primarily populated by antibiotic-resistant Gram-negative bacteria and included *E. coli*.^2^ Lack of treatment options will usher in the post-antibiotic era, meaning that routine infections can become lethal and standard invasive medical procedures carry a much higher level of risk. Therefore, strategies that aim to address these threats have high significance. We pose, and there is wide agreement in the community^3–12^, that the lack of robust and widely adoptable tools to measure the accumulation of molecules into bacteria has severely hampered antibiotic drug discovery. The Golden Era of antibiotics leveraged naturally abundant small molecules that were readily identified using traditional methods; since the end of the era, this methodology has proven to be much more difficult to be further mined for new antibiotics. The next phase of antibiotic drug discovery could potentially leverage the wealth of proteomics, genomics, and metabolomics data to design small molecule agents that are potent and of high specificity. To accomplish this, the field fundamentally needs guiding principles describing the molecular determinants of small molecule permeation into Gram-negative bacterial cells akin to the Lipinski’s rules of 5 (Ro5).

Gram-negative bacterial pathogens pose a significant challenge to antibiotic development due to their distinctive cell wall structure, particularly the outer membrane (OM), which restricts the penetration of small molecules into the periplasmic space (and then further into other subcellular compartments). It is currently understood that small molecules capable of traversing the OM either possess specific physicochemical properties that enable crossing of this barrier by passive diffusion or hijacking porins or other membrane embedded proteins (**Fig. 1a**). Typically, minimum inhibitory concentration (MIC) analyses have been used as a proxy for drug accumulation. However, this approach is problematic because MIC reflects antimicrobial activity, which is more indicative of target engagement, rather than directly measuring intracellular concentrations. It is therefore crucial to develop robust and widely adopted live cell assays that can independently measure the accumulation of small molecules across the OM, beyond just relying on MIC values. Developing such methods will provide more accurate insights into the accumulation behavior of potential molecules and ultimately aid in improving antibacterial drug development.^13^

**Figure 1.**
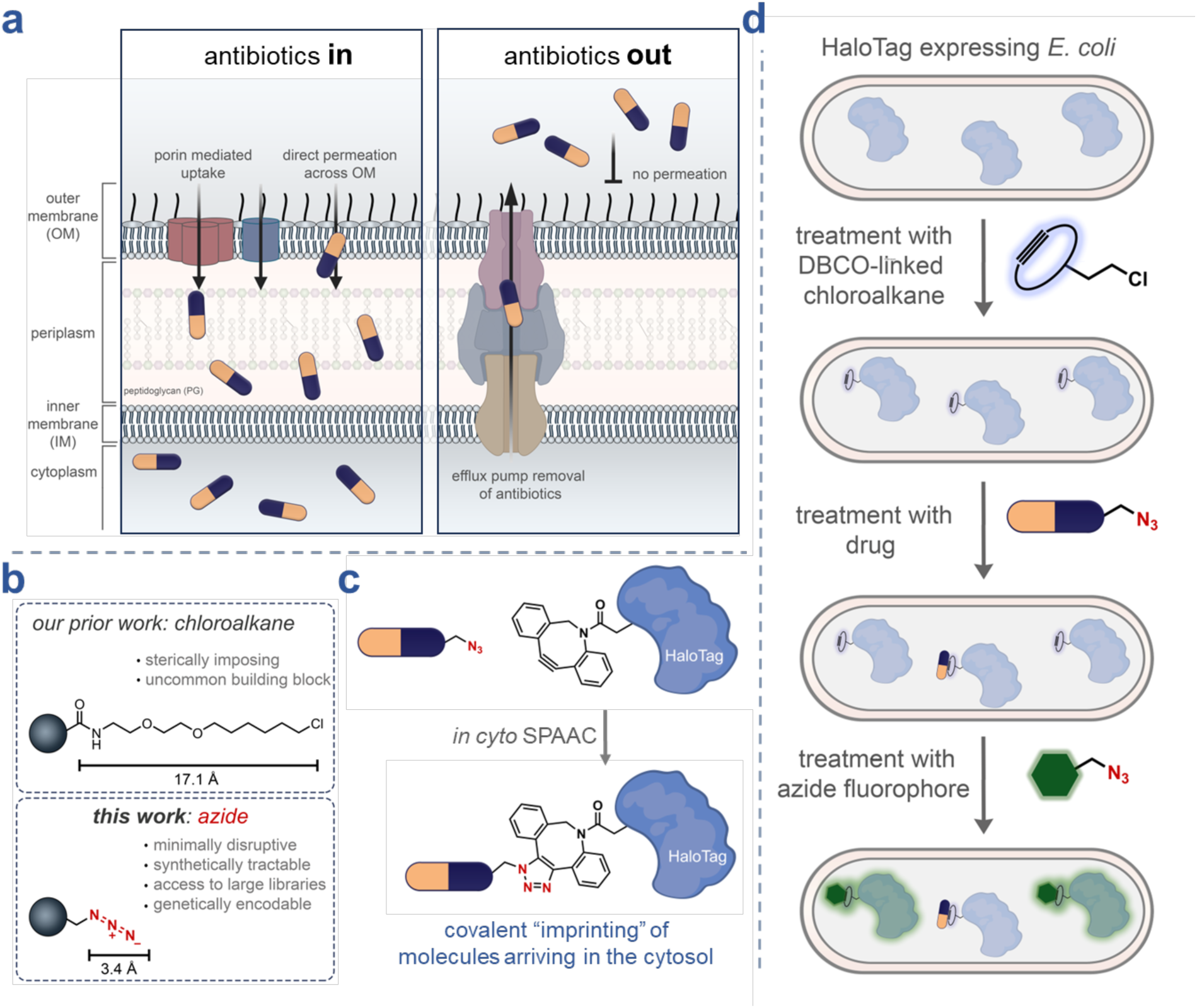
**a**, Schematic representation of the primary factors that can influence the level of molecules inside Gram-negative bacteria, including *E. coli*. There are several modes of permeation past the bilayers that surround Gram-negative bacteria, and these routes can differ based on the composition of the small molecule. **b**, A direct comparison in molecular size between the chloroalkane tag that is compatible with HaloTag and the smaller azide tag. **c**, Once compounds arrive in the cytosol of *E. coli*, they will be positioned to engage with the strained alkyne landmark that is found in the cytosol *via* a SPAAC reaction. This covalent register of arrival marks the subcellular localization of the drug within the cell. **d**, Overall workflow of CHAMP in *E. coli*. First, bacteria carrying a plasmid that expresses HaloTag in the cytosol are treated with a chloroalkane-modified strained alkyne that effectively converts the reactivity of HaloTag from chloroalkane to azide. The exposure of live cells to compounds of varying composition, which are azide tagged, (‘pulse step’) can be used to assess the level of cytosolic arrival. In the case of a molecule that reaches high levels of cytosolic arrival, there will be less strained alkyne to react with the subsequent exposure to a fluorescent azide (‘chase step’). In the case of molecules with high levels of accumulation, the cellular fluorescence level is expected to be low. The opposite should be observed for molecules with low levels of cytosolic accumulation.

The field has consistently regarded liquid chromatography-tandem mass spectrometry (LC-MS/MS) as a premier technique for quantifying drug accumulation.^14–17^ Indeed, devoid of the need for a chemical tag, this method presents significant advantages for measuring the uptake of molecules in bacteria. However, mass spectrometry, while advantageous for analyzing whole cell association, has principal limitations as currently reported: (a) restricted throughput capacity and (b) would ambiguously define molecule location unless additional, careful fractionation methods are included to show periplasmic or cytoplasmic accumulation, ^18^ and (c) needs compounds to be mass active. This lack of subcellular localization is also a limitation of spectroscopic measurements, such as those using fluorescent tags or inherent fluorescence. Despite these challenges, recent *tour de force* efforts by the Hergenrother laboratory^19,20^ have successfully employed LC-MS/MS to develop the eNTRy rules, highlighting the significance of primary amines in the accumulation of small molecules in *E. coli*. Yet, this level of throughput has been difficult to achieve by others in the field. Consequently, limited throughput in analyzing the cytoplasmic accumulation of molecules remains a significant challenge. Without a clear understanding of the exact parameters that drive the accumulation of molecules into the cytosol of *E. coli*, efforts to broadly test and prioritize drug leads will be hindered.

In recent years, the Kritzer lab introduced a groundbreaking method for quantifying cytosolic accumulation of molecules in mammalian cells. This method, called the ChloroAlkane Penetration Assay (CAPA), involves applying a chloroalkane-tagged ‘target molecule’ to HaloTag-expressing mammalian cells^21–22^ The level of cytosolic accumulation is then inferred from the fluorescence signal produced by a chloroalkane-fluorophore ‘chase molecule’. HaloTag, which can be expressed as freely diffusing proteins in the cytoplasm, selectively reacts with chloroalkanes to form a stable covalent bond. Critically, the covalent and subcellular nature of the assay confirms the cytosolic arrival of molecules, rather than whole-cell association.^23–28^ For these reasons, this method is now widely regarded as the gold standard for measuring cytosolic accumulation in mammalian systems. Building on the success of CAPA in mammalian systems, we recently adapted this approach to measure cytosolic accumulation in diderm bacteria, such as *E. coli*, *Mycobacterium smegmatis, and Mycobacterium tuberculosis*, in a process we term ‘Bacterial CAPA’ (BaCAPA).^29^ To address (1) potential interference induced by the 15-atom chloroalkane tag in bacterial accumulation studies (**Fig. 1b**) and (2) limitations in the availability of chloroalkane-tagged test molecules, we developed an enhanced method in *E. coli* that substitutes the chloroalkane with a shorter 3-atom azide tag. The choice of the azide tag was based on its minimal size, relatively low impact on the physicochemical properties of the parent compound, and stability in biological systems.^30,31^ Moreover, there are well-established protocols for late-stage azide installation into complex molecules^32,33^ and in synthetic peptides, as well as metabolic incorporation within biomacromolecules, such as RNA, oligosaccharides, and proteins.^34–37^

Our team recently introduced the Peptidoglycan Accessibility Click-Mediated AssessmeNt (PAC-MAN) assay in mycobacteria.^38^ In PAC-MAN, a strained alkyne, dibenzocyclooctyne (DBCO), is metabolically incorporated^39,40^ into the peptidoglycan layer of diderm bacteria to monitor the passage of molecules through the OM. In this work, we adapted our previous workflow by treating HaloTag-expressing *E. coli* cells with a chloroalkane-DBCO reagent that anchors a strained alkyne to cytosolic HaloTag. Azide-tagged test compounds that reach the bacterial cytosol are then covalently imprinted to HaloTag *via* strain-promoted azide-alkyne cycloaddition (SPAAC)^41,42^ reactions (**Fig. 1c**). This step is followed by treatment with an azide-tagged fluorophore, after which cellular fluorescence is measured using a flow cytometer or a plate reader in 96- or 384-well plates. Fluorescence levels are inversely proportional to the extent of SPAAC reactions between DBCO and the test azides in the cytosol, providing a straightforward method to determine apparent cytoplasmic arrival (**Fig. 1d**). The irreversible covalent reaction between chloroalkane-DBCO and HaloTag anchors DBCO in the cytosol and offers a method for determining subcellular localization. This adaptation, termed the ‘CHloroalkane Azide Membrane Penetration’ (CHAMP) assay, focuses on measuring cytosolic accumulation rather than whole-cell association. It operates at a throughput level that, to our knowledge, has not been previously achieved on such a scale. We demonstrated the capability to make more than 5000 measurements of approximately 1500 unique molecules in various biological contexts.

We first optimized the assay parameters to maximize the dynamic range of the fluorescence signals and adapted the assay for compatibility with high throughput screening platforms. A small panel of azide-tagged molecules was used to benchmark the testing phase, followed by testing a series of antibiotics with varying levels of biological activity against *E. coli*. Next, this approach was validated by evaluating the impact of OM permeabilization in *E. coli* on the accumulation of azide-modified drugs and analyzing the reactivity of these azides with DBCO-modified beads. Furthermore, the assay was adapted for high-throughput screening platforms, with the datasets coupled with structural modifications that were previously proposed to impact accumulation in *E. coli*. We analyzed a highly diverse set of azide-tagged molecules, including a 404-member and a 1152-member library, to evaluate their cytosolic uptake in *E. coli* under four unique biological contexts: wildtype, *tolC*-null *E. coli* (the major efflux pump gene), porin overexpression (highly permeable/hyperporinated), and OM permeabilization induced by antibiotic treatment. These thousands of unique measurements, all acquired in under a month, were analyzed to deliver a first-in-class profiling platform in *E. coli*.

### Optimization of HaloTag Expression in *E. coli*

Optimizing key conditions in the CHAMP assay is essential before using it as a method for assessing molecule accumulation in bacterial cells. These conditions include HaloTag expression, anchoring of DBCO-chloroalkane onto HaloTag, choice of fluorophore, and conditions for measuring the accumulation of the test molecules. First, *E. coli* cells were transformed with a plasmid encoding HaloTag (see SI for details). The expression of functional HaloTag was confirmed by treating cells with a chloroalkane-linked rhodamine 110 (R110cl) and subsequently conducting flow cytometry analysis (**Fig. 2a**). Given the nature of the dye, it was expected to have a high level of cellular accumulation. The observed cellular fluorescence intensity correlated well with the concentrations of IsoPropyl β-D-1-ThioGalactopyranoside (IPTG) used during the protein expression phase (**Fig. 2b**). Of significance, the cellular fluorescence intensities initiated at a low value in the absence of IPTG, suggesting that the expression of HaloTag was responsible for capturing the chloroalkane-linked fluorophore in the cytosolic space of *E. coli*. The signal intensities exhibited exponential growth with increasing IPTG concentration before reaching a saturation plateau.

**Figure 2.**
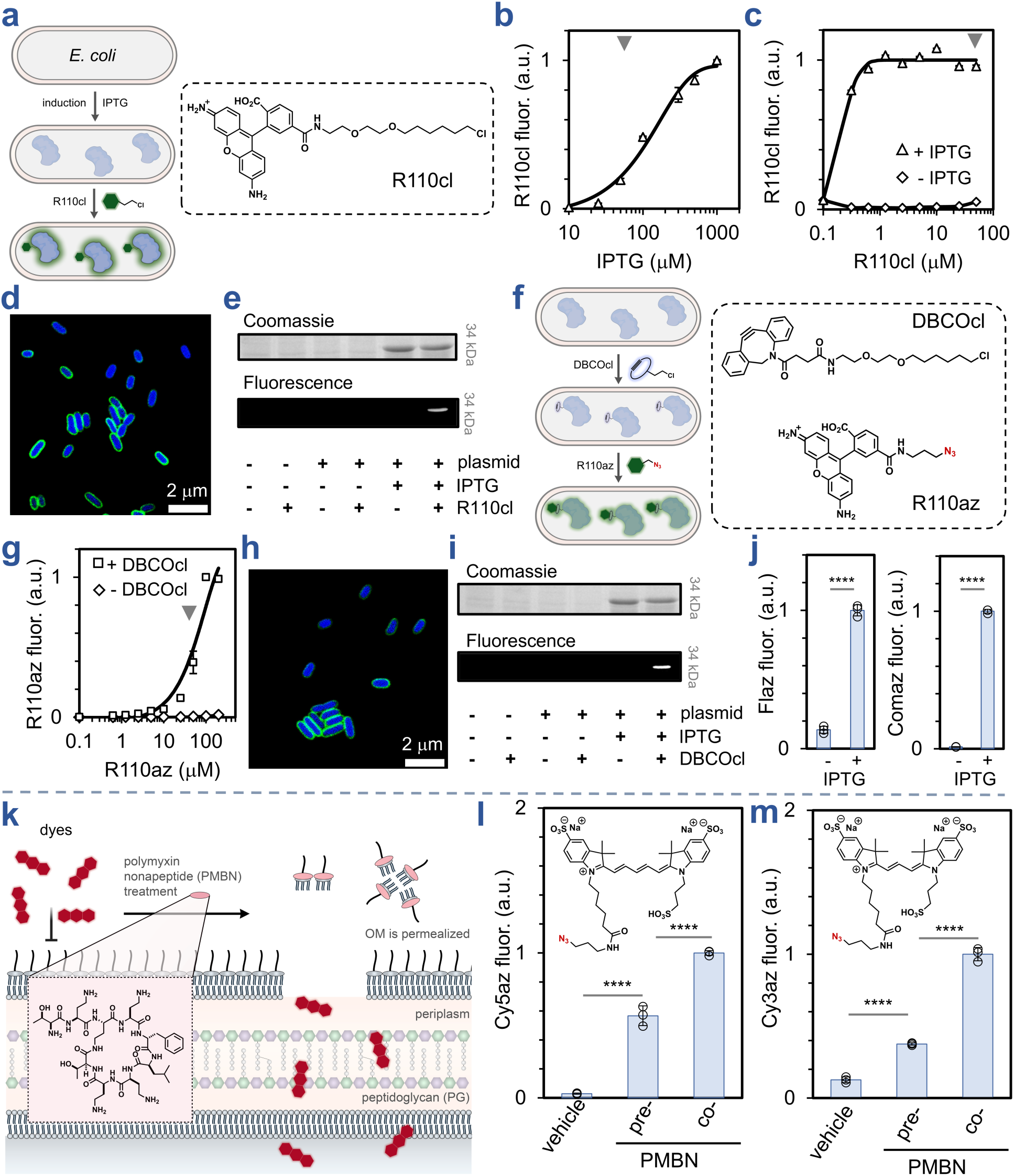
**a**, Scheme of the analysis of HaloTag expression in *E. coli*. HaloTag-expressing cells were incubated with R110cl, which should react with HaloTag in the cytosol. The chemical structure of R110cl is shown on the right. **b**, Cellular assay with *E. coli* carrying the HaloTag-expressing plasmid titrated with varying levels of IPTG. **c**, Cellular assay with HaloTag-expressing *E. coli* titrated with the ligand R110cl. **d**, Fluorescence confocal images of D-Lys(FITC) labeled *E. coli* expressing HaloTag and treated with Comcl. **e**, SDS-PAGE analysis (Coomassie and fluorescence) of *E. coli* carrying the HaloTag-expressing plasmid or the empty vector in the presence/absence of either IPTG or R110cl. **f**, Scheme of the analysis of HaloTag expression in *E. coli* with the CHAMP format. HaloTag-expressing cells were incubated with DBCOcl, which should react with HaloTag in the cytosol, followed by treatment with R110az. The chemical structures of DBCOcl and R110az are shown on the right. **g**, Cellular assay with HaloTag-expressing *E. coli* after treatment with DBCOcl then titrated the fluorophore R110az. **h**, Fluorescence confocal images of D-Lys(FITC) labeled *E. coli* expressing HaloTag, treated with DBCOcl, then reacted with Comaz. **i**, SDS-PAGE analysis (Coomassie and fluorescence) of *E. coli* carrying the HaloTag-expressing plasmid in the presence/absence of: IPTG, DBCOcl and treated with R110az. **j**, Cellular assay with HaloTag-expressing *E. coli* after treatment with DBCOcl then treatment with the fluorophore Flaz (left) and Comaz (right). **k**, Schematic representation of how PMBN can permeabilize the OM of *E. coli* and facilitate greater apparent accumulation of less permeable molecules. **l**, Cellular assay with HaloTag-expressing *E. coli* after treatment with DBCOcl then treated with the fluorophore Cy5az. Cells were treated with vehicle, pre-incubated with PMBN, or co-incubated with PMBN. **m**, Cellular assay with HaloTag-expressing *E. coli* after treatment with DBCOcl then treated with the fluorophore Cy3az. Cells were treated with vehicle, pre-incubated with PMBN, or co-incubated with PMBN. Data are represented as mean +/- SD (n = 3) of technical replicates. *P*-values were determined by a two-tailed *t*-test (* denotes a *p*-value < 0.05, ** < 0.01, ***<0.001, ns = not significant).

Additionally, with the goal of optimizing the concentration of the anchoring chloroalkane ligands in the CHAMP assay, cells expressing high levels of HaloTag were subjected to varying concentrations of R110cl. Cellular fluorescence intensities were low in the absence of R110cl, while they experienced significant enhancement at 0.3 μM of R110cl, followed by a dose-dependent increase that plateaued at approximately 3 μM (**Fig. 2c**). The localization of the fluorescence signals within the cells were additionally assessed using confocal imaging (**Fig. 2d** and **Fig. S1**). To confirm that the accumulation measurements in this study reflect cytoplasmic localization of molecules, the cytoplasmic compartment of *E. coli* cells was delineated by labeling the cell wall with D-Lys(FITC)-OH, which labels the peptidoglycan layer.^43^ Consequently, the cytosolic probe for confocal imaging was changed to coumarin chloroalkane (Comcl), which fluoresces in a distinct spectral window, thereby enabling clear spatial separation of cytosolic and periplasmic signals. Moreover, fluorescence imaging of an SDS-PAGE gel containing cellular components from induced cells revealed a fluorescent band at approximately 30 kDa, consistent with the expected molecular weight of HaloTag (**Fig. 2e**). Together, these results confirm the capability of HaloTag to selectively anchor molecules tagged with a chloroalkane linker in *E. coli*.

### DBCO-chloroalkane anchoring within HaloTag

To mitigate the potential impact of a longer chloroalkane linker, we employed an azide tag that could then undergo an *in cyto* SPAAC reaction^30,31^ with DBCO. To accomplish this, we envisioned that we could anchor a DBCO epitope within HaloTag by pre-treating cells with DBCOcl (**Fig. 2f**). HaloTag-bound DBCO can then be leveraged to assess the accumulation of azide-bearing target molecules in the cytosol of *E. coli*. Several experiments were conducted to refine the procedure. First, *E. coli* cells expressing HaloTag were incubated with DBCOcl to install the DBCO reporter within the cytoplasm. Empirical assessments were conducted to determine the optimal concentration for DBCOcl anchoring, employing an approach reminiscent of BaCAPA experiments. Cells were incubated with varying concentrations of DBCOcl followed by the treatment with R110cl (**Fig. S2**). Our findings indicated that the maximum levels of DBCO installation were achieved at approximately 50 μM of DBCOcl, exhibiting an EC_50_ value of ∼18.5 μM.

Subsequently, we conducted a baseline CHAMP experiment by treating *E. coli* cells with the azide-tagged rhodamine 110 (R110az) to reveal the anchored DBCO handles within individual cells. A titration of R110az revealed a concentration-dependent increase in fluorescence intensities, establishing the optimal concentration of R110az for subsequent experiments (**Fig. 2g**). Satisfyingly, a large dynamic range in fluorescence of up to 30-fold was observed in cells treated with both DBCOcl and R110az compared to controls. In the absence of DBCOcl treatment, cellular fluorescence levels were similar to those of untreated cells, indicating that R110az has minimal retention in bacterial cells without a covalent SPAAC reaction. Cellular fluorescence levels were also near background levels in non-induced cells that were also treated with DBCOcl and R110az, suggesting negligible retention of DBCOcl or cytosolic DBCO modified with R110az (**Fig. S3**).

Confocal microscopy confirmed that the fluorescence signal was distributed throughout the cytoplasm (**Fig. 2h** and **Fig. S4**) and fluorescent gel imaging also showed a band matching in the expected molecular weight range of HaloTag (**Fig. 2i**). These results are consistent with a site selective anchoring of the DBCO and subsequent SPAAC with an azido-fluorophore at the site of HaloTag. When HaloTag was expressed in a smooth *E. coli* strain (ATCC 25922 with intact lipopolysaccharides on its outer surface) there was a similar dynamic range (**Fig. S5**). These data highlight the potential ability to translate CHAMP to a system with more complex surface composition and suggest that CHAMP can be applied more broadly across most types of bacteria that can be genetically manipulated. In the case of smooth *E. coli*, it is important to consider how the system can work in this context because surface biomacromolecules (including *O*-antigen) can alter accumulation profiles due to the potential impact of *O*-antigens in steric shielding of porins.^44,45^ To verify the viability of the *E. coli* cells after inducing HaloTag expression and subjecting cells to DBCOcl treatment, we conducted a SYTOX Green analysis.^46^ In live prokaryotic cells, SYTOX Green is excluded from cells with intact permeability barriers.^47^ Our results revealed that the expression of HaloTag and incubation of these cells with DBCOcl and its occupancy by HaloTag in *E. coli* did not compromise the integrity of the cytoplasmic membrane (**Fig. S6**). This finding emphasizes the suitability of utilizing CHAMP for investigating molecular permeation to the cytosol without causing disruption to the cell’s permeability barriers.

We next sought to assess the compatibility of CHAMP with other azide-bearing fluorophores. Specifically, two more azide-bearing fluorophores, fluorescein (Flaz) and coumarin (Comaz), were tested. Incubation of cells after DBCOcl treatment with either fluorophore resulted in a significant increase in the cellular fluorescence (**Fig. 2j**). Cells incubated with Flaz exhibited a considerably lower fluorescence intensity compared to R110az-treated cells. Nevertheless, those results confirm the versatility of the assay for using various azide-bearing fluorophores. Recognizing that the assay is effectively completed at the end of the incubation with the test molecule, we proposed that higher signal-to-noise ratios could be achieved by removing the permeation barrier to the azido-fluorophores. Notably, formaldehyde fixation has been previously shown to outperform alcohol-based fixation in terms of increasing permeability in *E. coli*.^48^ Our results indicated that fixation performed prior to the incubation with fluorophores led to an overall increase in fluorescence levels for all three tested molecules (**Fig. S7**). This observation suggests that the dyes themselves could face significant permeation challenges imposed by the intact envelope of *E. coli*. Among the three dyes, the most favorable signal-to-noise ratio in fixed cells was achieved with Flaz treatment, resulting in a substantial increase in cellular fluorescence with minimum background staining.

### Influence of OM permeabilization on accumulation of Cy3az and Cy5az

We then applied CHAMP to analyze the impact of OM permeabilizers. Of significance, the permeation of small molecules in Gram-negative bacteria is purported to be primarily impeded by the OM.^49–52^ This formidable barrier has been a focal point of previous investigations aimed to identify compounds capable of disrupting the OM integrity. These types of molecules can serve as antibiotic adjuvants to enhance the effectiveness of antibiotics or circumvent resistance mechanisms through co-treatment. A prominent example in this category is polymyxin B nonapeptide (PMBN),^53^ a nonapeptide fragment derived from polymyxin B (**Fig. 2k**). PMBN is of particular interest due to its ability to permeabilize the OM at low concentrations. In our hands, PMBN displayed no significant impact on the colony forming unit (CFU) count when administered at concentrations of 5 μM and 10 μM, consistent with previous studies demonstrating its low toxicity at these concentrations (**Fig. S8**).^54^

To analyze the impact of OM permeabilization by PMBN on the accumulation of molecules, we selected the sulfonated cyanine series as model compounds. These dyes contain an azide group, specifically Cy5az and Cy3az, whose permeabilities are limited by their sulfonate groups (**Fig. 2l-m**). We hypothesized that OM permeabilization by PMBN would enhance the accumulation of the dyes, thereby increasing cellular fluorescence. Briefly, live *E. coli* cells were either pre-exposed to low concentrations of PMBN or co-incubated with PMBN and dye to assess the accumulation of Cy5az and Cy3az. Our data revealed a striking 19-fold increase in cellular fluorescence when PMBN was pre-incubated with the cells, and an even more substantial 34-fold increase when the cells were co-incubated with PMBN and Cy5az (**Fig. 2l**). A full range of permeabilization by PMBN was observed by 2.5 μM with an EC_50_ of 0.8 μM (**Fig. S9**). This pattern was mirrored with the analogous but smaller fluorophore, Cy3az, signifying that OM permeabilization assisted the accumulation of slightly less hydrophobic dyes as well (**Fig. 2m**). Given the substantial dynamic range that was observed, we propose that this assay could also be adapted for detection using a microplate reader for high throughput screening. Indeed, our results showed that the dynamic range was mostly retained in this format (**Fig. S10**), which should provide a more direct path to screening campaigns with diverse molecular libraries for OM permeabilizers.

### Accumulation of test small molecules and antibiotics

We set out to establish the feasibility of CHAMP in monitoring the accumulation of azide-tagged molecules into *E. coli* cells. For this initial demonstration, *E. coli* cells with DBCOcl anchored on expressed HaloTag were titrated with a test azide-bearing molecule (**Fig. 3a**), followed by treatment with R110az and cellular fluorescence levels were measured *via* flow cytometry. A concentration-dependent reduction in cellular fluorescence was observed, revealing an EC_50_ of ∼14 μM. The fluorescence intensity reached a maximum value at approximately 50 μM of the test molecule. These findings confirm the capability of the optimized CHAMP assay for investigating the accumulation of molecules in *E. coli*. With these results in hand, we shifted our focus to testing a series of azide-tagged small molecules featuring diverse structural motifs, anticipating that these variations would influence their accumulation. In the first subset, **1-3**, we tested the potential impact of amidation of carboxylic acids (**Fig. 3b**). Indeed, we observed that carboxamide, **1p**, had a higher level of apparent accumulation relative to its carboxylic acid counterpart **1n** (**Fig. 3c**). Appreciating the potential for an internal hydrogen bond due to the amino group in the β-position of *C*-terminal 2,4-diaminobutyric acid and the potentially confounding behavior of **1p**/**1n**, we extended the amino group further from the β-carbon in **2n**/**2p**. The apparent accumulation level of **2p** was significantly more pronounced, which may reflect the greater contribution of the positively charged lysine side chain, as it cannot readily form an internal hydrogen bond with the *C*-terminal group. While compounds **1n**/**1p**/**2n**/**2p** are theoretically expected to be primarily positively charged or net neutral under physiological pH, we sought to test if the pattern would hold true for a similar compound lacking a basic amino group. Our results showed that carboxamide in **3p** still led to higher levels of cytosolic accumulation in *E. coli* relative to the net negative **3n**. This small panel of compounds demonstrates how specific structural variations can readily tested for their potential ability to drive accumulation in *E. coli*, which could be crucial in designing prodrugs that mask carboxylic acids to enhance their permeability in similar pathogenic bacteria.

**Figure 3.**
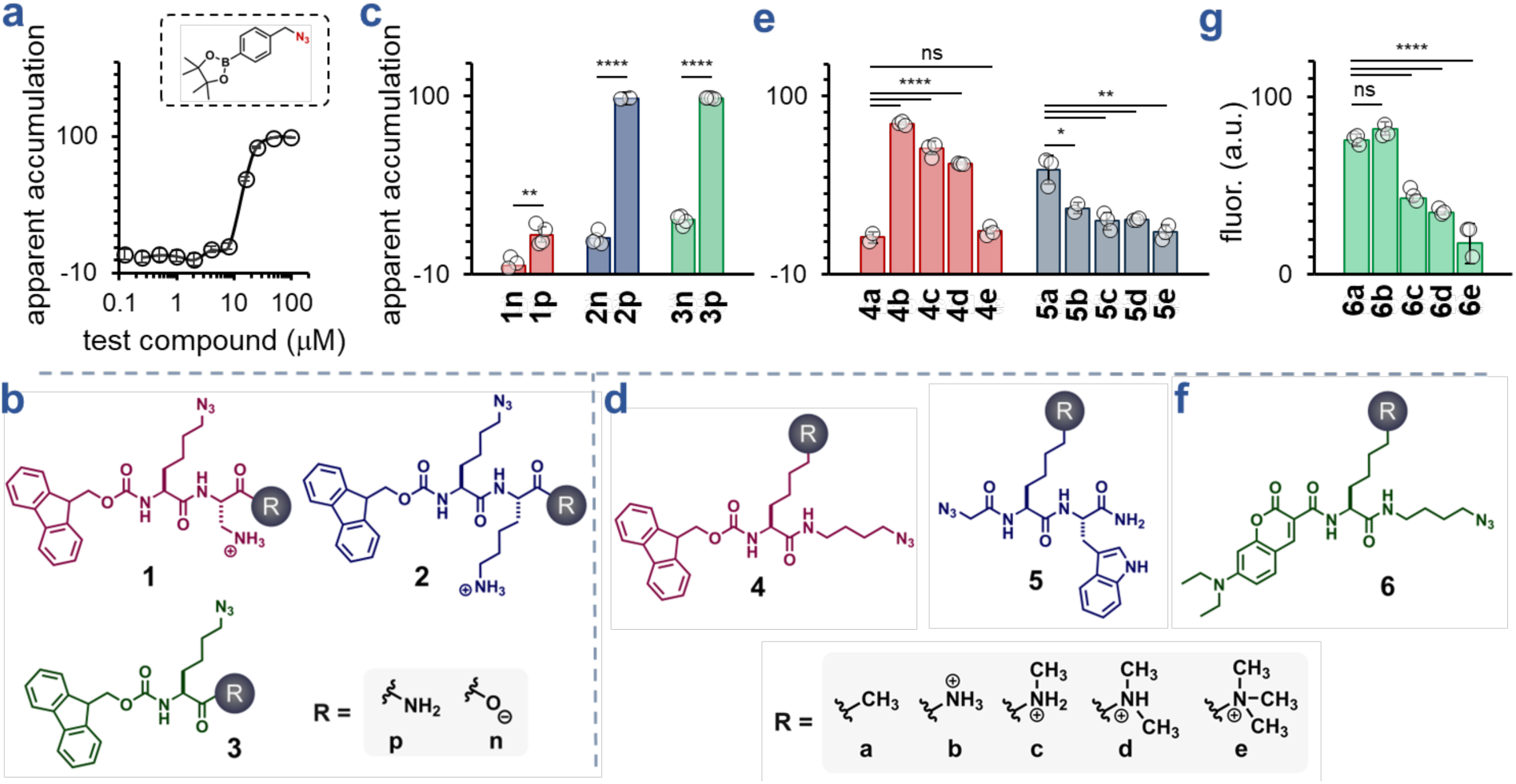
**a**, CHAMP analysis of *E. coli* titrated with an azide-tagged test molecule; the graph was obtained by plotting (1 – fold change, FC) against concentration, such that a higher (1 – FC) value corresponds to greater molecular accumulation **b**, Chemical structures of the **1**-**3** series. **c**, CHAMP analysis of *E. coli* with 50 μM of **1n**, **1p**, **2n**, **2p**, **3n**, and **3p**. For all CHAMP assays, *E. coli* cells carrying the HaloTag-expressing plasmid were induced with IPTG, treated with DBCOcl, then incubated with the azide-tagged test molecules for 1 h in PBS at 37 °C. **d**, Chemical structures of the **4**-**5** series. **e**, CHAMP analysis of *E. coli* with 50 μM of series **4** and **5**. **f**, Chemical structures of the series **6**. **g**, Direct cellular fluorescence assay with HaloTag-expressing *E. coli* after treatment with DBCOcl then 50 μM of series **6**. Data are represented as mean +/- SD (n = 3) of technical replicates. *P*-values were determined by a two-tailed *t*-test (* denotes a *p*-value < 0.05, ** < 0.01, ***<0.001, ns = not significant).

The Hergenrother group has used an established method in the field, mass spectrometry, to empirically demonstrate that primary amines can serve as privileged functional groups to improve accumulation in *E. coli* as part of the eNTRy rules.^17,19,20^ To benchmark *E. coli* CHAMP with these structural parameters, we made three additional compound panels, **4**-**6** (**Fig. 3d**). Series **4** has a similar general scaffold to our established molecules (**3**) that exhibited high levels of accumulation, while the molecules (**4a**-**4e**) feature five different configurations of the side chains ranging from an unfunctionalized terminal hydrocarbon to amino groups that vary in their degrees of methylation. Within this series, it was observed that the substitution of a methyl group in **4a** for a primary amino group in **4b** resulted in a large increase in apparent accumulation (**Fig. 3e**). The alkylation of the amino group progressively reduced accumulation levels in **4b**-**4e**. These results show marked agreement with the eNTRy rules. In fact, the trimethyl **4e** had a similar accumulation profile as the compound **4a** that does not have an amino group. To test the generalizability, a subsequent series, **5**, was synthesized that retained much of the same features to test the generalizability. Interestingly, the hydrocarbon terminated **5a** displayed higher apparent accumulation than any other analogs with an amino group, regardless of methylation states.

Next, we pivoted to benchmark the assay using an alternative strategy. Instead of using a pulse-chase modality to measure apparent accumulation, the test compounds in **6** all have a small fluorophore incorporated as part of the molecule (**Fig. 3f**). The goal was to determine whether the pulse-chase method produces results consistent with those obtained from a single-step direct fluorescence analysis. Similar to the **4** and **5** series, the methylation of the amino groups in series **6** led to lower accumulation levels (**Fig. 3g**). The absence of DBCOcl that installed the landmark and IPTG led to near background levels of fluorescence, which suggests that the cellular fluorescence observed in series **6** is due to engagement with the landmark and reported on cytosolic arrival (**Fig. S11**). Within the same series, we reasoned that we could evaluate the CHAMP-based levels of accumulation using a pulse-chase workflow. Consistently, we observed that the pattern of accumulation across **6a**-**6e** was nearly identical for a direct measurement and as CHAMP-based analysis with a non-overlapping fluorophore (**Fig. S12**). Together, these findings align well with the eNTRy rules and underscore the advantages of having a robust and straightforward method for measuring cytosolic accumulation. Such a method can significantly enhance the throughput of identifying structural determinants of accumulation.

We posited that CHAMP would serve as a versatile platform for unraveling the molecular determinants of accumulation related to elements found within the OM, which can include both the membrane bilayer and the proteins embedded within this barrier. To achieve this, we sought to compare the apparent accumulation of azide-tagged antibiotics (**Fig. 4a**). Our first goal was to evaluate the accumulation of azide-tagged antibiotics across biological contexts related to efflux pumps (**Fig. 4b**).^55^ Efflux pumps can greatly reduce the concentrations of antibiotics by excreting them to the extracellular space. Of the 36 known or putative efflux pumps in *E. coli*,^56^ AcrAB-TolC is the best characterized efflux pump and has been described as the most prevalent in decreasing susceptibility to antibiotics.^57,58^ Genetic deletion of *tolC* is sufficient to disrupt the tripartite formation of the AcrAB-TolC complex.^59,60^ To test CHAMP in this context, we adopted an antibiotic that has been previously described as a canonical substrate of AcrAB-TolC. Novobiocin belongs to the class of aminocoumarin antibiotics and it inhibits bacterial DNA gyrase.^61^ Given its cytosolic target, novobiocin needs to permeate past the OM in order to reach the target gyrase. Deletion and mutations to *tolC* can dramatically alter the MIC of novobiocin, which has been ascribed to its substrate recognition by this powerful efflux pump.^62^

**Figure 4.**
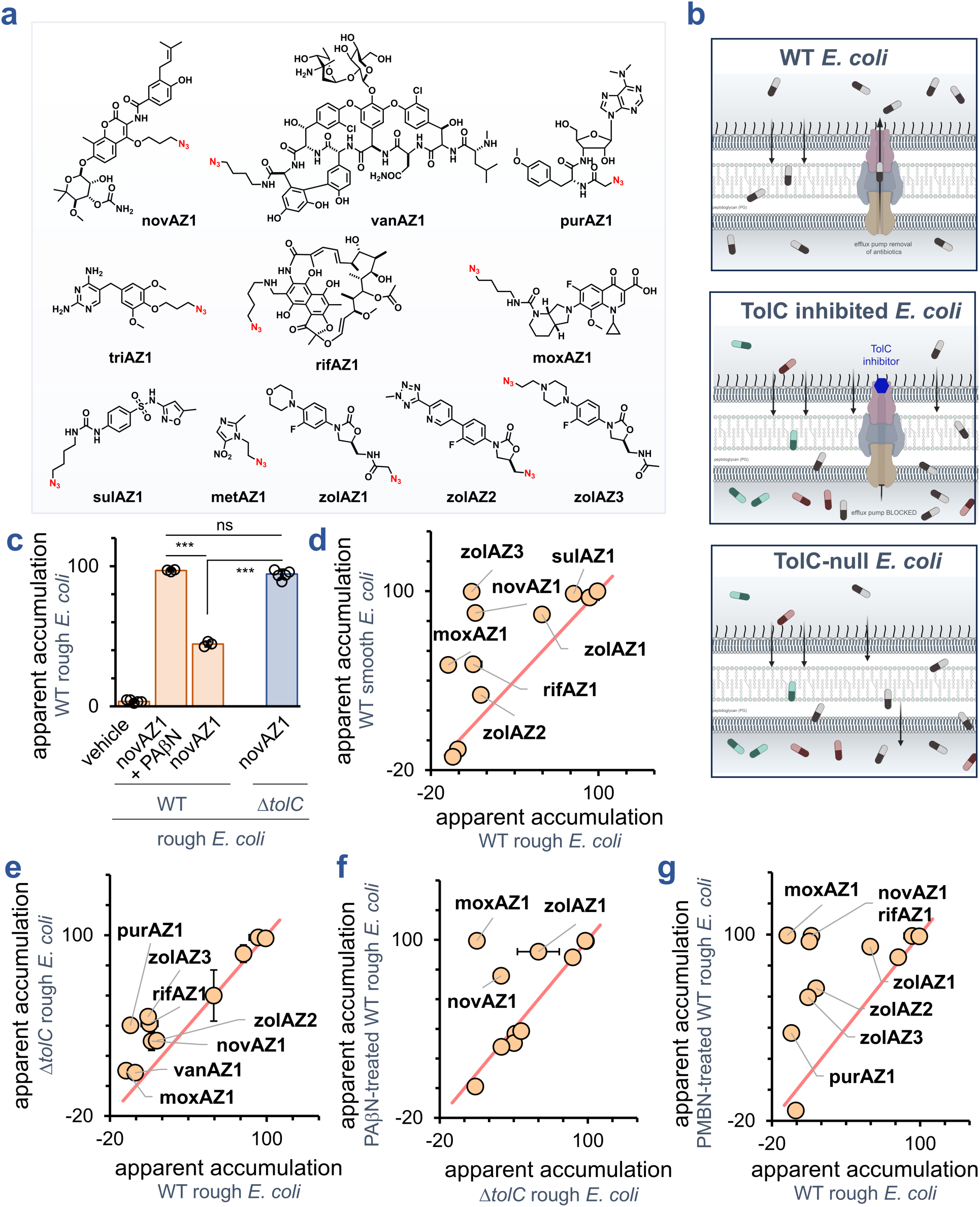
**a**, Chemical structure of the 11 azide-tagged antibiotics. **b**, Schematic representation of the three biological contexts that were tested with the modified antibiotics. This includes WT *E. coli*, D*tolC E. coli*, and WT *E. coli* treated with the small molecule TolC inhibitor PAβN. **c**, CHAMP analysis of *E. coli* with 50 μM of novAZ1 in WT *E. coli* or D*tolC E. coli* in the presence of absence of PAβN. **d**, Comparison of apparent accumulation of 11 azide-tagged antibiotics in *E. coli* using CHAMP across two biological contexts: WT smooth *E. coli* and WT rough *E. coli*. **e**, Comparison of apparent accumulation of 11 azide-tagged antibiotics in *E. coli* using CHAMP across two biological contexts: D*tolC E. coli* and WT rough *E. coli*. **f**, Comparison of apparent accumulation of 11 azide-tagged antibiotics in *E. coli* using CHAMP across two biological contexts: PAβN co-incubated WT rough *E. coli* and D*tolC E. coli*. **g**, Comparison of apparent accumulation of 11 azide-tagged antibiotics in *E. coli* using CHAMP across two biological contexts: PMBN co-incubated WT rough *E. coli* and WT rough *E. coli*. For all CHAMP assays, *E. coli* cells carrying the HaloTag-expressing plasmid were induced with IPTG, treated with DBCOcl, then incubated with the azide-tagged test molecules for 1 h in PBS at 37 °C. Data are represented as mean +/- SD (n = 3) of technical replicates. *P*-values were determined by a two-tailed *t*-test (* denotes a *p*-value < 0.05, ** < 0.01, ***<0.001, ns = not significant).

An azide-modified novobiocin derivative was synthesized, novAZ1, that minimally altered the chemical configuration of the original antibiotic (**Fig. 4a**). In WT *E. coli*, CHAMP analysis revealed that novAZ1 had a low level of apparent accumulation relative to *tolC*-null *E. coli* (**Fig. 4c**), which is an indication that TolC could reduce the effective concentration of novAZ1 in the cytosol of WT *E. coli*. These results are consistent with novAZ1 being a substrate of TolC. A parallel strategy was also employed to test the viability of CHAMP in measuring substrate recognition in *E. coli*. It is well established that the function of TolC can be disrupted with small molecule inhibitors such as phenylalanine-arginine b-naphthylamide (PAβN).^63^ To this end, a CHAMP analysis was performed for novAZ1 with and without the co-incubation of PAβN. Our results showed that the use of a small molecule chemical inhibitor of TolC led to an elevated level of apparent accumulation of novAZ1. Collectively, these experiments validate novAZ1 as a substrate of AcrAB-TolC in the context of the CHAMP assay. In the future, we will explore how CHAMP can be paired with high-throughput screens to discover next-generation TolC inhibitors (or that of other efflux pumps).

With these results in hand, we set out to widen our library of azide-tagged antibiotics and measure their apparent accumulation in *E. coli* in four different contexts that operate at the OM. The antibiotics covered a wide range of mechanisms of action and cellular targets. The azide groups were added *via* late-stage modifications and the chemistry used varied depending on the functional groups available to perform edits to install the azide epitope. In all, 11 azide-tagged antibiotics were included in this sub panel, representing one of the most comprehensive analyses reported to date. Analysis of the accumulation profiles revealed that CHAMP was readily able to discern the relative cytosolic accumulation levels of a wide range of molecules. Of note, there was no detectable accumulation of vancomycin azide (vanAZ1) within the cytosolic space of *E. coli*, consistent with expectations for this organism. Vancomycin exerts its activity by binding to the D-Ala-D-Ala terminus of the lipid II peptidoglycan precursor, which is presented on the outer leaflet of the inner membrane.^64^ However, *E. coli*, and, more broadly, Gram-negative bacteria, are intrinsically resistant to vancomycin due to the impermeability of their outer membrane, and only become susceptible when this barrier is disrupted.^65^ Nonetheless, the CHAMP assay, as employed in this study, provides information that excludes cytoplasmic accumulation of vancomycin but remains agnostic to its potential presence in the periplasm. Several antibiotics act within the periplasmic space of *E. coli* and other Gram-negative bacteria, including β-lactam antibiotics, which target periplasmic penicillin-binding proteins (PBPs) involved in peptidoglycan synthesis, and glycopeptide antibiotics such as vancomycin, which inhibit cell wall cross-linking.^66,67^ The current inability of the CHAMP platform to discriminate periplasmic localization represents a limitation of the assay in its present form. Accordingly, vancomycin serves as an effective negative control in our CHAMP assay for assessing the intracellular accumulation of other antibiotics. Other molecules that were smaller in size (e.g., trimethoprim, triAZ1, and sulfonamides, sulAZ1) showed effective accumulation into the cytosol of *E. coli*. Among molecules with high accumulation levels were oxazolidinones. A similar profile was also observed for smooth *E. coli* that has its full set of *O*-antigen (**Fig. 4d**), which indicates that the *O*-antigen chains do not appear to significantly hinder the accumulation levels of these specific molecules.

Next, accumulation analysis *via* CHAMP was performed across the entire series of antibiotics using either the TolC-null *E. coli* strain or the small molecule AcrAB-TolC inhibitor. Interestingly, a number of antibiotic derivatives, including puromycin, showed differential accumulation in *tolC* deleted strains relative to the parental WT strain (**Fig. 4e**). For some of these types of structures, the differential accumulation in *tolC*-null strains had yet to be analyzed. Additionally, accumulation profiles for the antibiotic derivatives were also tested with WT cells treated with the small molecule inhibitor, PAβN. As expected, the profile of accumulation across the series of antibiotics was not identical (**Fig. 4f**). This difference can be attributed to many factors including the disruption to the OM in the absence of TolC and/or the polypharmacology of most small molecules including PAβN. Testing a broader range of antibiotics could start to reveal how cells respond to genetic or small molecule stressors in the context of drug accumulation in *E. coli*. These results demonstrate the potential of utilizing CHAMP to confirm inhibitors of the TolC efflux pump and, most importantly, offer a means to readily compare the apparent accumulation of antibiotic derivatives in strains with genetic alterations that can potentially modulate the accumulation of drugs.

Another aspect of the OM that CHAMP could investigate is how modifying its overall integrity affects the accumulation of modified antibiotics. This includes OM-embedded channels that facilitate passage across the OM and the chemical permeabilization of this critical bilayer. We projected that for molecules whose accumulation was impeded by the OM, there would be an increase in apparent accumulation after the passage was improved. As described before, PMBN was chosen as a chemical permeabilizer of the OM. Critically, a high number of antibiotics had a shifted apparent accumulation in *E. coli* cells upon the co-treatment with PMBN (**Fig. 4g**). These results clearly demonstrate that the treatment with OM permeabilizers can alter the level of, otherwise less permeable, molecules into *E. coli*. Alternatively, we tested a genetic strategy to increase the accumulation levels in *E. coli* in a strain with overexpression of the porin FhuA. ^68^ Our CHAMP results using the antibiotics panel clearly show that, for some antibiotics, there is a shift in their accumulation profile upon hyperporination. Interestingly, the two methods of improving passage across the OM were distinct and driven by their modes of action.

Whereas PMBN may have a broader impact, and, theoretically, also impact the cytoplasmic membrane integrity, ^69^ the substrate scope of FhuA is likely to be much narrower because it is a porin. In this work, an open and non-selective variant of FhuA that does not discriminate based on hydrophilicity, designated “pore” was used; the pore is localized to the outer membrane. The substrate size range for this strain is not clear but *E. coli* K12 BW25113 expressing this variant of FhuA was shown to be sensitive to vancomycin with a 16-fold reduction in MIC relative to the wild type strain.^70^ Nonetheless, these analyses provide extensive description of the changing landscape of accumulation in four biological contexts that can impact passage and retention within bacteria. To the best of our knowledge, this is one of the most comprehensive analyses of a series of antibiotic-derivatives within biological contexts that could impact accumulation.

### High throughput library analyses in *E. coli*

Our investigation shifted to evaluating larger sets of molecules in *E. coli*. To this end, we assembled a library of 404 commercially available azide-bearing compounds (see SI). This collection encompassed a broad spectrum of characteristics, including hydrophobicity, rigidity, hydrogen bond donors/acceptors, molecular weight, charge, and polarity. Recognizing the prevalence of azide groups in various steric and electronic configurations, we pondered whether these chemical attributes could influence their reactivity with DBCO. To gauge the reaction progress of DBCO-azide within the stipulated assay timeframe, we empirically assessed the reaction completion employing DBCO- modified beads (**Fig. S13**). A workflow akin to live cell analysis was executed, albeit devoid of the permeability barriers inherent to the cellular outer and inner membranes. Interestingly, it was discerned that not all azides exhibited satisfactory reactivity within the designated timespan of the assay, signifying that azide reactivity necessitates careful consideration (**Fig. S14**). It is noteworthy that previous investigations have also documented variations in reactivity profiles between azides and strained alkynes, including DBCO.^71^ Aromatic azides have been found to have lower reactivity in SPAAC reactions with nearly a sevenfold reduced reactivity of phenyl azide in comparison with aliphatic azides even where the aliphatic azide conformation is in the form of azidotoluene. As for alkyl azides, the following trend in reactivity has been observed: primary > secondary > tertiary. This statement is supported by prior studies.^72^ It was quickly recognized that variations in the intrinsic reactivities of azides would constrain the analysis to only those azides exhibiting comparable reactivity, thereby narrowing the chemical diversity and reducing the overall screening breadth achievable with the CHAMP platform.

In light of the challenges posed by the variable reactivities of azides from a large library that are outsourced, we embarked on an alternative approach. To mitigate the influence of azide reactivity, we employed a differential scan, examining the same set of molecules in two closely related biological contexts that exhibit phenotypic divergence. This allowed us to isolate the change in apparent accumulation and attribute it solely to the underlying phenotypic differences. We posited that this method would be apt for unraveling the molecular determinants imposed by the OM of *E. coli*. To illustrate this, our first test case was the analysis of the accumulation of a 404-molecule library in WT *E. coli* with and without the treatment with PMBN (**Fig. 5a**). As before, the apparent accumulation of small molecules, for which the OM served as the primary barrier to entry into the cytosol, is expected to change upon co-incubation with PMBN, as this barrier will be disrupted. Conversely, molecules with high propensity for accumulation in intact WT *E. coli* cells would be expected to have minimal differences in their apparent accumulation with PMBN treatment. From our analysis, there was a consistent shift in apparent accumulation across the majority of molecules tested upon the co-incubation with PMBN (**Fig. 5b**). These results demonstrate the significant barrier that the OM poses to cytosolic arrival and highlight the feasibility of conducting large-scale analyses across various biological contexts. CHAMP can effectively inform us about the differences in the behavior of molecules within these contexts. Further, the difference in the accumulation profile was analyzed across a range of physicochemical parameters including hydrogen bonding numbers, rotatable bonds, molecular weight, topological polar surface area, ClogP, and logS (**Fig. 5c**). A larger increase in apparent accumulation following PMBN treatment should theoretically indicate that a specific parameter is implicated in reducing accumulation to the cytosol. These results highlight how high-volume screening can rapidly reveal patterns of molecular features that influence cytosolic accumulation. In this key demonstration, more than 800 individual measurements were made (>400 in wild-type and >400 in PMBN-treated cells), making it the largest reported screen in *E. coli* to date.

**Figure 5.**
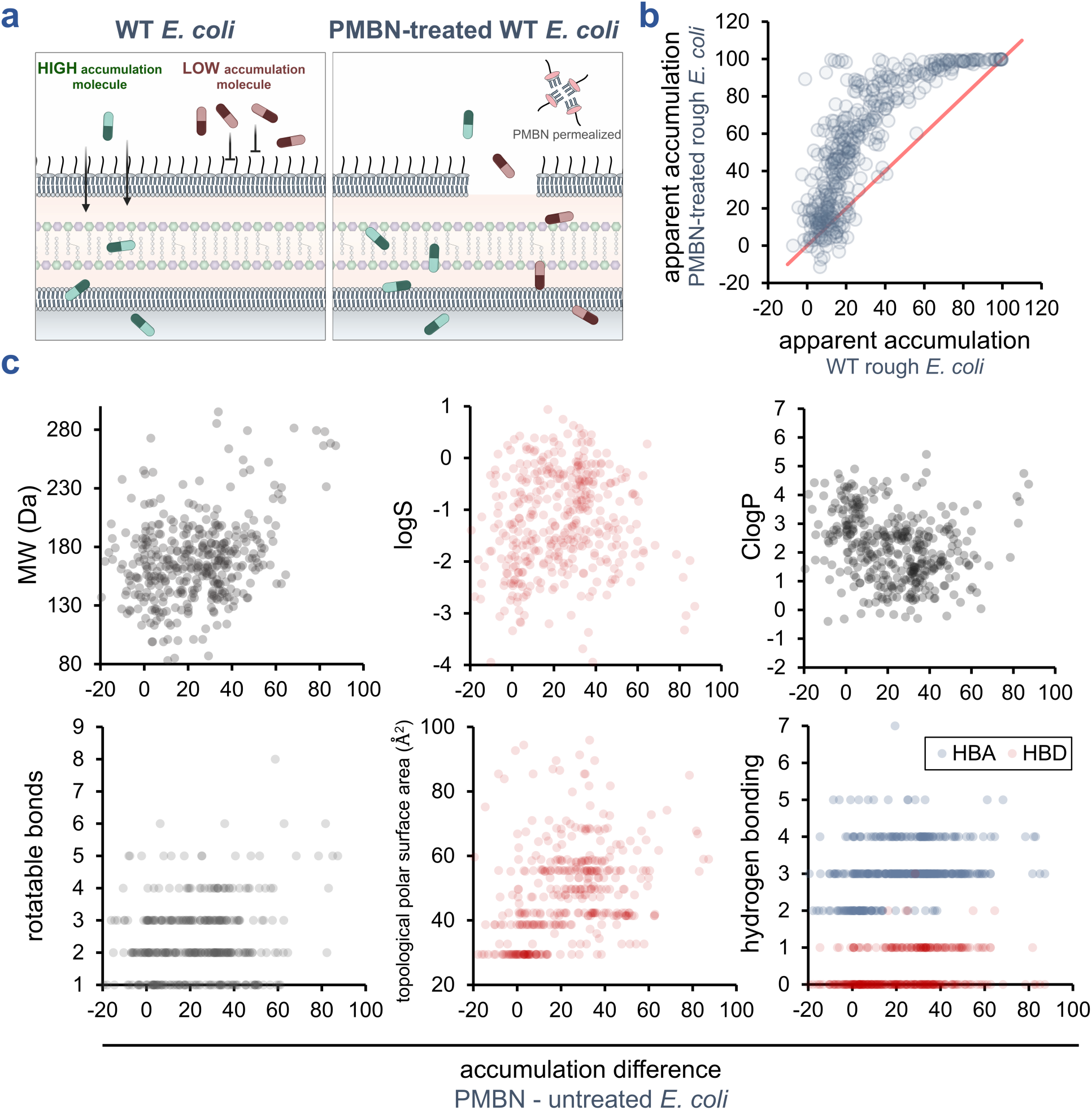
**a**, Schematic representation of the scenario of a high accumulation molecule and a low accumulation molecule in intact WT *E. coli* relative to WT *E. coli* treated with PMBN. The disruption to the membrane barriers upon PMBN treatment should primarily alter the accumulation profile of molecules whose apparent accumulations are low in intact WT *E. coli*. **b**, Comparison of apparent accumulation of 400+ small molecules in *E. coli* using CHAMP across two biological contexts: PMBN co-incubated WT rough *E. coli*and WT rough *E. coli*. **c**, Analysis of the 8 different physicochemical parameters that can potentially impact accumulation profiles of the 400+ small molecules (HBA denotes hydrogen bond acceptor; HBD denotes hydrogen bond donor). For all CHAMP assays, *E. coli* cells carrying the HaloTag-expressing plasmid were induced with IPTG, treated with DBCOcl, then incubated with the azide-tagged test molecules for 1 h in PBS at 37 °C.

As a complement to using a site-selective structural modification series (such as the amino methylation), the late stage tagging of antibiotics, and the 404 commercial library of azides, we next set out to evaluate the accumulation profile of a larger non-commercial library of small molecules. Each molecule in this library is azide-tagged, making it compatible with CHAMP ^31^ These molecules are diverse: aromatic and aliphatic, covering a wide range of chemical space and spanning a range of molecular weights. Our objective was to thoroughly examine the feasibility of a large-scale scan that has not been reported to this point. To further demonstrate the scalability of CHAMP, we screened a library comprising 1152 azide-bearing compounds that was recently described by the Dong and Sharpless groups in their strategy to install azide tags onto molecules using a diazotizing reagent.^32^ A benchmarking analysis was performed, spanning a range of cellular fluorescence levels: high, low, and intermediate using a 96-well plate format. Our experiments affirmed the remarkable reproducibility of CHAMP, underscored by a Z’ score of 0.954 (**Fig. S15**).

The entire 1152-member library was subjected to bead analysis under the same conditions as the cellular assay. Our results showed that there was a range of reactivities that were observed within the timescale of this assay, owing to the differences in the environments of the azide groups in a large library (**Fig. S16**). To start, WT *E. coli* cells were subjected to *E. coli* CHAMP analysis and the results were compared with the reactivity on the beads (**Fig. 6a**). In this analysis, there is a marked shift in the plot towards the upper left quadrant of the plot. This shift is highly indicative that their engagement with the DBCO landmark is greatly improved upon the removal of the physical barrier (e.g., OM and cytosolic membrane). Subsequently, we applied CHAMP to the complete array of the large library of molecules in four biological contexts: WT *E. coli*, WT *E. coli* pre-incubated with PMBN, hyperporinated *E. coli*, and hyperporinated D*tolC E. coli* strains. Our objective was to empirically demonstrate that CHAMP has a throughput capability that is unlike any technology to date, and, when paired with a large library, it provides a workflow to rapidly analyze cytosolic accumulation in *E. coli*. As with the smaller set of molecules, there was a general shift in apparent accumulation in WT *E. coli* upon the pre-treatment with PMBN (**Fig. 6b**). These results demonstrate that chemical disruption to the membrane barriers can broadly improve the cytosolic residency of molecules across a diverse range of physicochemical properties. Hyperporinated *E. coli* could provide a passage point to molecules, as in the case of many antibiotics. In comparison to WT *E. coli*, there was a general increase in apparent accumulation of molecules in the hyperporinated *E. coli* strain (**Fig. 6c**). Interestingly, there was general agreement on the two modalities of reduced barriers (**Fig. 6d**), yet not all compounds behaved the same way. This deviation could be, in part, due to the nature of substrate recognition by porins. Alternatively, the potential for the cytosolic membrane disruption by PMBN could reveal a slightly different set of molecules that arrive in the cytosol. With the goal of leveraging CHAMP to more broadly assess the substrate recognition by TolC, we compared the library response in hyperporinated *E. coli* to hyperporinated Δ*tolC E. coli* (**Fig. 6e**). This was an attempt to reveal compounds that are TolC substrates but have no access to the sites of recognition due to lack of accumulation. In this comparison, there was surprisingly general agreement in the behavior of the molecule in the presence and absence of TolC, which could indicate that the reduced barrier in hyperporinated cells could overwhelm the efflux capacity of TolC.

**Figure 6.**
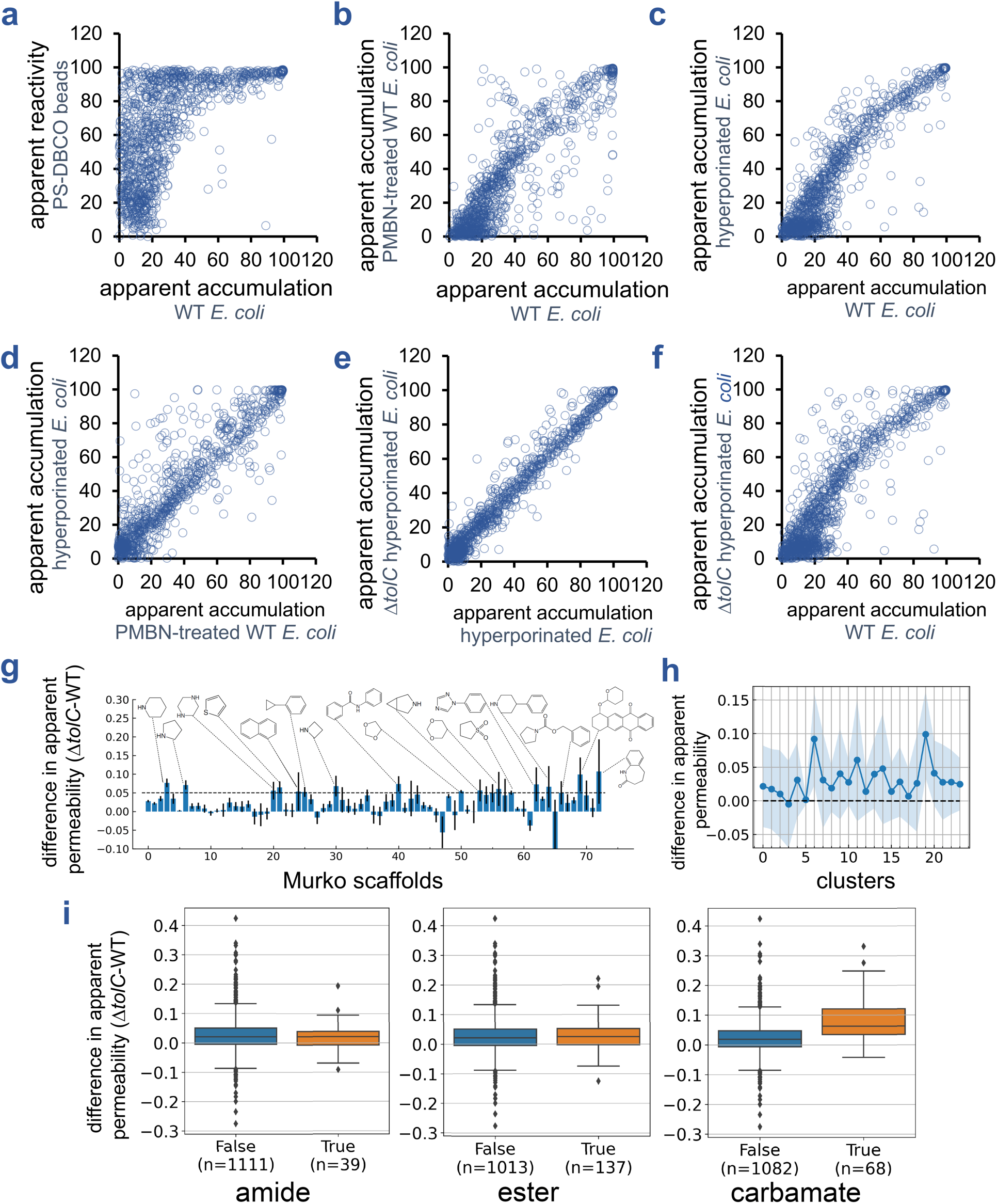
**a-f**, Comparison of apparent accumulation of 1152 small molecules in *E. coli* using CHAMP across two stated biological contexts. **g**, Average permeability shift for compounds in each Murcko scaffold. The scaffolds with large permeability shift (>0.05) are annotated with the scaffold chemical structure. **h**, Molecular clustering results in 24 clusters, the average permeability shift for each cluster is shown, shades indicate s.t.d. The clusters 6 and 19 have the lower shaded region also above 0. The representative molecules from these two clusters are annotated below. **i**, Permeability shift for compounds with or without (left) amide group, (middle) ester group, (right) carbamate group. Note that pure carbamate group or pure ester group here exclude the connected carbamate motif. For all CHAMP assays, *E. coli* cells carrying the HaloTag-expressing plasmid were induced with IPTG, treated with DBCOcl, then incubated with the azide-tagged test molecules for 1 h in PBS at 37 °C.

Using the comparison between WT and Δ*tolC* (**Fig. 6f**), we sought to broadly explore how the structure of the molecule could impact TolC recognition. We examined the correlation between compound efflux and physicochemical properties (**Fig. S17**). A few properties are shown to correlate with high statistical significance including hydrogen bond acceptor, conformational flexibility (number of rotatable bonds), amide bond, polar surface area, polarizability, saturated carbon atoms, larger molecular size, and less globularity correlate with higher chance of chemicals being recognized by TolC. There was a large range of chemical structures whose accumulation profiles were found to be especially altered by the deletion of *tolC* (**Fig. S18**). As part of the effort to understand how much reactivity weighs in on the analysis, we performed correlation analysis with different reactivity cutoff values to filter out compounds with low reactivity. Surprisingly, the reactivity cutoff from 0 to 0.4 does not have a significant influence on correlation values. We theorize that there may be two principal factors at play. First, by performing the differential scan of wild-type and *tolC* deletion mutant, the contribution from reactivity is minimized. Secondly, while the reactivity may still amplify the difference between the wild-type and the Δ*tolC* mutant, the poorer reactivity means a smaller difference in the differential scan. For this reason, such data points make smaller contributions to the final correlation value during the correlation calculation.

Next, we set out to identify motifs that correlate with compound efflux with the understanding that this would help extract substructure features that make a compound more likely to be recognized by TolC. We initially employed the Bemis-Murcko scaffold method to identify specific scaffolds whose accumulation profiles were particularly sensitive to the presence of TolC. The Bemis-Murcko scaffold is a method that treats rings and the connection between the rings (linkers) as the scaffold. The remaining fragments of the molecule are then considered as side chains. This library of 1152 compounds contains 158 unique scaffolds in total, with some of them much more enriched, e.g., the top 2 scaffolds have more than 10 compounds in each scaffold (**Fig. 6g**). We selected the first 73 scaffolds having more than 1 member in each scaffold and analyzed the average accumulation shift upon *tolC* deletion. We used a cutoff of 0.05 and depicted scaffolds where the average accumulation shift is more than 0.05. Such scaffolds include piperidine, pyrrolidine, piperazine, thiophene, naphthalene, azetidine, etc. Piperazine-based compounds have been shown to be potential modulators of efflux pump AcrAB-TolC.^73,74^ Thiophene antibacterials are known to be substrates of AcrAB-TolC in *E. coli*.^75^ Arylpiperidines have been screened as potentiators of novobiocin activity in *E. coli* cells by high-throughput assays.^76^

Though Murcko scaffolding represents the skeleton of a molecule, it is possible that side chains may also play an important role in substrate recognition by the TolC-related efflux pumps that are not captured in the scaffold-based analysis. Therefore, we also performed a chemical clustering analysis of the library. Chemical clustering is an approach that finds similarities among the molecules and groups them by similarity. This is also a common approach^77^ in drug discovery campaigns to prioritize compounds, select representative molecules, and derive smaller datasets. In our case, we grouped the molecules and investigated whether specific chemical clusters have a larger accumulation shift upon *tolC* deletion. We distributed the library into 24 clusters through hierarchical clustering. We then performed hierarchical clustering. The accumulation shift profile for molecules in each cluster showed two clusters (6 and 19) that have significantly higher apparent accumulation in the absence of TolC (**Fig. 6h**). Cluster 6 contains many compounds with the carbamate and ester functional group, while the cluster 19 contains doxorubicin and a similar derivative. We further developed a hypothesis that the carbamate motif may contribute to compound efflux, though it is not clear if the carbamate or the ester group alone is enough for that. To validate this hypothesis, we calculated accumulation shift upon *tolC* deletion for combinations of these functional groups (**Fig. 6i**). Only carbamate-based compounds were more populated within this dataset. Together, these results highlight the types of analyses that can be conducted and the queries that can be parameterized using a large volume of accumulation profiles in pathology-relevant biological contexts.

## DISCUSSION AND CONCLUSION

The use of MIC values as a proxy for molecular accumulation has long been a feature of antibiotic drug development. Drug accumulation is necessary but not sufficient for activity, making MICs an imperfect proxy for OM permeability that may generate broadly inaccurate structure-OM accumulation relationships. For example, some of the most widely used antibiotics have complex mechanisms of action that include downstream metabolic dysfunction.^78^ As well, structural changes to small molecules affect *both* their OM accumulation and interactions with target molecules. Therefore, it is extremely challenging (and potentially problematic) to attempt to decouple the inherent relationship between accumulation and toxicity solely *via* MICs. Additionally, anti-virulence agents or adjuvants, such as OM disruptors, typically lack intrinsic antimicrobial activity and are not always suitable for MIC analyses. CHAMP adds a tool to the existing methods used to measure accumulation of molecules in the cytosol.

It would be beneficial for the community to be able to query beyond the chemical space occupied by existing antibiotics. A wide range of prior efforts have focused on creative ways to measure the accumulation of small molecules, including coupling of periplasmic accumulation to b-lactamase activity,^79^ split luciferase,^14^ modification of antibiotics with fluorescent handles,^80^ LC-MS-based quantification,^15,41,42,81,82^ electrophysiology using planar lipid bilayers, ^83,84^ and microfluorospectrometry. ^85–87^ Our laboratories have also recently described a luciferase-based assay using D-cysteine tagged antibiotics.^88^ Despite advances in understanding OM accumulation for certain molecular groups, current approaches have inherent limitations. These limitations preclude the comprehensive analysis of structurally diverse chemical libraries. The lack of this type of analysis poses a significant challenge to establishing guiding principles for structure-accumulation relationships and achieving high predictive power based on chemical structure. Properly elucidating how chemical structure influences accumulation and local concentration – independently of MIC values – is crucial, as these structure-accumulation relationships are fundamental to drug discovery efforts.

Gram-negative bacteria, such as *E. coli*, have an OM and an inner plasma membrane that act as formidable barriers that severely restrict the movement of molecules into and out of the cell. The selective permeability of these membranes can make it challenging for antibiotics to reach intracellular targets. Being able to readily measure intracellular accumulation levels could potentially expedite drug discovery and development. Yet, this type of analysis continues to be a major challenge for the field. In developing CHAMP, we have leveraged the efficient and selective reaction of HaloTag with chloroalkane-modified molecules. Prior efforts using HaloTag in bacteria focused on the tracking of protein fusions^89,90^ and the role of cationic peptides on membrane permeabilization.^91^

We describe a straightforward and efficient platform, CHAMP, that effectively converts HaloTag expressed in the cytosol of *E. coli* into a strained alkyne. Test molecules tagged with an azide group are imprinted onto HaloTag upon their arrival in the cytosol. As a final step, the level of accumulation is determined by the number of unoccupied sites available for an azide-tagged fluorophore. We demonstrated that fluorescence can be readily measured using flow cytometry or standard plate readers, making it compatible with high-throughput screens. The robustness of the assay was confirmed empirically, and the throughput was exemplified by screening over 1,000 individual test molecules. Additionally, we demonstrated that CHAMP can be deployed to report on OM permeabilizers, and we are currently using it to screen for adjuvants that enhance the accumulation levels of antibiotics. Finally, we showed that the high-throughput data from CHAMP can be enhanced with emerging data science and cheminformatics tools to facilitate the discrimination of molecular profiles that increase recognition by the efflux pump AcrAB-TolC. We acknowledge that there are limitations to the CHAMP assay. Principally, in this iteration, the target pathogen must be amenable to genetic manipulation, and the molecule must be tagged with an azide. There is evidence that azides undergo modification within the biological milieu,^92,93,94^ a process shown to occur considerably more slowly for alkyl azides than for aryl azides, with cytosolic glutathione contributing to the reduction of azides to amines in *E. coli*. Although the SPAAC DBCO-azide reaction is relatively slow, it proceeds several orders of magnitude faster than the competing glutathione-mediated reduction of azides in biological systems, making this competing process less disruptive to the overall fluorescence readout in our CHAMP assays. Since all azides in the panel are exposed to the same intracellular glutathione environment, the CHAMP assay provides a reliable measure of relative accumulation across different azides under comparable conditions. An additional consideration is the potential impact of impurities in one of the small molecule-azide libraries, which may skew permeability of the molecules under test. Nevertheless, to minimize any confounding effects from solvent interactions, the concentration of DMSO, known to disrupt membrane integrity, was maintained below 1% in all working solutions, in accordance with the guidelines of the Clinical and Laboratory Standards Institute and previous findings by Tunçer et al.^95^

For antibiotics that do not efficiently cross the OM, an alternative strategy may involve co-treatment with an adjuvant molecule that disrupts the OM, thereby promoting the passage of molecules to their targets. The most widely used OM permeabilizer is PMBN, a fragment of the colistin family of antibiotics. Our results indicate that the OM is a significant barrier to the permeability of a subset of molecules, as its disruption led to a significant increase in fluorescence from azide-tagged dyes. Intriguingly, pre-treatment of cells with PMBN suggests that the disruption of OM integrity in *E. coli* caused by PMBN can persist even after the compound is washed away. Most importantly, we demonstrated that CHAMP could analyze the ability of molecules to permeabilize the OM of *E. coli* by directly measuring the increased permeability of other highly impermeable molecules, such as antibiotics. Given the capacity to adapt the assay to a plate-reading format, we will explore how it can be implemented in future high-throughput screening campaigns for novel OM permeabilizers.

For molecules that navigate past the OM, the next challenge is retention. In drug-resistant bacteria, efflux pumps can reduce the accumulation of molecules that are structurally unrelated.^96–98^ While it is generally understood that efflux pumps often play an important role in drug resistant phenotypes, it remains poorly defined how molecular structure drives recognition by efflux pumps. Assignment of substrates to specific efflux pumps has been made primarily by correlative analysis between expression levels and antibiotic susceptibility. Antibiotic susceptibility correlations have many disadvantages that can lead to mis-assignments. The deletion of porins can be a valuable tool to study recognition of substrates. We showed that CHAMP can be used in combination with the deletion of a primary efflux pump, AcrAB-TolC, to comprehensively test a large library of molecules for their potential recognition. In the future, we plan to start with EKO-35, a strain that has 35 of the efflux pumps deleted out of the genome,^99^ and then compare this strain to one in which a single efflux pump is introduced. This method will allow for the more precise assignment of structural preference among the various efflux pumps in *E. coli*.

Another major cellular machinery that can modulate the concentration of small molecules is porins imbedded within the OM of *E. coli*. Most FDA-approved antibiotics are hydrophilic compounds that are believed to penetrate OMs *via* porins.^100–103^ In *E. coli*, cefoxitin has been purported to primarily enter cells *via* OmpF and OmpC whereas ampicillin and ertapenem permeate through OmpC. The importance of porins for small molecule permeation is exemplified by the overexpression of the OM siderophore transporter FhuA, which effectively acts like an open and nonselective porin for the uptake of molecules. ^68,70^ Hyperporinated *E. coli* becomes highly susceptible to large antibiotics like vancomycin, novobiocin, and erythromycin. For many other molecules, the entry mechanism has not been fully established. The balance between porin-mediated entry and efflux pump mediated ejection out of the cell will ultimately dictate the effective concentration of molecules inside *E. coli*. Despite its medical significance, it has proven to be extremely challenging to precisely examine the role that each component has on permeation profiles.

## ACKNOWLEDGEMENT

This study was supported by the NIH grant 1R01AI178975-01 (M.M.P., W.I, and S.S.), R35GM124893 (M.M.P.), R01AI179080-01 (M.M.P., W.I, and S.S.) and the NSF grant MCB-2111728 (W.I.).

## SUPPORTING INFORMATION

Additional figures, tables, and materials/methods are included in the supporting information file.

## Reporting summary

Further information on data collection and sample size is available in the Reporting Summary linked to this article.

## Data availability

The datasets generated and/or analyzed during the current study are available within this article and its Supplementary Information and can be downloaded with no restrictions at https://doi.org/10.5281/zenodo.17434250

## Code availability

All raw data supporting the findings of this study are available within the article and its Supplementary Information files. Flow cytometry data is available in a spreadsheet that can be downloaded at https://doi.org/10.5281/zenodo.17434250 and is available to the public with no restrictions.

## Author contributions

Conceptualization: M.M.P, W.I., and M.S.S.; Methodology: M.M.P, W.I., M.S.S., and G.M.O.; Investigation: G.M.O., Z.L., and S.F.; Formal Analysis: G.M.O., Z.L., S.F., and M.S.G.; Resources: G.M.O., Z.L., M.D.C., R.D., Y.H., B.E.D., T.G., K.B.S., and J.D.; Writing-Original draft: M.M.P., M.S.S., W.I., and GM.O.; Writing-Review and Editing: M.M.P. and G.M.O.; Visualization: M.M.P.; Supervision: M.M.P, W.I., and M.S.S.; Project Administration and Funding Acquisition: M.M.P, W.I., and M.S.S.

## REFERENCES

1 Marston, H. D., Dixon, D. M., Knisely, J. M., Palmore, T. N. & Fauci, A. S. Antimicrobial Resistance. JAMA 316, 1193–1204 (2016). 10.1001/jama.2016.11764 2553454 [pii]

2 Kadri, S.S. Key Takeaways From the U.S. CDC’s 2019 Antibiotic Resistance Threats Report for Frontline Providers. Crit Care Med. 10.1097/CCM.0000000000004371. (2020) doi: 10.1097/CCM.0000000000004371

3 Six, D. A., Krucker, T. & Leeds, J. A. Advances and challenges in bacterial compound accumulation assays for drug discovery. Curr Opin Chem Biol 44, 9–15 (2018). 10.1016/j.cbpa.2018.05.005

4 Ferreira, R. J. & Kasson, P. M. Antibiotic Uptake Across Gram-Negative Outer Membranes: Better Predictions Towards Better Antibiotics. ACS Infect Dis 5, 2096–2104 (2019). 10.1021/acsinfecdis.9b00201

5 Hancock, R. E. The bacterial outer membrane as a drug barrier. Trends Microbiol 5, 37–42 (1997). 10.1016/S0966-842X(97)81773-8

6 Cama, J., Henney, A. M. & Winterhalter, M. Breaching the Barrier: Quantifying Antibiotic Permeability across Gram-negative Bacterial Membranes. J Mol Biol 431, 3531–3546 (2019). 10.1016/j.jmb.2019.03.031

7 Zhao, S. et al. Defining new chemical space for drug penetration into Gram-negative bacteria. Nat Chem Biol 16, 1293–1302 (2020). 10.1038/s41589-020-00674-6

8 Prajapati, J. D., Kleinekathofer, U. & Winterhalter, M. How to Enter a Bacterium: Bacterial Porins and the Permeation of Antibiotics. Chem Rev 121, 5158–5192 (2021). 10.1021/acs.chemrev.0c01213

9 Delcour, A. H. Outer membrane permeability and antibiotic resistance. Biochim Biophys Acta 1794, 808–816 (2009). 10.1016/j.bbapap.2008.11.005

10 Zgurskaya, H. I., Lopez, C. A. & Gnanakaran, S. Permeability Barrier of Gram-Negative Cell Envelopes and Approaches To Bypass It. ACS Infect Dis 1, 512–522 (2015). 10.1021/acsinfecdis.5b00097

11 Vergalli, J. et al. Porins and small-molecule translocation across the outer membrane of Gram-negative bacteria. Nat Rev Microbiol 18, 164–176 (2020). 10.1038/s41579-019-0294-2

12 Bolla, J. M. et al. Strategies for bypassing the membrane barrier in multidrug resistant Gram-negative bacteria. FEBS Lett 585, 1682–1690 (2011). 10.1016/j.febslet.2011.04.054

13 Vergalli, J., Refregiers, M., Ruggerone, P., Winterhalter, M. & Pages, J. M. Advances in methods and concepts provide new insight into antibiotic fluxes across the bacterial membrane. Commun Biol 7, 1508 (2024). 10.1038/s42003-024-07168-4

14 Hadjicharalambous, A. et al. Investigating Penetration and Antimicrobial Activity of Vector-Bicycle Conjugates. ACS Infect Dis 10, 2381–2389 (2024). 10.1021/acsinfecdis.3c00427

15 Iyer, R. et al. Evaluating LC-MS/MS To Measure Accumulation of Compounds within Bacteria. ACS Infect Dis 4, 1336–1345 (2018). 10.1021/acsinfecdis.8b00083

16 Widya, M. et al. Development and Optimization of a Higher-Throughput Bacterial Compound Accumulation Assay. ACS Infect Dis 5, 394–405 (2019). 10.1021/acsinfecdis.8b00299

17 Geddes, E. J., Li, Z. & Hergenrother, P. J. An LC-MS/MS assay and complementary web-based tool to quantify and predict compound accumulation in E. coli. Nat Protoc 16, 4833–4854 (2021). 10.1038/s41596-021-00598-y

18 Prochnow, H. et al. Subcellular Quantification of Uptake in Gram-Negative Bacteria. Anal. Chem. 91, 3, 1863–1872 (2019). https://pubs.acs.org/doi/10.1021/acs.analchem.8b03586#Abstract

19 Geddes, E. J. et al. Porin-independent accumulation in Pseudomonas enables antibiotic discovery. Nature 624, 145–153 (2023). 10.1038/s41586-023-06760-8

20 Richter, M. F. et al. Predictive compound accumulation rules yield a broad-spectrum antibiotic. Nature 545, 299–304 (2017). 10.1038/nature22308

21 Mientkiewicz, K. M., Peraro, L. & Kritzer, J. A. Parallel Screening Using the Chloroalkane Penetration Assay Reveals Structure-Penetration Relationships. ACS Chem Biol 16, 1184–1190 (2021). 10.1021/acschembio.1c00434

22 Peraro, L. et al. Cell Penetration Profiling Using the Chloroalkane Penetration Assay. J Am Chem Soc 140, 11360–11369 (2018). 10.1021/jacs.8b06144

23 Deprey, K. & Kritzer, J. A. Quantitative measurement of cytosolic penetration using the chloroalkane penetration assay. Methods Enzymol 641, 277–309 (2020). 10.1016/bs.mie.2020.03.003

24 Los, G. V. & Wood, K. The HaloTag: a novel technology for cell imaging and protein analysis. Methods Mol Biol 356, 195–208 (2007). 10.1385/1-59745-217-3:195

25 Los, G. V. et al. HaloTag: a novel protein labeling technology for cell imaging and protein analysis. ACS Chem Biol 3, 373–382 (2008). 10.1021/cb800025k

26 Hoelzel, C. A. & Zhang, X. Visualizing and Manipulating Biological Processes by Using HaloTag and SNAP-Tag Technologies. Chembiochem 21, 1935–1946 (2020). 10.1002/cbic.202000037

27 England, C. G., Luo, H. & Cai, W. HaloTag technology: a versatile platform for biomedical applications. Bioconjug Chem 26, 975–986 (2015). 10.1021/acs.bioconjchem.5b00191

28 Benink, H. A. & Urh, M. HaloTag technology for specific and covalent labeling of fusion proteins. Methods Mol Biol 1266, 119–128 (2015). 10.1007/978-1-4939-2272-7_8

29 Ongwae, G. M. et al. Measurement of Small Molecule Accumulation into Diderm Bacteria. ACS Infect Dis 9, 97–110 (2023). 10.1021/acsinfecdis.2c00435

30 Jewett, J. C. & Bertozzi, C. R. Cu-free click cycloaddition reactions in chemical biology. Chem Soc Rev 39, 1272–1279 (2010). 10.1039/b901970g

31 Agard, N. J., Prescher, J. A. & Bertozzi, C. R. A strain-promoted [3 + 2] azide-alkyne cycloaddition for covalent modification of biomolecules in living systems. J Am Chem Soc 126, 15046–15047 (2004). 10.1021/ja044996f

32 Meng, G. et al. Modular click chemistry libraries for functional screens using a diazotizing reagent. Nature 574, 86–89 (2019). 10.1038/s41586-019-1589-1

33 Guo, J., Wang, X., Ni, C., Wan, X. & Hu, J. SulfoxFluor-enabled deoxyazidation of alcohols with NaN(3). Nat Commun 13, 2752 (2022). 10.1038/s41467-022-30132-x

34 Krasheninina, O. A., Thaler, J., Erlacher, M. D. & Micura, R. Amine-to-Azide Conversion on Native RNA via Metal-Free Diazotransfer Opens New Avenues for RNA Manipulations. Angew Chem Int Ed Engl 60, 6970–6974 (2021). 10.1002/anie.202015034

35 Dieterich, D. C., Link, A. J., Graumann, J., Tirrell, D. A. & Schuman, E. M. Selective identification of newly synthesized proteins in mammalian cells using bioorthogonal noncanonical amino acid tagging (BONCAT). Proc Natl Acad Sci U S A 103, 9482–9487 (2006). 10.1073/pnas.0601637103

36 Kiick, K. L., Saxon, E., Tirrell, D. A. & Bertozzi, C. R. Incorporation of azides into recombinant proteins for chemoselective modification by the Staudinger ligation. Proc Natl Acad Sci U S A 99, 19–24 (2002). 10.1073/pnas.012583299

37 Saxon, E. & Bertozzi, C. R. Cell surface engineering by a modified Staudinger reaction. Science 287, 2007–2010 (2000). 10.1126/science.287.5460.2007

38 Liu, Z. et al. A Metabolic-Tag-Based Method for Assessing the Permeation of Small Molecules Across the Mycomembrane in Live Mycobacteria. Angew Chem Int Ed Engl 62, e202217777 (2023). 10.1002/anie.202217777

39 Apostolos, A. J. et al. Facile Synthesis and Metabolic Incorporation of m-DAP Bioisosteres Into Cell Walls of Live Bacteria. ACS Chem Biol 15, 2966–2975 (2020). 10.1021/acschembio.0c00618

40 Pidgeon, S. E. et al. L,D-Transpeptidase Specific Probe Reveals Spatial Activity of Peptidoglycan Cross-Linking. ACS Chem Biol 14, 2185–2196 (2019). 10.1021/acschembio.9b00427

41 Spangler, B. et al. Molecular Probes for the Determination of Subcellular Compound Exposure Profiles in Gram-Negative Bacteria. ACS Infect Dis 4, 1355–1367 (2018). 10.1021/acsinfecdis.8b00093

42 Spangler, B., Yang, S., Baxter Rath, C. M., Reck, F. & Feng, B. Y. A Unified Framework for the Incorporation of Bioorthogonal Compound Exposure Probes within Biological Compartments. ACS Chem Biol 14, 725–734 (2019). 10.1021/acschembio.9b00008

43 Kuru, E. et al. In Situ probing of newly synthesized peptidoglycan in live bacteria with fluorescent D-amino acids. Angew Chem Int Ed Engl. 7;51(50):12519–23 (2012). doi: 10.1002/anie.201206749.

44 Feigman, M. S. et al. Synthetic Immunotherapeutics against Gram-negative Pathogens. Cell Chem Biol 25, 1185–1194 e1185 (2018). 10.1016/j.chembiol.2018.05.019

45 Lee, J., Patel, D. S., Kucharska, I., Tamm, L. K. & Im, W. Refinement of OprH-LPS Interactions by Molecular Simulations. Biophys J 112, 346–355 (2017). 10.1016/j.bpj.2016.12.006

46 Mortimer, F. C., Mason, D. J. & Gant, V. A. Flow cytometric monitoring of antibiotic-induced injury in Escherichia coli using cell-impermeant fluorescent probes. Antimicrob Agents Chemother 44, 676–681 (2000). 10.1128/AAC.44.3.676-681.2000

47 Roth, B. L., Poot, M., Yue, S. T. & Millard, P. J. Bacterial viability and antibiotic susceptibility testing with SYTOX green nucleic acid stain. Appl Environ Microbiol 63, 2421–2431 (1997). 10.1128/aem.63.6.2421-2431.1997

48 Zhu, L., Rajendram, M. & Huang, K. C. Effects of fixation on bacterial cellular dimensions and integrity. iScience 24, 102348 (2021). 10.1016/j.isci.2021.102348

49 Stokes, J. M. et al. Pentamidine sensitizes Gram-negative pathogens to antibiotics and overcomes acquired colistin resistance. Nat Microbiol 2, 17028 (2017). 10.1038/nmicrobiol.2017.28

50 Klobucar, K. et al. Chemical Screen for Vancomycin Antagonism Uncovers Probes of the Gram-Negative Outer Membrane. ACS Chem Biol 16, 929–942 (2021). 10.1021/acschembio.1c00179

51 Vaara, M. Agents that increase the permeability of the outer membrane. Microbiol Rev 56, 395–411 (1992). 10.1128/mr.56.3.395-411.1992

52 Ejim, L. et al. Combinations of antibiotics and nonantibiotic drugs enhance antimicrobial efficacy. Nat Chem Biol 7, 348–350 (2011). 10.1038/nchembio.559

53 Poirel, L., Jayol, A. & Nordmann, P. Polymyxins: Antibacterial Activity, Susceptibility Testing, and Resistance Mechanisms Encoded by Plasmids or Chromosomes. Clin Microbiol Rev 30, 557–596 (2017). 10.1128/CMR.00064-16

54 French, S. et al. Potentiation of Antibiotics against Gram-Negative Bacteria by Polymyxin B Analogue SPR741 from Unique Perturbation of the Outer Membrane. ACS Infect Dis 6, 1405–1412 (2020). 10.1021/acsinfecdis.9b00159

55 Du, D. et al. Multidrug efflux pumps: structure, function and regulation. Nat Rev Microbiol 16, 523–539 (2018). 10.1038/s41579-018-0048-6

56 Nishino, K. & Yamaguchi, A. Analysis of a complete library of putative drug transporter genes in Escherichia coli. J Bacteriol 183, 5803–5812 (2001). 10.1128/JB.183.20.5803-5812.2001

57 Koronakis, V., Eswaran, J. & Hughes, C. Structure and function of TolC: the bacterial exit duct for proteins and drugs. Annu Rev Biochem 73, 467–489 (2004). 10.1146/annurev.biochem.73.011303.074104

58 Zgurskaya, H. I., Krishnamoorthy, G., Ntreh, A. & Lu, S. Mechanism and Function of the Outer Membrane Channel TolC in Multidrug Resistance and Physiology of Enterobacteria. Front Microbiol 2, 189 (2011). 10.3389/fmicb.2011.00189

59 Fralick, J. A. Evidence that TolC is required for functioning of the Mar/AcrAB efflux pump of Escherichia coli. J Bacteriol 178, 5803–5805 (1996). 10.1128/jb.178.19.5803-5805.1996

60 Gerken, H. & Misra, R. Genetic evidence for functional interactions between TolC and AcrA proteins of a major antibiotic efflux pump of Escherichia coli. Mol Microbiol 54, 620–631 (2004). 10.1111/j.1365-2958.2004.04301.x

61 Gellert, M., O’Dea, M. H., Itoh, T. & Tomizawa, J. Novobiocin and coumermycin inhibit DNA supercoiling catalyzed by DNA gyrase. Proc Natl Acad Sci U S A 73, 4474–4478 (1976). 10.1073/pnas.73.12.4474

62 Augustus, A. M., Celaya, T., Husain, F., Humbard, M. & Misra, R. Antibiotic-sensitive TolC mutants and their suppressors. J Bacteriol 186, 1851–1860 (2004). 10.1128/JB.186.6.1851-1860.2004

63 Renau, T. E. et al. Inhibitors of efflux pumps in Pseudomonas aeruginosa potentiate the activity of the fluoroquinolone antibacterial levofloxacin. J Med Chem 42, 4928–4931 (1999). 10.1021/jm9904598

64 Molinari, H. et al. Structure of vancomycin and a vancomycin/D-Ala-D-Ala complex in solution. Biochemistry 29, 9, 2271–2277 (1990). 10.1021/bi00461a010

65 Vaara, M. et al. Susceptibility of gram-negative bacteria to polymyxin B nonapeptide. Antimicrobial Agents and Chemotherapy 25(6):701–705. 10.1128/aac.25.6.701

66 Typas, A., Banzhaf, M., Gross, C. A. & Vollmer, W. From the regulation of peptidoglycan synthesis to bacterial growth and morphology. Nat. Rev. Microbiol. 10, 123–136 (2012).

67 Breukink, E. & de Kruijff, B. Lipid II as a target for antibiotics. Nat. Rev. Drug Discov. 5, 321–332 (2006).

68 Braun, M., Killmann, H., Maier, E., Benz, R. & Braun, V. Diffusion through channel derivatives of the Escherichia coli FhuA transport protein. Eur J Biochem 269, 4948–4959 (2002). 10.1046/j.1432-1033.2002.03195.x

69 Sabnis, A. et al. Colistin kills bacteria by targeting lipopolysaccharide in the cytoplasmic membrane. Elife 10 (2021). https://doi.org/ARTNe6583610.7554/eLife.65836

70 Krishnamoorthy, G. et al. Breaking the Permeability Barrier of Escherichia coli by Controlled Hyperporination of the Outer Membrane. Antimicrob Agents Chemother 60, 7372–7381 (2016). 10.1128/AAC.01882-16

71 Dommerholt, J. et al. Highly accelerated inverse electron-demand cycloaddition of electron-deficient azides with aliphatic cyclooctynes. Nat Commun 5, 5378 (2014). 10.1038/ncomms6378

72 Zimmerman, ES. et al. Production of Site-Specific Antibody–Drug Conjugates Using Optimized Non-Natural Amino Acids in a Cell-Free Expression System. Bioconjugate Chem. 25, 2, 351–361 (2014)

73 Bohnert, J. A. et al. Novel Piperazine Arylideneimidazolones Inhibit the AcrAB-TolC Pump in Escherichia coli and Simultaneously Act as Fluorescent Membrane Probes in a Combined Real-Time Influx and Efflux Assay. Antimicrob Agents Chemother 60, 1974–1983 (2016). 10.1128/AAC.01995-15

74 Compagne, N. et al. Optimization of pyridylpiperazine-based inhibitors of the Escherichia coli AcrAB-TolC efflux pump. Eur J Med Chem 259, 115630 (2023). 10.1016/j.ejmech.2023.115630

75 Chan, P. F. et al. Thiophene antibacterials that allosterically stabilize DNA-cleavage complexes with DNA gyrase. Proc Natl Acad Sci U S A 114, E4492–E4500 (2017). 10.1073/pnas.1700721114

76 Thorarensen, A. et al. 3-Arylpiperidines as potentiators of existing antibacterial agents. Bioorg Med Chem Lett 11, 1903–1906 (2001). 10.1016/s0960-894x(01)00330-4

77 Armitage, E. G. et al. Metabolic Clustering Analysis as a Strategy for Compound Selection in the Drug Discovery Pipeline for Leishmaniasis. ACS Chem Biol 13, 1361–1369 (2 Hadjicharalambous, A.018). 10.1021/acschembio.8b00204

78 Keren, I., Wu, Y., Inocencio, J., Mulcahy, L. R. & Lewis, K. Killing by bactericidal antibiotics does not depend on reactive oxygen species. Science 339, 1213–1216 (2013). 10.1126/science.1232688 339/6124/1213 [pii]

79 Cho, H., Uehara, T. & Bernhardt, T. G. Beta-Lactam Antibiotics Induce a Lethal Malfunctioning of the Bacterial Cell Wall Synthesis Machinery. Cell 159, 1300–1311 (2014). 10.1016/j.cell.2014.11.017

80 June, C. M. et al. A fluorescent carbapenem for structure function studies of penicillin-binding proteins, beta-lactamases, and beta-lactam sensors. Anal Biochem 463, 70–74 (2014). 10.1016/j.ab.2014.07.012

81 Zhou, Y. et al. Thinking outside the “bug”: a unique assay to measure intracellular drug penetration in gram-negative bacteria. Anal Chem 87, 3579–3584 (2015). 10.1021/ac504880r

82 Davis, T. D., Gerry, C. J. & Tan, D. S. General platform for systematic quantitative evaluation of small-molecule permeability in bacteria. ACS Chem Biol 9, 2535–2544 (2014). 10.1021/cb5003015

83 Cama, J. et al. Quantification of Fluoroquinolone Uptake through the Outer Membrane Channel OmpF of Escherichia coli. Journal of the American Chemical Society 137, 13836–13843 (2015). 10.1021/jacs.5b08960

84 Ghai, I., Winterhalter, M. & Wagner, R. Probing transport of charged beta-lactamase inhibitors through OmpC, a membrane channel from E. coli. Biochem Biophys Res Commun 484, 51–55 (2017). https://doi.org/S0006-291X(17)30126-2 [pii]10.1016/j.bbrc.2017.01.076

85 Kascakova, S., Maigre, L., Chevalier, J., Refregiers, M. & Pages, J. M. Antibiotic transport in resistant bacteria: synchrotron UV fluorescence microscopy to determine antibiotic accumulation with single cell resolution. PLoS One 7, e38624 (2012). 10.1371/journal.pone.0038624 PONE-D-12-05250 [pii]

86 Cinquin, B. et al. Microspectrometric insights on the uptake of antibiotics at the single bacterial cell level. Sci Rep 5, 17968 (2015). 10.1038/srep17968 srep17968 [pii]

87 Vergalli, J. et al. Spectrofluorimetric quantification of antibiotic drug concentration in bacterial cells for the characterization of translocation across bacterial membranes. Nat Protoc 13, 1348–1361 (2018). 10.1038/nprot.2018.036

88 Dash, R., Holsinger, K. A., Chordia, M. D., Gh, M. S. & Pires, M. M. Bioluminescence-Based Determination of Cytosolic Accumulation of Antibiotics in Escherichia coli. ACS Infect Dis 10, 1602–1611 (2024). 10.1021/acsinfecdis.3c00684

89 Ke, N., Landgraf, D., Paulsson, J. & Berkmen, M. Visualization of Periplasmic and Cytoplasmic Proteins with a Self-Labeling Protein Tag. J Bacteriol 198, 1035–1043 (2016). 10.1128/JB.00864-15

90 Clark, S. A., Singh, V., Vega Mendoza, D., Margolin, W. & Kool, E. T. Light-Up “Channel Dyes” for Haloalkane-Based Protein Labeling in Vitro and in Bacterial Cells. Bioconjug Chem 27, 2839–2843 (2016). 10.1021/acs.bioconjchem.6b00613

91 Yang, Z. & Weisshaar, J. C. HaloTag Assay Suggests Common Mechanism of E. coli Membrane Permeabilization Induced by Cationic Peptides. ACS Chem Biol 13, 2161–2169 (2018). 10.1021/acschembio.8b00336

92 Handlon AL, Oppenheimer NJ. Thiol reduction of 3’-azidothymidine to 3’-aminothymidine: kinetics and biomedical implications. Pharm Res. 5(5):297–9 (1988). doi: 10.1023/a:1015926720740. PMID: 3244639.

93 Zhang C, Dai P, Vinogradov AA, Gates ZP, and Pentelute LB. Site-Selective Cysteine–Cyclooctyne Conjugation. Angew. Chem. Int. Ed. 57, 6459 (2018).

94 Bertozzi CR, Sletten EM. Bioorthogonal Chemistry: Fishing for Selectivity in a 94 Sea of Functionality. Chem. Commun.49, 11007–11022 (2013)

95 Tunçer et al., Modulation of bacterial membranes and cellular macromolecules by dimethyl sulfoxide: A dose-dependent study providing novel insights. Int. J. Biol. Macromol. 267(Pt 2):131581(2024). doi:10.1016/j.ijbiomac.2024.131581]

96 Nikaido, H. & Pages, J. M. Broad-specificity efflux pumps and their role in multidrug resistance of Gram-negative bacteria. FEMS Microbiol Rev 36, 340–363 (2012). 10.1111/j.1574-6976.2011.00290.x

97 Pages, J. M., James, C. E. & Winterhalter, M. The porin and the permeating antibiotic: a selective diffusion barrier in Gram-negative bacteria. Nat Rev Microbiol 6, 893–903 (2008). 10.1038/nrmicro1994

98 Krishnamoorthy, G. et al. Synergy between Active Efflux and Outer Membrane Diffusion Defines Rules of Antibiotic Permeation into Gram-Negative Bacteria. mBio 8 (2017). 10.1128/mBio.01172-17

99 Teelucksingh, T. et al. A genetic platform to investigate the functions of bacterial drug efflux pumps. Nat Chem Biol 18, 1399–1409 (2022). 10.1038/s41589-022-01119-y

100 Isabella, V. M. et al. Toward the rational design of carbapenem uptake in Pseudomonas aeruginosa. Chem Biol 22, 535–547 (2015). 10.1016/j.chembiol.2015.03.018 S1074-5521(15)00112-X [pii]

101 Iyer, R. et al. Whole-Cell-Based Assay To Evaluate Structure Permeation Relationships for Carbapenem Passage through the Pseudomonas aeruginosa Porin OprD. ACS Infect Dis 3, 310–319 (2017). 10.1021/acsinfecdis.6b00197

102 Sugawara, E., Kojima, S. & Nikaido, H. Klebsiella pneumoniae Major Porins OmpK35 and OmpK36 Allow More Efficient Diffusion of beta-Lactams than Their Escherichia coli Homologs OmpF and OmpC. Journal of Bacteriology 198, 3200–3208 (2016). https://doi.org/JB.00590-16 [pii]10.1128/JB.00590-16

103 Tran, Q. T. et al. Structure-kinetic relationship of carbapenem antibacterials permeating through E. coli OmpC porin. Proteins 82, 2998–3012 (2014). 10.1002/prot.24659

