## Supporting Information for "Click-Based Determination of Accumulation of Molecules in *Escherichia coli*"

### TABLE OF CONTENTS

#### SUPPLEMENTARY FIGURES

| | |
|---|---|
| **Figure S1.** Confocal fluorescence images of *E. coli* cells expressing HaloTag and labeled with D-Lys(FITC) and treated with Comcl | S5 |

| | |
|---|---|
| **Figure S2.** Concentration dependent competition with DBCOcl in BaCAPA | S6 |
| **Figure S3.** Concentration dependent response of DBCOcl treatment | S7 |
| **Figure S4.** Confocal fluorescence images of *E. coli* cells expressing HaloTag, labeled with D-Lys(FITC) and treated with DBCOCl, Comaz. | S8 |
| **Figure S5.** CHAMP using smooth strain of *E. coli* | S9 |
| **Figure S6.** Cellular integrity examination with SYTOX green after DBCOcl treatment | S10 |
| **Figure S7.** Impact of formaldehyde fixation on fluorescence signal levels | S11 |
| **Figure S8.** Colony-forming unit (CFU) analysis of *E. coli* treated with PMBN | S12 |
| **Figure S9.** Concentration dependent permeabilization by PMBN | S13 |
| **Figure S10.** Uptake of fluorophores in *E. coli* in the presence/absence of PMBN | S14 |
| **Figure S11.** Determination of the extent of non-specific labelling following incubation with the coumarin series **6a**-**6e** | S15 |
| **Figure S12.** Comparison between direct readouts of fluorescence and the readout using CHAMP for the coumarin series **6a**-**6e** | S16 |
| **Figure S13.** Schematic representation of the DBCO-modified beads assay | S17 |
| **Figure S14.** Reactivity profile of the azido library | S18 |
| **Figure S15.** High throughput benchmarking for CHAMP | S19 |
| **Figure S16.** Reactivity profile of the 1152 azido compounds in the library | S20 |
| **Figure S17.** Reactivity cutoff parameter scan on physicochemical property correlations | S21 |
| **Figure S18.** Analysis of compounds that are more significantly impacted by TolC | S22 |
| **Table S1.** Pearson R correlation between physicochemical properties and compound efflux | S23 |
| **BIOLOGICAL METHODS** | |
| **Table S2.** Bacterial strains and plasmids used in this study | S24 |
| Plasmid construction for HaloTag expression in *E. coli* | S24 |
| HaloTag protein amino acid sequence. | S25 |
| HaloTag protein expression and bacterial cell culture | S25 |
| One-step assay with rhodamine chloroalkane (R110cl). | S25 |
| Three-step assay: - DBCOcl incubation, azido-tagged small molecule (pulse), and Flaz dye (chase) | S26 |
| Two-step assay without fixation prior to chase step: - DBCOcl incubation, azido-tagged dye | S27 |
| Two-step assay with fixation prior to chase step: - DBCOcl incubation, formaldehyde fixation, and azido-tagged dye | S27 |
| Assay with PMBN: - DBCOcl incubation, azido-tagged dye coincubation with PMBN | S27 |
| 1152 azido-tagged test molecules library screen | S28 |

| | |
|---|---|
| Cell preparation for confocal microscopy. | S29-S31 |
| Novobiocin azide (**novAZ1**) accumulation assay in WT *E. coli* BW25113 and ΔtolC *E. coli* BW25113 with or without efflux inhibitor PAβN | S31 |
| **SYNTHESIS AND CHARACTERIZATION OF ANTIBIOTICS/SMALL MOLECULES** | S32 |
| **General information about materials and instruments** | S32-S35 |
| Synthesis and characterization of **DBCOcl** | S35-S38 |
| Synthesis and characterization of **vanAZ1** | S39-S42 |
| Synthesis and characterization of **rifAZ1** | S43-S45 |
| Synthesis and characterization of **triAZ1** | S46-S47 |
| Synthesis and characterization of **moxAZ1** | S48-S53 |
| Synthesis and characterization of **novAZ1** | S54-S58 |
| Synthesis and characterization of **purAZ1** | S59-S63 |
| Synthesis and characterization of **zolAZ1** | S64-S67 |
| Synthesis and characterization of **zolAZ2** | S68-S69 |
| Synthesis and characterization of **zolAZ3** | S70-S71 |
| Synthesis and characterization of **metAZ1** | S72-S73 |
| Synthesis and characterization of **sulAZ1** | S74-S77 |
| **General methods for synthesis of peptide probes** | S78-S79 |
| Synthesis and characterization of **1p** and **1n** | S80-S88 |
| Synthesis and characterization of **2p** and **2n** | S89-S98 |
| Synthesis and characterization of **3p** | S99-S102 |
| Synthesis and characterization of **4a**, **4b**, **4c**, **4d**, and **4e** | S103-S124 |
| Synthesis and characterization of **5a**, **5b**, **5c**, **5d**, and **5e** | S125-S149 |
| Synthesis and characterization of **6a**, **6b**, **6c**, **6d**, and **6e** | S150-S178 |
| 1152 Library of azido-tagged molecules (Dong/Sharpless Library) | S179-S180 |
| 404 Library of azido-tagged molecules (Enamine/Sigma) | S181-S183 |
| **REFERENCES** | S184 |


**SUPPLEMENTARY FIGURES**

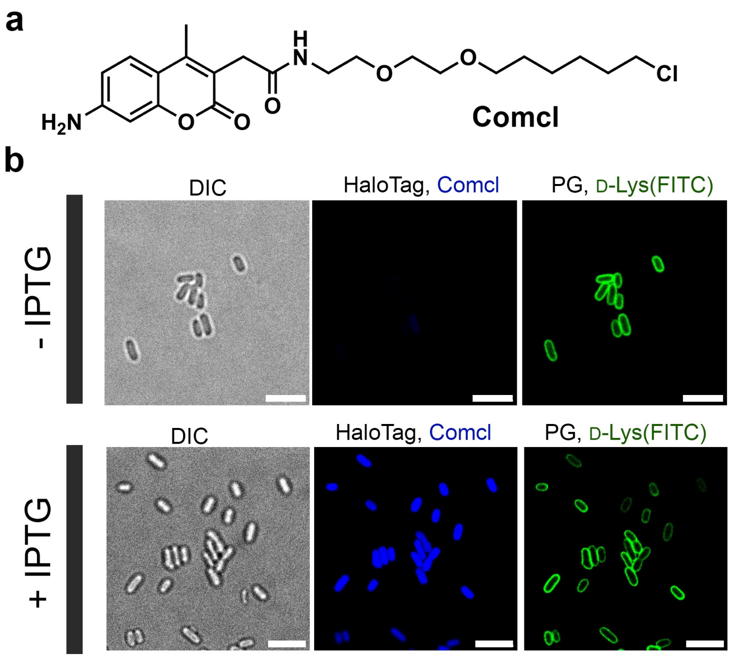


**Figure S1. Confocal fluorescence images of *E. coli* cells expressing HaloTag, labeled with D-Lys(FITC) and treated with coumarin chloroalkane to produce a blue cytosolic signal.** *E. coli* cells harboring the plasmid pQE-30_HaloTag (Amp) were cultured in LB medium supplemented with 500 μM D-Lys(FITC) and incubated at 37 °C with shaking for 2 h. Protein expression was induced with 1 mM isopropyl β-D-1-thiogalactopyranoside (IPTG), and cells were incubated for 1 h to allow expression of the HaloTag protein. Next, cells were washed three times with phosphate-buffered saline (PBS) and incubated with 50 μM coumarin chloroalkane (Comcl structure shown on panel **a**) for 30 min at 37 °C with shaking. Cells were subsequently washed three times with PBS, fixed and imaged. Excitation was performed using 405 nm (Coumarin) and 488 nm (FITC) laser lines; panel **b** shows the images in different channels. Scale bar: 2 μm.

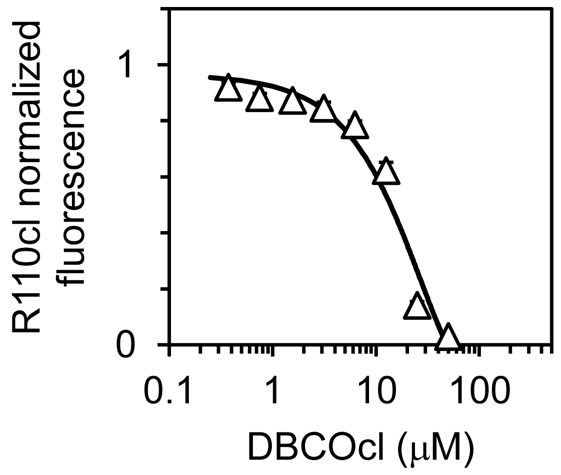


**Figure S2. Concentration dependent competition with DBCOcl in BaCAPA.** *E. coli* cells expressing HaloTag protein were incubated with increasing concentrations of DBCOcl for 30 min followed by treatment with 50 μM of R110cl for 1 h. Data are represented as mean +/- SD of technical replicates (n = 3 independent samples in a single experiment).

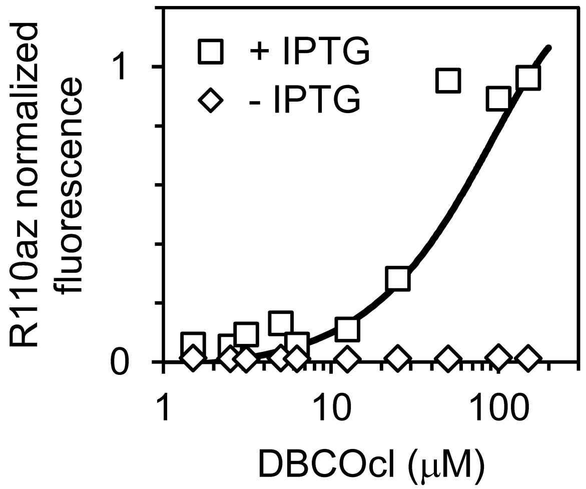


**Figure S3. Concentration dependent response of DBCOcl treatment.** *E. coli* cells were treated with or without IPTG for inducing the expression of HaloTag protein, followed by incubation with increasing concentrations of DBCOcl for 30 min and treatment with 50 μM of R110az for 1 h. Data are represented as mean +/- SD of technical replicates (n = 3 independent samples in a single experiment).

**a**

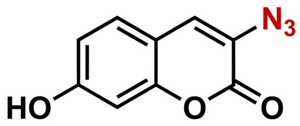


**b**

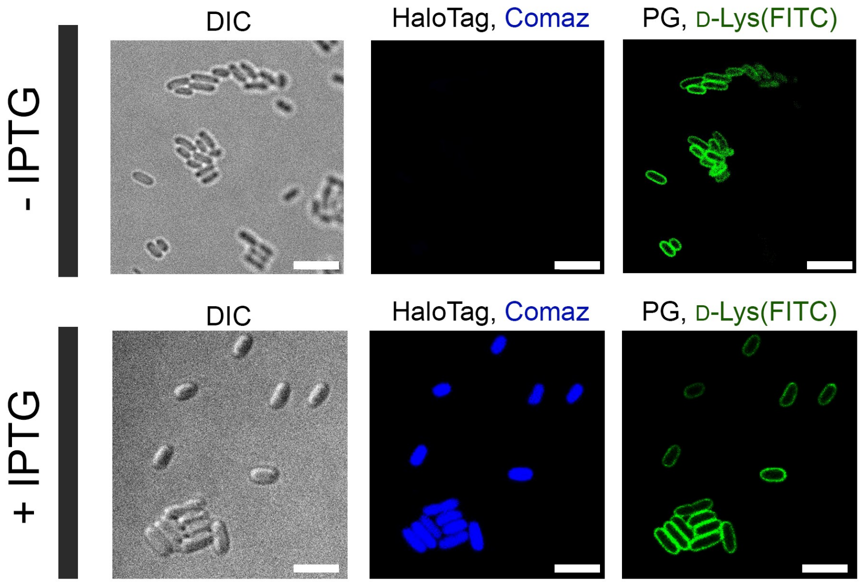


**Figure S4. Confocal fluorescence images of *E. coli* cells expressing HaloTag, labeled with D-Lys(FITC) and treated with DBCOCl, which subsequently reacted with Comaz.** *E. coli* cells harboring the plasmid pQE-30_HaloTag (Amp) were cultured in LB medium supplemented with 500 μM D-Lys(FITC) and incubated at 37 °C with shaking for 2 h. Protein expression was induced with 1 mM isopropyl β-D-1-thiogalactopyranoside (IPTG), and cells were incubated for 1 h to allow expression of the HaloTag protein. Uninduced cultures were maintained as negative controls. Next, cells were washed three times with phosphate-buffered saline (PBS) and incubated with 50 μM DBCOcl for 30 min at 37 °C with shaking. Cells were subsequently washed three times with PBS, fixed with 4% (w/v) formaldehyde for 30 min, washed again, and incubated with 50 μM Comaz (structure shown on panel **a**) for 1 h at 37 °C with shaking. Finally, cells were washed three times and mounted on 1% (w/v) agarose pads for imaging by confocal microscopy. Excitation was performed using 405 nm (Coumarin) and 488 nm (FITC) laser lines; panel **b** shows the images in different channels. Scale bar: 2 μm.

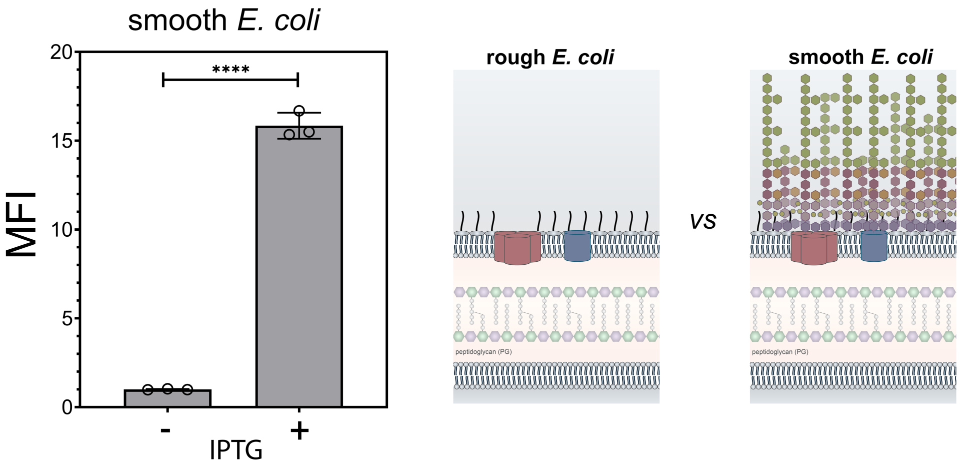


**Figure S5. CHAMP using smooth strain of *E. coli.*** Flow cytometry analysis of smooth *E. coli* strain (ATCC 25922) in the presence/absence of IPTG using the CHAMP workflow with R110az. *E. coli* ATCC 25922 has intact lipopolysaccharides on its outer surface (right). Data are represented as mean +/- SD of technical replicates (n = 3 independent samples in a single experiment). P-values were determined by a two-tailed t-test (**** denotes a p-value <0.0001).

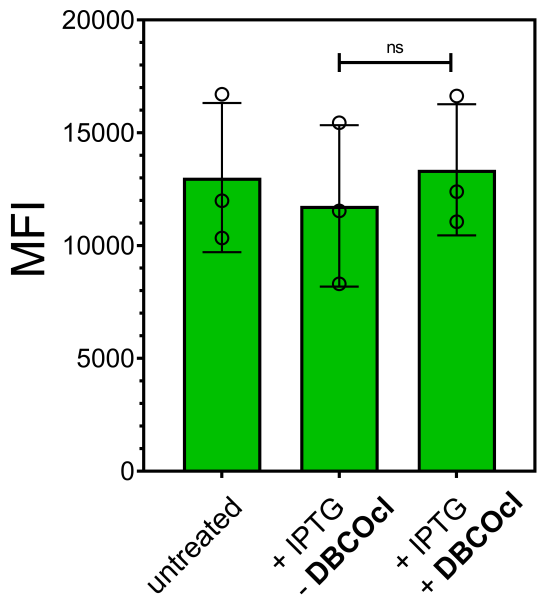


**Figure S6. Cellular viability examination with SYTOX green after DBCOcl treatment.** *E. coli* induced to express HaloTag protein were treated with DBCOcl followed by treatment with 5 μM of SYTOX Green stain for 15 min; uninduced cells were used as a negative control. Cellular fluorescence was measured by flow cytometry. Data are represented as mean +/- SD of technical replicates (n = 3 independent samples in a single experiment). P-values were determined by a two-tailed t-test (ns = not significant).

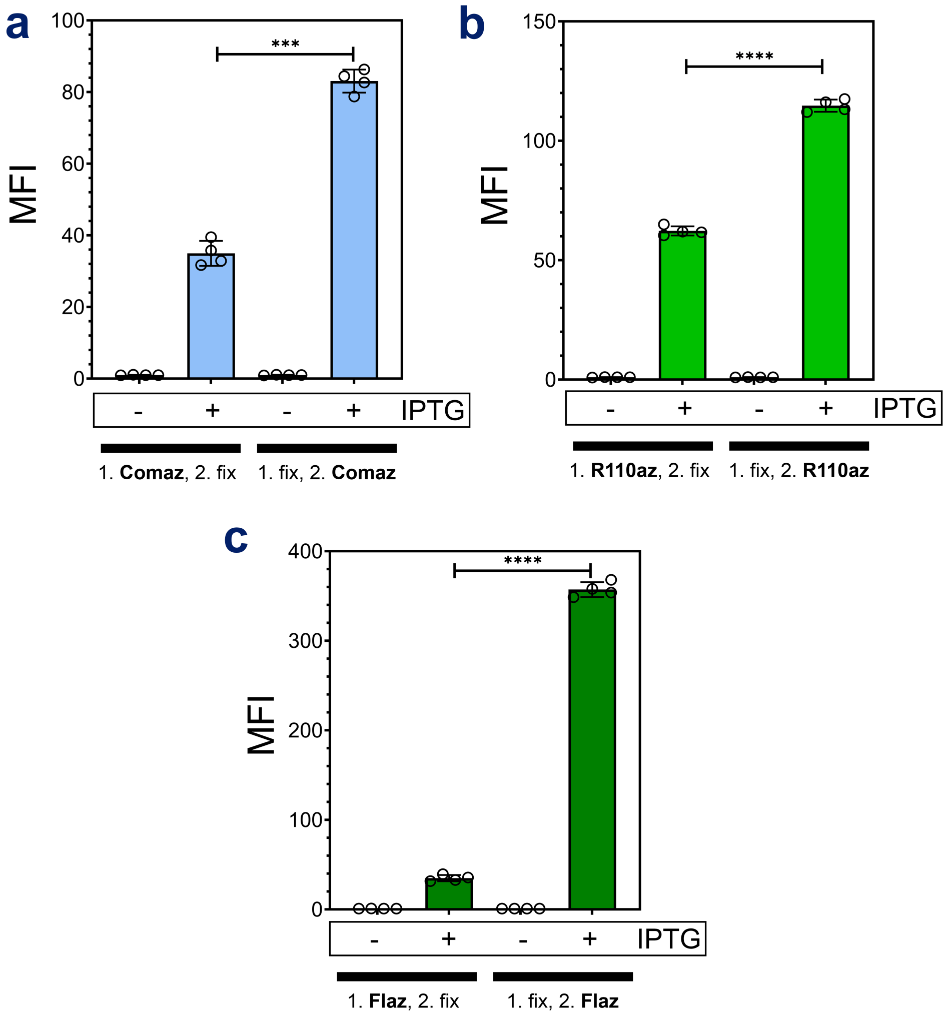


**Figure S7. Impact of formaldehyde fixation on fluorescence signal levels.** For formaldehyde treatment, cells were incubated with 4% formaldehyde in PBS for 30 min at room temperature followed by 3X wash with PBS. Dye treatment of Comaz (**a**), R110az (**b**), and Flaz (**c**) was performed at 25 μM for 1 h at room temperature either before or after fixation. Cellular fluorescence intensities were measured by flow cytometry. Data are represented as mean +/- SD of technical replicates (n = 3 independent samples in a single experiment). P-values were determined by a two-tailed t-test (*** denotes a p-value <0.001, **** <0.0001).

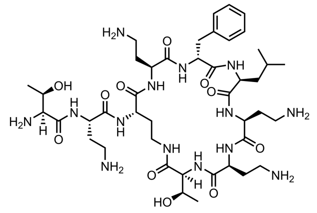

PMBN

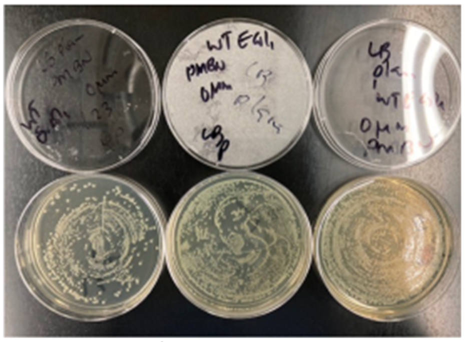

0 μM PMBN

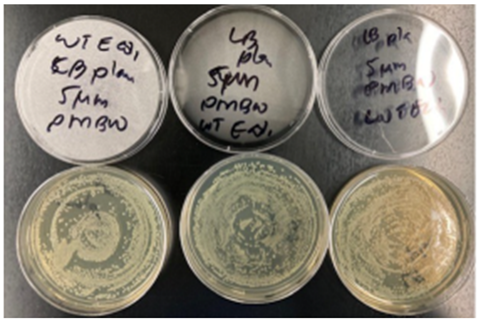

5 μM PMBN

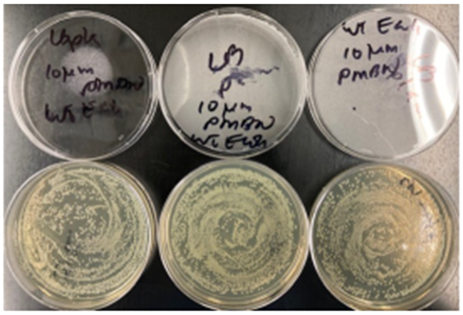

10 μM PMBN

**Figure S8. Colony-forming unit (CFU) analysis of *E. coli* treated with PMBN.** Top left: structure of PMBN. *E. coli* cells that were induced with IPTG were treated with vehicle or 5 or 10 μM of PMBN for 1 h at 37 °C in PBS. Next, cells were spread on agar/LB plates which were placed in a 37 °C incubator for 15 h. Images of three individual sets of agar plates are shown.

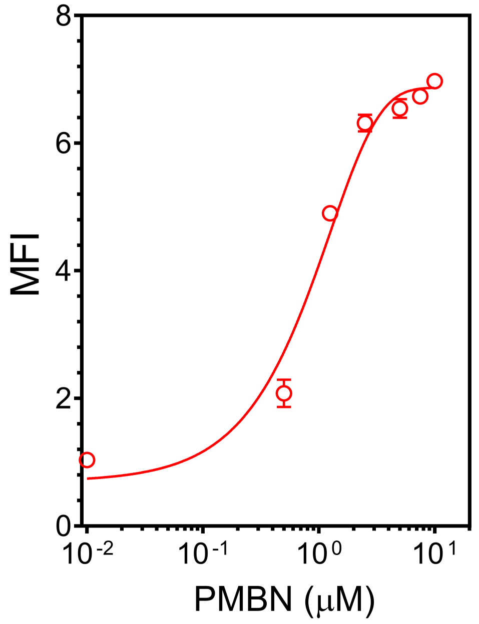


**Figure S9. Concentration dependent permeabilization by PMBN.** *E. coli* cells were treated with increasing concentrations of PMBN, fixed with formaldehyde, and treated with 25 μM of Flaz. Data is represented as mean +/- SD of technical replicates (n = 3 independent samples in a single experiment).

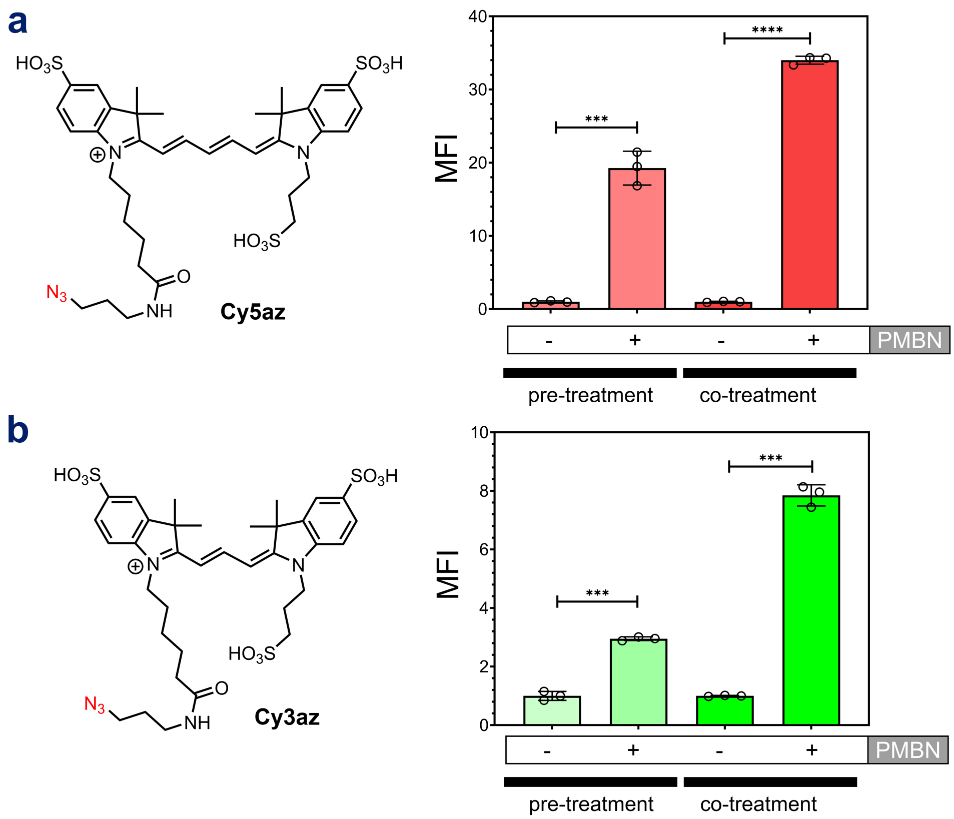


**Figure S10. Uptake of fluorophores in *E. coli* in the presence/absence of PMBN.** *E. coli* cells were treated with either 30 min co-incubation of 50 μM Cy5az or 50 μM Cy3az along with 10mM PMBN, or stepwise incubation with pre-treatment of PMBN for 30 min, followed with either fluorophore for 30 min. Cellular fluorescence intensities were measured by the plate reader with excitation at 649nm and 550nm respectively. Data are represented as mean +/- SD of technical replicates (n = 3 independent samples in a single experiment). P-values were determined by a two-tailed t-test (* denotes a p-value < 0.05, ** < 0.01, ***<0.001, ns = not significant).

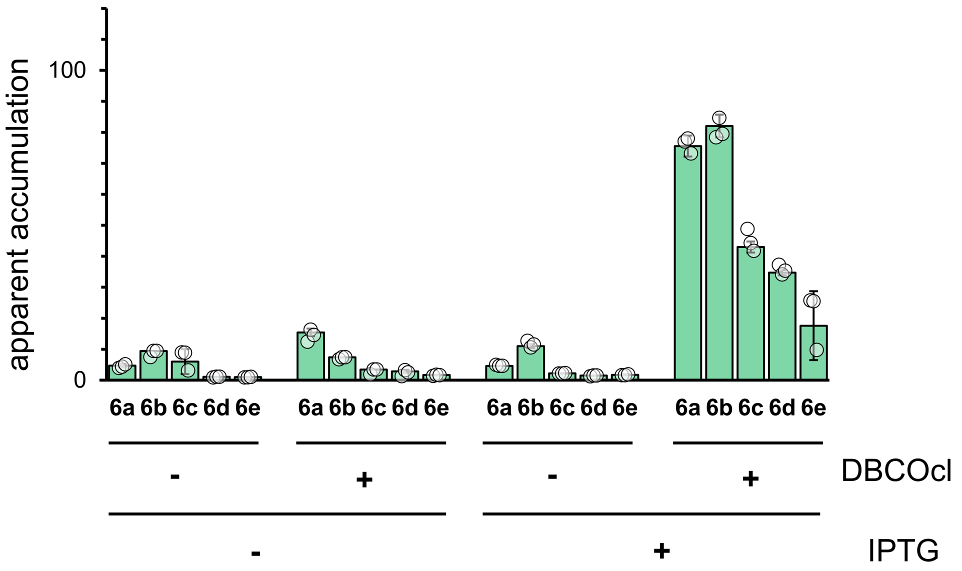


**Figure S11. Determination of the extent of non-specific labelling following incubation with the coumarin series 6a-6e.** Cells were induced to express HaloTag and treated with DBCOcl or a PBS control followed by incubation with the coumarin series **6a** – **6e**; uninduced cells were used as a negative control. Data are represented as mean +/- SD of technical replicates (n = 3 independent samples in a single experiment).

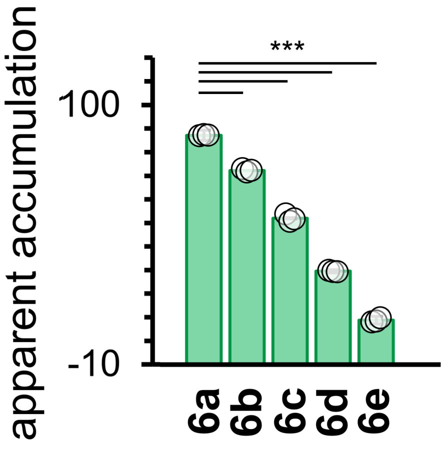


**Figure S12. Comparison between direct readouts of fluorescence and the readout using CHAMP for the coumarin series 6a-6e.** The accumulation profile across **6a**-**6e** was nearly identical between the direct readout and the readout using the usual CHAMP data analysis. Data are represented as mean +/- SD of technical replicates (n = 3 independent samples in a single experiment).

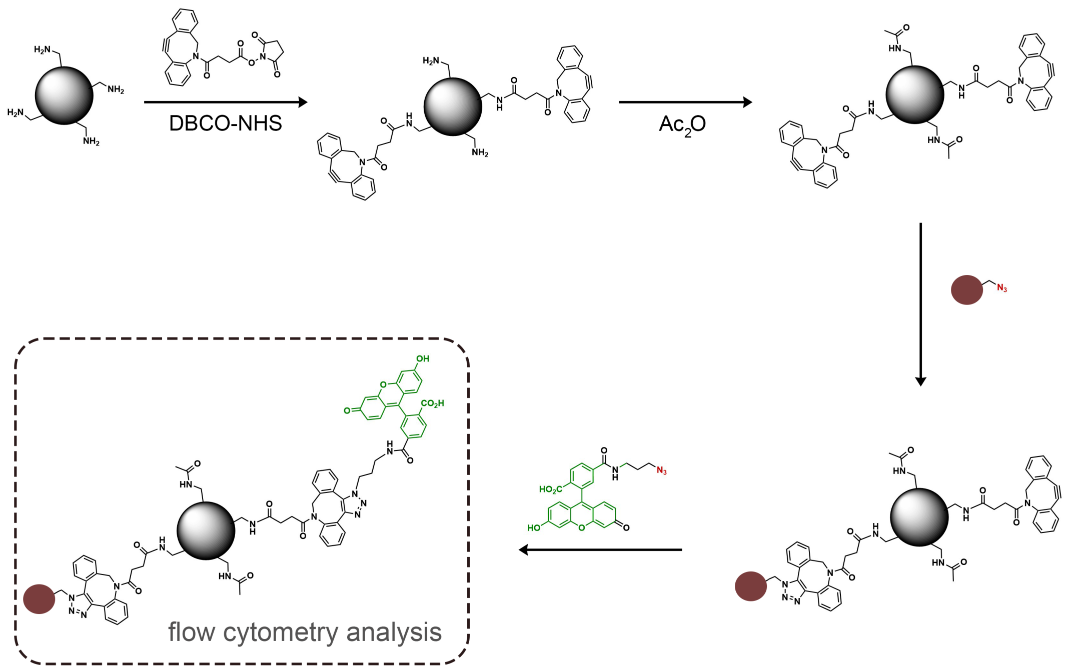


**Figure S13. Schematic representation of the DBCO-modified beads assay.** Amino-terminated polystyrene beads were reacted with NHS-activated DBCO. Free amines that remained unreacted were capped with acetic anhydride. DBCO-modified beads were incubated with the panel of 50 μM azide-modified molecules for 2 h at 37 $^{o}$C in PBS, followed by 1 h incubation with 50 μM Flaz, and analyzed by flow cytometry. The fraction of starting MFI is the ratio of the mean fluorescence intensity of the beads with the treatment of an individual molecule relative to the treatment with DMSO.

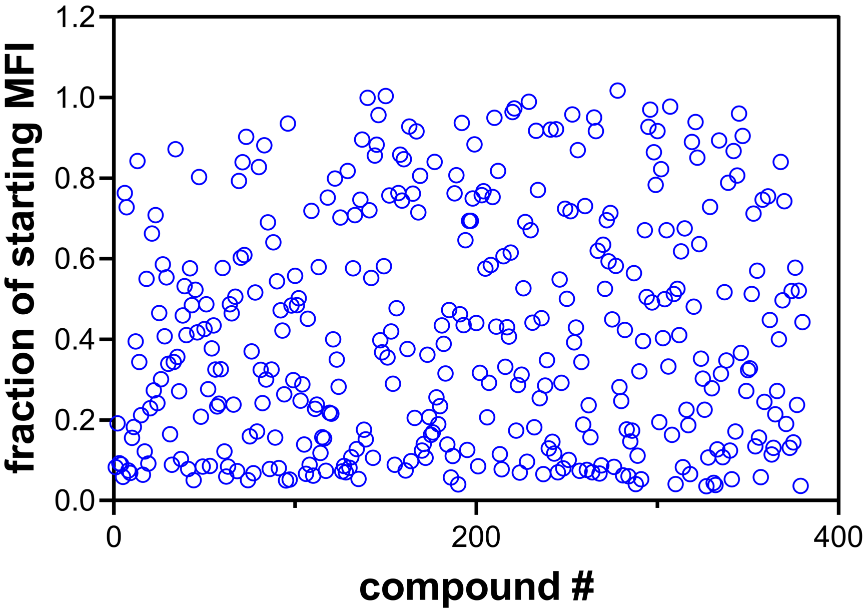


**Figure S14. Reactivity profile of the azido library.** DBCO-modified beads were incubated with 50 μM azide-modified molecules for 1 h at 37 °C in PBS, followed by 1 h incubation with 50 μM Flaz. Fluorescence intensities of the beads were measured by flow cytometry. The fraction of starting MFI is the ratio of the mean fluorescence intensity of the beads with the treatment of an individual molecule relative to the treatment of DMSO.

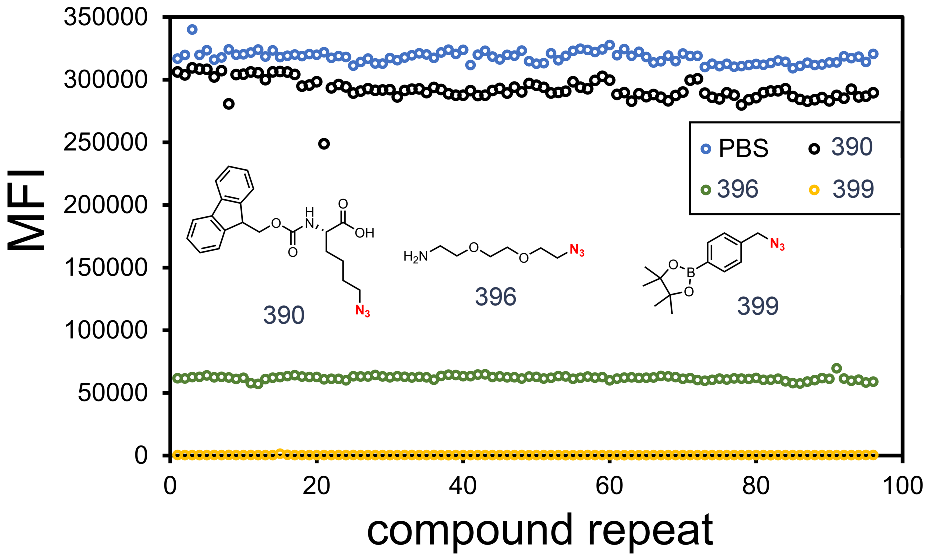


**Figure S15. High throughput benchmarking for CHAMP.** CHAMP was performed in a 96-well plate format that included test molecules with high, low, and intermediate accumulation levels. These data resulted in a Z' score of 0.954.

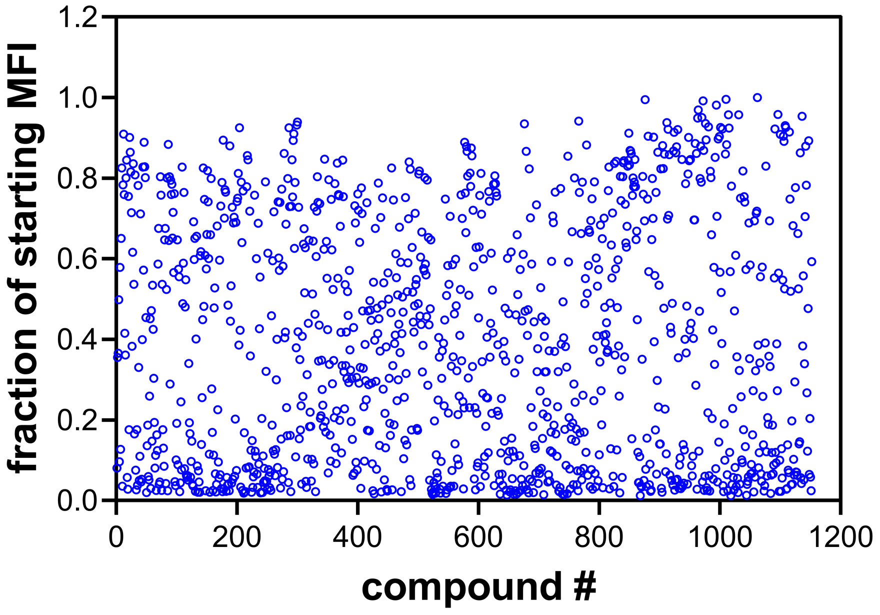


**Figure S16. Reactivity profile of the 1152 azido compounds in the library.** DBCO-modified beads were incubated with 50 μM azide-modified molecules for 2 h at 37 °C in PBS, followed by 1 h incubation with 50 μM Flaz. Fluorescence intensities of the beads were measured by flow cytometry. The fraction of starting MFI is the ratio of the mean fluorescence intensity of the beads with the treatment of an individual molecule relative to the treatment of DMSO.

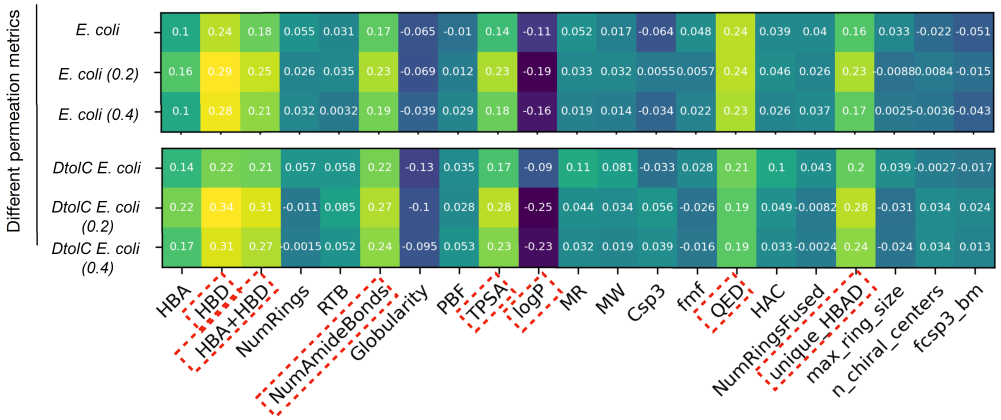


**Figure S17. Reactivity cutoff parameter scan on physicochemical property correlations.** A heatmap of the Pearson R correlation constants is shown, annotated with the Pearson R values. For the rows, the top rows represent no data filtering; 0.2/0.4 represents filtering out datapoints that have reactivity values less than 0.2/0.4. HBA: hydrogen bond acceptor; HBD: hydrogen bond donor; HBA+HBD: sum of number of HBA and HBD; NumRings: number of rings; RTB: number of rotatable bonds; NumAmideBonds: number of amide bonds; Globularity: division of volume by surface area; PBF: plane of best-fit; TPSA: topological polar surface area; logP**:** partition between water and octanol; MR: molecular refractivity, a measure of polarizability; MW: molecular weight; Csp3: fraction of sp3 carbons; fmf: fraction of atoms belonging to Murcko framework; QED: quantitative estimate of drug-likeness; HAC: heavy atom count; NumRingsFused: number of rings fused; unique_HBAD: non-redundant hydrogen bond acceptors and donors; max_ring_size: the size of maximum ring; n_chiral_centers: number of chiral centers; fcsp3_bm: fraction of sp3 carbon atoms in the Bemis-Murcko scaffold, a measure of saturation of scaffold carbon atoms.

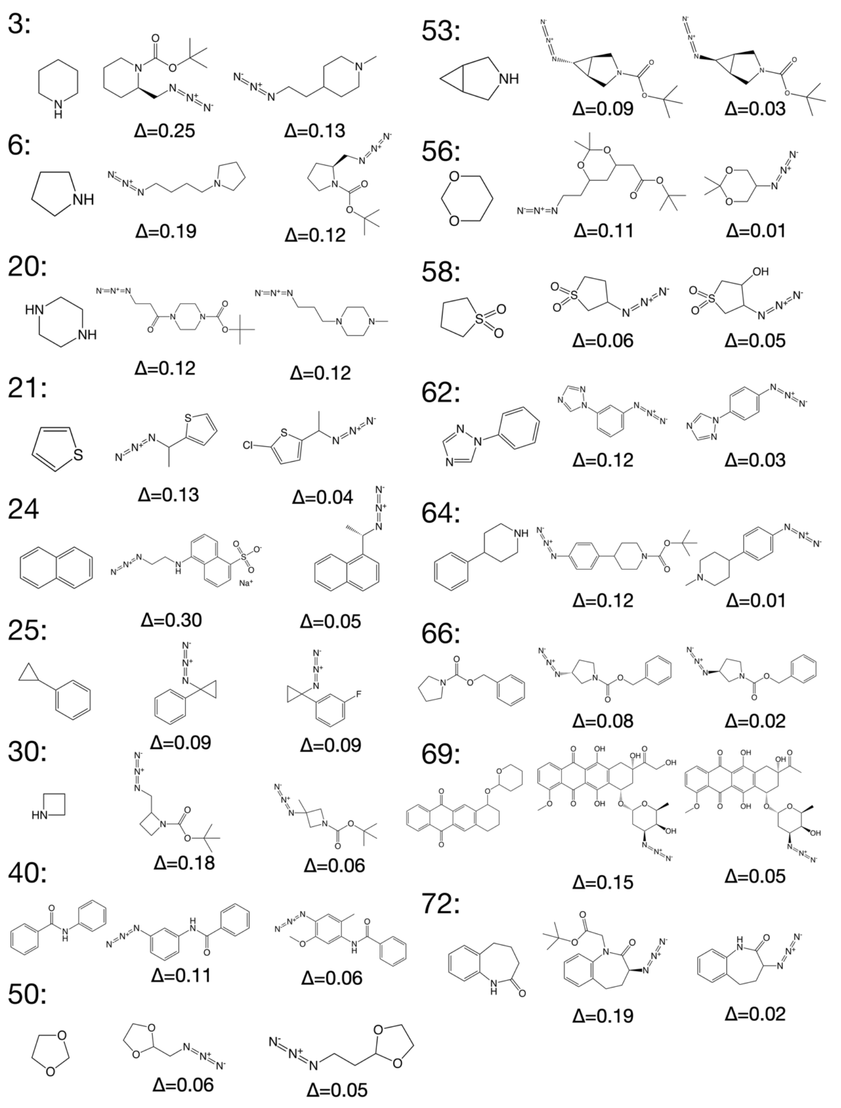


**Figure S18. Analysis of compounds that are more significantly impacted by TolC.** Two separate examples of compounds within each scaffold demonstrating large shifts in apparent accumulation related to TolC.

**Table S1.** Pearson R correlation between physicochemical properties and compound efflux.

| Properties | HBA | HBD | HBA+HBD | NumRings | RTB | Num Amide Bonds | Globularity |
|---|---|---|---|---|---|---|---|
| Pearson R<br>p-value | 0.17<br>$3x10^{-7}$ | -0.052<br>0.1 | 0.11<br>$8x10^{-4}$ | 0.041<br>0.2 | 0.12<br>$4x10^{-4}$ | 0.20<br>$2x10^{-9}$ | -0.27<br>$4x10^{-16}$ |

| Properties | PBF | TPSA | logP | MR | MW | Csp3 | fmf |
|---|---|---|---|---|---|---|---|
| Pearson R<br>p-value | 0.20<br>$2x10^{-9}$ | 0.15<br>$6x10^{-6}$ | 0.084<br>0.013 | 0.27<br>$5x10^{-16}$ | 0.28<br>$9x10^{-17}$ | 0.12<br>$4x10^{-4}$ | -0.055<br>0.01 |

| Properties | QED | HAC | Num Rings Fused | unique_HBAD | max_ring_size | N_chiral_centers | fcsp3_bm |
|---|---|---|---|---|---|---|---|
| Pearson R<br>p-value | -0.064<br>0.06 | 0.27<br>$3x10^{-16}$ | 0.041<br>0.2 | 0.16<br>$1x10^{-6}$ | 0.041<br>0.2 | 0.084<br>0.01 | 0.14<br>$2x10^{-5}$ |

*Physicochemical properties with Pearson R correlation value > 0.1 and p-value < 0.01 are highlighted in grey background color.*

**HBA**: hydrogen bond acceptor; **HBD**: hydrogen bond donor; **HBA+HBD**: sum of number of HBA and HBD; **NumRings**: number of rings; **RTB**: number of rotatable bonds; **NumAmideBonds**: number of amide bonds; **Globularity**: division of volume by surface area; **PBF**: plane of best-fit; **TPSA**: topological polar surface area; **logP:** partition between water and octanol; **MR**: molecular refractivity, a measure of polarizability; **MW**: molecular weight; **Csp3**: fraction of sp3 carbons; **fmf**: fraction of atoms belonging to Murcko framework; **QED**: quantitative estimate of drug-likeness; **HAC**: heavy atom count; **NumRingsFused**: number of rings fused; **unique_HBAD**: non-redundant hydrogen bond acceptors and donors; **max_ring_size**: the size of maximum ring; **n_chiral_centers**:

number of chiral centers; **fcsp3_bm**: fraction of sp3 carbon atoms in the Bemis-Murcko scaffold, a measure of saturation of scaffold carbon atoms.

**Biological methods.**

**Table S2.** Bacterial strains and plasmids used in this study

| **Strain and Plasmid** | **Relevant characteristics** | **Source or reference** |
|---|---|---|
| *E. coli* BW25113 | WT K12 Keio parent strain; *E. coli* background | Horizon Discovery Ltd |
| *E. coli* BW25113 *attTn7::mini Tn7T Gmr fhuAC/4L* | K12 Keio parent strain; constitutive expression of the FhuA pore (non-selective variant of FhuA); gentamicin 20 μg/ml) | Brown Lab, McMaster |
| *E. coli* BW25113 Δ*tolC::kanR* | K12 Δ*tolC*, *Kan*$^R$ (Keio collection, *tolC* deletion strain; kanamycin,50 μg/ml) | Horizon Discovery Ltd |
| *E*. coli BW25113 Δ*tolC::kanR attTn7::mini Tn7T Gmr fhuAC/4L* | K12 Δ*tolC*, *Kan*$^R$; constitutive expression of the FhuA pore (non-selective variant of FhuA) *Gent*$^R$ | Brown Lab, McMaster |
| *E. coli* ATCC25922 | Wild type, smooth, O-antigen | ATCC |
| pQE-30 | *T5* promoter, *Amp*$^R$ | Genscript |

**Plasmid construction for HaloTag expression in *E. coli*.** For expression of the HaloTag protein in WT *E. coli*, (smooth, ATCC25922) and in different variants of *E. coli* K12 (BW25113), HaloTag was cloned onto an expression vector pQE-30 (Genscript) and codon-optimized for expression in *E. coli*; the molecular cloning and DNA synthesis was performed by Genscript.

**HaloTag protein amino acid sequence.**
MASAEIGTGFPFDPHYVEVLGERMHYVDVGPRDGTPVLFLHGNPTSSYVWRNIIPHVA
PTHRCIAPDLIGMGKSDKPDLGYFFDDHVRFMDAFIEALGLEEVVLVIHDWGSALGFH
WAKRNPERVKGIAFMEFIRPIPTWDEWPEFARETFQAFRTTDVGRKLIIDQNVFIEGTL

PMGVVRPLTEVEMDHYREPFLNPVDREPLWRFPNELPIAGEPANIVALVEEYMDWLH
QSPVPKLLFWGTPGVLIPPAEAARLAKSLPNCKAVDIGPGLNLLQEDNPDLIGSEIARW
LSTLEISGGGGGSGGGGSGGGGSGS

**HaloTag protein expression and bacterial cell culture.** For expression of the HaloTag protein in WT *E. coli*, (smooth, ATCC25922) and in different variants of *E. coli* K12 (BW25113), the expression plasmid Halo_pQE-30 was transformed into *Escherichia coli* and grown on an LB/agar plate with ampicillin (100 μg/mL) at 37 ºC overnight. Colonies were picked and grown first in 5mL LB broth with ampicillin at 37 ºC overnight. A flask containing 35 mL of LB media was then inoculated with the overnight culture at a ratio of 1:100 in the presence of 100 μg/mL ampicillin and cells were grown at 37 ºC for 2 h, or until the optical density at 600 nm reached 0.2 when cultures were induced with 1 mM Isopropyl β-d-1-thiogalactopyranoside (IPTG) at 37 °C for 2 h in a shaker incubator. Cells were then pelleted at 3005*g* for 3 min. The pellet was washed three times with 1X PBS pH 7.4 following which cells were resuspended in 35 mL of 1X PBS for the assay. The expression and identity of the protein was confirmed by SDS-PAGE.

**One-step assay with rhodamine chloroalkane (R110cl).** 3 mL of HaloTag expressing cell culture was transferred to 15 mL conical tubes and pelleted by centrifuging. The cells were washed 3X with 3 mL of 1x PBS and finally re-suspended in 3 mL of 1x PBS. The cells were pipetted into wells in a 96-well plate in triplicate, pelleted down by centrifuging for 3 min at 2700*g* and resuspended using 5 μM R110cl. The cells were incubated for 30 min at 37°C, centrifuged for 3 min, washed 1X with PBS and fixed with 4% formaldehyde in PBS for 30 min statically at room temperature. The cells were subjected to analysis by flow cytometry on the Attune NxT Acoustic Focusing Cytometer (Invitrogen) by exciting using the blue laser with emission at 525 nm for acquisition.

**CHAMP assay development**

**Three-step assay: - DBCOcl incubation, azido-tagged small molecule (pulse), and Flaz dye (chase).** 35 mL of HaloTag expressing fresh cell culture was washed with PBS and transferred to 50 mL conical tubes and pelleted by centrifuging. The cells were re-suspended in 35 mL of a 50 μM solution of DBCOcl in PBS and transferred to an Erlenmeyer flask for incubation at 37 °C for 30 min in a shaker incubator. The DBCOcl imprinted cells were centrifuged to pellet at 3000*g* using HERAEUS Multicentrifuge X1 centrifuge (Thermofisher Scientific), washed 3X with PBS and resuspended in the same volume of PBS as the original cell suspension. 90 μL of this cell culture was added into each well of a pre-prepared 96 well plate containing 10 μL of the azido-tagged test compound from each 500 μM working solution in PBS. The plate was then covered with plastic film and incubated for 1 h at 37 °C in a shaker incubator. Each azido-tagged test molecule was replicated in three wells. Control experiment wells were either on the same plate or a separate plate in the same shaker incubator for 1 h at 37°C, and each well contained 10 μL of PBS with 90 μL of DBCO-labeled cell culture (both IPTG induced and non-induced cells), in triplicate. In the end, the plate was then spun at 2700*g* for 3 min using a plate holder centrifuge to pellet the cells, the supernatant was discarded, and the cells were washed 1X with PBS. Next, cells in both the experimental wells and the control wells were fixed with 4% formaldehyde in PBS for 30 min and incubated without shaking at room temperature on the benchtop. Following formaldehyde fixation, the cells were centrifuged and washed 1X with PBS. In the last chase step, 100 μL of 50 μM Flaz was added to the washed cells in each well, resuspended by pipetting up and down, and incubated at 37°C for 60 min in a shaker incubator. Finally, the cells in the plate were pelleted by centrifuging in a plate centrifuge, washed 1X with PBS, resuspended in 200 μL of PBS, and subjected to analysis by flow cytometry.

**Two-step assay without fixation prior to chase step: - DBCOcl incubation, azido-tagged dye.** DBCO labeled cells were prepared as described earlier and resuspended in the same volume of PBS as the original cell suspension. 90 μL of the cell culture was added into each well containing 10 μL of the azido-tagged dye at 500 μM in 96-well plates, covered with plastic film, and incubated for 1 h at 37 °C in a shaker incubator. Each azido-tagged molecule or dye was replicated in three wells. The plate was then spun at 2700*g* for 3 min using a plate holder centrifuge, the supernatant was discarded, and the cells were washed 1X with PBS. Next, cells were fixed with 4% formaldehyde in PBS for

30 min statically at room temperature on the benchtop. Following formaldehyde fixation, the cells were washed 1X with PBS and resuspended in 200 μL of PBS for analysis by flow cytometry.

**Two-step assay with fixation prior to chase step: - DBCOcl incubation, formaldehyde fixation, and azido-tagged dye.** DBCO labeled cells prepared as described earlier were washed 3X with PBS and resuspended in 4% formaldehyde in PBS and incubated statically at room temperature on the benchtop for 30 min. Following formaldehyde fixation, the cells were washed 1X with PBS and resuspended in the same volume of PBS as was in the original cell suspension. For the azido-tagged dyes screen, 90 μL of the cell culture was added into wells containing 10 μL of the azido-tagged dye at 500 μM in 96-well plates, covered with plastic film, and incubated for 1 h at 37 °C in a shaker incubator. Each azido-tagged dye test was replicated in three wells. The plate was then spun at 2700*g* for 3 min, the supernatant was discarded, and the cells were resuspended in 200 μL of PBS for analysis by flow cytometry.

**Assay with PMBN: - DBCOcl incubation, azido-tagged dye coincubation with PMBN** DBCO labeled cells were prepared as described earlier and washed 3X with PBS and resuspended in PBS containing 5 μM PMBN. For the azido-tagged small molecule (three-step) or azido-tagged dyes screen, 90 μL of the cell culture was added into each well containing 10 μL of the azido-tagged small molecule/dye at 500 μM in 96-well plates, covered with plastic film, and incubated for 1 h at 37 °C in a shaker incubator. Each azido-tagged molecule or dye was replicated in three wells. The plate was then spun at 2700*g* for 3 min using a plate holder centrifuge, the supernatant was discarded, and the cells were washed 1X with PBS for fixing, as described earlier, before analysis by flow cytometry (two-step), or for the chase step with an azido-tagged dye (three-step), as described earlier.

**1152 azido-tagged test molecules library screen**

The 1152-molecule azido-tagged library in the form of 96-well mother plates (total 12 plates) was obtained from the Dong Lab at the Institute of Translational Medicine, Zhangjiang Institute for Advanced Study (Shanghai Jiao Tong University) and stored at -

20 °C. Each well contained 50 μL of individual test molecules at 5 mM in DMSO. Prior to the assay, the original 96-well plates (denoted as the "mother plates") were allowed to warm up to room temperature and spun down for 1 min at 2700*g,* to allow the stock solution to settle at the bottom of each well, using a plate holder centrifuge. 10 μL of the DMSO stock was drawn from each well and dispensed to a new set of plates (denoted as the "daughter plates"). Then 90 μL PBS was added to the daughter plates to make a 100 μL solution of each test molecule at 500 μM working concentration. 10 μL of each test molecule from the daughter plates was transferred to a new 96-well plate (the "working plates").

DBCO labeled cells were prepared as described earlier and poured into a reservoir. 90μL of DBCO-labeled cell suspension was drawn from the reservoir and dispensed to the working plates prepared above using a 96-multichannel pipette dispenser (Sequence®). The plates were covered with plastic films and incubated for 1 h at 37 °C in a shaker incubator. The plates were then spun at 2700*g* for 3 min using a plate holder centrifuge. The supernatant was discarded, and the cells were washed 1X with PBS. Next, cells were fixed with 4% formaldehyde in PBS for 30 min statically at room temperature on the benchtop. Following formaldehyde fixation, the cells were washed 1X with PBS. For the final chase step, 100 μL of 50 μM Flaz was added to each well, resuspended and incubated at 37°C for 1 h in a shaker incubator. Lastly, the cells were pelleted by centrifuging the 96-well plate in a plate holder centrifuge, washed 1X with PBS, resuspended in 200 μL of PBS, and analyzed by flow cytometry.

**Cell preparation for confocal microscopy**

DBCO labeled cells prepared as described earlier were washed 3X with PBS and resuspended in the same volume of PBS as was drawn from the original cell suspension. For the azido-tagged small molecules screen, 90 μL of the cell culture was added into wells containing 10 μL of the azido-tagged test compound at 500 μM in 96-well plates, covered with parafilm, and incubated for 1 h at 37 °C in a shaker incubator. Each azido-tagged dye test was replicated in three wells. Control experiment wells were either on the same plate or a separate plate in the same incubator for 1 h at 37 °C, and each well contained 10 µL of PBS with 90 µL of DBCO-labeled cell culture (both IPTG induced and non-induced cells), in triplicate. The plate was then spun at 2700*g* for 3 min in a plate holder centrifuge, the supernatant was discarded, and the cells were washed 1X with 1X PBS.

Next, cells in both the experimental wells and the control wells were fixed with 4% formaldehyde in 1X PBS for 30min statically incubated at room temperature on the benchtop. Following formaldehyde fixation, the cells were washed 1X with PBS and for the chase step, 100 µL of 50 µM Flaz was added into each well, resuspended by pipetting up and down, and incubated at 37°C for 60 min in a shaker incubator. Lastly, the cells were pelleted by centrifuging the 96-well plate in a plate centrifuge, washed 1X with PBS, resuspended in 200 µL of PBS. For confocal microscopy, the fixed cells were treated with 5 µg/mL of TAMRA-tagged wheat germ agglutinin (Vector Laboratories, RL-1022) for 30 min at 4 °C. Glass microscope slides were spotted with a 1% agar pad, and 5 µL of the sample were deposited onto the agar. Samples were covered with a glass coverslip and imaged using a Zeiss 880/990 multiphoton Airyscan microscopy system (40× oil-immersion lens) equipped with 488 nm and 550 nm lasers. Images were obtained and analyzed via Zeiss Zen software. We acknowledge the Keck Center for cellular imaging and the usage of the Zeiss 880/980 multiphoton Airyscan microscopy system (PI-AP: NIH-OD025156).

**Confocal fluorescence images of *E. coli* cells expressing HaloTag, labeled with D-Lys(FITC) and treated with coumarin chloroalkane to produce a blue cytosolic signal.** *E. coli* cells harboring the plasmid pQE-30_HaloTag (Amp) were cultured overnight in LB medium. The overnight culture was diluted 1:100 into fresh LB medium supplemented with 500 µM D-Lys(FITC) and incubated at 37 °C with shaking until the optical density at 600 nm ($OD_{600}$) reached approximately 1.2 to allow the incorporation of D-Lys(FITC) in the cell wall. Protein expression was induced with 1 mM isopropyl β-D-1-thiogalactopyranoside (IPTG), and cells were incubated for 1 h to allow expression of the

HaloTag protein. Uninduced cultures were maintained as negative controls. Next, cells were washed three times with phosphate-buffered saline (PBS) and incubated with 50 μM coumarin chloroalkane (Comcl) for 30 min at 37 °C with shaking. Cells were subsequently washed three times with PBS, fixed with 4% (w/v) formaldehyde for 30 min, washed again and mounted on 1% (w/v) agarose pads for imaging by confocal microscopy. Confocal fluorescence imaging was performed using a Leica STELLARIS 8 Super-Resolution Microscope equipped with a 100× oil-immersion objective. Image acquisition and analysis were carried out using ImageJ software. Excitation was performed using 405nm (Coumarin) and 488 nm (FITC) laser lines. We acknowledge the Keck Center for cellular imaging (PI-AP: NIH-OD025156).

**Confocal fluorescence images of *E. coli* cells expressing HaloTag, labeled with D-Lys(FITC) and treated with DBCOcl, which was subsequently reacted with Comaz to generate a blue cytosolic signal.** *E. coli* cells harboring the plasmid pQE-30_HaloTag (Amp) were cultured overnight in LB medium. The overnight culture was diluted 1:100 into fresh LB medium supplemented with 500 μM D-Lys (FITC) and incubated at 37 °C with shaking until the optical density at 600 nm ($OD_{600}$) reached approximately 1.2 to allow the incorporation of D-Lys (FITC) in the cell wall. Protein expression was induced with 1 mM isopropyl β-D-1-thiogalactopyranoside (IPTG), and cells were incubated for 1 h to allow expression of the HaloTag protein. Uninduced cultures were maintained as negative controls. Next, cells were washed three times with phosphate-buffered saline (PBS) and incubated with 50 μM DBCOcl for 30 min at 37 °C with shaking. Cells were subsequently washed three times with PBS, fixed with 4% (w/v) formaldehyde for 30 min, washed again, and incubated with 50 μM Comaz for 1 h at 37 °C with shaking. Finally, cells were washed three times and mounted on 1% (w/v) agarose pads for imaging by confocal microscopy. Confocal fluorescence imaging was performed using a Leica STELLARIS 8 Super-Resolution Microscope equipped with a 100× oil-immersion objective. Image acquisition and analysis were carried out using ImageJ software. Excitation was performed using 405nm (Coumarin) and 488 nm (FITC) laser lines. We acknowledge the Keck Center for cellular imaging (PI-AP: NIH-OD025156).

**Novobiocin azide (novAZ1) accumulation assay in WT *E. coli* BW25113 and Δ*tolC* *E. coli* BW25113 with or without efflux inhibitor PAβN.** DBCO labeled cells for the accumulation assay were prepared as described earlier. For each strain of *E. coli*, cells were divided in two equal groups with one set containing 25 μL of 200μM novAZ1 and 25 μL of PBS as a control; and the other set containing 25 μL of 200 μM novAZ1 and 25 μL of 64 μg/mL PAβN. Next, 50 μL of DBCO-labeled WT *E. coli* cell suspension or Δ*tolC E. coli* suspension was added to the wells in volumes needed to bring the final concentration of novAZ1 and PAβN to 50 μM and 16 μg/mL respectively. These cells were incubated for 1 h, then pelleted by centrifugation, and the supernatant was removed. Cells were washed 3X in PBS and resuspended for the chase step with R110az as described earlier. Finally, the cells were centrifuged, washed 1X with PBS, fixed with 4% formaldehyde statically at room temperature, and were subjected to flow cytometry analysis. A similar method was followed for all azido-tagged antibiotics used in this study.

#### Synthesis and characterization of azido-tagged antibiotics and test molecules used in this study

##### General information about materials and instruments:

Chemicals, reagents, and solvents used for the synthesis were purchased from standard sources such a Sigma-Aldrich (St. Louis, MO, USA), Fisher Scientific (Hampton, NH, USA), VWR (Randor, PA, USA), Chem-Impex (Wood Dale, IL, USA) and Alfa Aesar (Ward Hill, MA, USA). Most antibiotics precursors used in this study were procured from AK Scientific (Union City, CA, USA). Linezolid was purchased from Toronto Research Chemicals (Cat #: D195600) and Novobiocin sodium 90% (HPLC) was purchased from AK Scientific, Inc. 2-(2-((6-Chlorohexyl)oxy)ethoxy)ethanamine hydrochloride was purchased from AmBeed, DBCO-NHS (Cat.#: BP-22231) was purchased from BroadPharm. Rhodamine azide (R110 azide, 6- isomer, R110az), fluorescein azide (FAM azide, 6-isomer, Flaz), sulfo-Cyanine3 azide (Cy3az), and sulfo-Cyanine5 azide (Cy5az) were all purchased from Lumiprobe®. Rhodamine chloroalkane (R110cl) and coumarin chloroalkane were purchased from Promega® with trade name HaloTag(R) R110 Direct (TM) Ligand and HaloTag® Coumarin Ligand respectively. Coumarin-azide (Comaz) and FITC-labeled D-lysine were purchased from Sigma Aldrich. The 24-member library of azido-tagged molecules was purchased from Enamine and Sigma. These compounds were used as is after acquisition, purity of compounds was >95%. The stock solutions of these compounds were made in DMSO at 50 mM and stored at -20 $^{o}$C. The working solutions of these compounds (500 μM) in PBS were made freshly from the mother stock prior to assays.

The 1152-member azido-tagged library was obtained from Dong Lab at the Institute of Translational Medicine, Zhangjiang Institute for Advanced Study (Shanghai Jiao Tong University).

All reagents and chemicals for synthesis were purchased and used without any further purification. Organic solvents were removed under vacuo using rotary evaporator. Thin layer chromatography analyses for monitoring reaction progress were performed on aluminum backed thin layer silica gel plates (Merck F254); plates were developed in appropriate solvents mentioned in each individual reaction and were observed under UV lamp. Column chromatography on silica gel (Supelco, 60 Å, 230-400 mesh, 40-63 μ particle size) was performed on a manually packed column with selected solvents for each individual sample.

Preparative reverse phase HPLC purification was performed on instruments equipped with Waters 1525 pumps and 2489 UV/Visible Detector on a Phenomenex Luna 10 µm C8(2) 100 Å (250 x 21.2 mm) or C18 columns using a 5 to 100% linear gradient of methanol in $H_2O$ or MeCN in water each containing 0.1% TFA at 10 mL/min. The HPLC fractions of the desired compounds were first concentrated under reduced pressure using a rotary evaporator. The concentrated aqueous solutions were frozen prior to lyophilization with Labconco Freezone 4.5 L lyophilizer (-84 $^{o}$C). The purity of the samples was ascertained by either TLC or analytical HPLC using a Phenomenex Luna 5 µm C8(2) 100 Å (250 x 4.6 mm) on the same instrument; using gradient elution in $H_2O$/MeCN or $H_2O$/MeOH with 0.01% TFA in each solvent at 1 mL/min prior to its use in biological assays. Some of the compounds having amino function isolated from HPLC may have been in trifluoroacetate salt form, and these were not neutralized prior to characterization by methods described below and were used as-is for bioassays.

$^1H$ and $^{13}C$-NMR spectra for final compounds and intermediates were acquired on a Varian 600 MHz spectrophotometer. All NMR spectra were processed and analyzed using MestreNova software. Residual solvent signal from $CDCl_3$, $CD_3CN$, $CD_3OD$ and DMSO-$d_6$ referenced to tetramethylsilane (TMS) were used as reference standards for defining chemical shifts. Chemical shifts are reported in ppm and coupling constants (J) are reported in Hertz [Hz]. Deuterated solvents were used as received from Cambridge Isotopes. Compounds isolated from HPLC and dried using the lyophilizer still showed presence of water in $^1H$-NMR analysis. This water may have been present in NMR solvents used or the compounds were hygroscopic and acquired moisture from atmosphere prior to obtaining $^1H$-NMR. Routine mass analysis ESI was performed on Advion Expression® CMS mass spectrometer using standard parameters for intermediates. For the analysis of fragmentation sensitive compounds, low fragmentation, low energy setup was used. MALDI-TOF mass spectra were obtained for certain high molecular weight compounds on the Shimadzu MALDI-8020 instrument with α-Cyano-4-hydroxycinnamic acid (α-CHCA) matrix. The observed molecular weights for compounds were represented as m/z. Additional LCMS analysis, which were performed on Shimadzu Prominence-i LCMS system equipped with LC-2030C 3D liquid chromatograph, autosampler, PDA and LCMS-2020 mass detector. Data analysis was performed with Lab Solutions software. High resolution electrospray ionization mass spectrometry (HRMS, ESI/MS) analyses were obtained in positive ion mode on an Agilent 6545B Q-TOF LC/MS equipped with 1260 Infinity II LC system with auto sampler and equipped with C18(2) column (Luna 5 µm 100Å 250 x 4.6 mm). Quantitative High resolution electrospray ionization mass spectrometry (HRMS, ESI/MS) analyses were obtained on an Agilent 6545B Q-TOF LC/MS equipped with 1260 Infinity II LC system with auto sampler and equipped with C18(2) column (Luna 5 µm 100Å 250 x 4.6 mm. Quantitative LCMS analysis was performed on Agilent 6545B Q-TOF as describe above in positive ion mode. For each antibiotics-azide tested, parameters were such as collision energy and fragmentation pattern (formation of defined daughter ions and their intensities were optimized). Prior to actual quantification of recovered antibiotics-azides from cell lysates, standard curves at lowest possible concentration were obtained. The LCMS/MS data was processed with Mass Hunter software, and subsequent quantification was done with Q-TOF MS quantitative software. UV spectroscopic analysis was performed on Genesys 50 (Thermo Scientific) UV-Visible Spectrophotometer.

Azide-antibiotics reported in the current studies such **vanAZ1**, **rifAZ1**, **triAZ1**, **metAZ1**, **sulAZ1**, **moxAZ1**, **purAZ1**, **novAZ1**, **zolAZ1, zolAZ2,** and **zolAZ3** are either reported by us or by others in the literature and were synthesized according to the known procedures and characterized thoroughly; purity was determined by HPLC analysis wherever possible prior to their use.

**Scheme 1: Synthesis of DBCO chloroalkane (DBCOcl).**

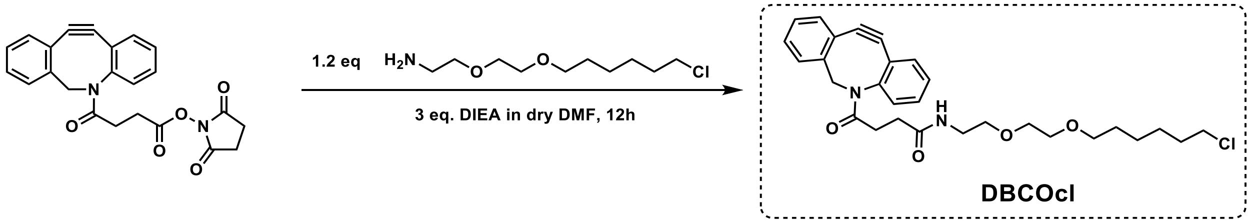


Following a published protocol, 41 mg of DBCO-NHS (0.1 mmol, BroadPharm) in dry DMF was mixed with 32 mg of 2-(2-((6-Chlorohexyl)oxy)ethoxy)ethanamine hydrochloride (0.12 mmol, AmBeed) into which was added 43 µL of DIPEA (0.24mmol).[1] The reaction mixture was stirred at room temperature for 1hr following which, DMF was removed by rotary evaporation of a DMF/toluene mixture. The crude residue was acidified and purified by RP-HPLC (Mobile phase A: 90% v/v water initial to 0% v/v water with 0.1% TFA; mobile phase B: 10% v/v MeCN -100% v/v MeCN with 0.1% over a 70min gradient) to afford 35 mg of an oily product (67% yield) which was found to be DBCOcl, the analytical data was in agreement with assigned structure. Calc.: $C_{29}H_{36}ClN_2O_4$ $[M+H]^+$ 511.2358; Obs. 511.2.

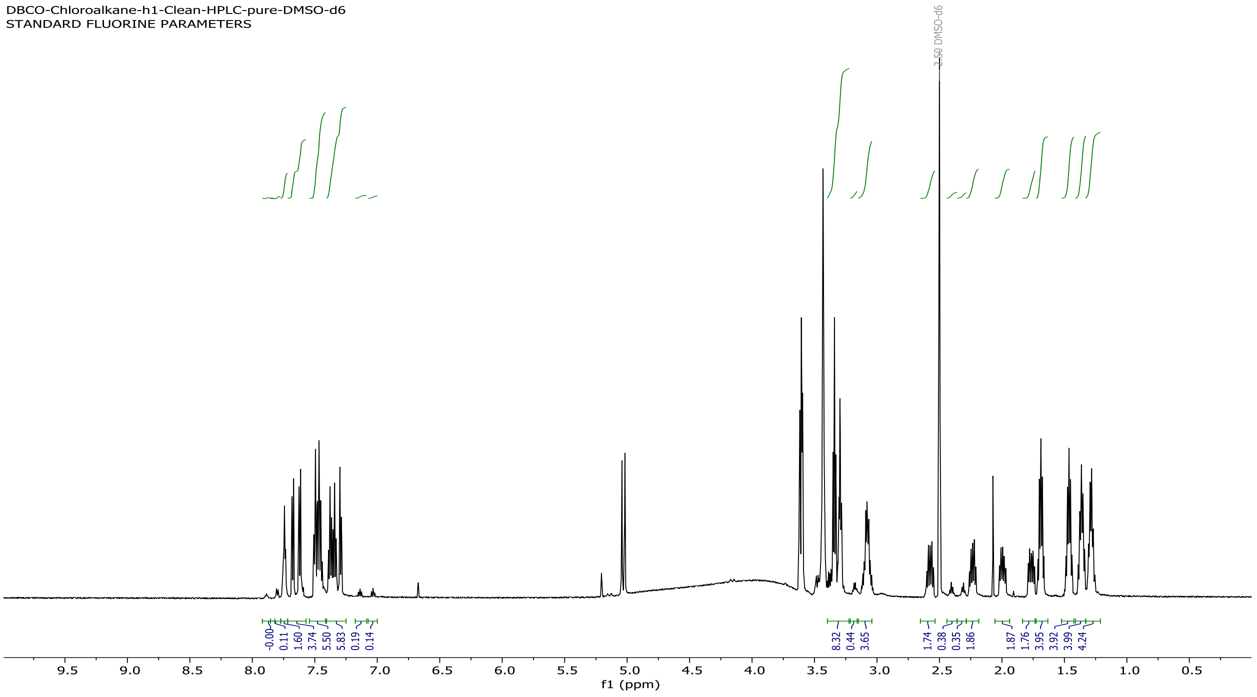


[1]NMR spectrum of DBCOcl.

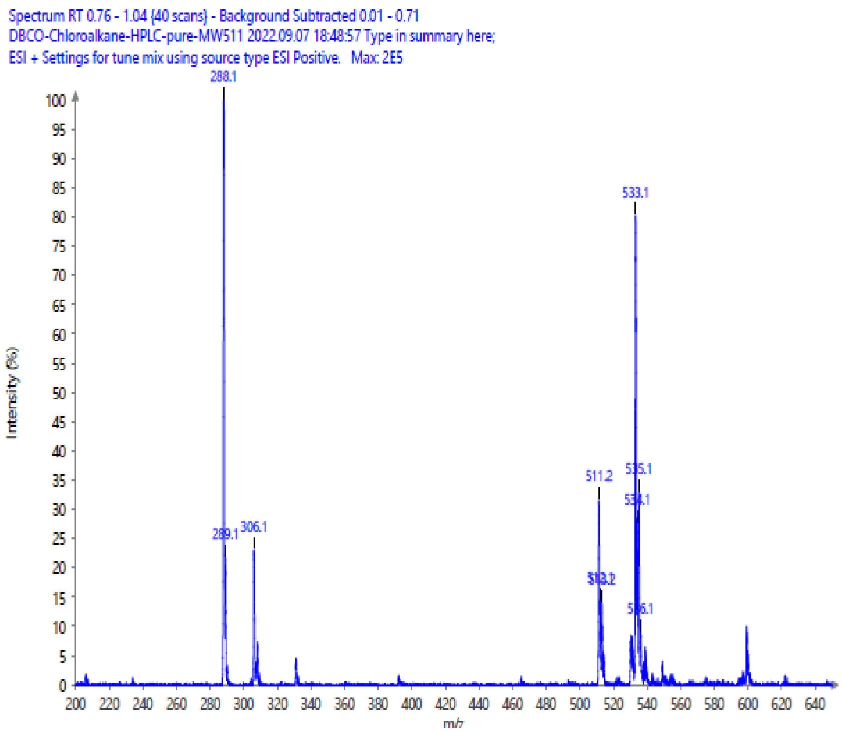


ESI-MS of DBCOcl.

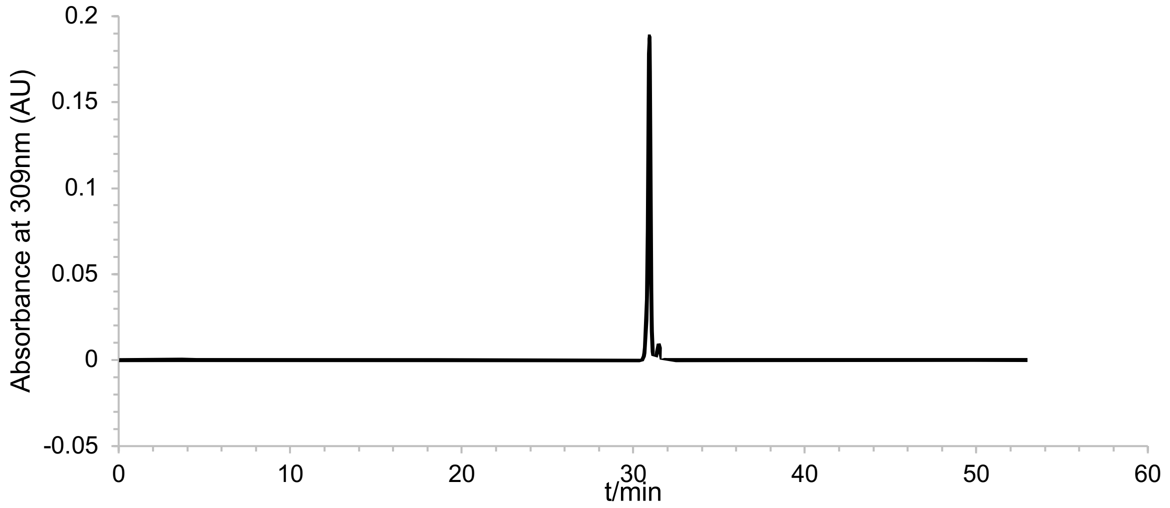


DBCOcl purity check analytical RP-HPLC detection at 309 nm.

#### Synthesis of vanAZ1.

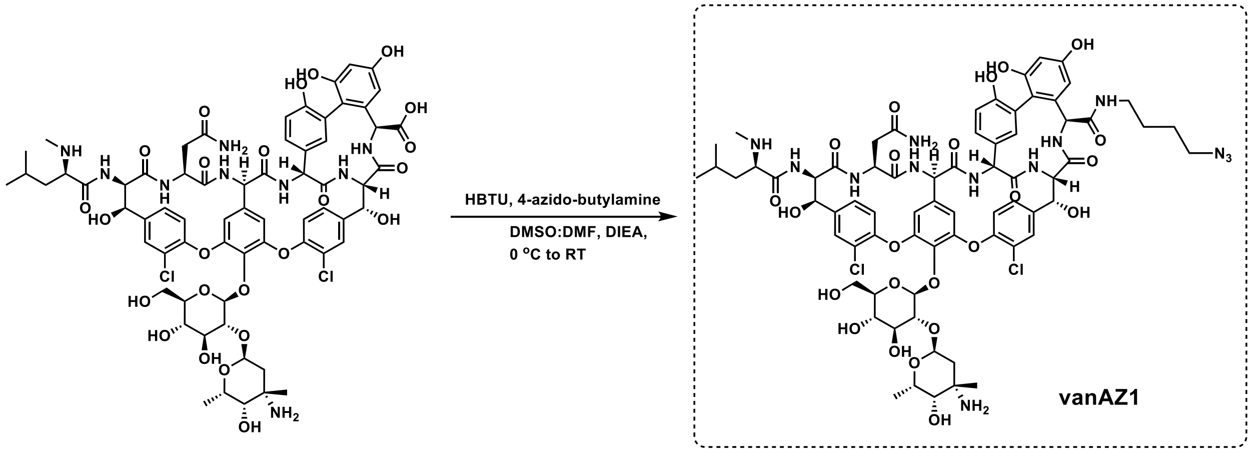


Following reported procedure[2] vancomycin-HCl (MW: 1485, 51.0 mg, 0.0343 mmol, Chem-Impex, Cat # 00315, Lot# 001204-1802003) was dissolved in 1:1 DMSO:DMF, 2 mL and to it was added 4-azido-butylamine (10.0 mL solution in $CH_2Cl_2$ of 1:1 conc, MW: 114.15, 5.0 mg, 0.043 mmol), the solution was cooled in an ice water bath and stirred. While stirring, a solution of HBTU (15.2 mg, 0.04 mmol), followed by DIEA (50 mL) was added. The ice bath was allowed to warm up to room temperature, and the mixture was stirred for 6 h. The crude reaction mixture was subjected to analytical HPLC and this analysis showed that the starting material was completely consumed ($R_t$ ~ 12 min) to form a new nonpolar product with Rt ~18 min. The reaction mixture was then diluted with Millipore DD water, 10.0 mL, and the mixture was filtered through 0.4 µ filter. The clear solution was injected to reverse phase preparative HPLC under the same conditions (80:20; water: methanol containing 0.1%TFA) to isolate single peak fractions. The fractions were analyzed by MALDI-TOF prior to combining; fractions showing the desired mass were combined and evaporated under reduced pressure with a rotary evaporator. The leftover solution was frozen in -80 °C freezer and lyophilized for 24 hrs to dry solid (37.5 mg, yield 79%), characterized by $^1$H-NMR and MALDI-TOF mass spectrometry. Calculated m/z 1543.5102 for $C_{70}H_{84}Cl_2N_{13}O_{23}$ $[M+H]^+$; MALDI-TOF observed m/z 1544.985.

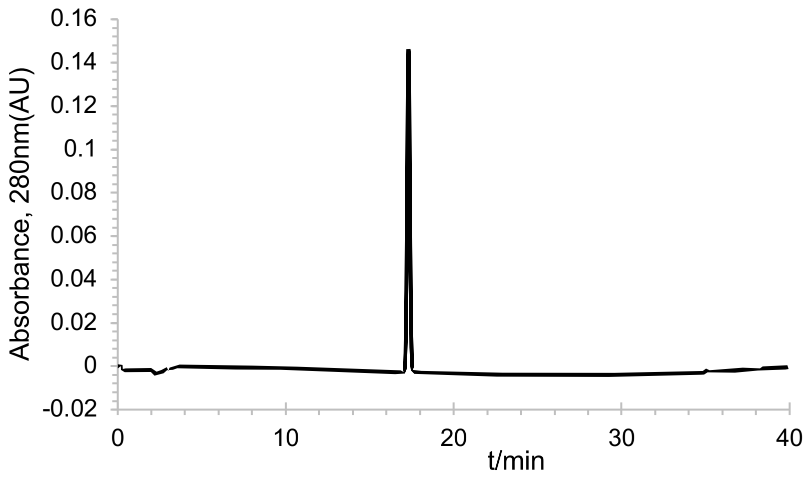


Analytical HPLC chromatogram of **vanAZ1**.

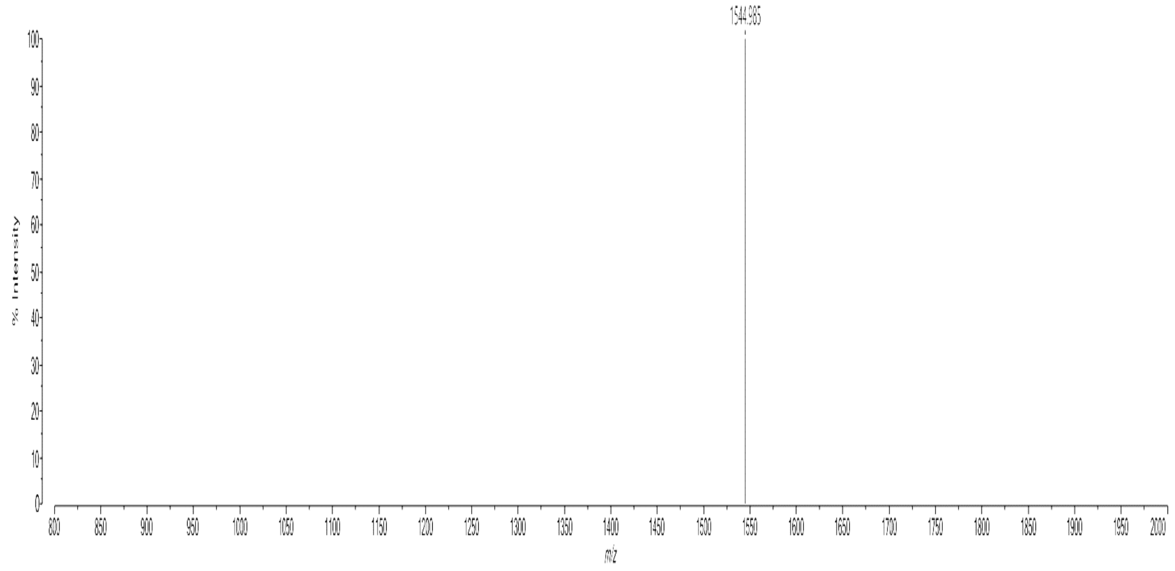


MALDI-TOF spectrum of **vanAZ1**.

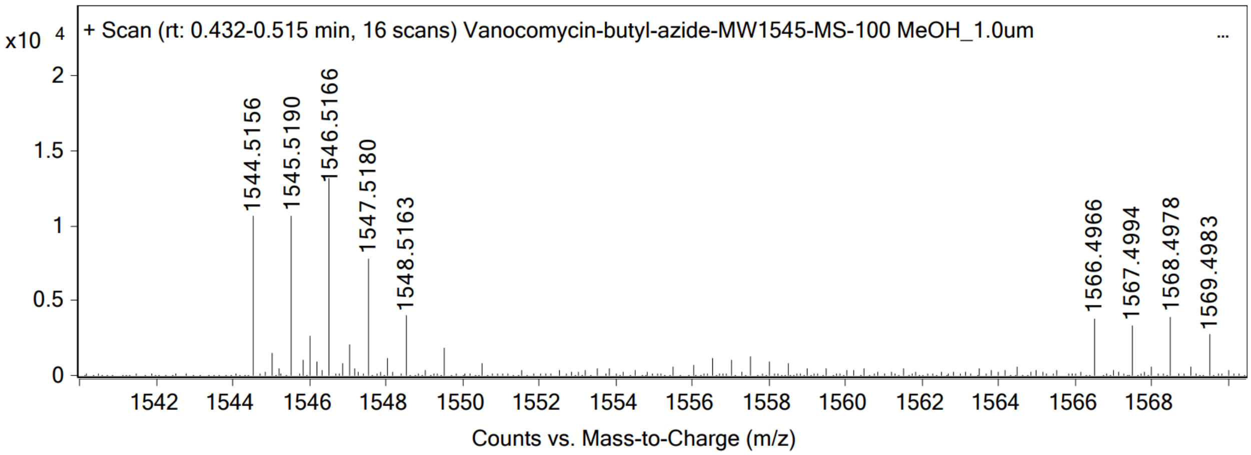


High resolution mass spectrum (ESI) of **vanAZ1**.

**Synthesis of rifAZ1.**

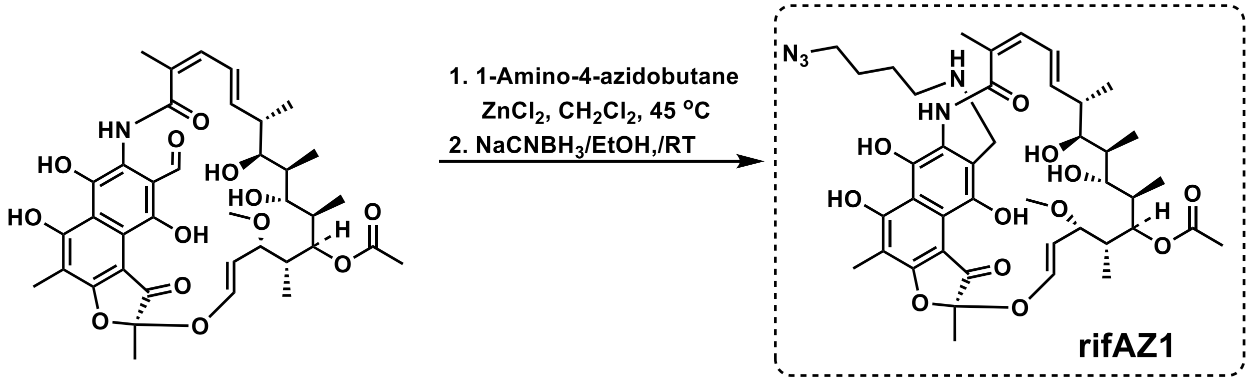


Rifamycin aldehyde (72.5 mg. 0.1 mmol) was dissolved in 5.0mL of anhydrous methylene chloride in 50 mL RB flask equipped with reflux condenser.[3] To this solution was added 1-amino-4-azidobutane (17.1 mg, 0.15 mmol), followed by $ZnCl_2$ (11.0 mg, at room temperature and the mixture was refluxed at 45 °C in an oil bath for 45 minutes. The mixture was then allowed to cool to room temperature. The solvents were evaporated, and the crude material was used as-is for reduction of iminium. The crude material was dissolved in ethanol (200% proof, 2.5 mL), sodium cyanoborohydride (10.7 mg, 0.17 mmol) was added to it and the mixture was stirred at room temperature for 2 h. TLC analysis of the crude material indicated complete conversion of the starting material ($R_t$ ~ 0.4) to a polar new compound (Rt ~0.2) in EtOAc:methanol (95:5) . The volatiles were removed under vacuo using a rotary evaporator. The remaining residue was dissolved in EtOAc and purified by column chromatography over silica gel using gradient ranging from ethyl acetate to 3% methanol EtOAc. Fractions showing homogeneity on TLC were combined and concentrated under reduced pressure to yield a red solid (65.1 mg, 79%). Further purification with reverse phase HPLC (80:20; Water:Methanol) afforded pure a compound that was used for the biological assay. The product was characterized by $^1$H-NMR and MALDI-TOF MS. Mass: m/z 824.4076 for $C_{42}H_{58}N_5O_{12}$ $[M+H]^+$ ; ESI MS observed m/z 824.4329.

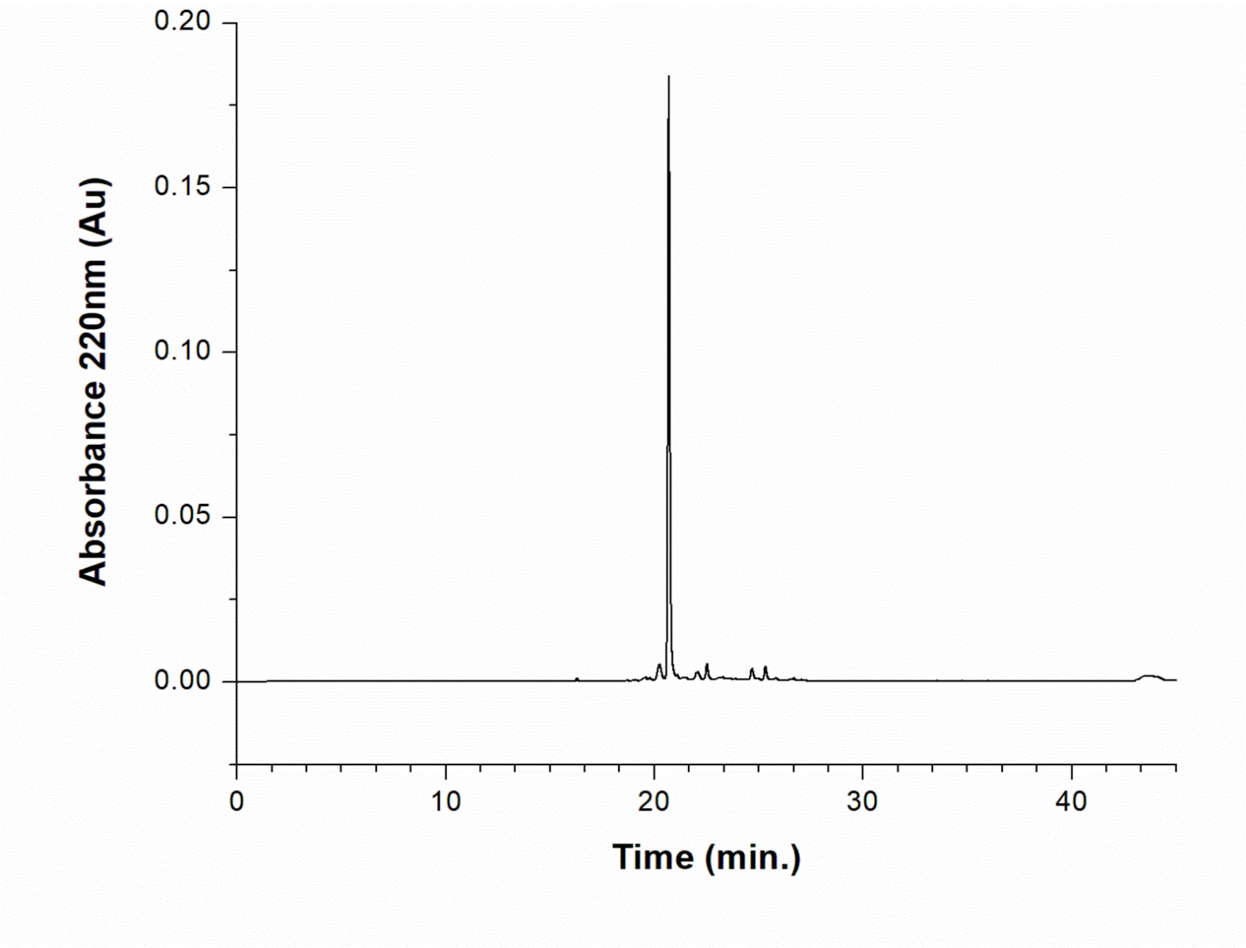


Analytical HPLC chromatogram of **rifAZ1**.

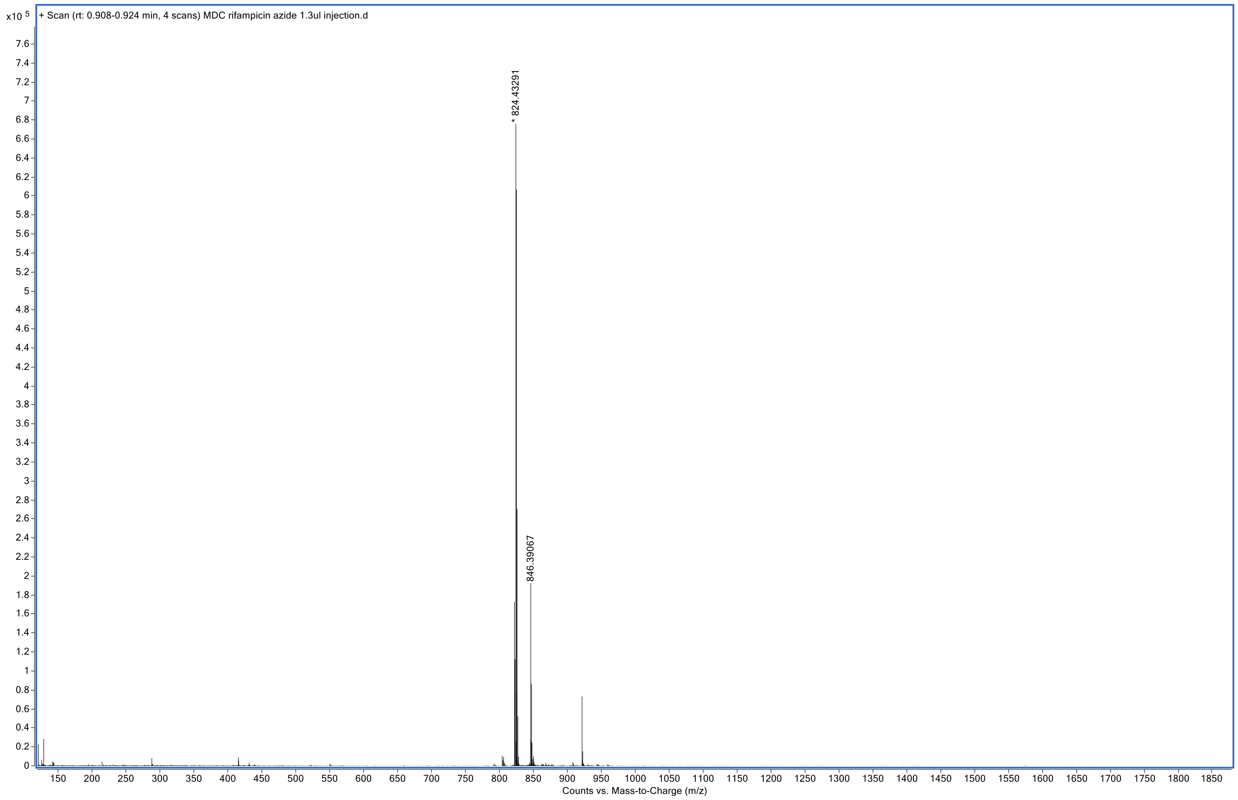


High resolution mass spectrum (ESI) of **rifAZ1**.

**Synthesis of triAZ1.**

Following a published protocol, **triAZ1** was synthesized and characterized as follows:[3]

NMR $^1$H (600 MHz, $CD_3OD$): δ 7.22 (s, 1H), 6.56 (s, 2H), 8.06 (d, J=6 Hz, 1H), 3.99 (t, J=6 Hz, 2H), 3.84 (s, 3H), 3.66 (s, 1H), 3.73 (dd, J= 3, 6 Hz, 1H), 3.58 (t, J= 3, 6 Hz, 1H), 1.91 (m, 2H). $^{13}$C NMR (151 MHz, $CDCl_3$) δ δ 166.2, 156.1, 155.0, 140.3, 137.0, 133.7, 111.0, 107.1, 71.0, 56.6, 33.0, 30.6. MS (ESI): m/z 360.5 for $C_{16}H_{22}N_7O_3$ $[M + H]^+$.

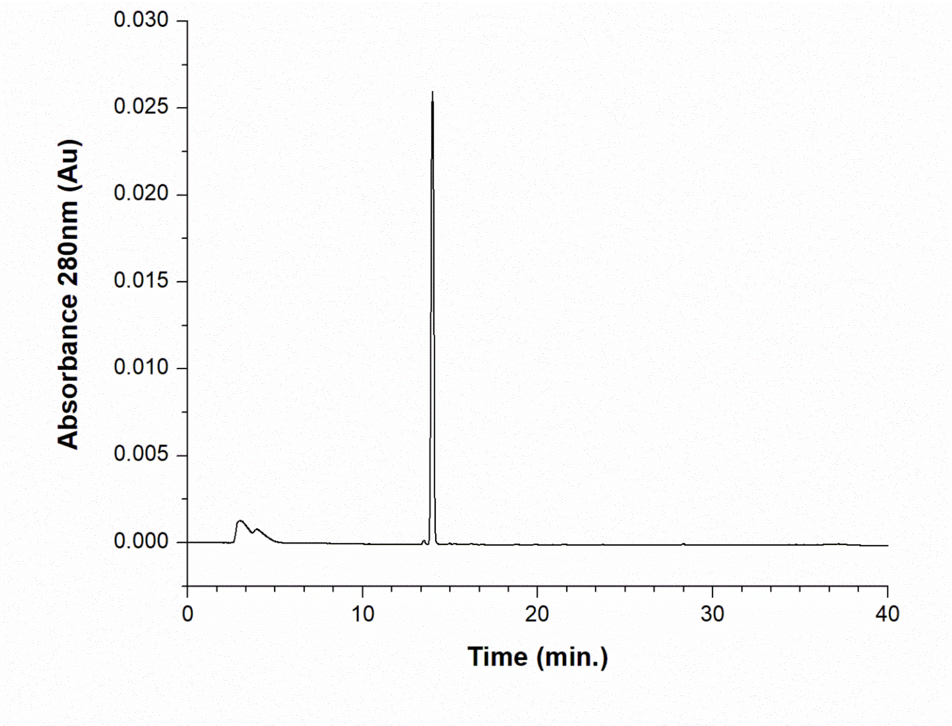


Analytical HPLC chromatogram of **triAZ1**.

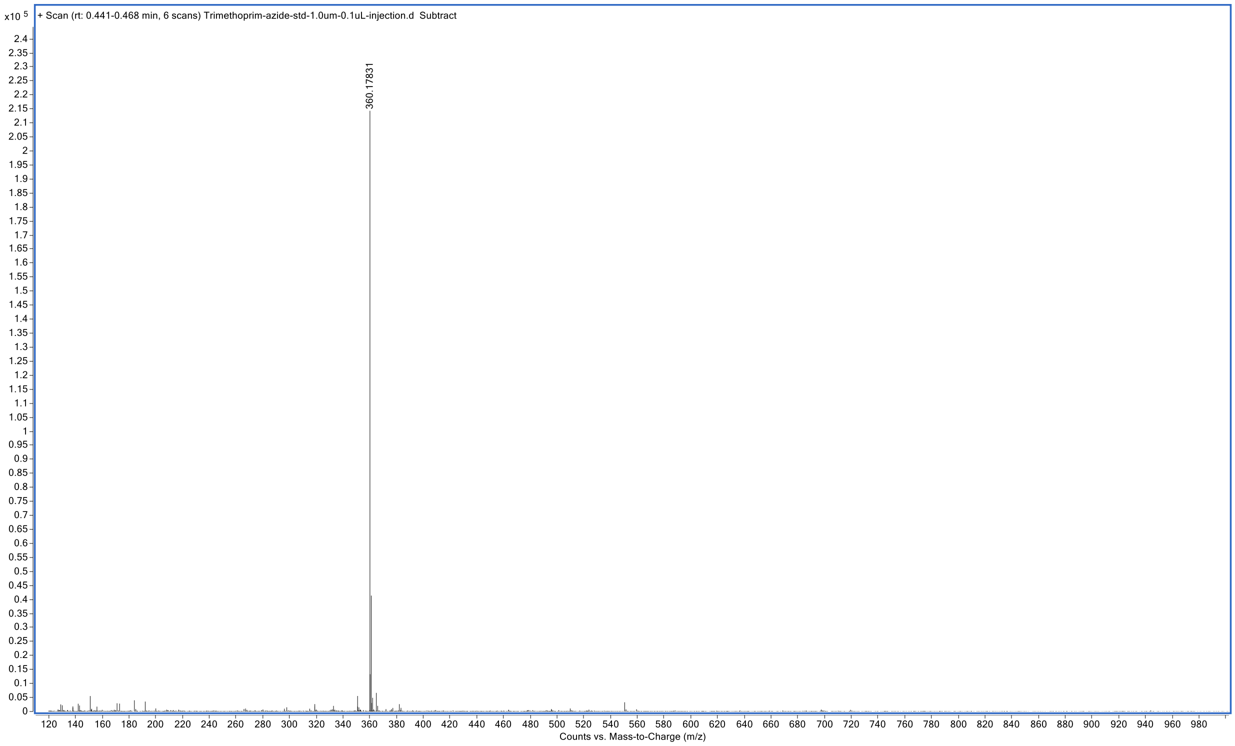


High resolution mass spectrum (ESI) of **triAZ1**.

**Synthesis of moxAZ1**.

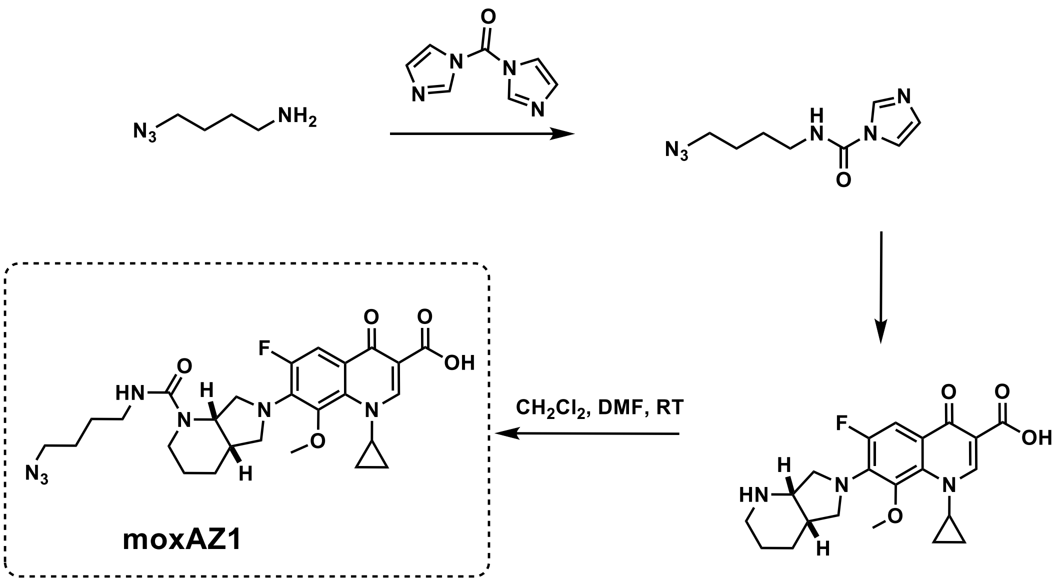


First 4-azidobutyl amine (114 mg, 1 mmol) in anhydrous $CH_2Cl_2$ (3 mL) was mixed with CDI (165.0 mg, 1 mmol). The mixture was stirred at room temperature overnight. The next day, 100 µL of the homogenous solution from this reaction (~0.33 mmol) was added to the clear solution of moxifloxacin (137 mg, 0.33 mol) in DMF and DIEA (50 µL). The mixture was further stirred at room temperature for 6 h. The reaction mixture was analyzed by TLC and ESI-mass for the formation of the product. At the end of the stirring, the reaction mixture was diluted with $CH_2Cl_2$ (20 mL), acidified with citric acid and washed with water (10 mL x 3), and finally with brine. The organic layer was separated and dried over $Na_2SO_4$ and was concentrated under reduced pressure using rotary evaporator. The residue was dissolved in a minimum amount of $CH_2Cl_2$ and loaded on a silica gel column for purification. Using initial $CH_2Cl_2$ and then $CH_2Cl_2$:MeOH (98:2), the desired compound was eluted. The TLC homogenous fractions were combined and concentrated under reduced pressure (109 mg, 61%). The compound was analyzed by analytical HPLC, characterized by high resolution mass spectroscopy and $^1$H-NMR as follows: HRMS observed m/z 542.2527; expected m/z 542.2522 for: $C_{26}H_{32}FN_7O_5$ $[M+H]^+$. $^1$H NMR (600 MHz, $CDCl_3$) δ 0.85 (m, 1H), 1.06-1.15 (m, 2H), 1.29 (m, 1H), 1.54 (m, 1H), 1.62 (m, 1H), 1.64-1.87 (m, 6H), 2.32 (m, 1H), 3.02 (d (m, 1H)t, J=2 and 6 Hz, 1H) 3.26-3.35 (m, 5H), 3.49 (t, J= 6 Hz, 1H), 3.57 (s, 3H, $CH_3$), 3.72 (brd, J= 6 Hz, 1H), 3.83 (dt, J= 1 and 6Hz, 1H), 3.97 (m, 1H), 4.10 (m, 1H), 4.80 (q, J= 6Hz, 1H), 7.74 (d, J=12 Hz, 1H), 8.76 (s,

1H). $^{13}$C NMR (151 MHz, cdcl$_3$) δ 8.74, 10.64, 24.16, 25.04, 26.44, 27.63, 35.78, 40.59, 40.61, 48.75, 51.30, 52.58, 56.46, 107.73, 107.96, 108.12, 118.81, 134.52, 137.36, 137.43, 141.08, 149.80, 152.97, 154.63, 157.95, 167.17, 176.79, 187.14.

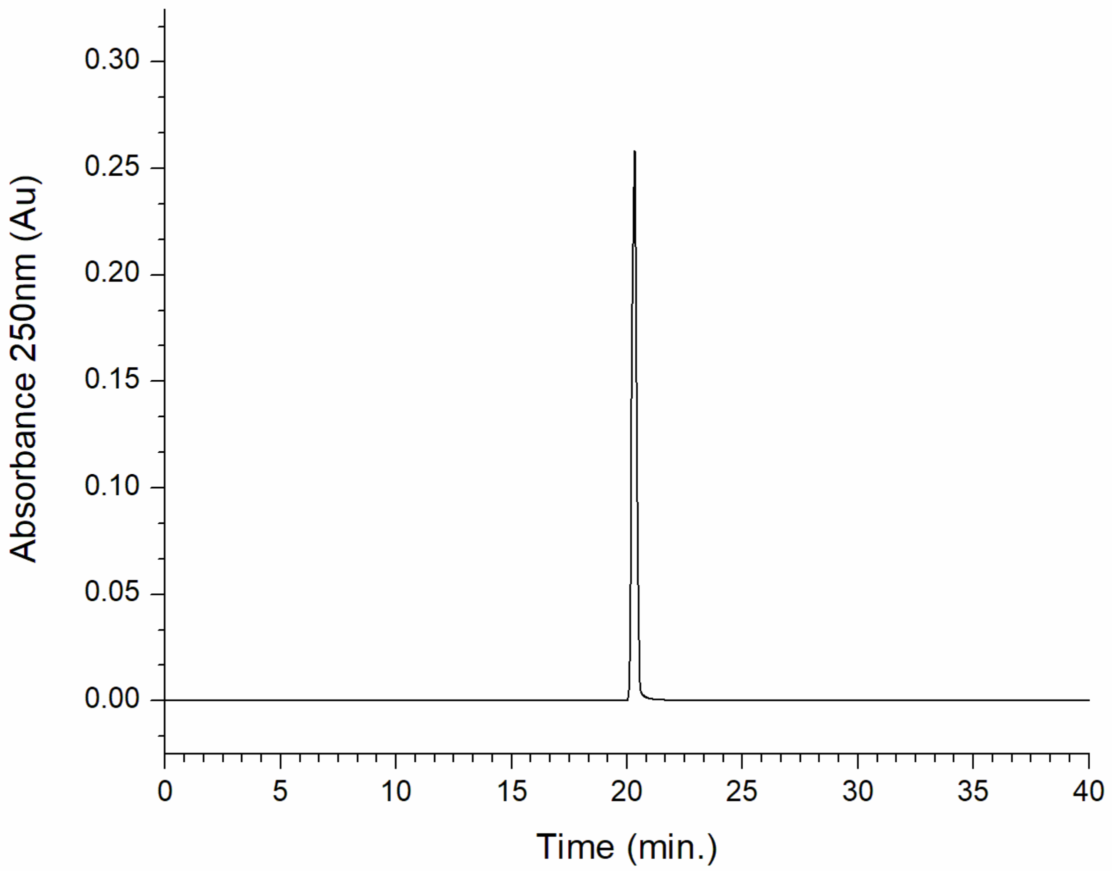

Analytical HPLC chromatogram of **moxAZ1**.

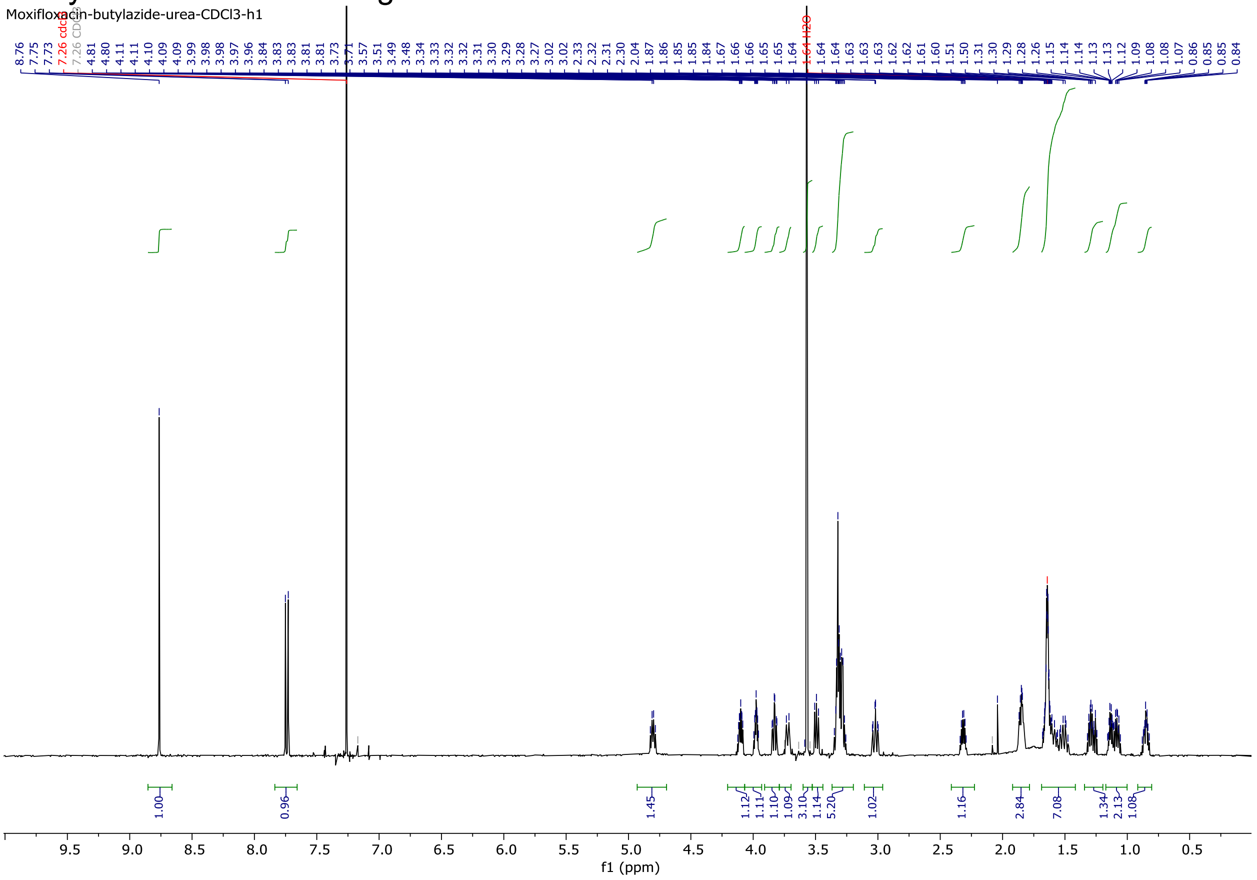


[1]H-NMR of **moxAZ1**.

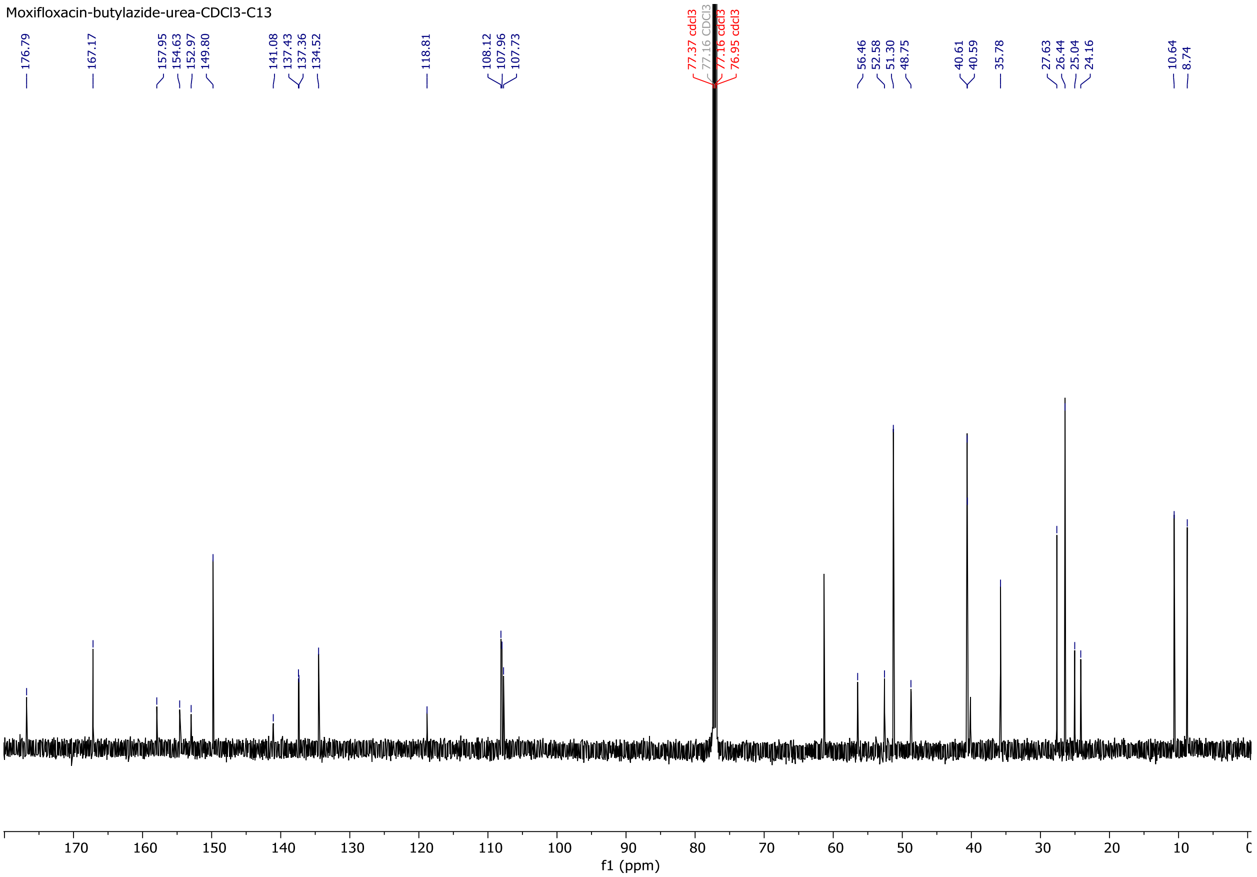


$^{13}C$-NMR of **moxAZ1**.

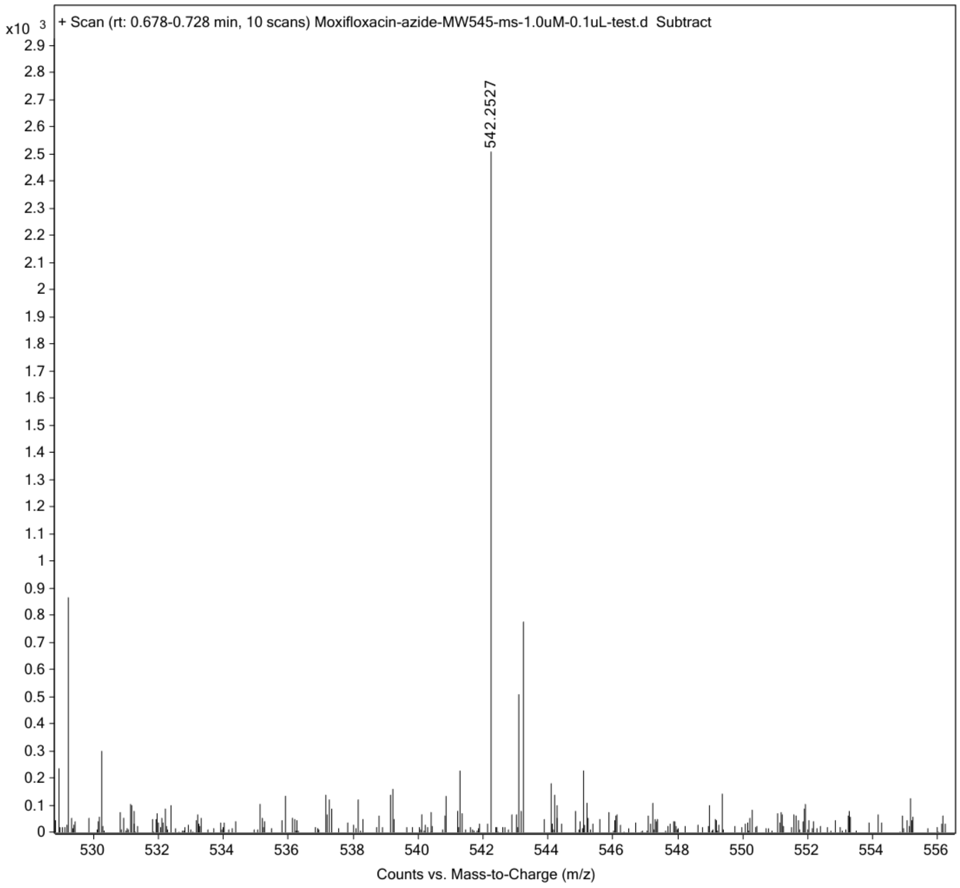


High resolution mass spectrum (ESI) of **moxAZ1**.

**Synthesis of novAZ1**.

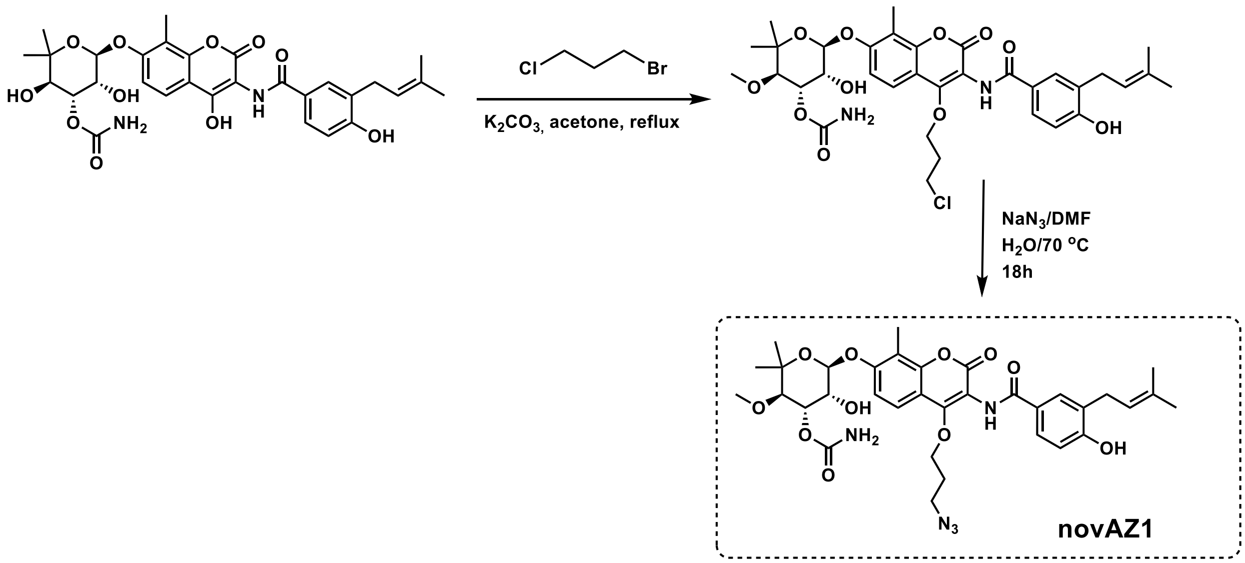


Novobiocin (122.4 mg, 0.2mmol, AK Scientific, Cat. # X5037) was suspended in acetone (10.0 mL) and to this solution, was added $K_2CO_3$ (500 mg) and 1-bromo-3-chloropropane (40 mg, 0.258 mmol). The mixture was heated to reflux with stirring for 4 hours during which time the reaction was monitored by TLC analysis to confirm the formation of the non-polar compound. The reaction mixture was then allowed to cool to room temperature and most of the acetone was removed under reduced pressure to afford a leftover residue that was dissolved in water and ethyl acetate (1:2 ratio, 30 mL total). The mixture was carefully acidified to pH 6.0 with dil. HCl (0.1M) following which the ethyl acetate layer was separated, washed with brine, and concentrated under reduced pressure. The crude material thus obtained was purified with silica gel column chromatography, and the compound was eluted with chloroform: methanol (9:1). The TLC-analyzed homogenous fractions were combined and concentrated under reduced pressure. The oily residue was triturated with ether to yield a white solid (95 mg, 69% yield). This chloropropyl ether was characterized by $^1$H-NMR and mass spectroscopy. MS (ESI): m/z 710.8 for $[M+Na]^+$.
The chloropropyl ether-novobiocin (69.0mg, 0.1 mmol) was dissolved in DMF: Water (9:1, 3mL). $NaN_3$ (26 mg, 4 mmol, 4.0eq.). The mixture was added to the solution and was heated at 70 °C for 16 hrs. The next day, TLC analysis showed a slightly more non-polar spot formed and mass analysis of the reaction mixture revealed complete conversion to azide. The reaction mixture was cooled to RT, diluted with ethyl acetate and water (1:1, 20 mL) and extracted. The organic layer was washed with water, brine and dried over

$Na_2SO_4$. Concentration under reduced pressure afforded the crude material, which was purified with silica gel column chromatography using chloroform: methanol (98:2), to yield the propyl-azide derivative of novobiocin as an off white solid (42 mg, 60%). NMR $^1H$ (600 MHz, $CD_3OD$): δ 7.77 (m, 2H, ArH), 7.71 (d, J = 12Hz, 1H, ArH), 7.25 (d, J = 12Hz, 1H, ArH), 6.86 (d, J = 6.0 Hz, 1H, ArH), 5.60 (s, 1H, OCHO), 5.33 (m, 2H), 4.53 (t, J = 6.0 Hz, 2H, O-$CH_2$) 4.25 (brs, 1H), 3.59 (d, J = 6 Hz, 1H), 3.56 (s, 3H, $OCH_3$), 3.51 (t, J = 6 Hz, 1H), 3.36 (d, J = 12 Hz, 1H), 2.32 (s, 3H, $ArCH_3$), 2.04 (m, 2H, $CH_2$), 1.75 (d, J = 6Hz, 6H, $CH_3$ x 2), 1.35 (s, 3H, $CH_3$),1.15 (s, 3H, $CH_3$). MS (ESI): m/z 696.2 and 718.2 and for $C_{16}H_{22}N_7O_3$ $[M + H]^+$ and $[M + Na]^+$ respectively ; observed 718.2701.

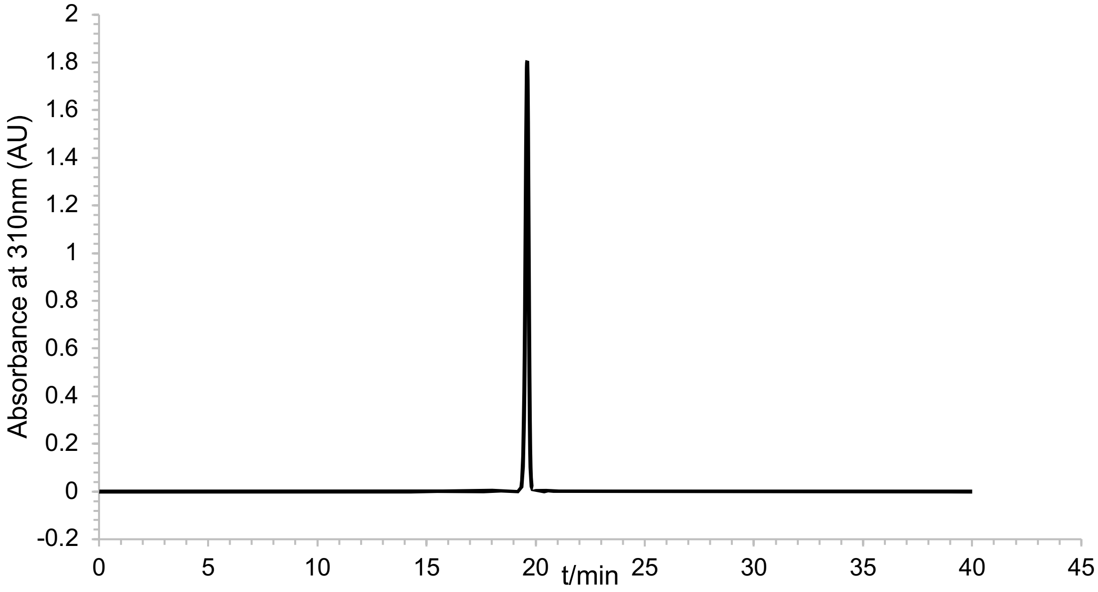

Analytical HPLC chromatogram of **novAZ1**.

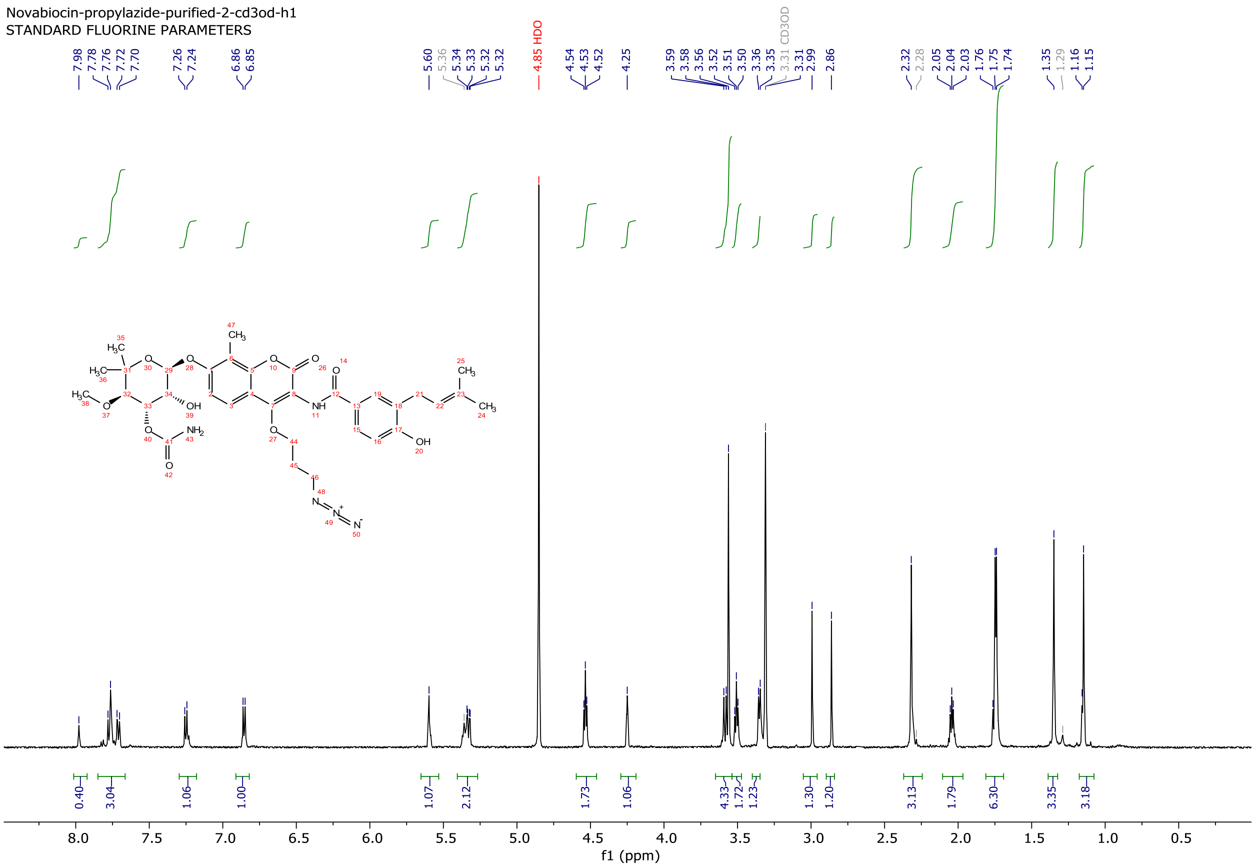


[1]H-NMR of **novAZ1**.

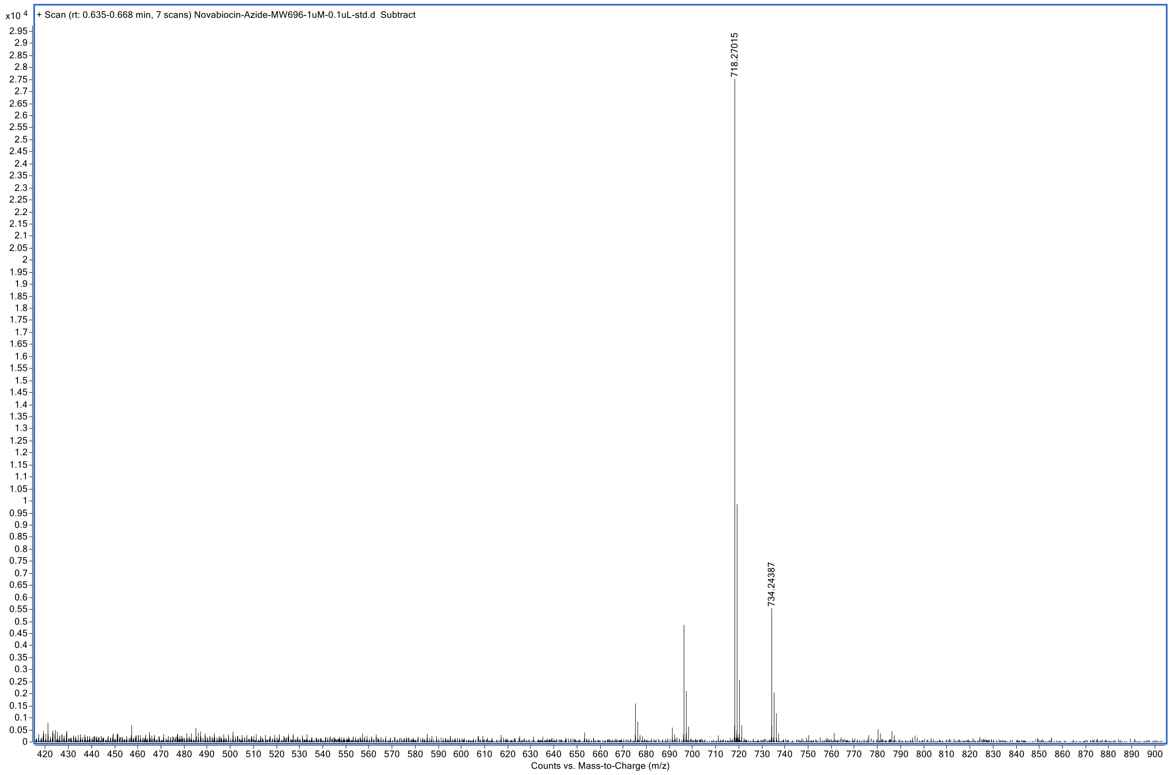


High resolution mass spectrum (ESI) of **novAZ1**.

**Synthesis of purAZ1**.

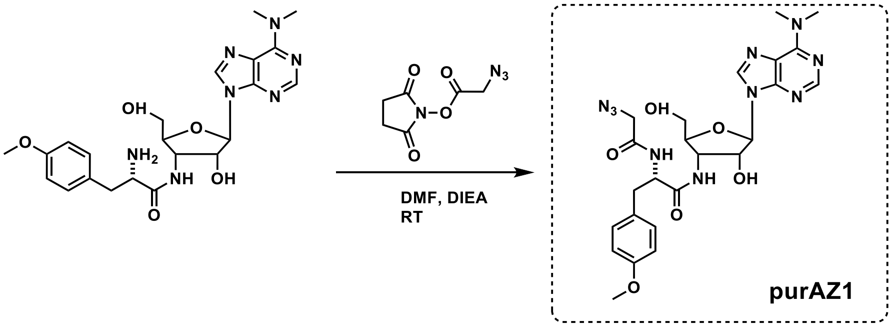


Puromycin (47.1mg, 0.1mmol) dissolved in dry DMF (1 mL), DIEA (50 µL, 37.1 mg, 0.29 mmol) was first added followed by NHS-ester of azido-acetic acid (24.5 mg, 0.12 mmol). The mixture was stirred under nitrogen at room temperature, and the reaction progress was followed by ESI MS analysis, observed m/z 555 for expected product. At the end of 12 h stirring starting material appears to be completely consumed, and the new product was formed. The mixture was then diluted with ether (10 mL) to allow oily residue to separate out at room temperature, the supernatant ether layer discarded. The oily residue was further triturated with 1 mL ether to remove most of the DMF, and the left-over oily residue was used for further purification. The residue was dissolved first dissolved in HPLC grade acetonitrile (1 mL), followed by dilution with 90:10 ($H_2O$:$CH_3CN$ containing 0.1%TFA, 8 mL). The solution was then subjected to preparative HPLC purification using starting gradient of 10% B ($CH_3CN$ containing 0.1%TFA), and reaching to 100% B in min. The fractions containing desired product were combined, concentrated under reduced pressure using rotary evaporator, the remaining aqueous solution was frozen to -80 $^{o}C$ and lyophilized to obtain the dry residue. The compound was analyzed by analytical HPLC, characterized by High resolution mass and $^{1}H$-NMR as follows: HRMS observed m/z 555.2419 $[M+H]^+$ and 577.2243 $[M+Na]^+$; expected m/z 555.2423 and 577.2242 for $C_{24}H_{30}N_{10}O_6$ $[M+H]^+$ and $[M+Na]^+$ respectively. $^{1}H$ NMR (600 MHz, $CD_3OD$) δ 8.55 (brs, 1H), 8.40 (d, J=6 Hz, 1H), 8.34 (s, 1H), 8.23 (d, J= 6 Hz, 1H), 7.79 (d, J= 6 Hz, 1H), 7.79 (d, J= 6 Hz, 1H), 7.39 (t, J= 6 Hz, 1H), 7.31 (t, J= 6 Hz, 1H), 7.16 (d, J= 12 Hz, 2H), 6.88 (d, J= 12 Hz,

2H), 6.27 (brd, J= 3Hz, 1H), 5.74 (dd, J= 6, 3 Hz, 1H), 4.52 (q, J= 6Hz, 1H), 4.37 (d, J= 6 Hz, 1H), 4.24 (brs, 1H), 4.21 (m, 1H), 3.98-4.06 (m, 2H), 3.84 (s, 3H), 3.78 (brs, 6H), 3.46 (brd, J= 6 Hz, 2H), 3.00 (dd, J= 6 Hz, 1H), 2.91 (dd, J= 6 Hz, 1H).

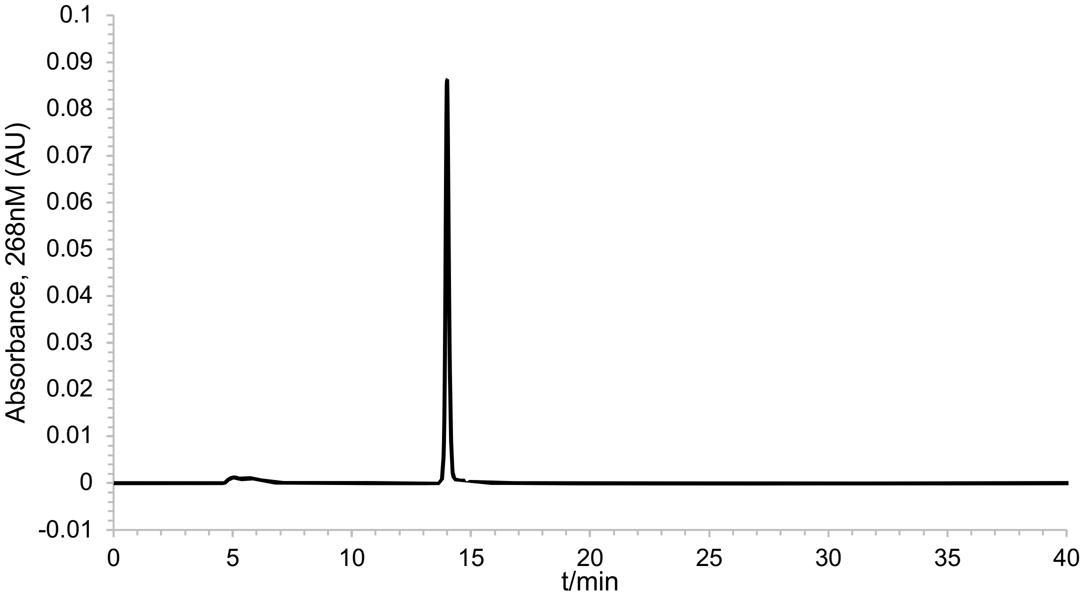


Analytical HPLC chromatogram of **purAZ1**.

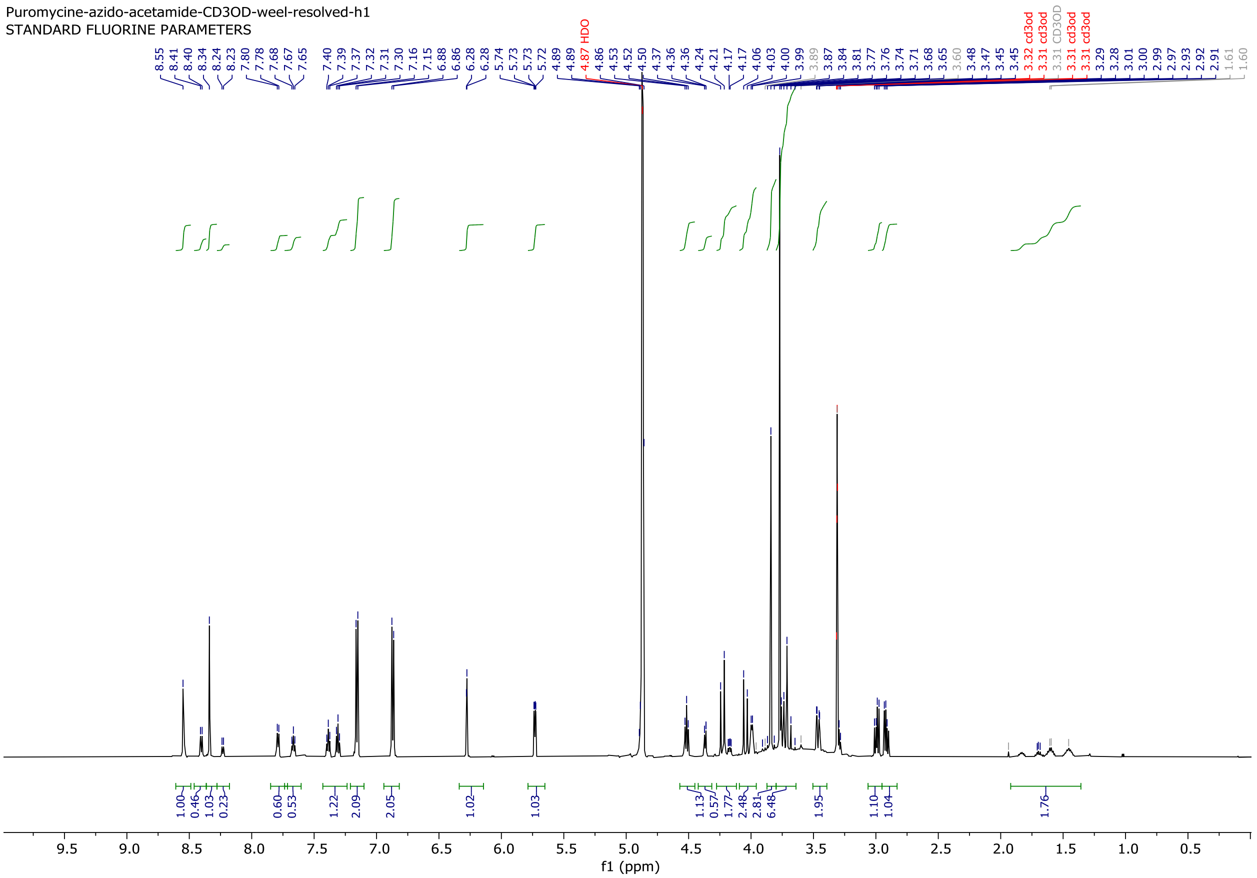


$^1$H-NMR of **purAZ1**.

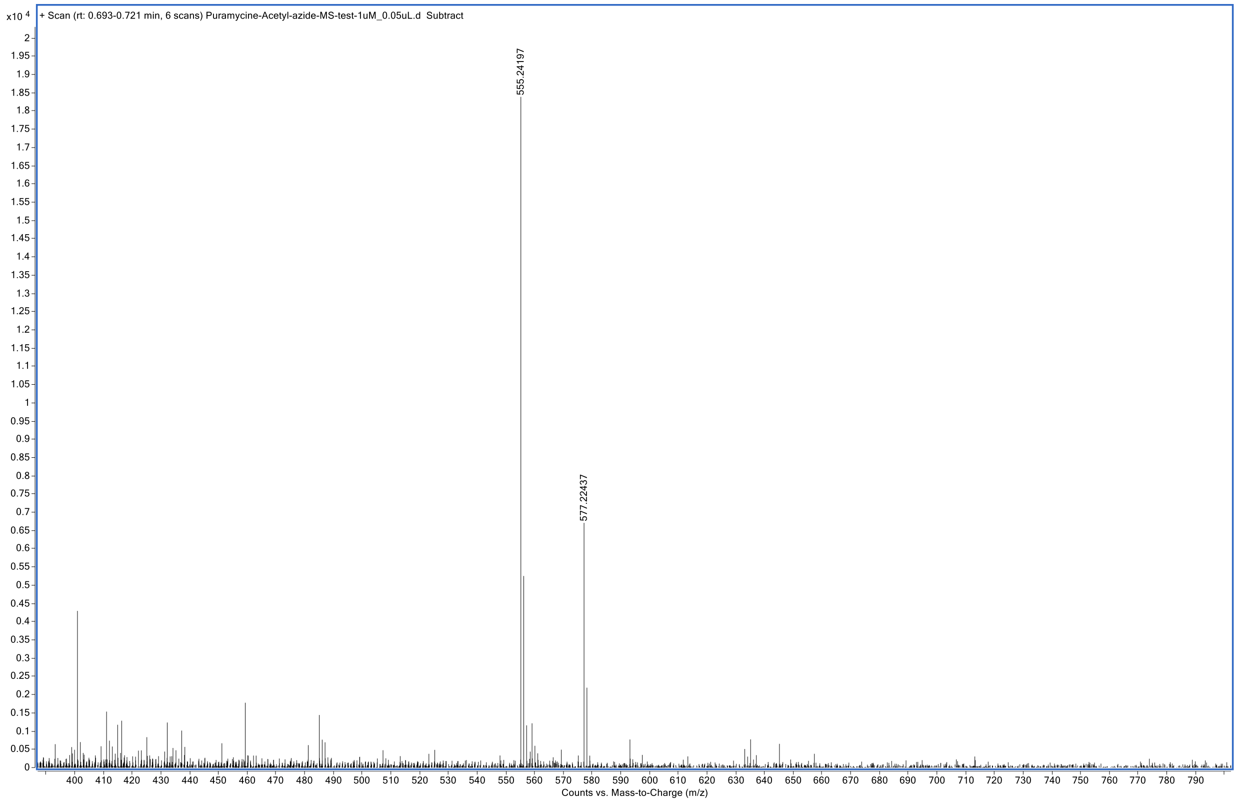


High resolution mass spectrum (ESI) of **purAZ1**.

**Synthesis of zolAZ1.**

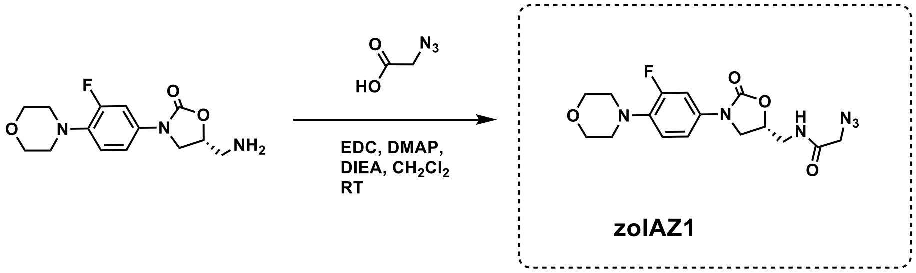


**zolAZ1**

Following a published protocol, **zolAZ1** was synthesized and characterized as follows:[4] To a solution of amine in $CH_2Cl_2$ (10 mL), EDCI.HCl (107 mg, 0.56 mmol) and 2-azidoacetic acid 24 (42 µl, 57 mg, 0.56 mmol) were added. The reaction mixture was then stirred for 3 h at 20 °C. The reaction mixture was adsorbed on silica gel and purified on a silica gel column ($CH_2Cl_2$ then $CH_2Cl_2$/EtOH: 96/4) to afford the compound 5 (134 mg, 0.34 mmol, yield: 76%) isolated as a white solid. NMR $^1$H (400 MHz, $CDCl_3$): δ 7.38 (dd, J = 2.6, 14.3 Hz, 1H), 7.05–7.01 (m, 2H), 6.88 (dd, J = 8.8, 9.4 Hz, 1H), 4.78–4.71 (m, 1H), 4.01 (t, J = 8.9 Hz, 1H), 3.96 (s, 2H), 3.84–3.81 (m, 4H), 3.75–3.56 (m, 3H), 3.02–3.00 (m, 4H). NMR $^{13}$C (100 MHz, $CDCl_3$): δ 168.0, 156.6, 156.8, 136.8, 132.9, 119.0, 114.1, 107.7, 71.7, 67.1, 52.5, 51.1, 47.9, 42.2. MS (ESI): m/z 379.2 $[M + H]^+$.

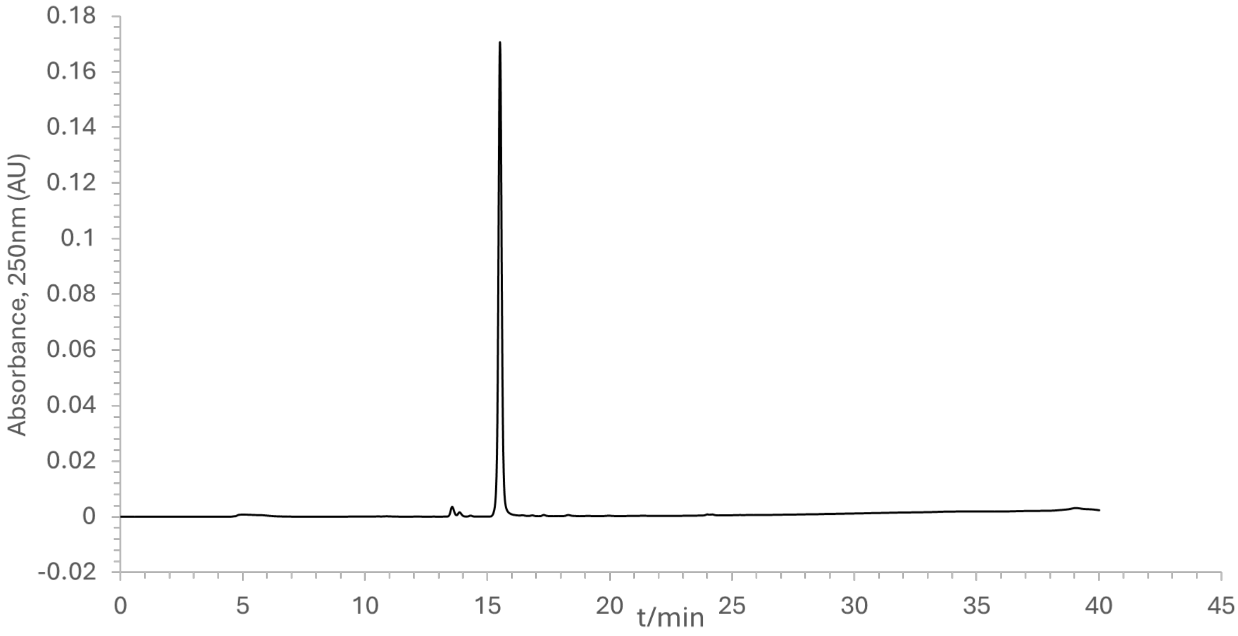


Analytical HPLC chromatogram of **zolAZ1**.

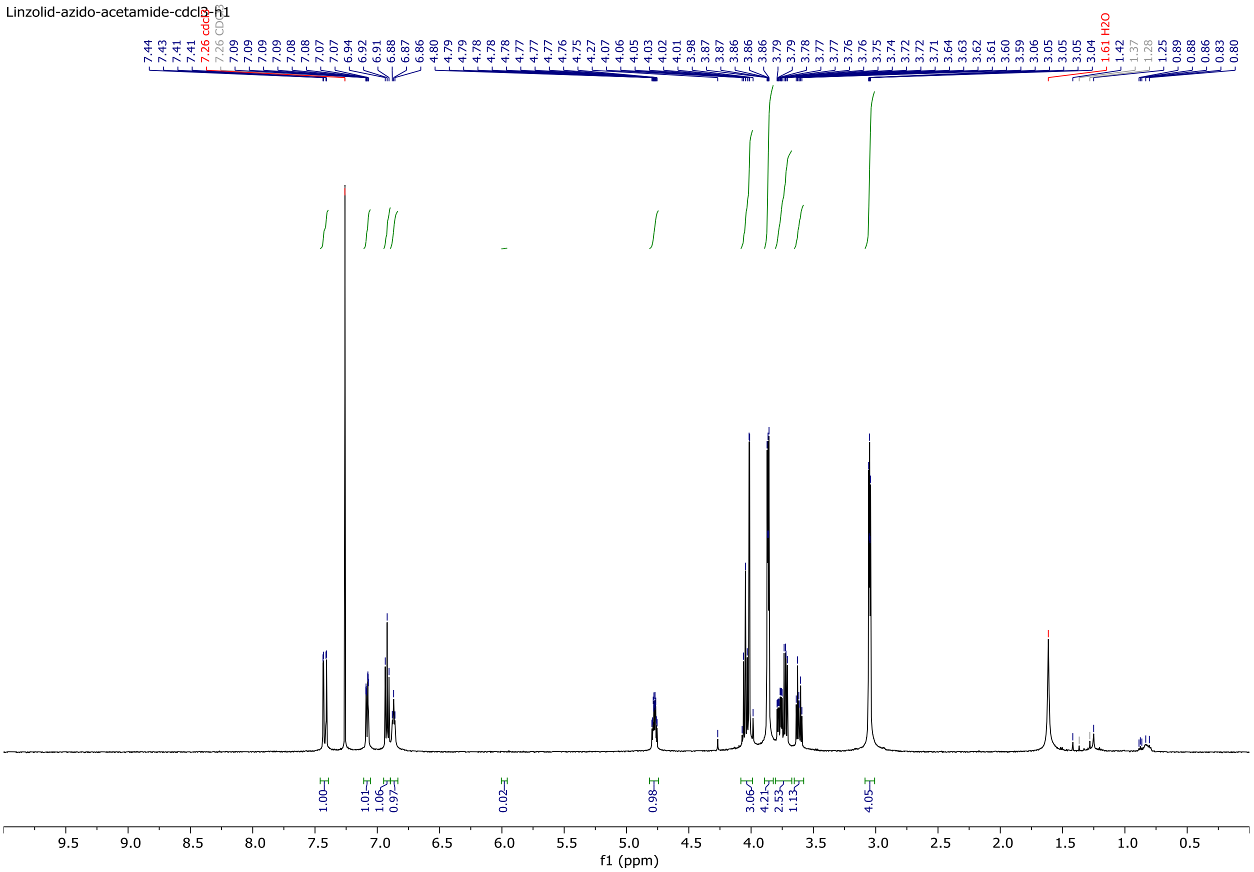

[1]H-NMR of **zolAZ1**.

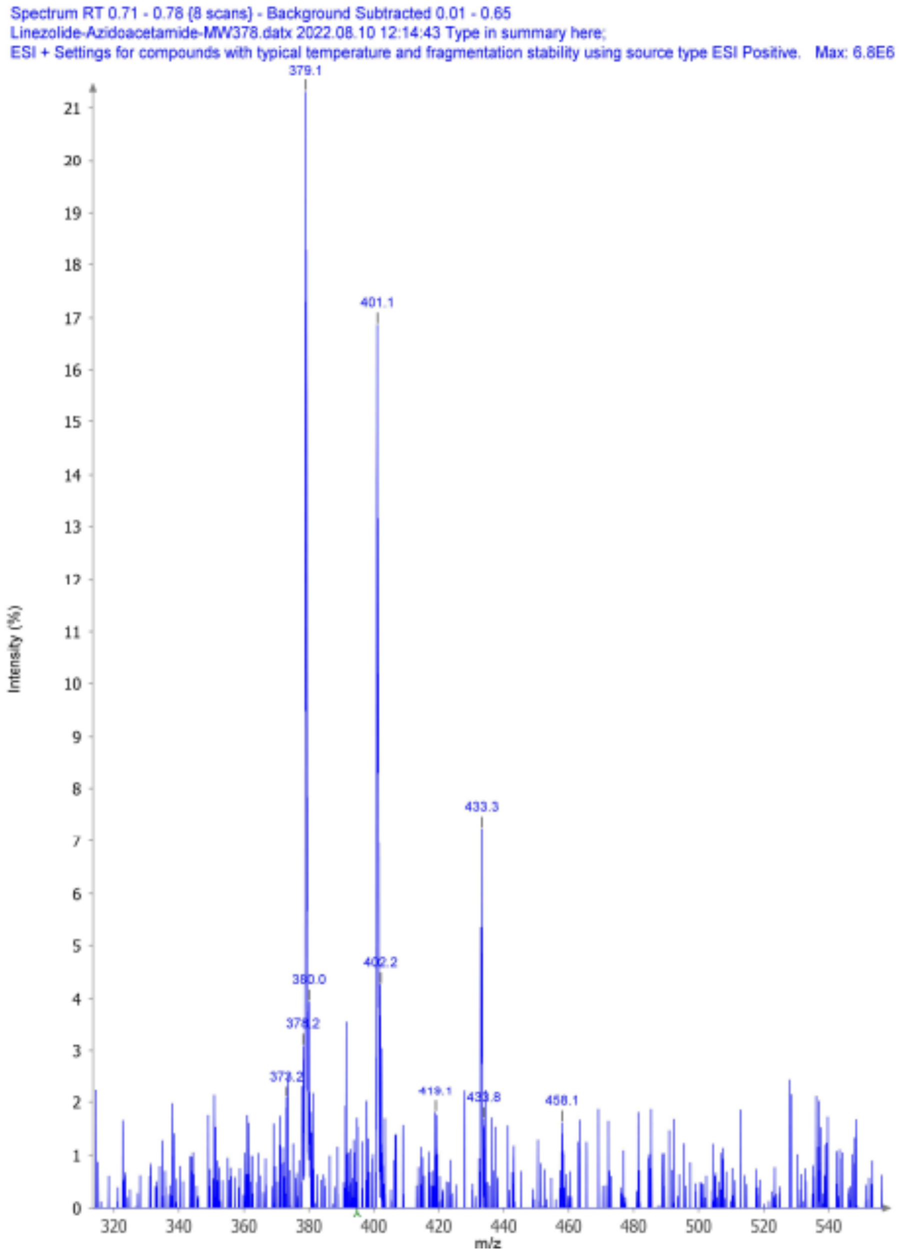


Mass spectrum (ESI) of **zolAZ1**.

**Synthesis of zolAZ2.**

Following a published protocol, **zolAZ2** was synthesized and characterized as follows:[5]

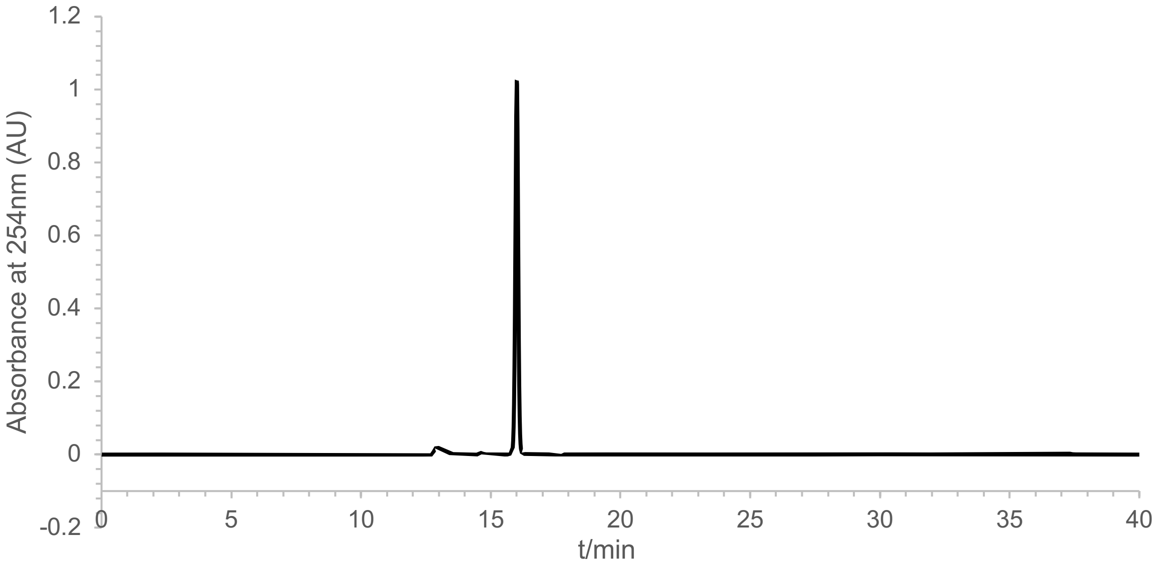


Analytical HPLC chromatogram of **zolAZ2**.

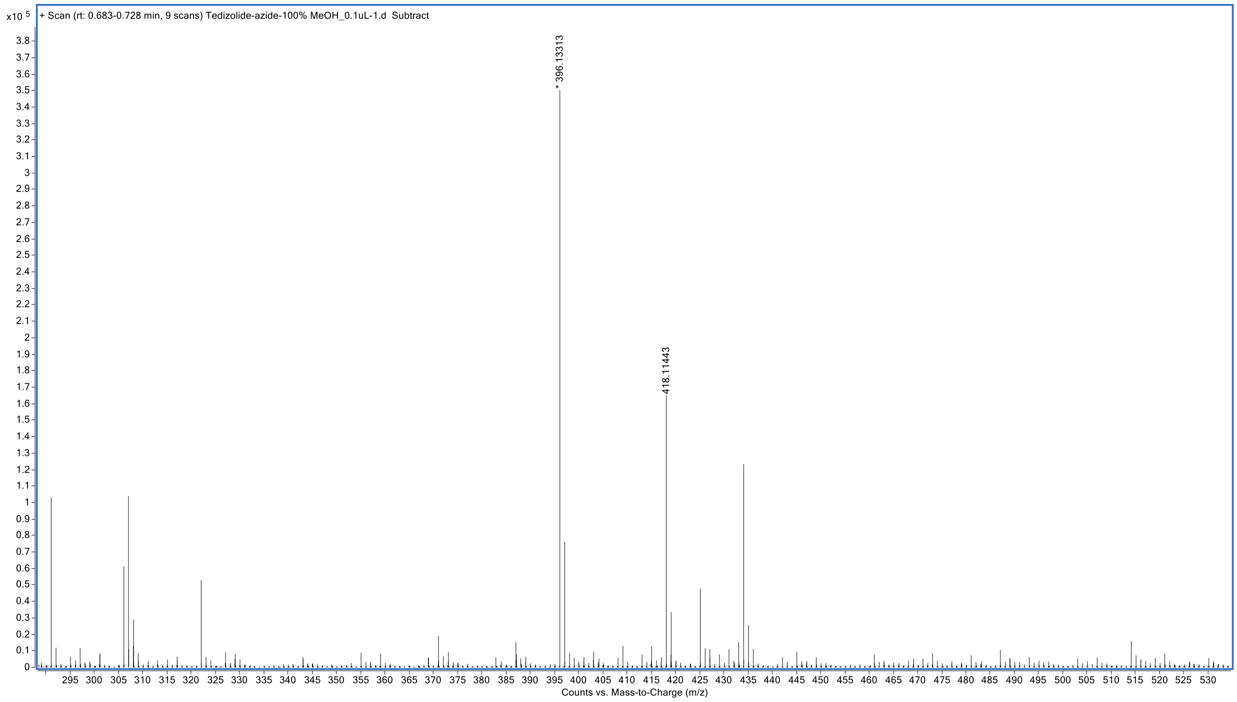


High resolution mass spectrum (ESI) of **zolAZ2**.

#### Synthesis of zolAZ3.

Following a published protocol, **zolAZ3** was synthesized and characterized as follows:[5]

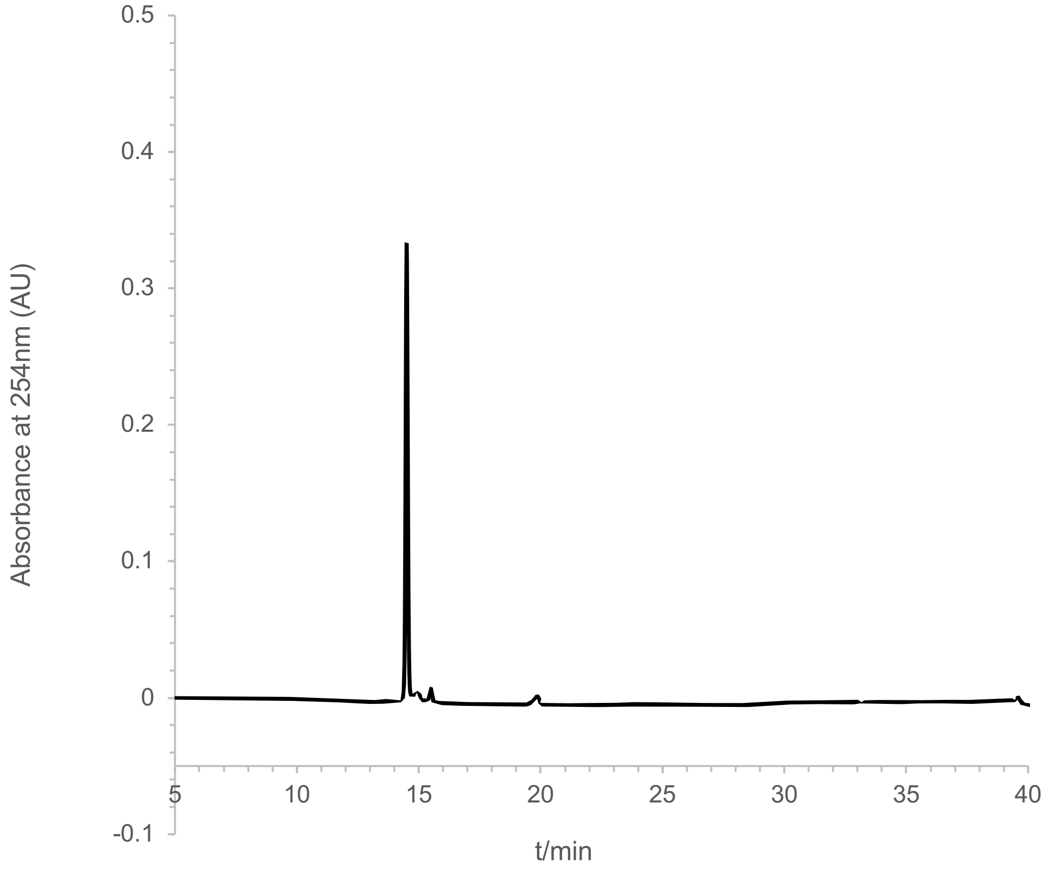


Analytical HPLC chromatogram of **zolAZ3**.

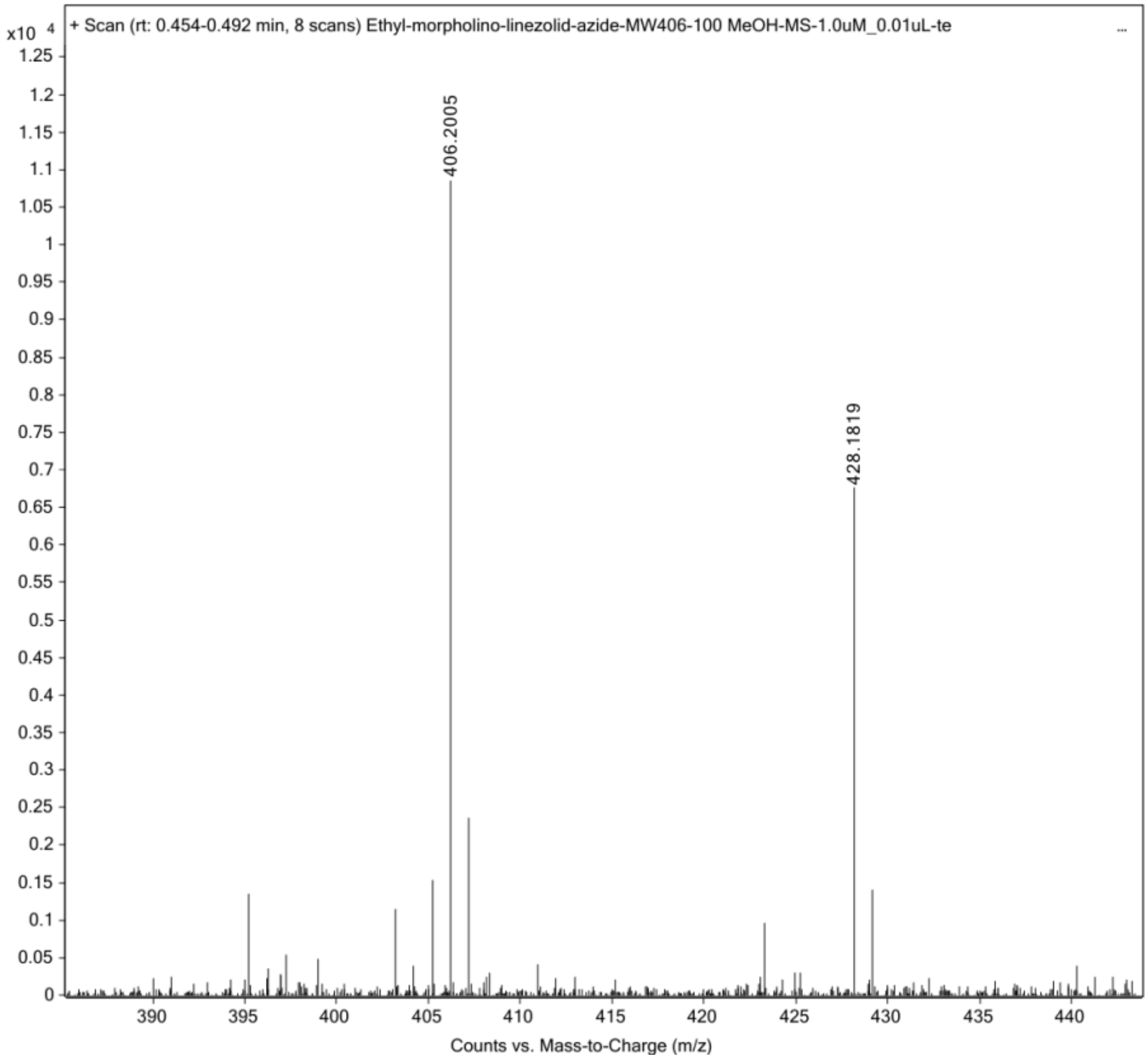


High resolution mass spectrum (ESI) of **zolAZ3**.

**Synthesis of metAZ1.**

Following a published protocol, **metAZ1** was synthesized and characterized as follows:[6]

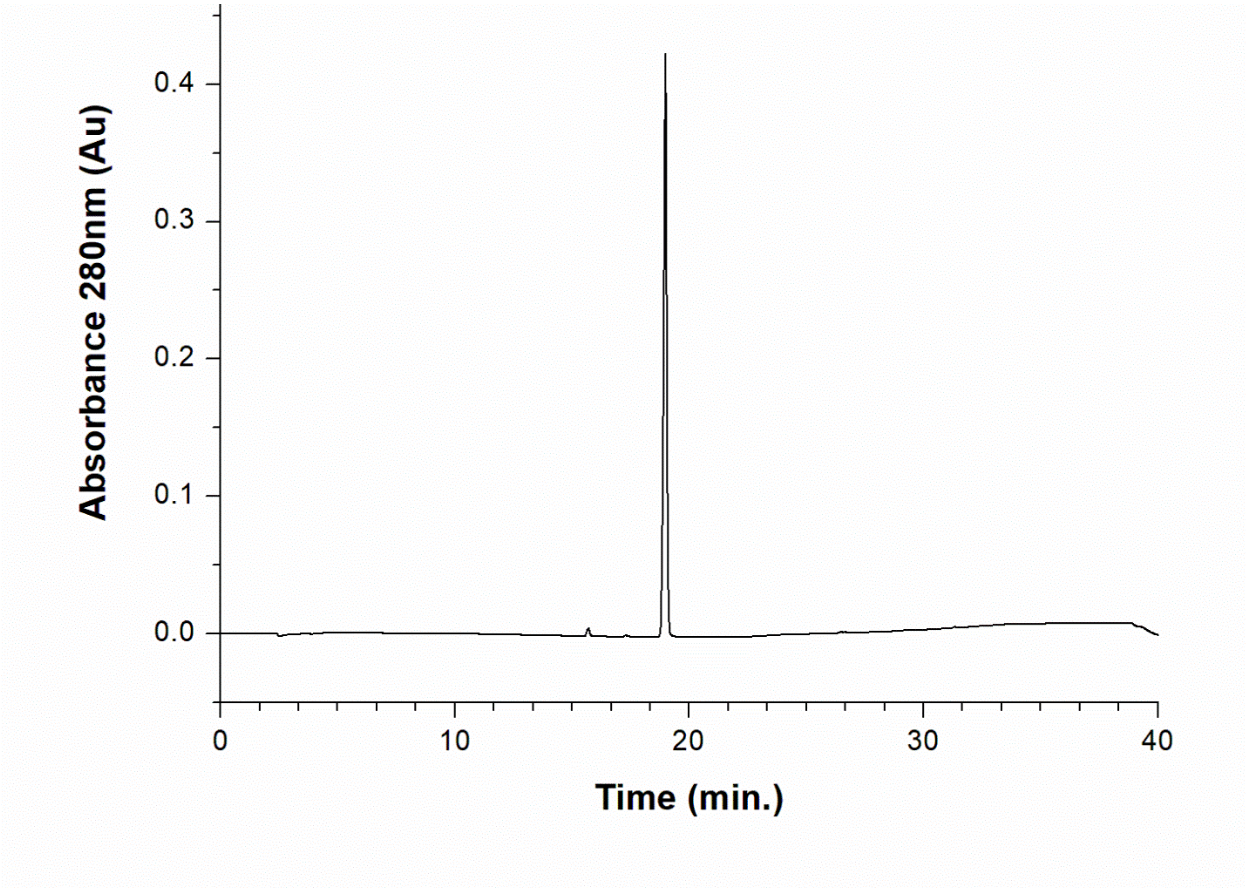


Analytical HPLC chromatogram of **metAZ1**.

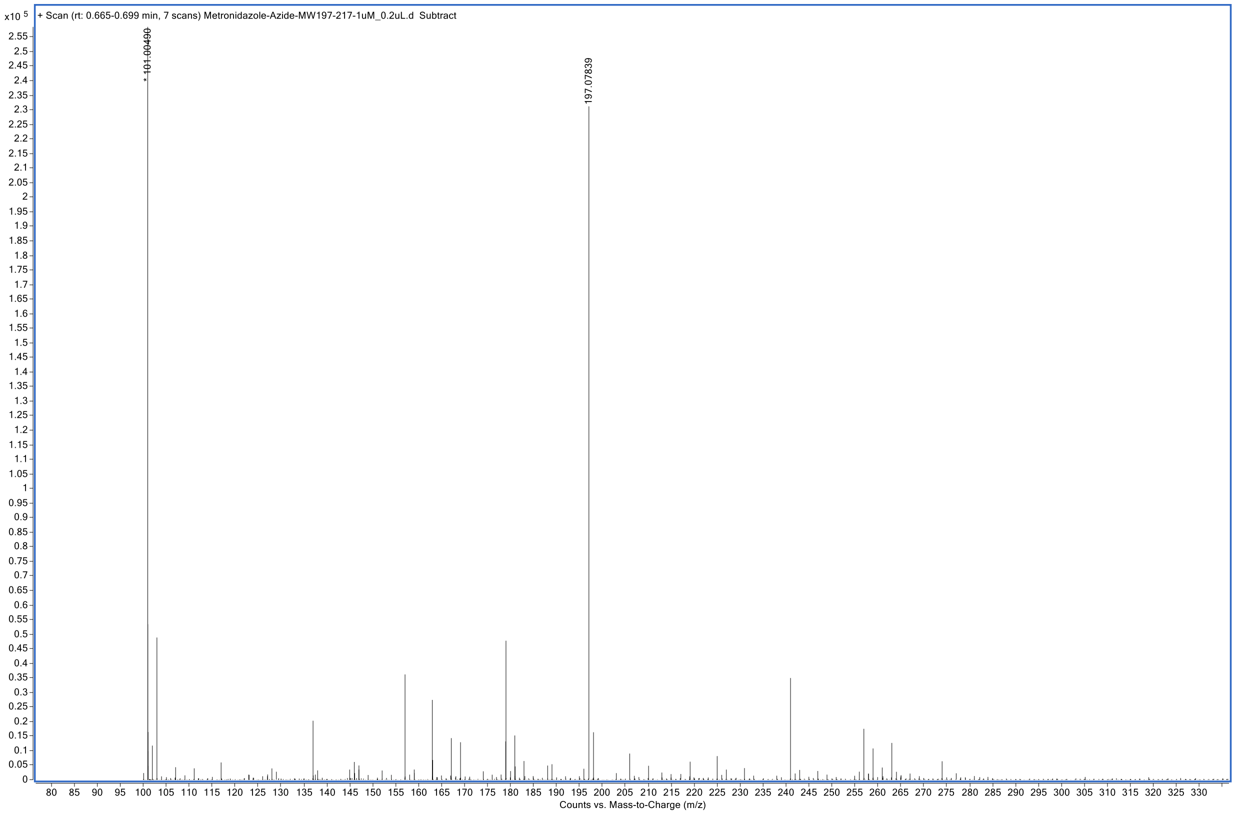


High resolution mass spectrum (ESI) of **metAZ1.**

**Synthesis of sulAZ1.**

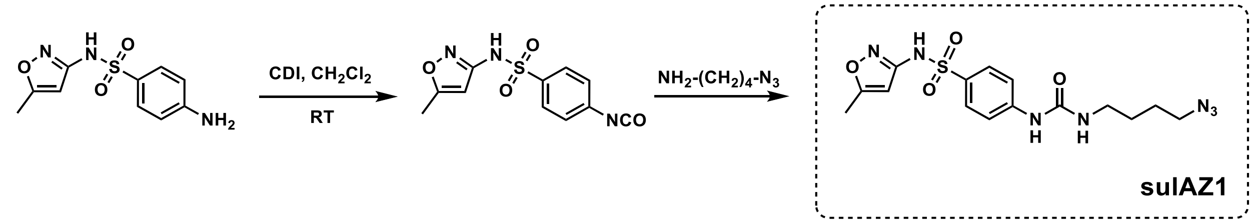


Following a published protocol, **sulAZ1** was synthesized and characterized as follows:[7] A solution of sulfamethoxazole (51.0 mg, 0.2 mmol) in anhydrous $CH_2Cl_2$ (5 mL) was mixed with CDI (40 mg, 0.25 mmol) at room temperature. The homogenous mixture was stirred overnight (~16 h). Next day 4-azidobutyl amine was added and the mixture was further stirred for 4hrs, TLC and ESI mass analysis indicated the new product formation (non-polar compound compared with starting material). The reaction was quenched with aq. citric acid solution, diluted with 10 mL $CH_2Cl_2$ and organic layer separated. Organic washed with water, brine and dried over $Na_2SO_4$. Concentration under reduced pressure using rotary evaporator followed by silica gel chromatography hexane:ethyl acetate (70:30) yielded the desired compound as solid (61.0 mg, 62%). The compound was analyzed by analytical HPLC, characterized only by High resolution mass and used as is: HRMS observed m/z 394.1296 $[M+H]^+$ and 416.1114 $[M+Na]^+$; expected m/z 394.1292 and 416.1111 for : $C_{15}H_{20}N_7O_4S$ $[M+H]^+$ and $[M+Na]^+$ respectively; observed 394.1296. $^1H$ NMR (600 MHz, $CD_3CN$) δ 1.53-1.62 (m, 4H), 2.30 (s, 3H, $CH_3$), 3.18 (q, J= 6Hz, 2H), 3.32 (t, J= 6 Hz, 2H), 6.11 (s, 1H), 7.53 (s, 1H), 7.56 (d, J= 6 Hz, 2H), 7.72 (d, J= 6 Hz, 2H).

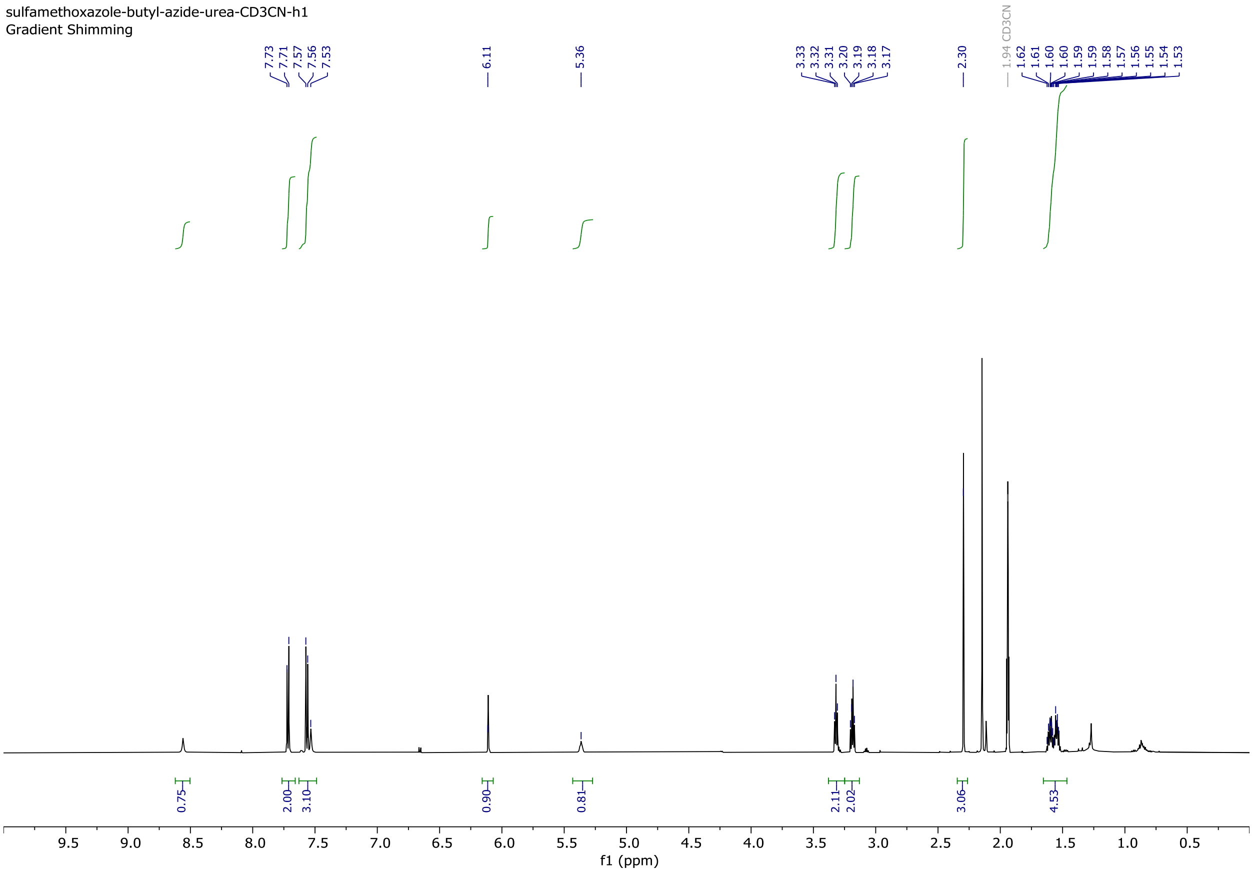


$^1$H-NMR of **sulAZ1**.

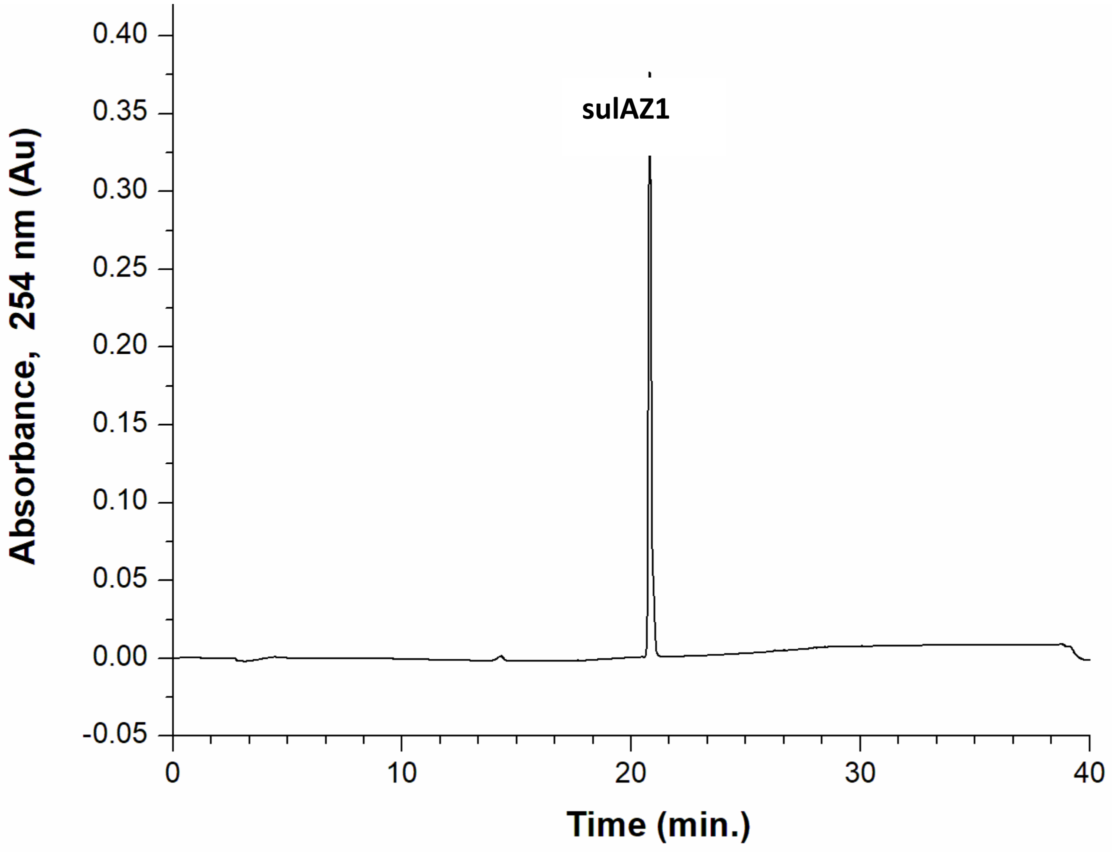


Analytical HPLC chromatogram of **sulAZ1**.

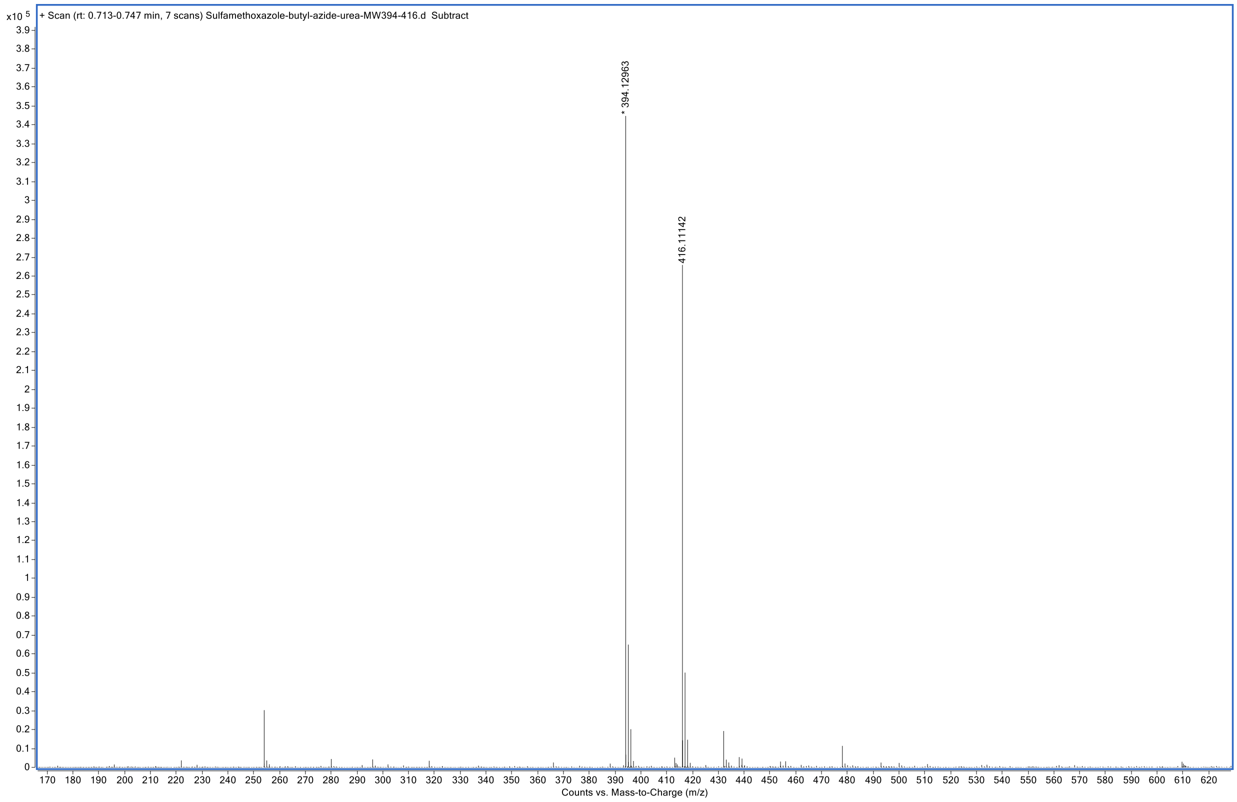


High resolution mass spectrum (ESI) of **sulAZ1**.

#### General methods for synthesis of peptide probes

All peptides were built by solid-phase peptide synthesis. Amidated C-terminus peptides were built starting from Fmoc-aminomethyl phenoxy resin (Rink amide resin) procured from Sigma-Aldrich (St. Louis, MO, USA); COOH terminated peptides were built on either chloro-trityl resin (Sigma-Aldrich) or Fmoc-amino acid preloaded on Wang resin procured from p2bioSystems (Louisville, KY 40223). Crude peptides were purified by preparative reverse phase HPLC purification instruments equipped with Waters 1525 pumps and 2489 UV/Visible Detector on a Phenomenex Luna 10 µm C8(2) 100 Å (250 x 21.2 mm) or C18 columns using a 5 to 100% linear gradient of methanol (MeOH) in $H_2O$ or acetonitrile (MeCN) in water each containing 0.1% TFA at 10 mL/min. The HPLC fractions of the desired compounds were first concentrated under reduced pressure using a rotary evaporator. The concentrated aqueous solutions were then frozen and lyophilized with Labconco Freezone 4.5 L lyophilizer (-84$^{o}$C). The purity of the samples was ascertained by analytical HPLC using a Phenomenex Luna 5 µm C8(2) or C18 columns of 100 Å (250 x 4.6 mm) on the same instrument; using gradient elution in $H_2O$/MeCN or $H_2O$ /MeOH with 0.01% TFA in each solvent at 1 mL/min prior to use in biological assays. Electrospray ionization (ESI)-based mass analysis was performed on Advion Expression® CMS mass spectrometer using standard parameters for intermediates. A low fragmentation and low energy setup were used to analyze fragmentation-sensitive compounds. Reaction progress and purification of products were analyzed with Additional LCMS analysis, which were performed on Shimadzu Prominence-i LCMS system equipped with LC-2030C 3D liquid chromatograph, autosampler, PDA and LCMS-2020 mass detector. Data analysis was performed with Lab Solutions software. MALDI-TOF mass spectra were obtained for certain high molecular weight compounds on the Shimadzu MALDI-8020 instrument with α-Cyano-4-hydroxycinnamic acid (α-CHCA) matrix. The observed molecular weights for compounds were represented as m/z. High resolution electrospray ionization mass spectrometry (HRMS, ESI/MS) analyses were obtained on an Agilent 6545B Q-TOF LC/MS equipped with 1260 Infinity II LC system with auto sampler and equipped with C18(2) column (Luna 5 µm 100Å 250 x 4.6 mm. Quantitative LCMS analysis was performed on Agilent 6545B Q-TOF as describe above in positive ion mode. For each antibiotics-azide tested, parameters were such as collision energy and fragmentation pattern (formation of defined daughter ions and their intensities were optimized). Prior to actual quantification of recovered antibiotics-azides from cell lysates, standard curve at lowest possible concentration were obtained. The LCMS/MS data was processed with Mass Hunter software, and subsequent quantification was done with Q-TOF MS quantitative software. UV spectroscopic analysis was performed on Genesys 50 (Thermo Scientific) UV-Visible Spectrophotometer.$^{1}$H and $^{13}$C-NMR spectra for final compounds and intermediates were acquired on a Varian 600MHz spectrophotometer. $^{1}$H and $^{13}$C-NMR spectra for final compounds and intermediates were acquired on a Varian 600MHz spectrophotometer. All NMR spectra were processed and analyzed using MestreNova software. Residual solvent signals from deuterated solvents were used as an internal standard with reference to tetramethylsilane (TMS) for defining chemical shifts. Chemical shifts are reported in ppm (d) and coupling constants (J) are reported in Hertz [Hz].

Fmoc-Lys(N3)-OH (**2n**) is commercially available and was purchased from aapptec (Louisville, KY 40228) and used without further purification.

**Synthesis of 1p.**

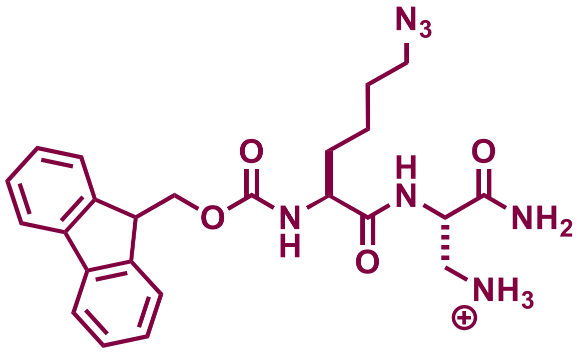


100 mg of Fmoc-aminomethyl)-phenoxy resin (Rink amide resin) of loading capacity 0.48 mmol/g was swollen in DCM after which the Fmoc protecting group was removed using 20% piperidine in DMF for 30 min. The resin was washed using alternating DCM and MeOH and finally rinsed with DMF. Fmoc-Dap (Boc)-OH was coupled to the resin using 4 equivalents of the amino acid and 4 equivalents each of Oxyma and DIC; the coupling reaction was done at room temperature with shaking for 2 h. The Fmoc group was similarly removed using 20% piperidine and following wash steps as described above, Fmoc-Lys($N_3$)-OH was similarly coupled to Dap (Boc) to make Fmoc-Lys($N_3$)-Dap(Boc)-Rink amide resin. The peptide was cleaved off the resin using 95% TFA:2.5% TIPS: 2.5% $H_2O$ and purified by preparative RP-HPLC on a on a Phenomenex Luna 10 µm C8(2) 100 Å (250 x 21.2 mm) using a 20 to 100% linear gradient of MeCN in $H_2O$ each containing 0.1% TFA at 10 mL/min. ESI-MS $[M+H]^+$ m/z calcd. 480.2354 for $C_{24}H_{30}N_7O_4$, obsd. 480.2356. $^1$H NMR (600 MHz, CD3CN) δ 1.41-1.44 (m,1H), 1.55-1.59 9M, 2H), 1.60-1.67 (m, 4H), 1.78-1.82 (m, 1H), 3.23 (dd, J= 1 and 6 Hz, 1H), 3.29 (t, J= 6Hz, 2H), 3.38 (dd, J= 1 and 6 Hz, 1H), 4.05 (brs, 1H), 4.23 (t, J= 6Hz, 1H), 4.31 (t, J= 6Hz, 1H), 4.35 (dd, J= 2 and 6Hz, 1H), 4.64 (brs, 1H), 6.14 (brs, 1H), 6.57 (brd, J= 6Hz, 1H), 7.07 (brs, 1H), 7.34 (t, J= 6Hz, 2H), 7.42 (t, J= 6Hz, 2H), 7.69 (brt, J= 6Hz, 2H), 7.83 (d, J= 6Hz, 2H), 8.20 (brd, J= 6Hz, 1H).

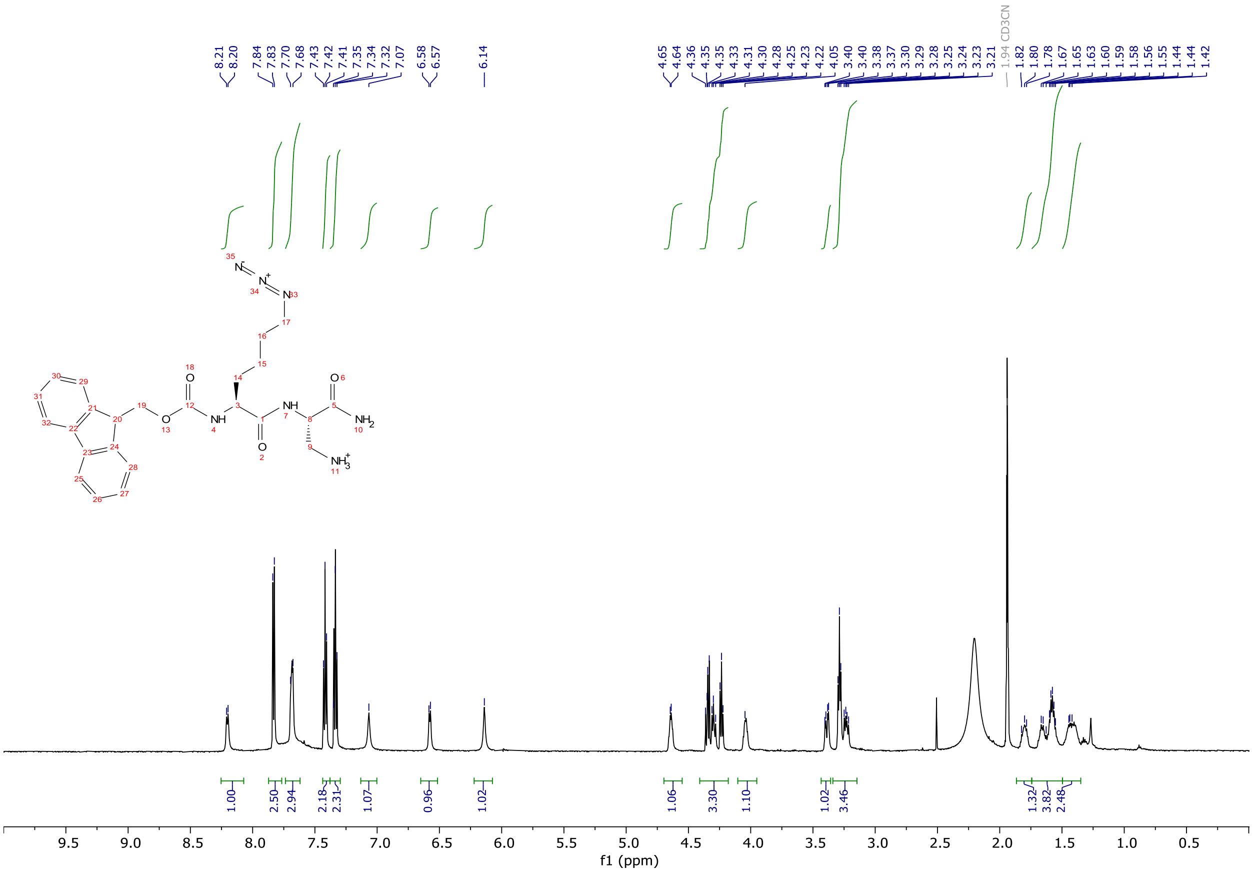

$^{1}$H-NMR of **1p** in $CD_3CN$.

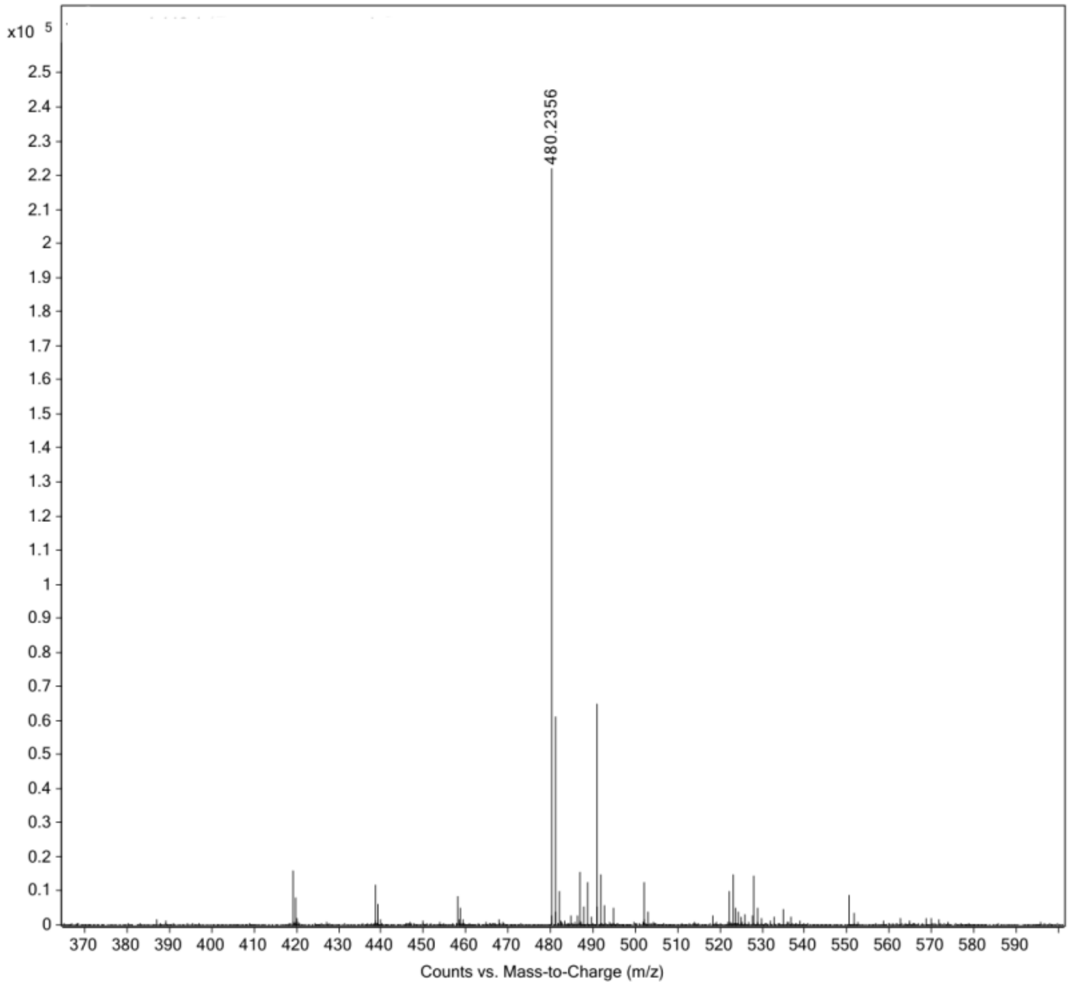


High resolution mass spectrum (ESI) of **1p**.

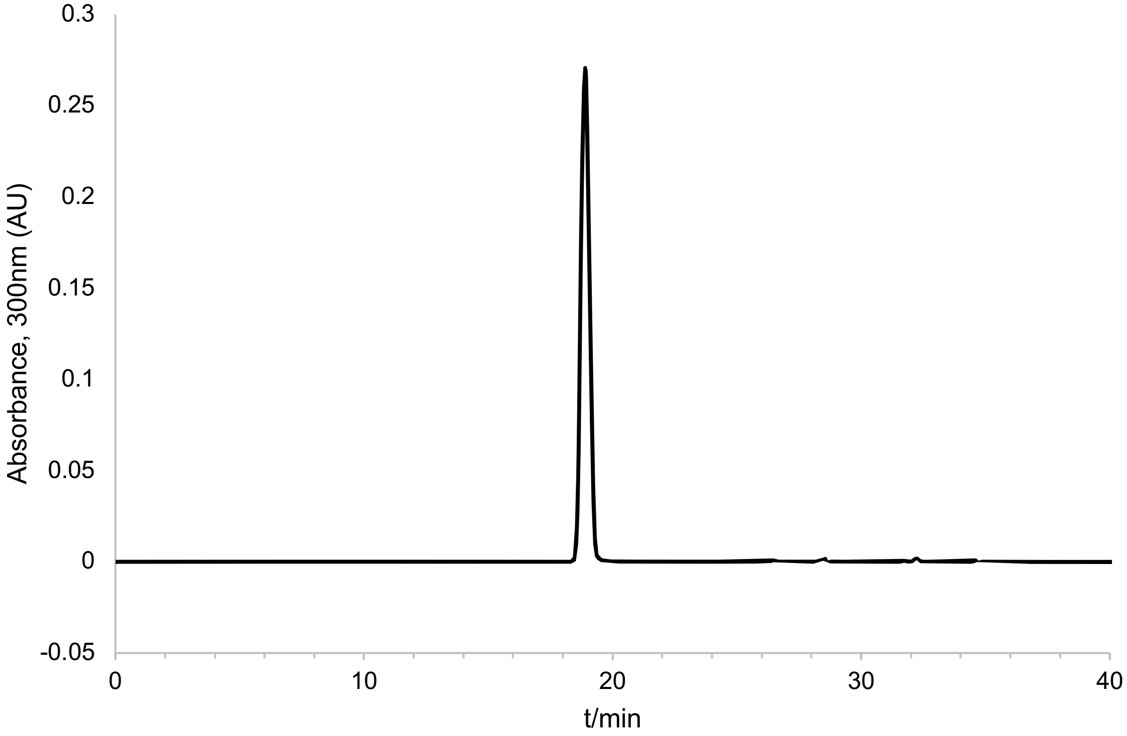


Analytical HPLC chromatogram of **1p**.

**Synthesis of 1n**.

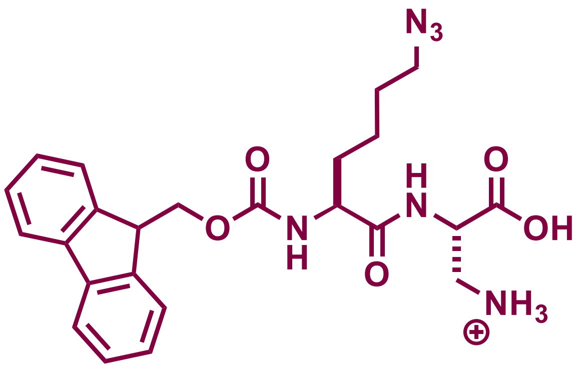


50 mg of 2-chlorotrityl chloride resin (Rink amide resin) of loading capacity 1.5 mmol/g was swollen in dry DCM in a peptide synthesis vessel. Fmoc-Dap (Boc)-OH was dissolved in dry DCM (1eq.) and added to the resin and shaken for 5 min after which 4 eq of DIEA was added and incubated for 1 h with shaking at room temperature. The resin was washed 3X with dry DCM, washed 1X with MeOH, and rinsed with DMF. Next, the Fmoc protecting group was removed using 20% piperidine in DMF for 30 min. The resin was washed using alternating DCM and MeOH and finally rinsed with DMF. Fmoc-Lys($N_3$)-OH was coupled to the resin using 4 equivalents of the amino acid and 4 equivalents each of Oxyma and DIC; the coupling reaction was done at room temperature with shaking for 2 h to make Fmoc-Lys($N_3$)-Dap (Boc)-trityl chloride resin. The peptide was cleaved off the resin using 95% TFA:2.5% TIPS: 2.5% $H_2O$ and purified by preparative RP-HPLC on a on a Phenomenex Luna 10 µm C8(2) 100 Å (250 x 21.2 mm) using a 20 to 100% linear gradient of MeCN in $H_2O$ each containing 0.1% TFA at 10 mL/min. ESI-MS $[M+H]^+$ m/z calcd. 481.2194 for $C_{24}H_{29}N_6O_5$, obsd. 481.2197. $^1$H NMR (600 MHz, CD3CN) δ 1.41-1.44 (m,1H), 1.55-1.59 9M, 2H), 1.60-1.67 (m, 4H), 1.78-1.82 (m, 1H), 3.23 (dd, J= 1 and 6 Hz, 1H), 3.29 (t, J= 6Hz, 2H), 3.38 (dd, J= 1 and 6 Hz, 1H), 4.05 (brs, 1H), 4.23 (t, J= 6Hz, 1H), 4.31 (t, J= 6Hz, 1H), 4.35 (dd, J= 2 and 6Hz, 1H), 4.64 (brs, 1H), 6.14 (brs, 1H), 6.57 (brd, J= 6Hz, 1H), 7.07 (brs, 1H), 7.34 (t, J= 6Hz, 2H), 7.42 (t, J= 6Hz, 2H), 7.69 (brt, J= 6Hz, 2H), 7.83 (d, J= 6Hz, 2H), 8.20 (brd, J= 6Hz, 1H).

$^{13}$C NMR (151 MHz, $CD_3CN$) δ 23.5, 29.0, 31.8, 42.1, 47.9, 50.9, 51.9, 56.2, 118.3, 121.0, 126.2, 126.2, 128.1, 128.8, 142.1, 144.9, 145.0, 157.8, 170.5, 174.6, 183.4.

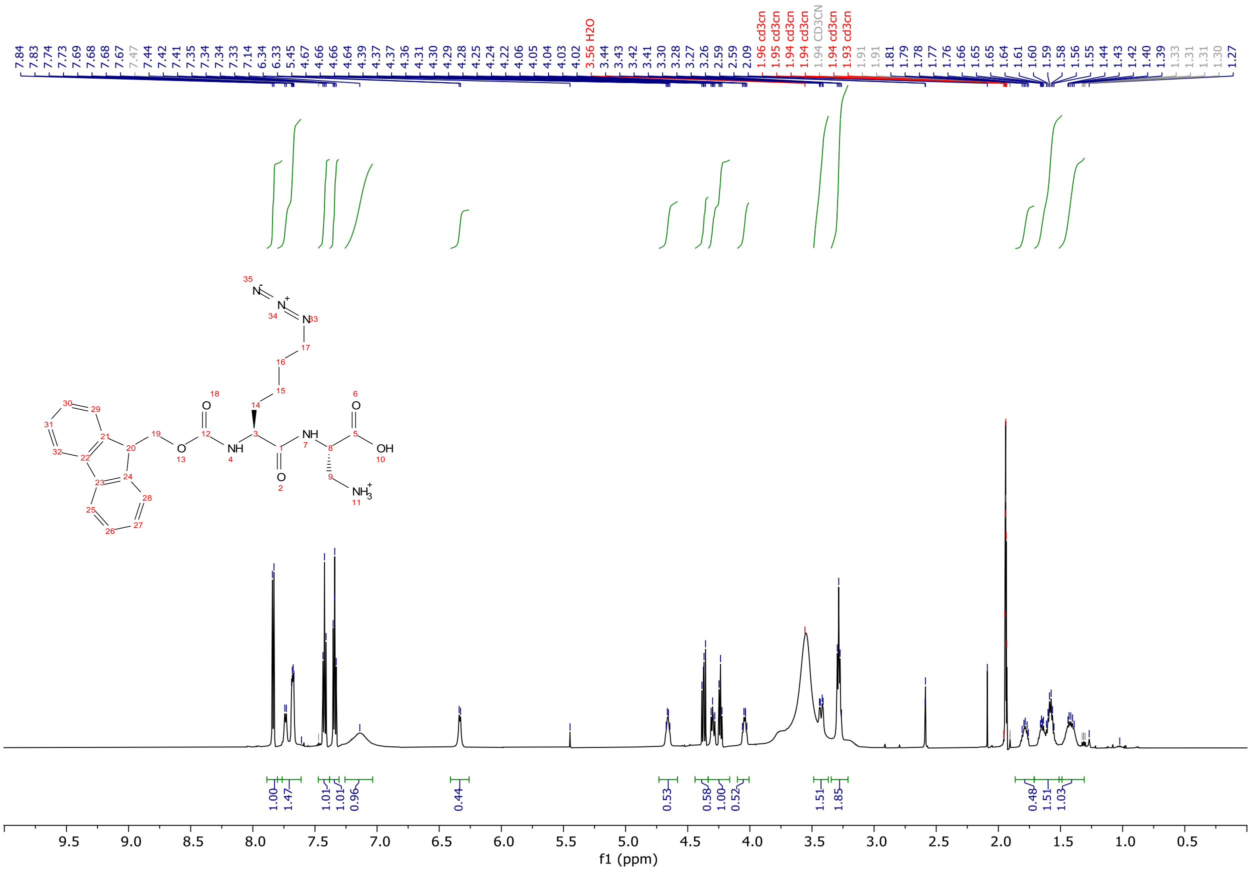

[1]H-NMR of **1n** in $CD_3CN$.

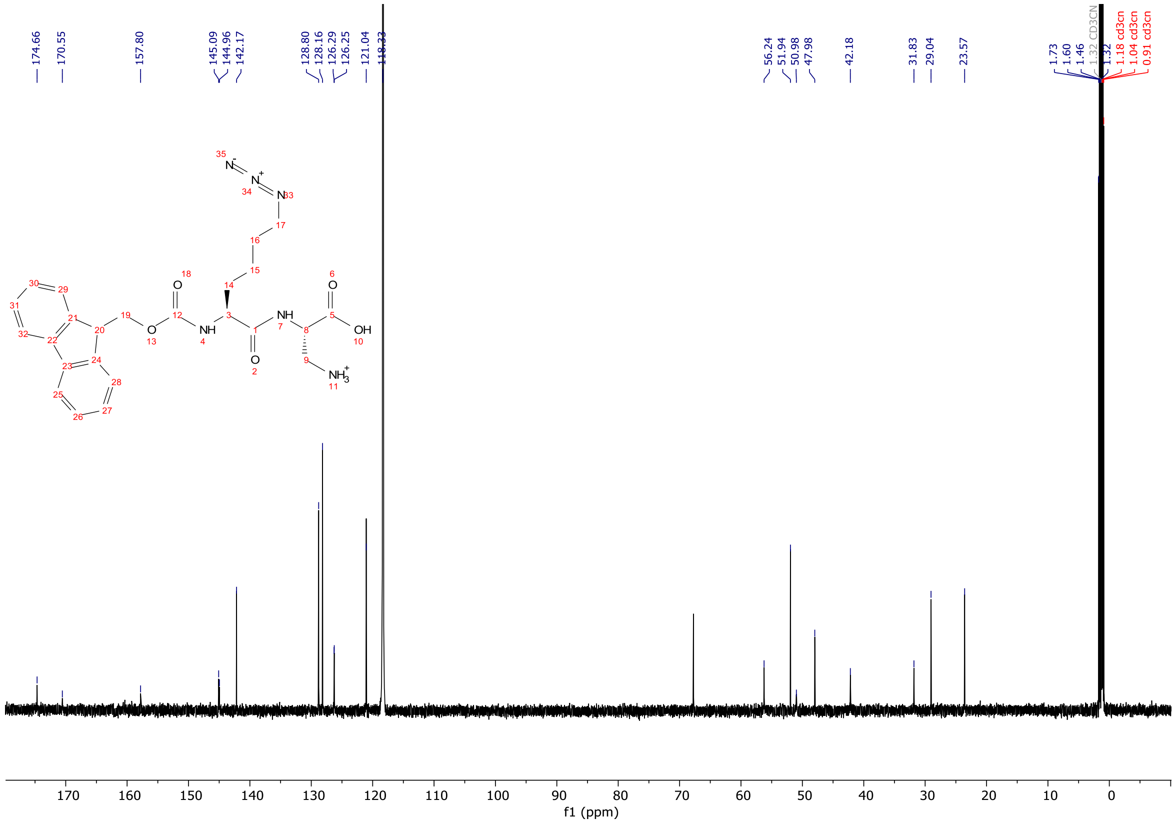

$^{13}$C-NMR of **1n** in $CD_3CN$.

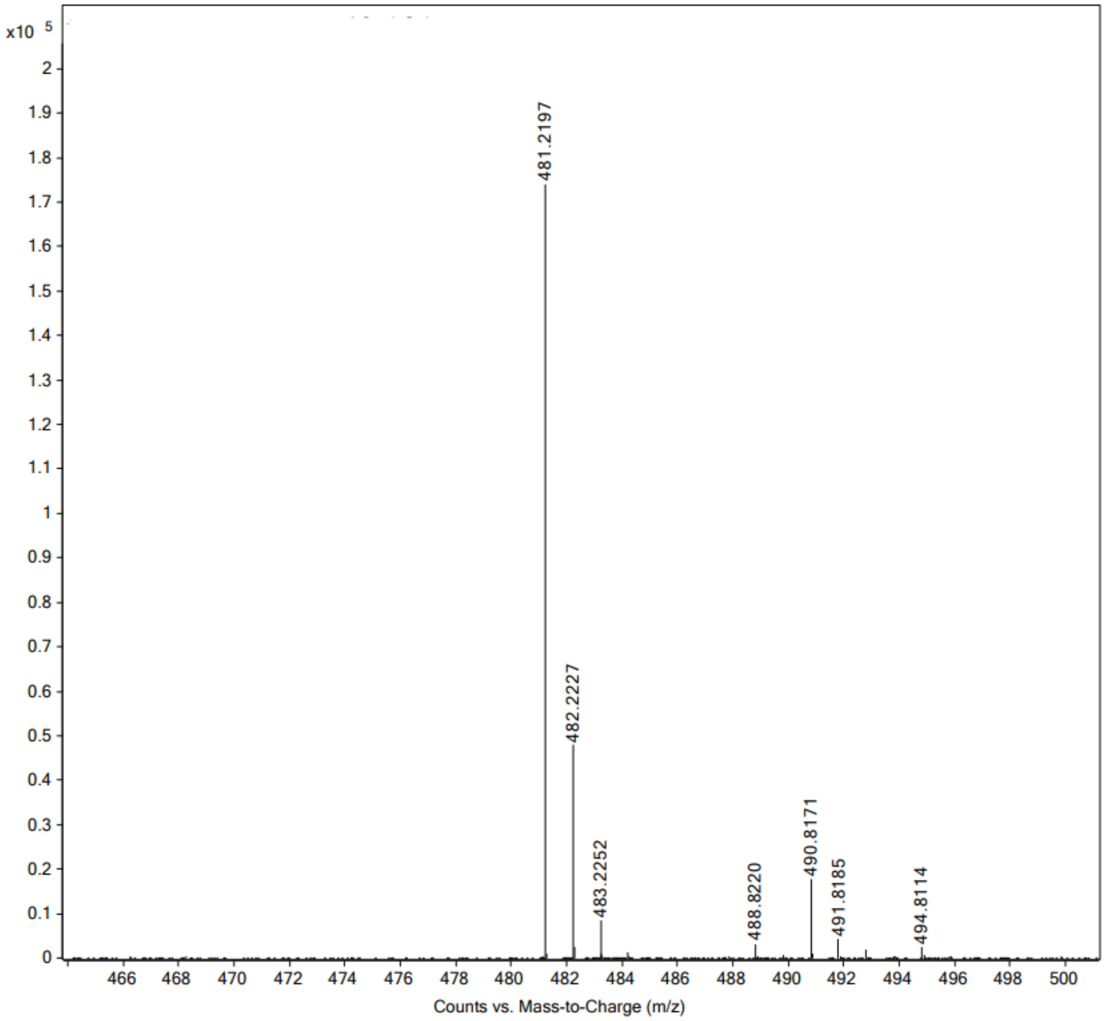


High resolution mass spectrum (ESI) of **1n**.

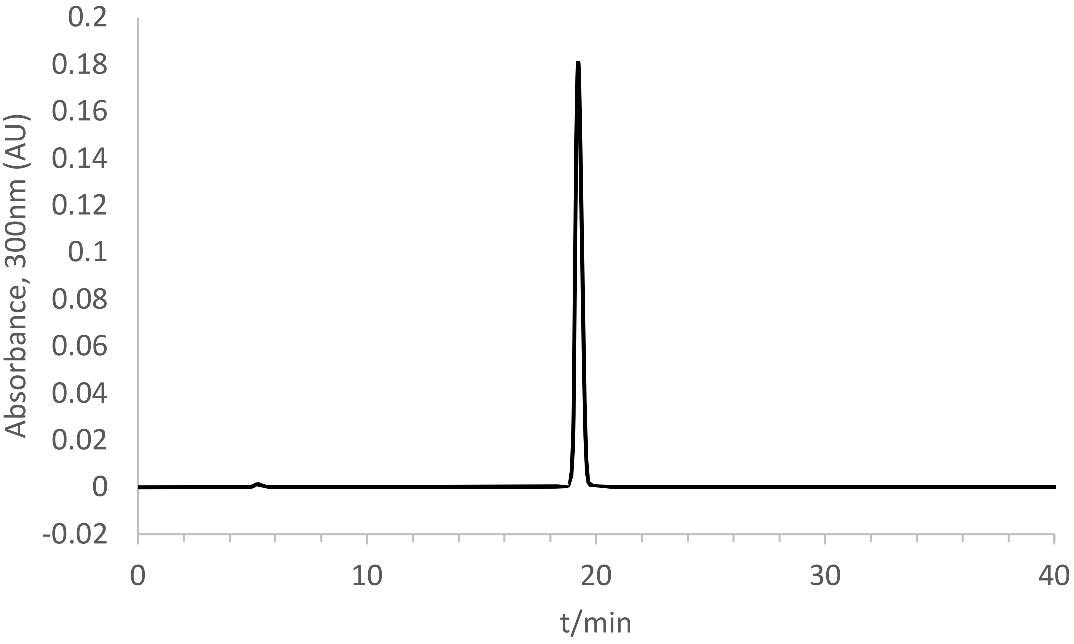


Analytical HPLC chromatogram of **1n**.

**Synthesis of 2p**.

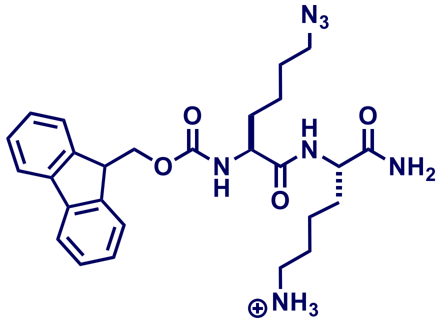


100 mg of Fmoc-(aminomethyl)-phenoxy resin (Rink amide resin) of loading capacity 0.48 mmol/g was swollen in DCM after which the Fmoc protecting group was removed using 20% piperidine in DMF for 30 min. The resin was washed using alternating DCM and MeOH and finally rinsed with DMF. Next, Fmoc-Lys (Boc)-OH was coupled to the resin using 4 equivalents of the amino acid and 4 equivalents each of Oxyma and DIC; the coupling reaction was done at room temperature with shaking for 2h. The Fmoc group was similarly removed using 20% piperidine and following wash steps as described above, Fmoc-Lys($N_3$)-OH was similarly coupled to Lys (Boc) to make Fmoc-Lys($N_3$)-Lys(Boc)-Rink amide resin. The peptide was cleaved off the resin using 95% TFA:2.5% TIPS: 2.5% $H_2O$ and purified by preparative RP-HPLC on a on a Phenomenex Luna 10 µm C8(2) 100 Å (250 x 21.2 mm) using a 20% to 100% linear gradient of MeCN in $H_2O$ each containing 0.1% TFA at 10 mL/min. ESI-MS $[M+H]^+$ m/z calcd. 522.2823 for $C_{27}H_{36}N_7O_4$, obsd. 522.2825 and $[M+Na]^+$ 544.2651. $^1H$ NMR (600 MHz, $CD_3CN$) δ 1.30-1.40 (m, 2H), 1.50-1.65 (m, 3H), 1.72-1.79 (m, 1H), 2.86 (m, 1H), 3.27 (t, J=6 Hz, 1H), 4.01 (dd, J=6, 8 Hz, 1H), 4.21- 4.24 (m, 1H), 4.28 (dd, J= 6Hz, 1H), 4.36 (dd, J= 6Hz, 1H), 7.33 (t, J= 6Hz, 2H), 7.41 (t, J= 6Hz, 2H), 7.66 (brt, J= 6Hz, 2H), 7.82 (d, J= 6Hz, 2H).

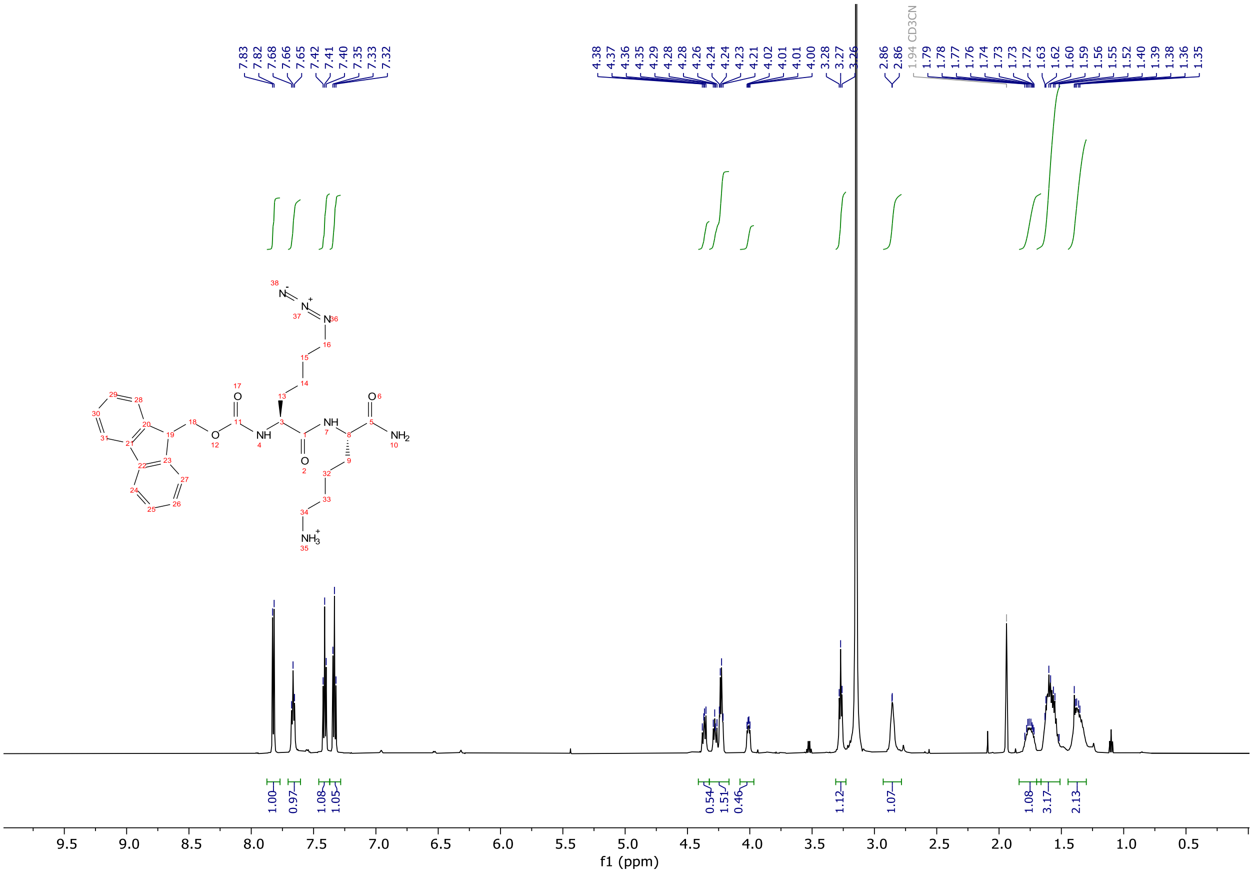

[1]H-NMR of **2p** in $CD_3CN$.

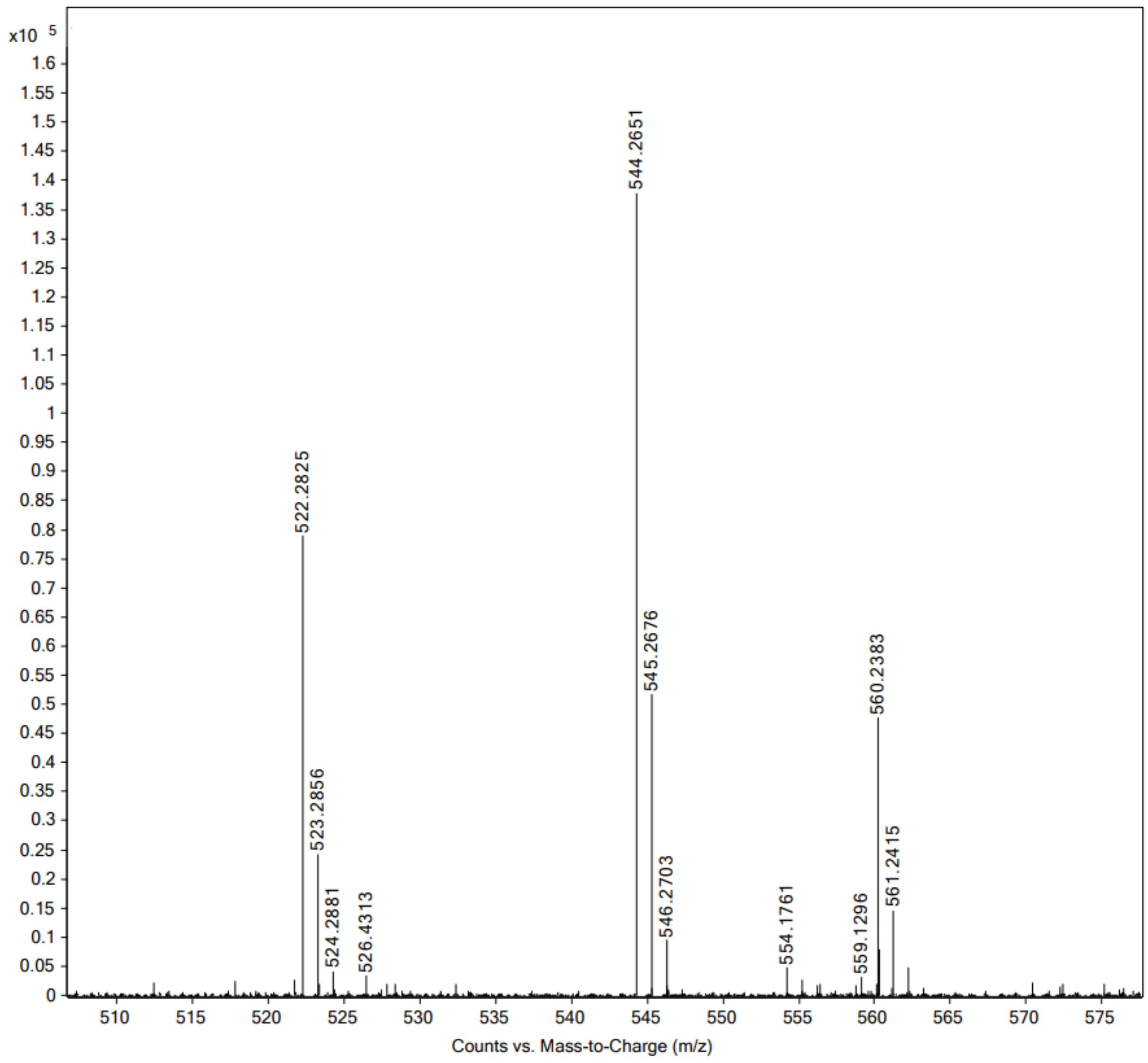


High resolution mass spectrum (ESI) of **2p**.

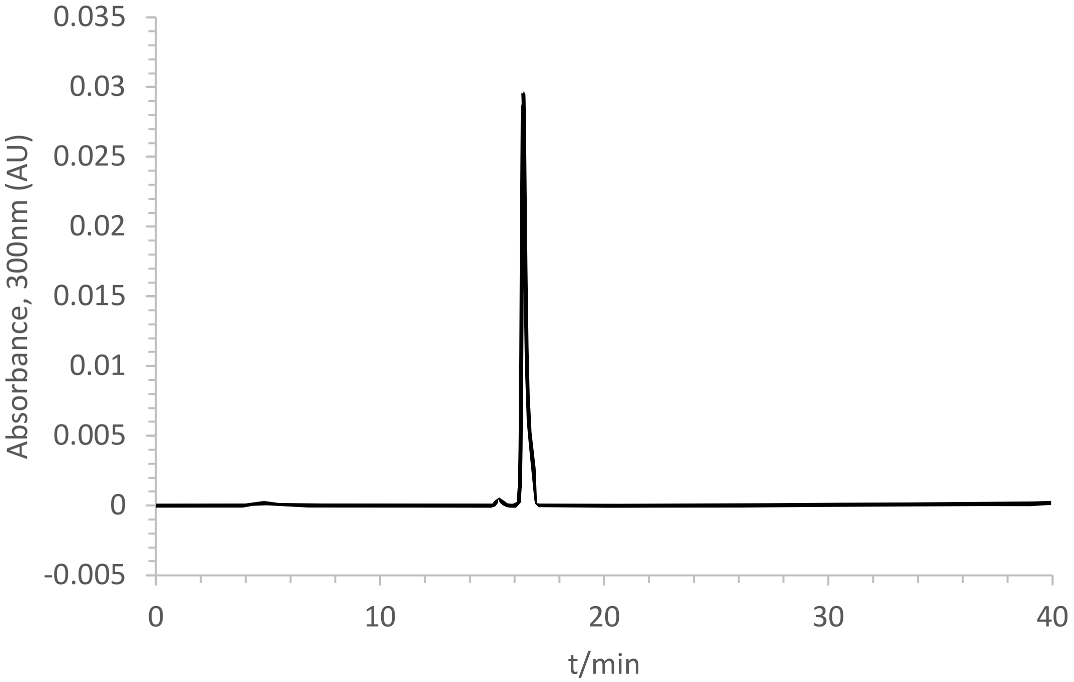


Analytical HPLC chromatogram of **2p**.

**Synthesis of 2n**.

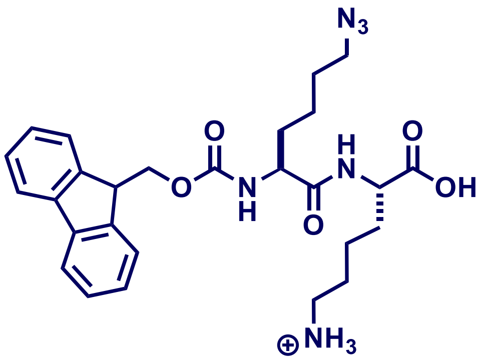


50 mg of 2-chlorotrityl chloride resin (Rink amide resin) of loading capacity 1.5 mmol/g was swollen in dry DCM in a peptide synthesis vessel. Fmoc-Lys (Boc)-OH was dissolved in dry DCM (1 eq.) and added to the resin and shaken for 5 min after which 2 eq of DIEA was added and incubated for 1h with shaking at room temperature. The resin was washed 3X with dry DCM, washed 1X with MeOH, and rinsed with DMF. Next, the Fmoc protecting group was removed using 20% piperidine in DMF for 30 min. The resin was washed using alternating DCM and MeOH and finally rinsed with DMF. Next, Fmoc-Lys(Boc)-OH was coupled to the resin using 4 equivalents of the amino acid and 4 equivalents each of Oxyma and DIC; the coupling reaction was done at room temperature with shaking for 2 h. The Fmoc group was similarly removed using 20% piperidine and Following wash steps as described above, Fmoc-Lys($N_3$)-OH was similarly coupled to Lys(Boc) to make Fmoc-Lys($N_3$)-Lys(Boc)-trityl chloride resin. The peptide was cleaved off the resin using 95% TFA:2.5% TIPS: 2.5% $H_2O$ and purified by preparative RP-HPLC on a on a Phenomenex Luna 10 µm C8(2) 100 Å (250 x 21.2 mm) using a 20 to 100% linear gradient of MeCN in $H_2O$ each containing 0.1% TFA at 10 mL/min. ESI-MS $[M+H]^+$ m/z calcd. 523.2663 for $C_{27}H_{35}N_6O_5$, obsd. 523.2669. $^1$H NMR (600 MHz, $CD_3CN$) δ 1.30-1.40 (m, 2H), 1.50-1.65 (m, 3H), 1.72-1.79 (m, 1H), 2.90 (t, J=6 Hz, 1H), 3.26 (t, J=6 Hz, 1H), 4.09 (dd, J=6, 8 Hz, 1H), 4.21- 4.24 (m, 1H), 4.28 (dd, J= 6Hz, 1H), 4.35 (dd, J= 6Hz, 1H), 6.31 (brs, 0.5H), 7.16 (brs, 1H), 7.33 (t, J= 6Hz, 2H), 7.41 (t, J= 6Hz, 2H), 7.67 (brt, J= 6Hz, 2H), 7.82 (d, J= 6Hz, 2H). $^{13}$C NMR (151 MHz, $CD_3CN$) δ 22.7, 23.5, 26.9, 29.0,

31.1, 32.4, 40.4, 47.9, 51.9, 53.0, 55.7, 67.4, 120.9, 120.9, 126.2, 126.2, 128.0, 128.1, 128.6, 142.0, 144.9, 145.0, 157.3.

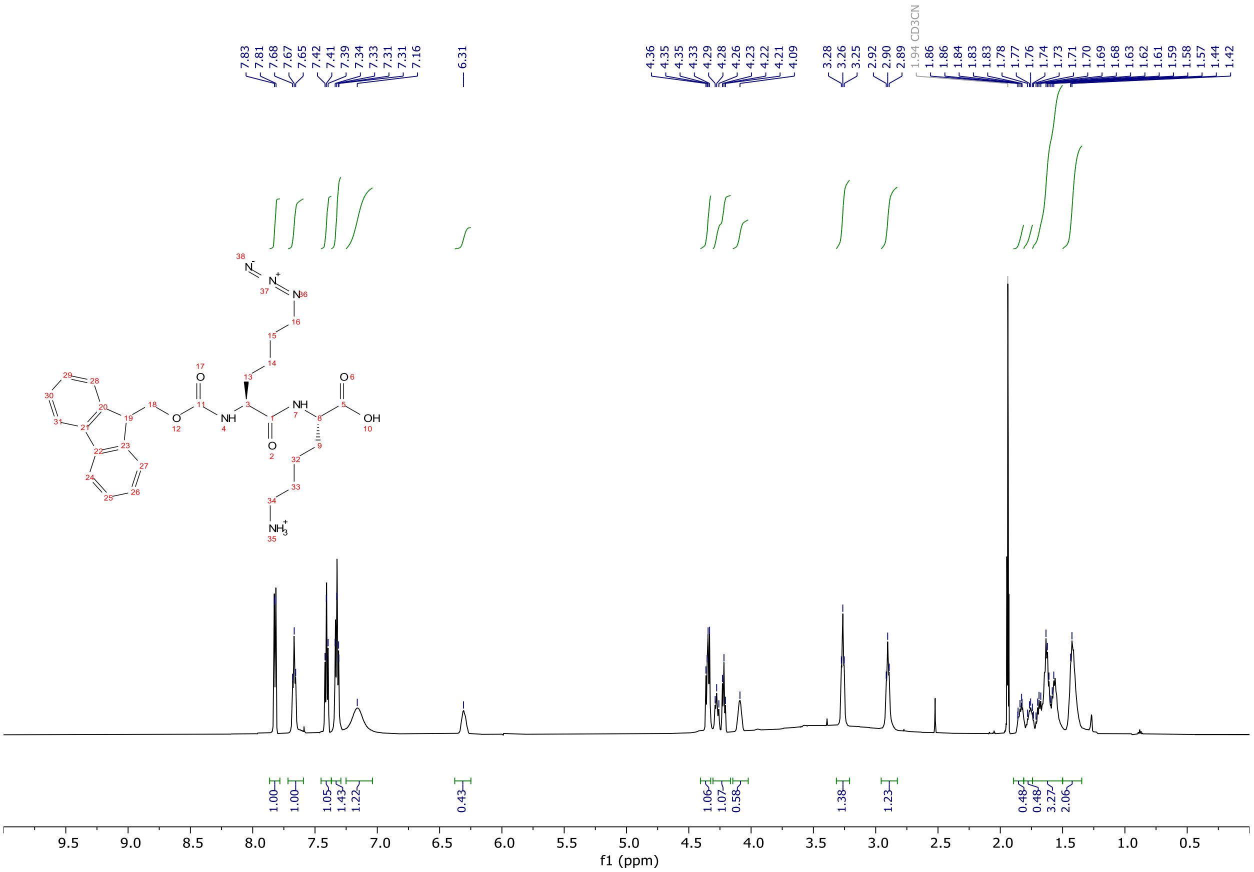

[1]H-NMR of **2n** in $CD_3CN$.

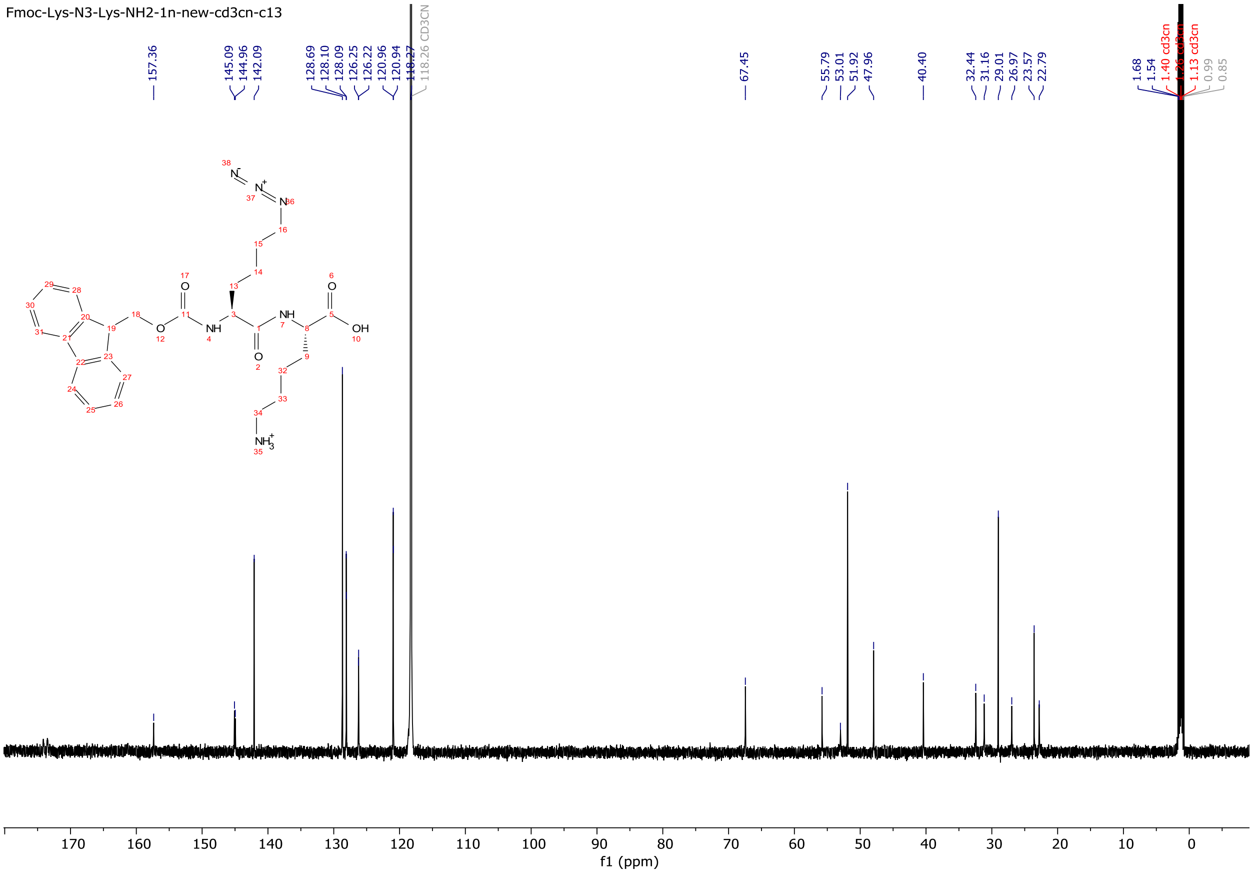


$^{13}$C-NMR of **2n** in $CD_3CN$.

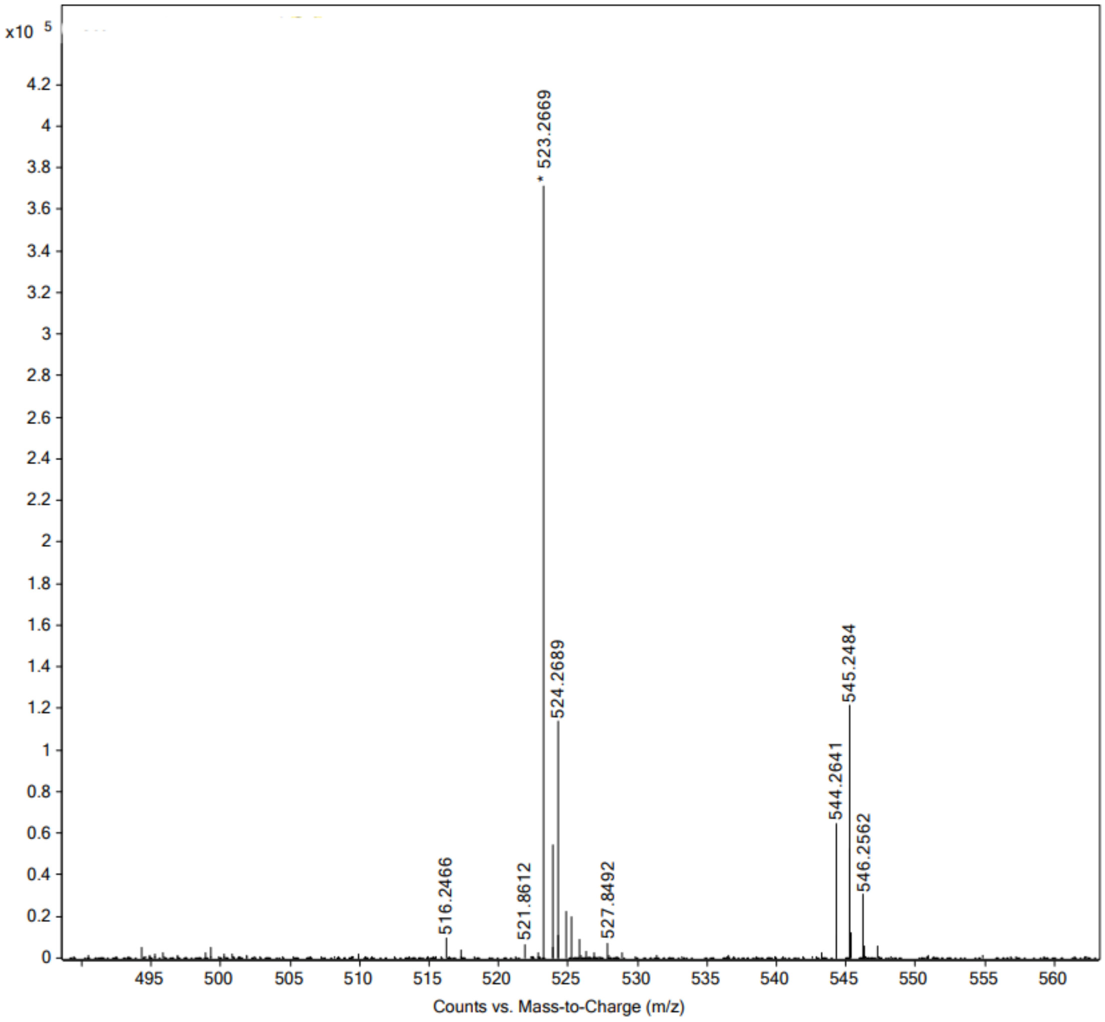


High resolution mass spectrum (ESI) of **2n**.

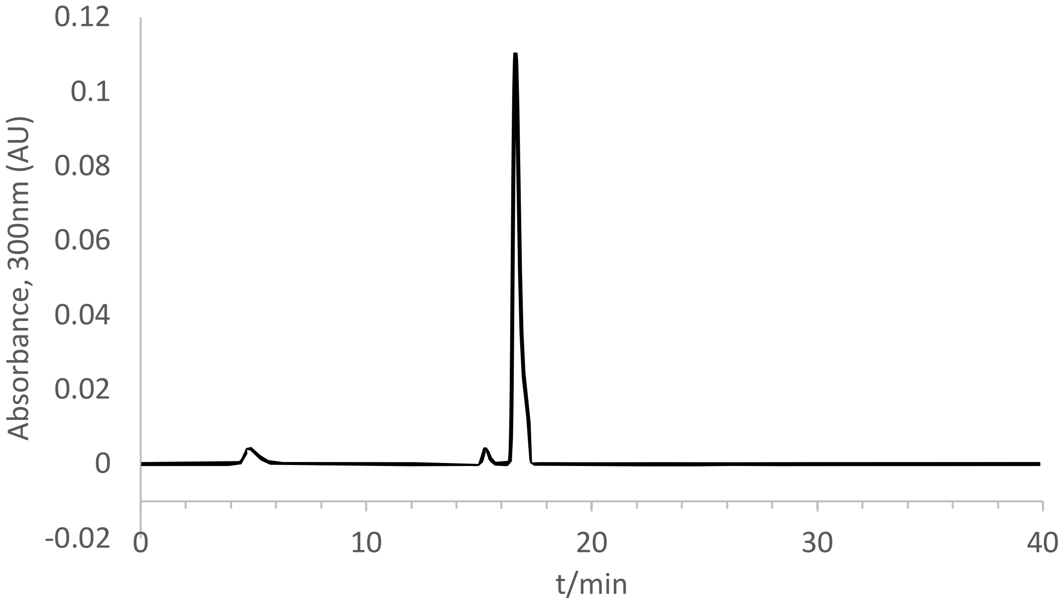


Analytical HPLC chromatogram of **2n**.

**Synthesis of 3p**.

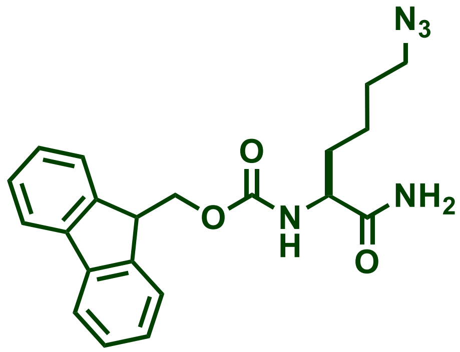


100 mg of Fmoc-aminomethyl)-phenoxy resin (Rink amide resin) of loading capacity 0.48 mmol/g was swollen in DCM after which the Fmoc protecting group was removed using 20% piperidine in DMF for 30 min. The resin was washed using alternating DCM and MeOH and finally rinsed with DMF. Fmoc-Lys($N_3$)-OH was coupled to the resin using 4 equivalents of the amino acid and 4 equivalents each of Oxyma and DIC; the coupling reaction was done at room temperature with shaking for 2 h. The amidated amino acid was cleaved off the resin using 95% TFA:2.5% TIPS: 2.5% $H_2O$ and purified by preparative RP-HPLC on a on a Phenomenex Luna 10 µm C8(2) 100 Å (250 x 21.2 mm) using a 20 to 100% linear gradient of MeCN in $H_2O$ each containing 0.1% TFA at 10 mL/min. ESI-MS $[M+H]^+$ m/z calcd. 394.1874 for $C_{21}H_{24}N_5O_3$ obsd. 394.1876. $^1H$ NMR (600 MHz, $CD_3CN$) δ 1.30-1.40 (m, 1H), 1.50-1.65 (m, 2H), 1.72-1.82 (m, 1H), 2.90 (brs), 3.29 (t, J=6 Hz, 1H), 3.99 (brs, 1H), 4.24 (t, J= 6Hz, 1H), 4.33 (t, J= 6Hz, 1H), 4.38 (dd, J= 6Hz, 1H), 5.78 (brs, 0.5H), 5.91 (brs, 0.5H), 6.33 (brs, 0.5H), 7.16 (brs, 1H), 7.34 (dt, J= 1 and 6Hz, 2H), 7.42 (t, J= 6Hz, 2H), 7.68 (brt, J= 6Hz, 2H), 7.83 (d, J= 6Hz, 2H).

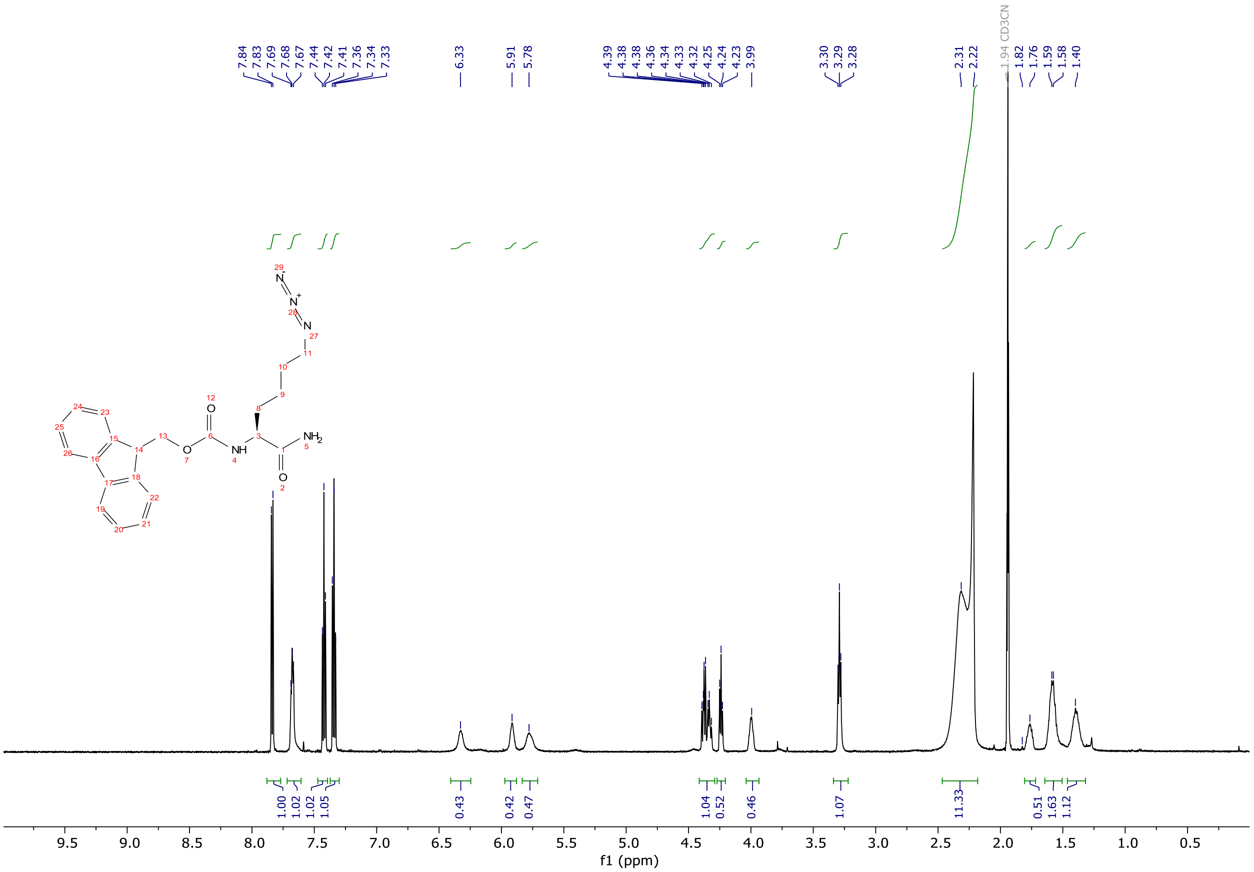

[1]H-NMR of **3p** in $CD_3CN$.

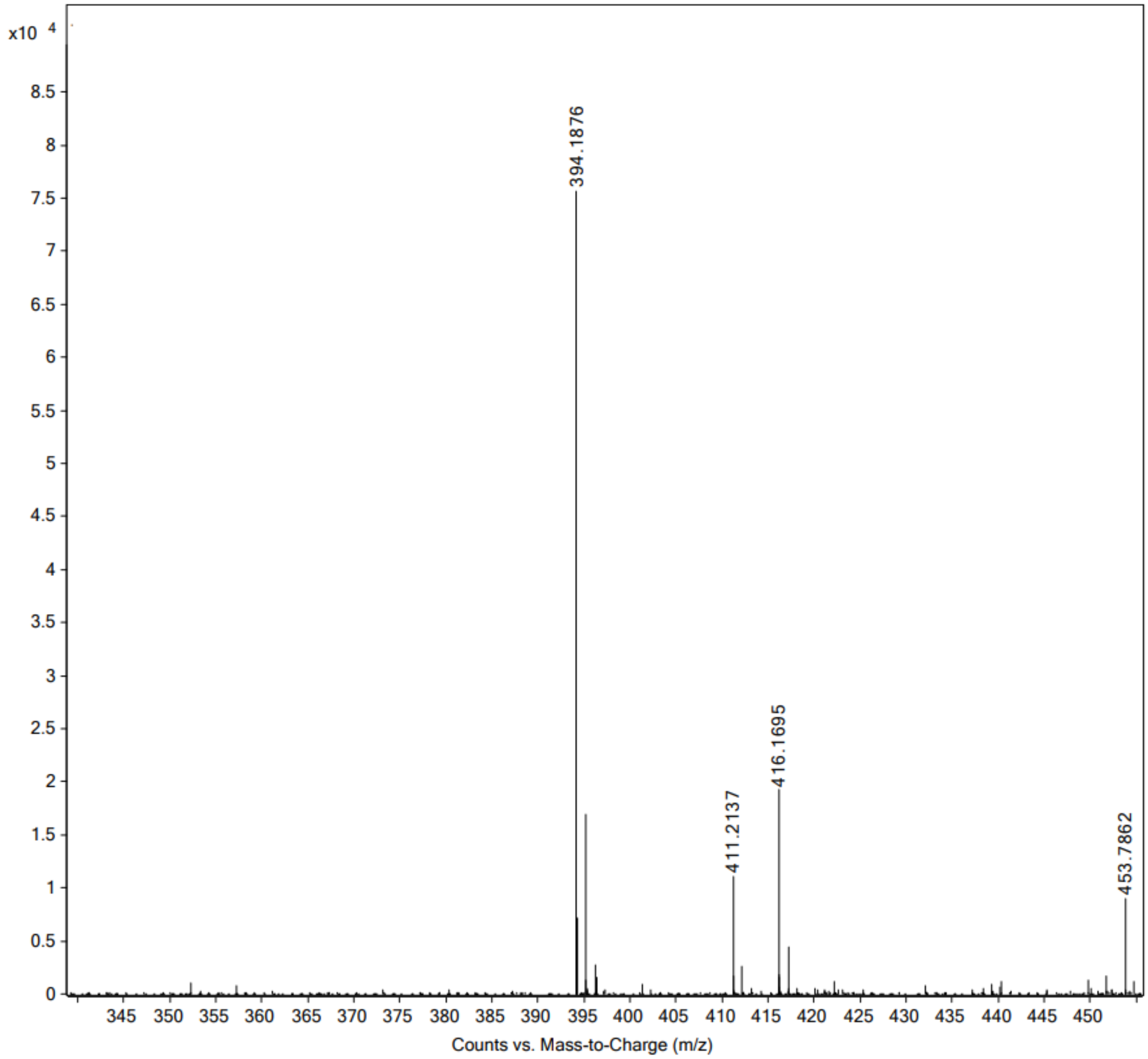


High resolution mass spectrum (ESI) of **3p**.

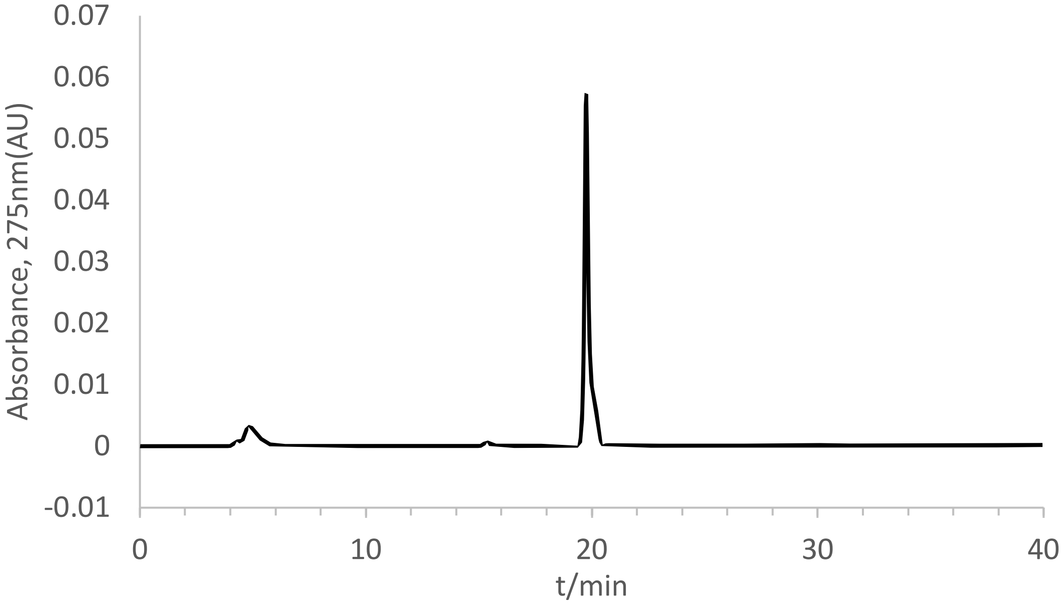


Analytical HPLC chromatogram of **3p**.

**Synthesis of 4a**.

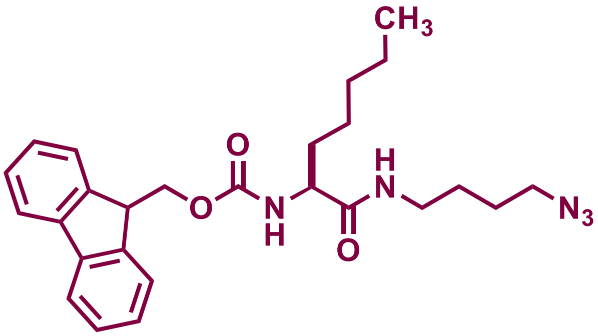


4-azidobutylamine was coupled to Fmoc-Nle-OH using 2 equivalents of the amino acid, 2 eq. of HBTU and 4 eq. DIEA in DMF. The coupling reaction was done at room temperature with agitation for 2 h to make Fmoc-Nle-butyl azide. Excess DMF solvent was removed by rotary evaporation as a DMF-toluene azeotrope until dry powder was obtained. The powder was resuspended in nanopure water to make a white suspension. The suspension was centrifuged at 4000 g for 10 min to separate the coupling reagents (aqueous layer) from the product (pellet). The pellet was resuspended in 20% HPLC grade acetonitrile in water (0.1% TFA) and injected into preparative RP-HPLC for purification on a Phenomenex Luna 10 µm C8(2) 100 Å (250 x 21.2 mm) using a 20 to 100% linear gradient of MeCN in $H_2O$ each containing 0.1% TFA at 10 mL/min. ESI-MS $[M+H]^+$ m/z calcd. 450.2500 for $C_{25}H_{32}N_5O_3$, obsd. 450.2500 $^1$H NMR (600 MHz, $CD_3CN$) δ 0.89 (t, 3H, CH3),1.20 (m,1H), 1.25-1.39 (m, 2H), 1.47-1.59 (m, 4H), 1.72 (m, 1H), 3.16 (dd, J= 6 and 9 Hz, 2H), 3.28 (t, J= 6 Hz, 1H), 3.95 (brs, 1H), 4.23 (t, J= 6Hz, 1H), 4.32 (brt, J= 6Hz, 1H), 4.36 (dd, J= 2 and 6Hz, 1H), 5.90 (brs, 1H), 6.66 (brs, 1H), 7.34 (t, J= 6Hz, 2H), 7.42 (t, J= 6Hz, 2H), 7.68 (brt, J= 6Hz, 2H), 7.83 (d, J= 6Hz, 2H).

.

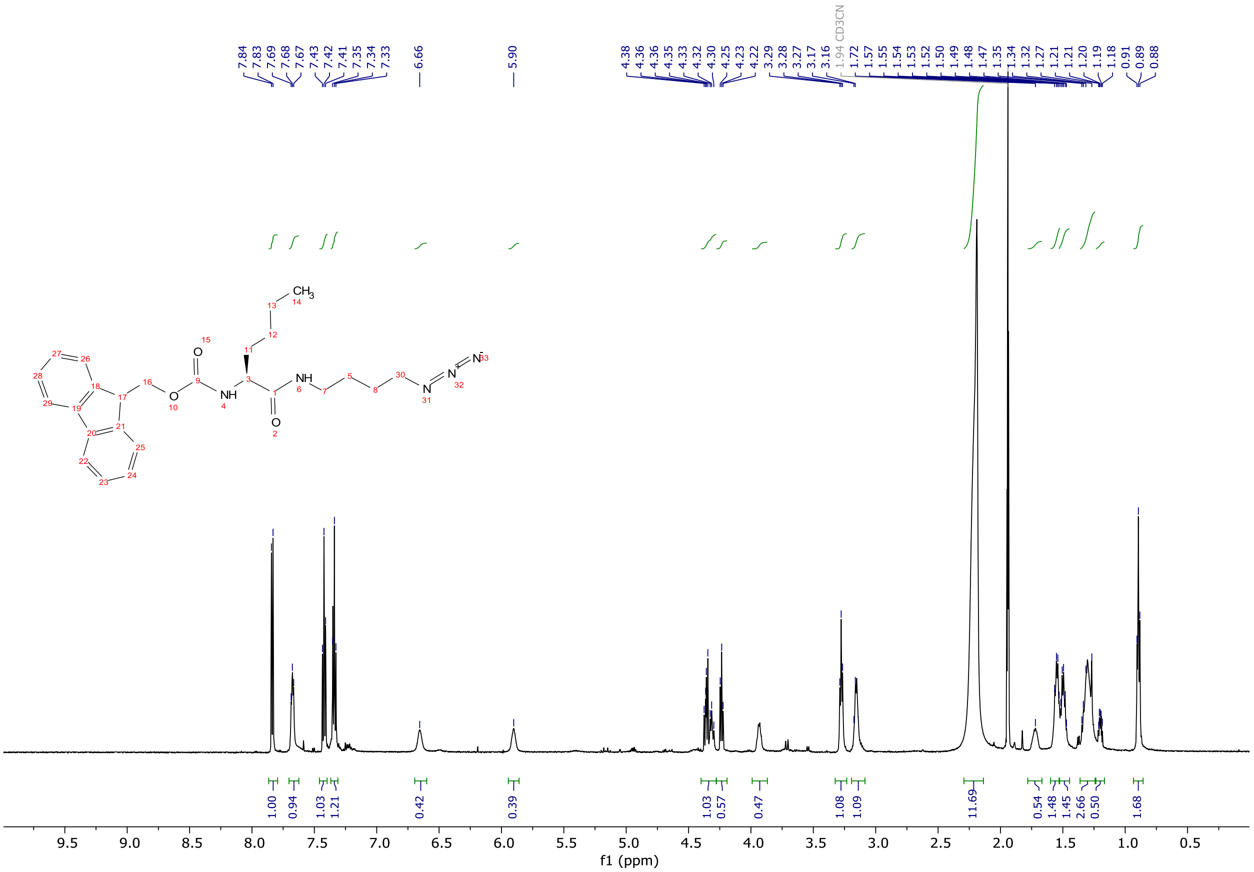

[1]H-NMR of **4a** in $CD_3CN$.

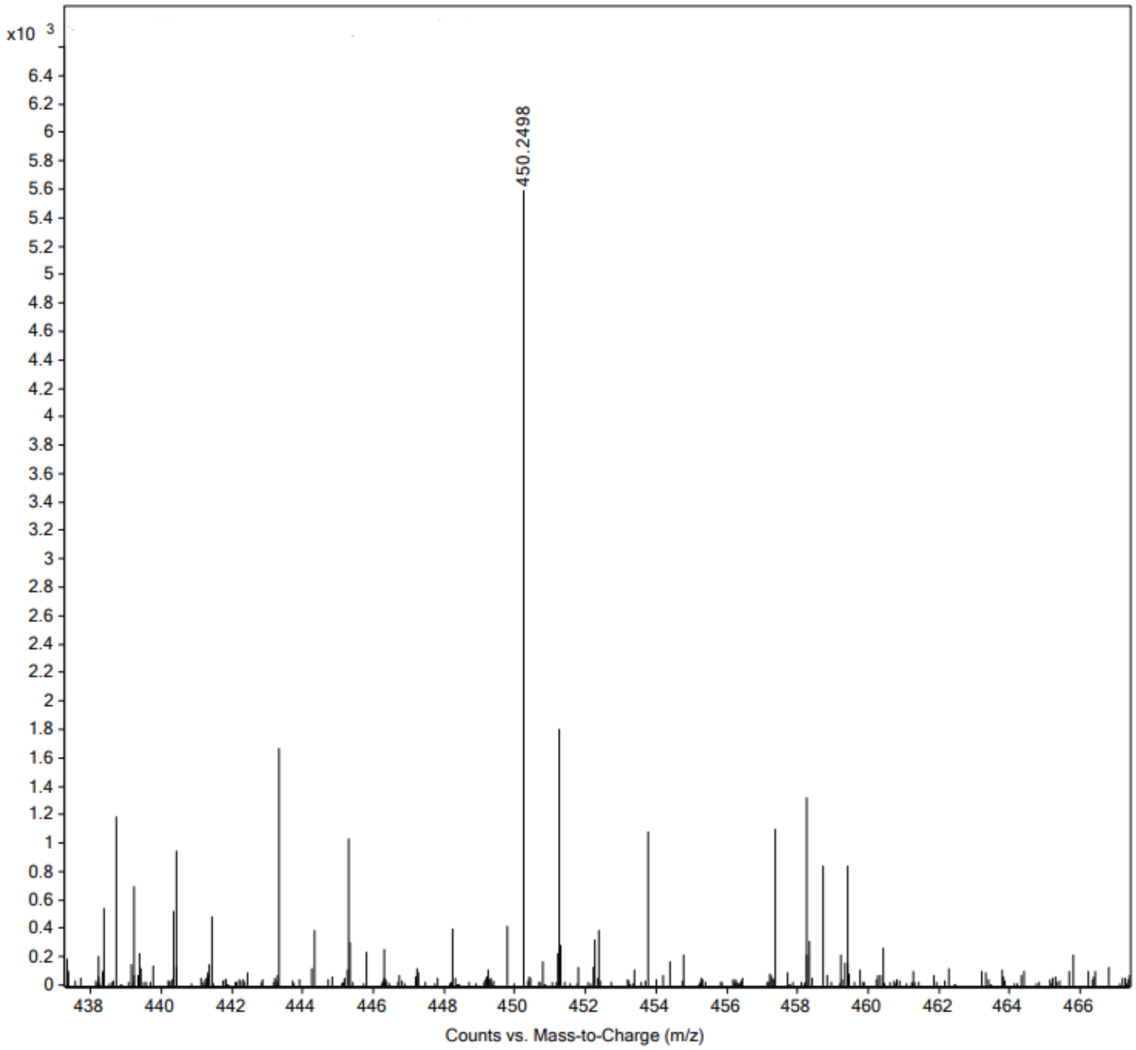


High resolution mass spectrum (ESI) of **4a.**

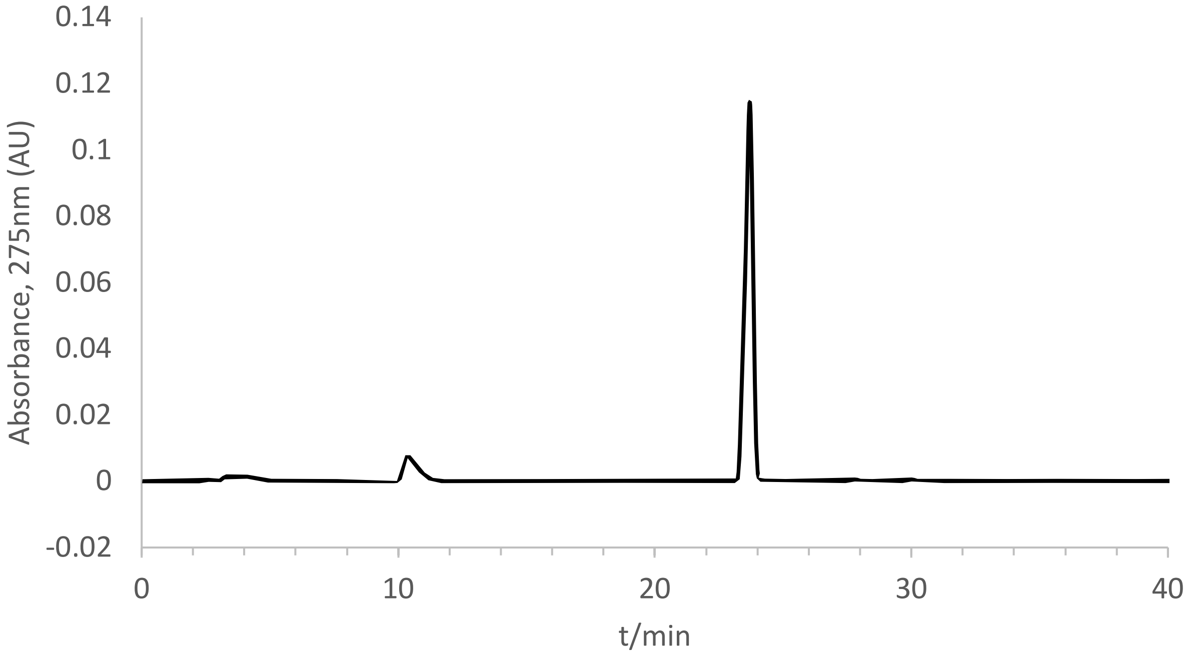


Analytical HPLC chromatogram of **4a**.

**Synthesis of 4b, 4c, 4d, and 4e or Fmoc-Lys $(Me)_x$- butyl azide where x = 0, 1, 2, 3**

4-azidobutylamine was coupled to either Fmoc-Lys (Boc)-OH or Fmoc-Lys (Boc, Me)-OH or Fmoc-Lys$(Me)_2$-OH.HCl or Fmoc-Lys$(Me)_3$-$OH^+Cl^-$ using 2 equivalents of the amino acid, 2 eq. of HBTU and 4 eq. DIEA in DMF. The coupling reaction was done at room temperature with agitation for 2 h to link the two building blocks by an amide bond. Excess DMF solvent was removed by rotary evaporation as a DMF-toluene azeotrope until dry powder was obtained. The powder was resuspended in nanopure water to make a white suspension. The suspension was centrifuged at 4000 g for 10 min to separate the coupling reagents (aqueous layer) from the product (pellet). The pellet was resuspended in 20% HPLC grade acetonitrile in water (0.1% TFA) and injected into preparative RP-HPLC for purification on a Phenomenex Luna 10 μm C8(2) 100 Å (250 x 21.2 mm) using a 20 to 100% linear gradient of MeCN in $H_2O$ each containing 0.1% TFA at 10 mL/min.

**4b.**

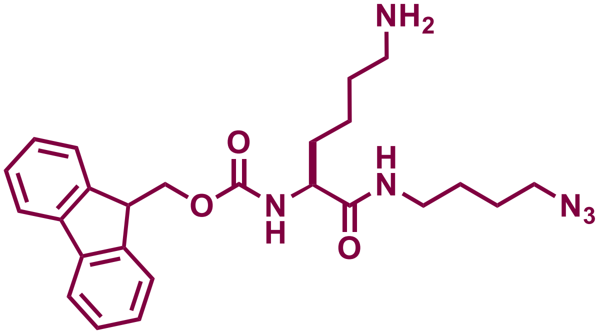


ESI-MS $[M+H]^+$ m/z calcd. 465.2609 for $C_{25}H_{33}N_6O_3$, obsd. 465.2609. $^1H$ NMR (600 MHz, $cdcl_3$) δ 1.30-171 (m, 3H), 2.91 (brs, 1H), 3.14 (m, 2H), 4.09 (t, J= 6Hz, 1H), 4.27 (m, 1H), 6.19 (brd, J=6 Hz, 1H), 7.23 (dt, J= 1 and 6Hz, 2H), 7.33 (t, J= 6Hz, 2H), 7.51 (brd, J= 6Hz, 2H), 7.69 (d, J= 6Hz, 2H), 7.99 (brs, 1H). $^{13}C$ NMR (151 MHz, $cdcl_3$) δ 22.15, 26.01, 26.39, 26.69, 31.90, 38.94, 39.46, 46.89, 50.81, 54.55, 67.28, 120.00, 125.04, 127.06, 127.80, 141.19, 143.62, 156.67, 161.57, 161.80, 172.27.

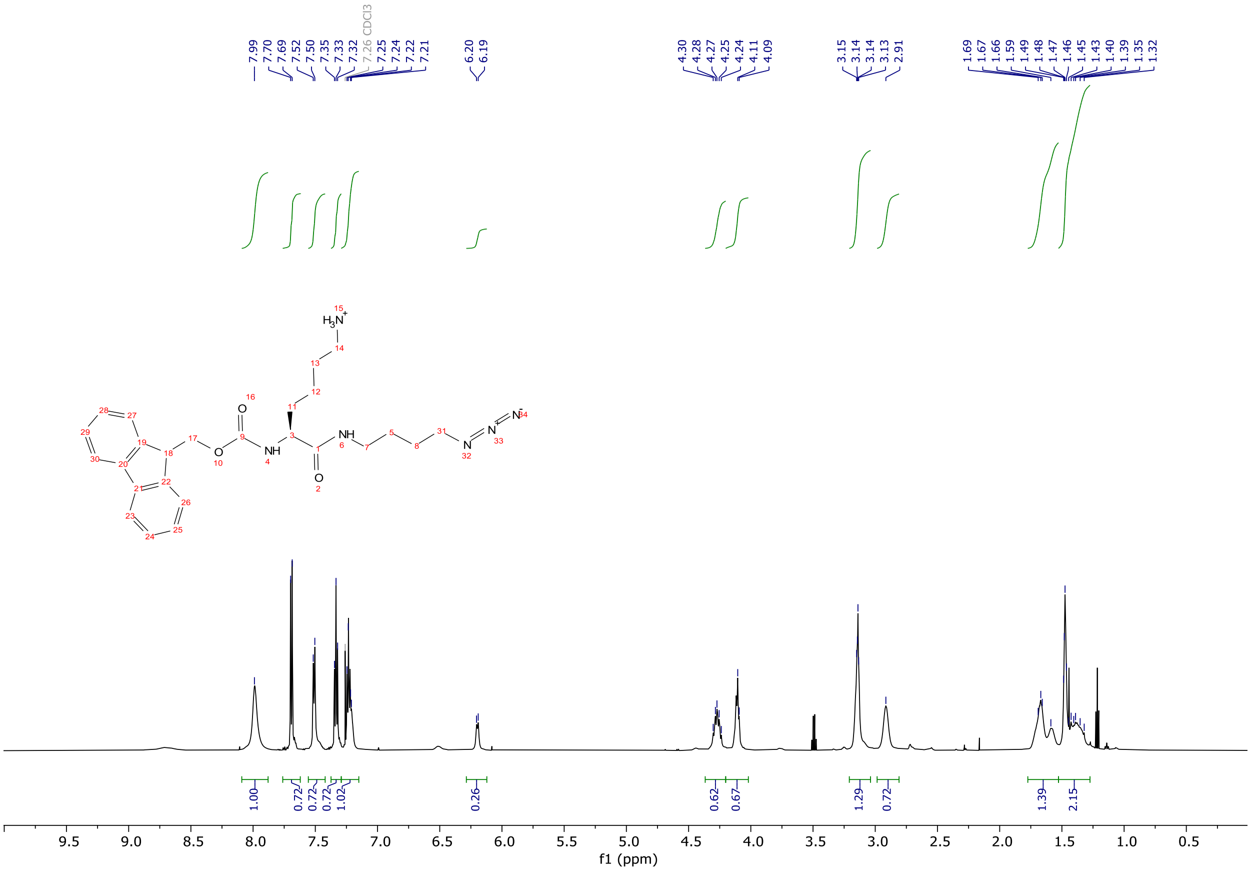

[1]H-NMR of **4b** in $CDCl_3$.

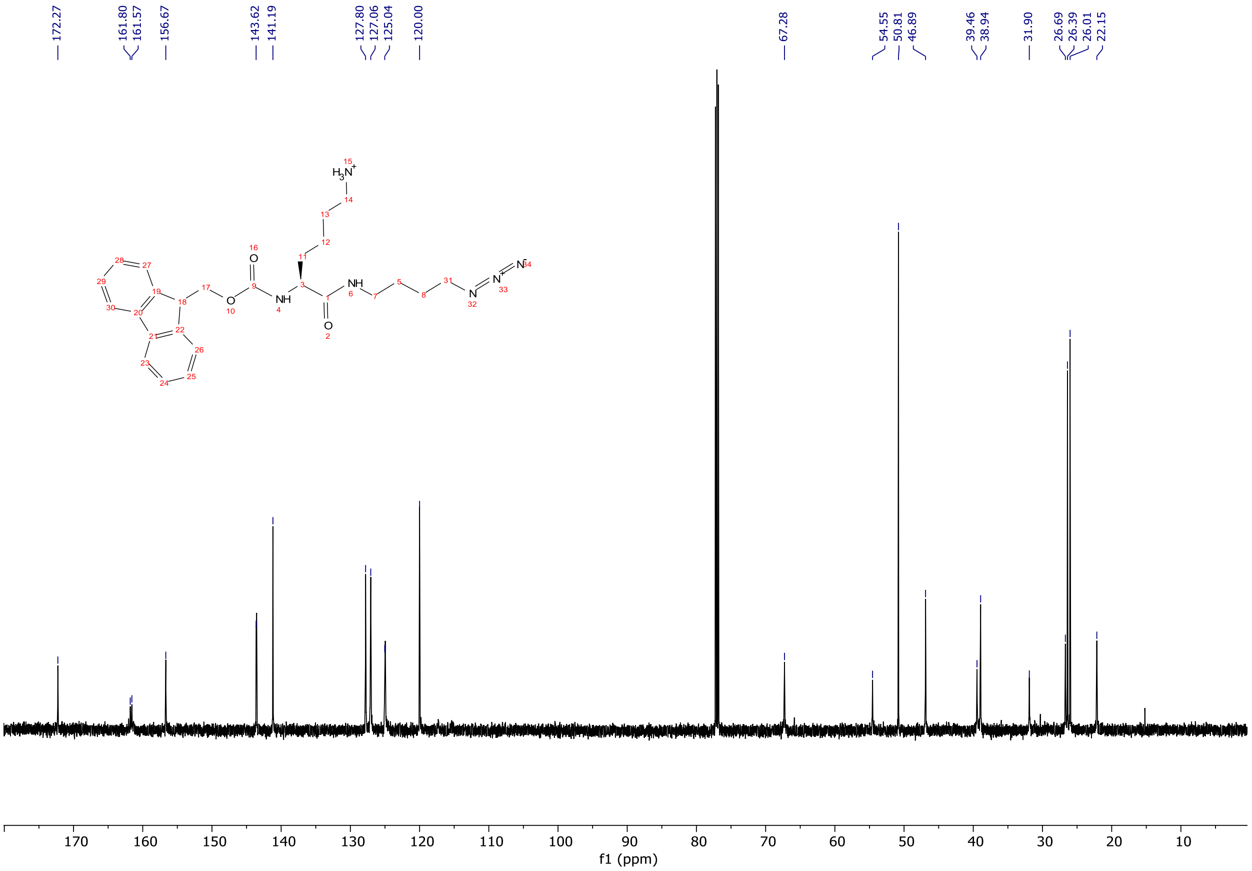


$^{13}$C-NMR of **4b** in $CDCl_3$.

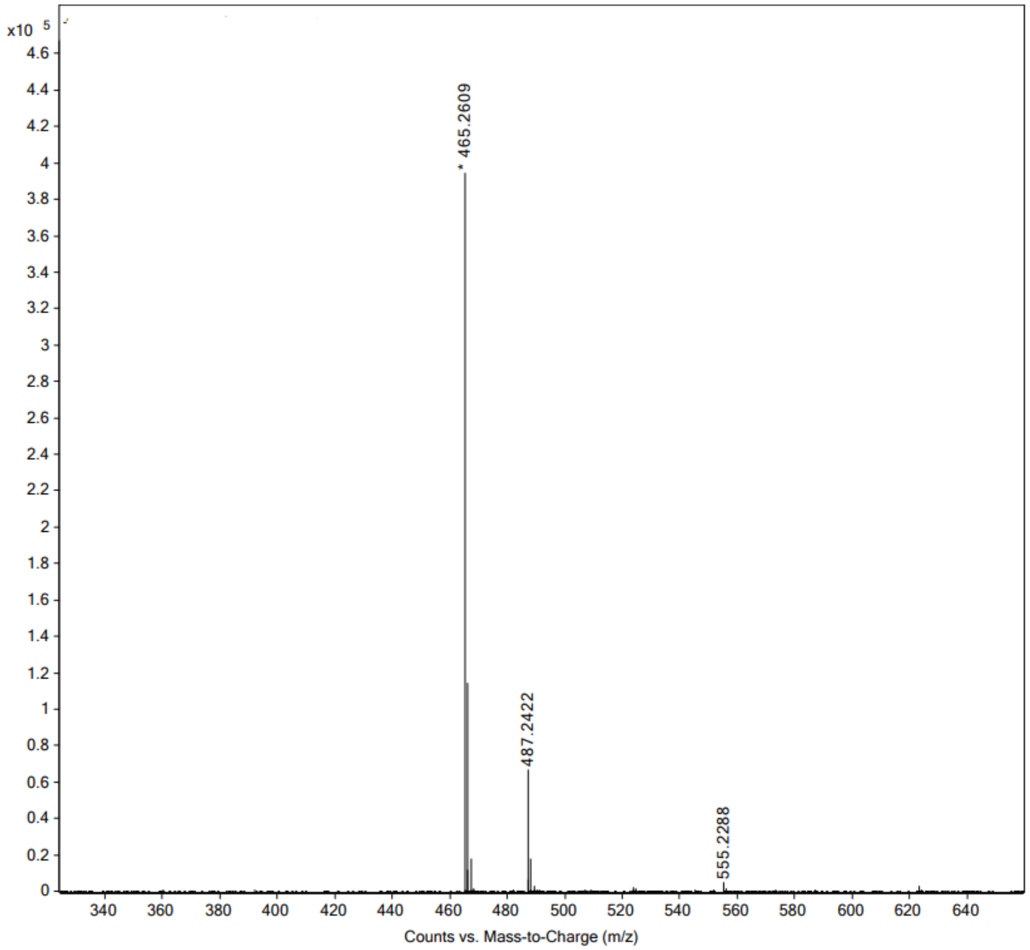


High resolution mass spectrum (ESI) of **4b**.

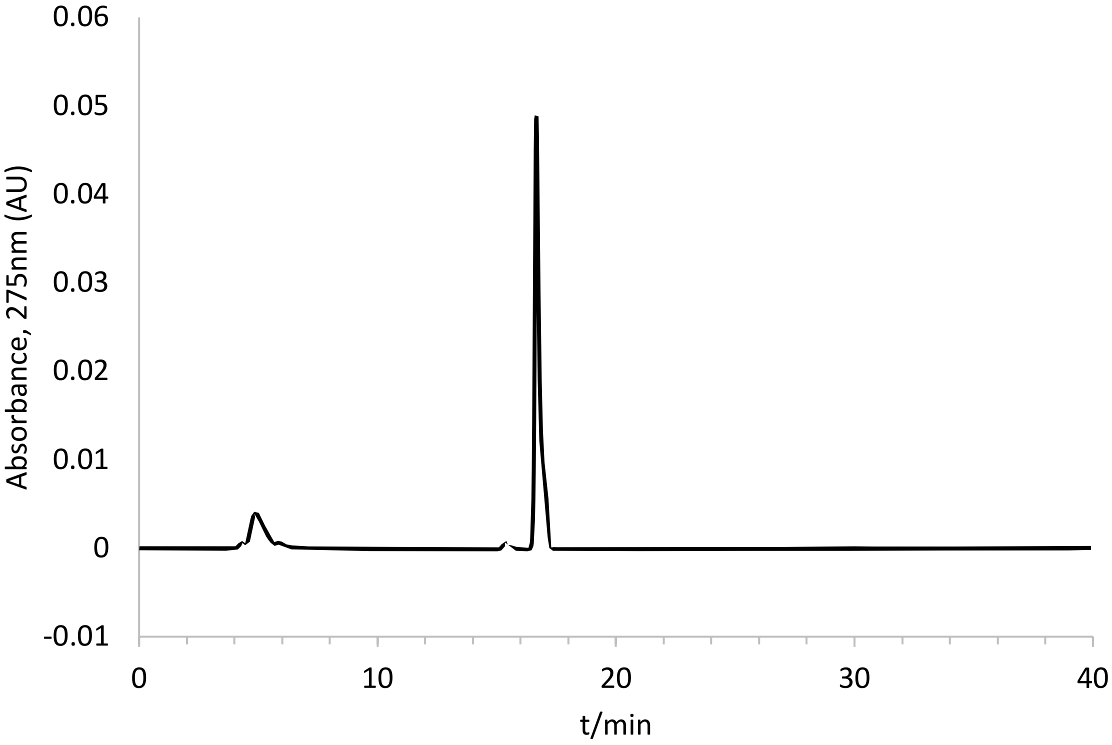


Analytical HPLC chromatogram of **4b**.

**4c.**

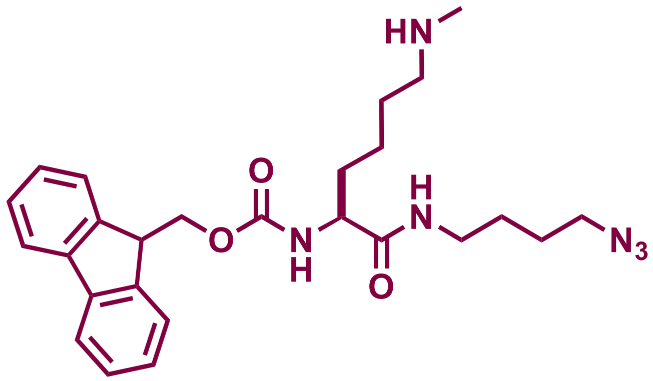

ESI-MS $[M+H]^+$ m/z calcd. 479.2765 for $C_{26}H_{35}N_6O_3$, obsd. 479.2779. $^1$H NMR (600 MHz, $CDCl_3$) δ 1.43 (m, 1H), 1.54 (br, 2H), 1.75-1.82 (m, 3H), 2.88 (m, 1H), 2.99 (m, 1H), 3.21 (brs, 1H), 4.17 (q, J= 6Hz, 2H), 4.30 (m, 1H), 6.01 (brd, J=6 Hz, 1H), 7.24 (brs, 1H), 7.27 (t, J= 6Hz, 2H), 7.38 (t, J= 6Hz, 2H), 7.54 (brd, J= 6Hz, 2H), 7.74 (d, J= 6Hz, 2H), 9.05 (brs, 1H), 9.39 (brs, 1H).

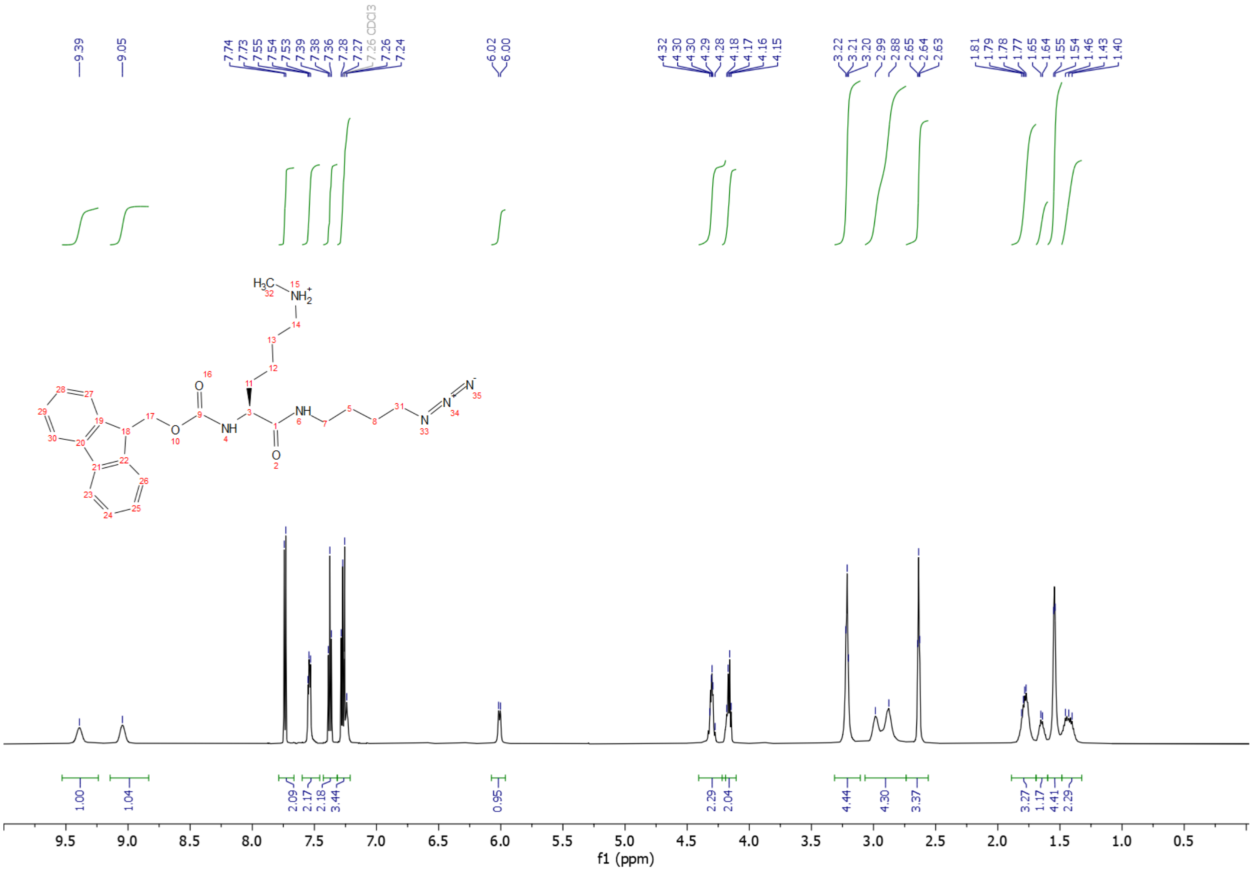

[1]H-NMR of **4c** in $CDCl_3$.

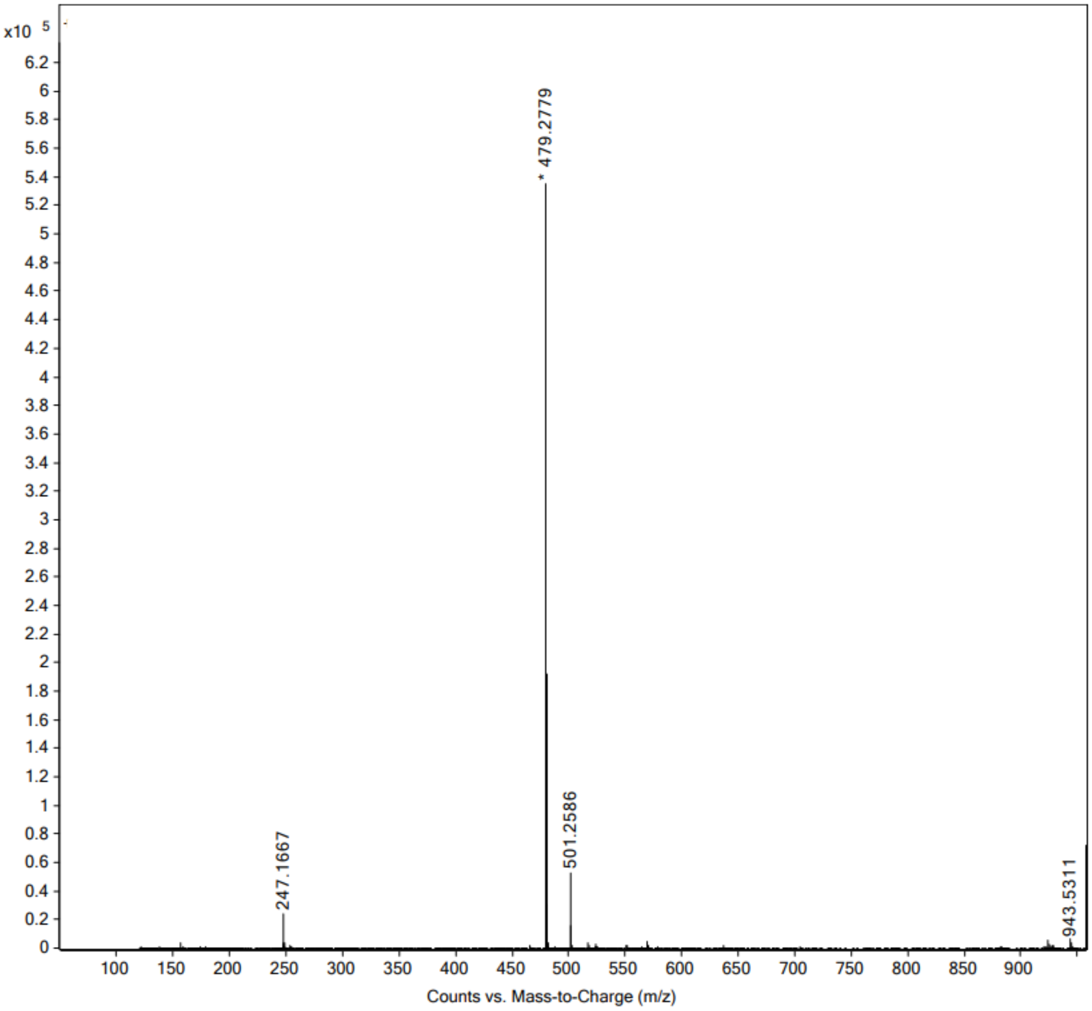


High resolution mass spectrum (ESI) of **4c**.

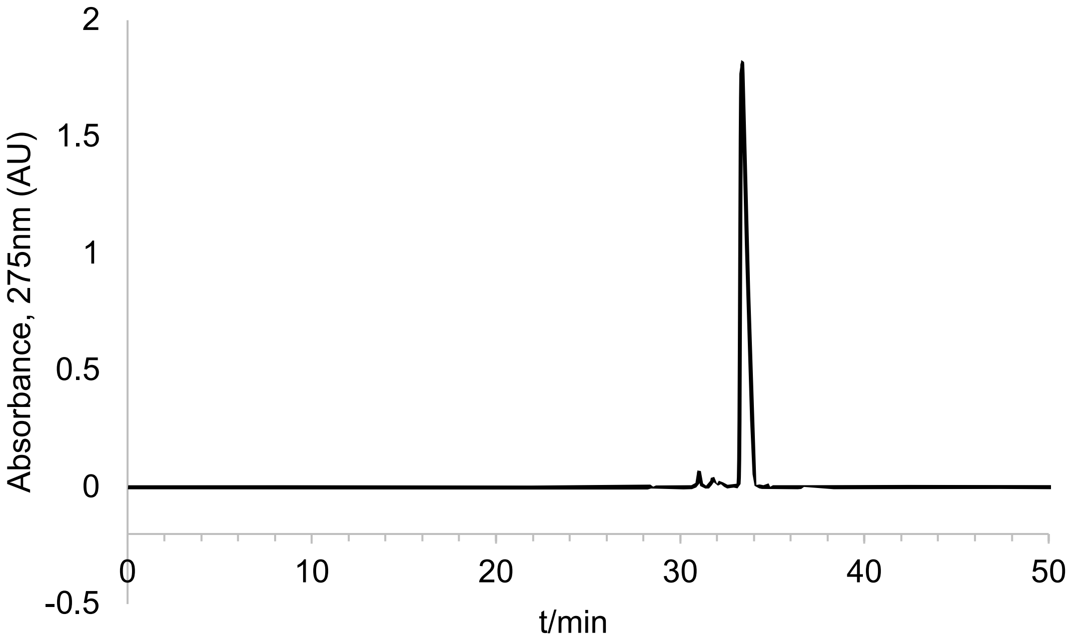


Analytical HPLC chromatogram of **4c.**

**4d.**

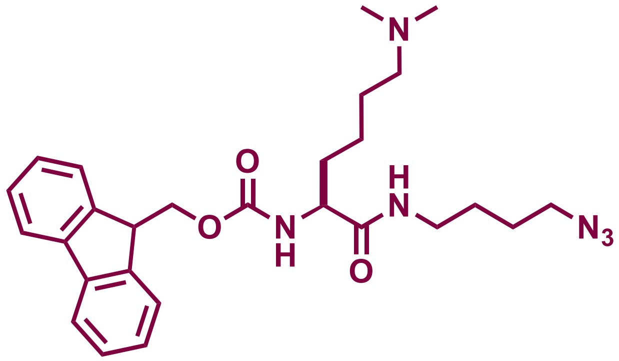


ESI-MS [M+H]+ m/z calcd. 493.2922 for $C_{27}H_{37}N_6O_3$, obsd. 493.2926. $^1$H NMR (600 MHz, cdcl$_3$) δ 1.45 (m, 1H), 1.59 (m, 2H), 1.70 (m, 1H), 1.82 (m, 2H), 2.82 (brs, 6H), 3.10 (brs, 1H), 4.20 (t, J = 6Hz, 1H), 4.21 (m, 1H), 4.32 (m, 1H), 6.00 (brs, 1H), 7.30 (t, J= 6 Hz, 2H), 7.40 (t, J= 6 Hz, 2H), 7.59 (t, J= 6 Hz, 2H), 7.76 (d, J= 6 Hz, 2H), 10.33 (brs, 1H), 11.13 (brs, 1H). $^{13}$C NMR (151 MHz, cdcl$_3$) δ 21.91, 23.52, 26.20, 26.55, 32.00, 39.11, 42.72, 43.47, 47.15, 51.05, 53.96, 57.71, 67.23, 114.89, 116.81, 120.10, 125.24, 127.18, 127.87, 141.37, 143.90, 156.58, 160.76, 161.01, 172.37.

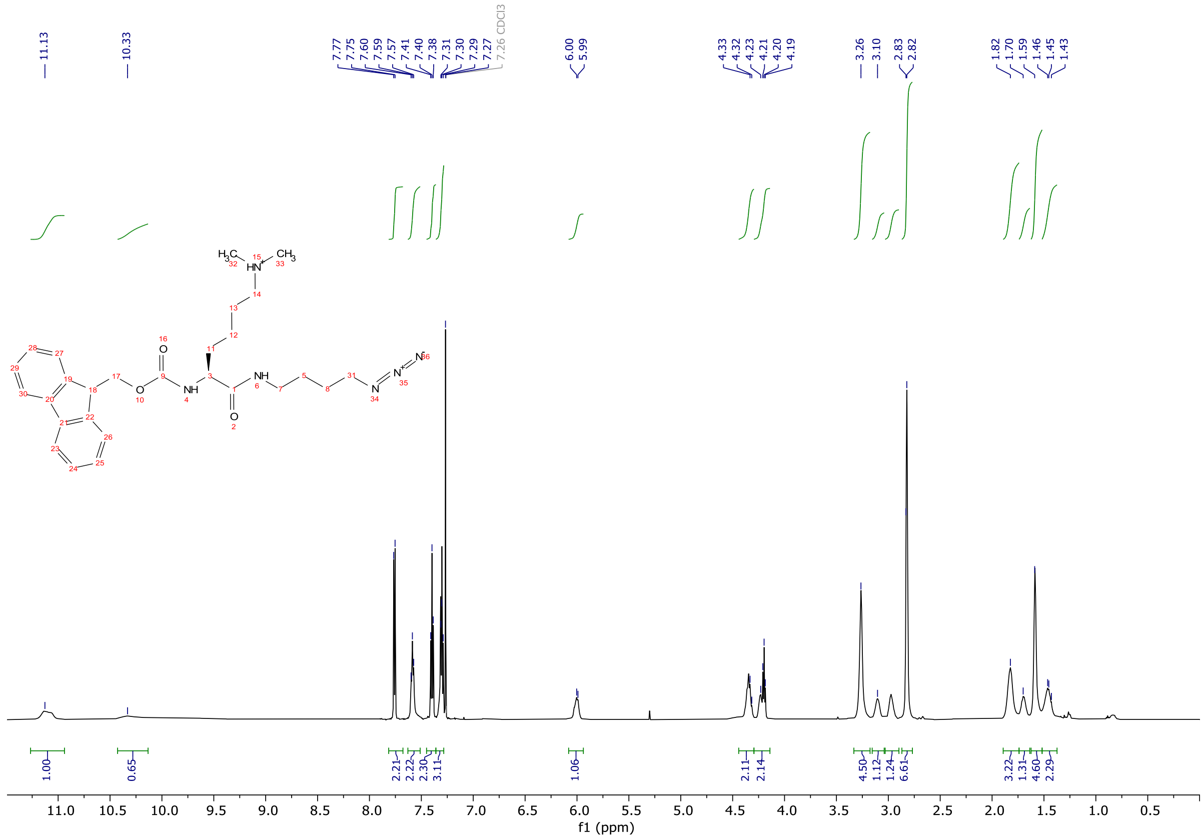

$^{1}$H-NMR of **4d** in $CDCl_3$.

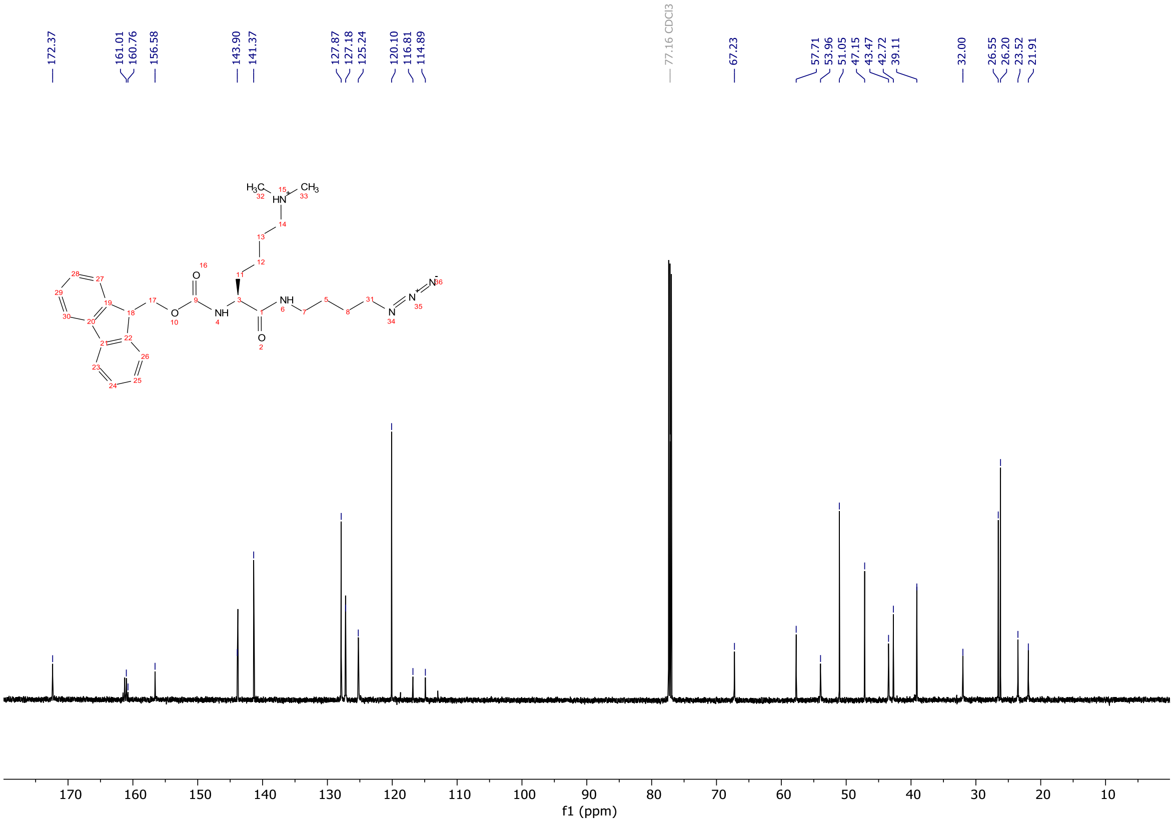

$^{13}C$-NMR of **4d** in $CDCl_3$.

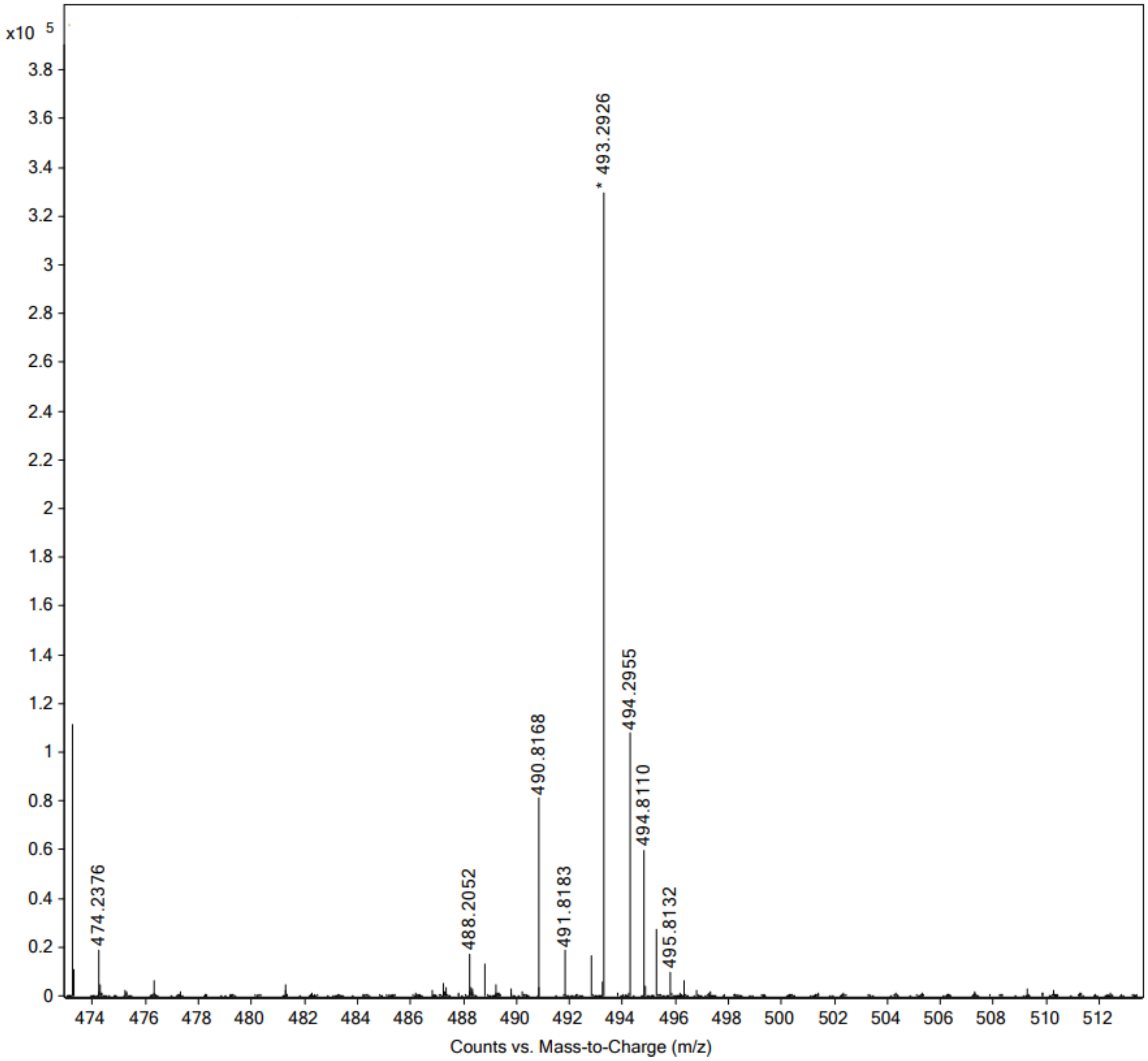


High resolution mass spectrum (ESI) of **4d**.

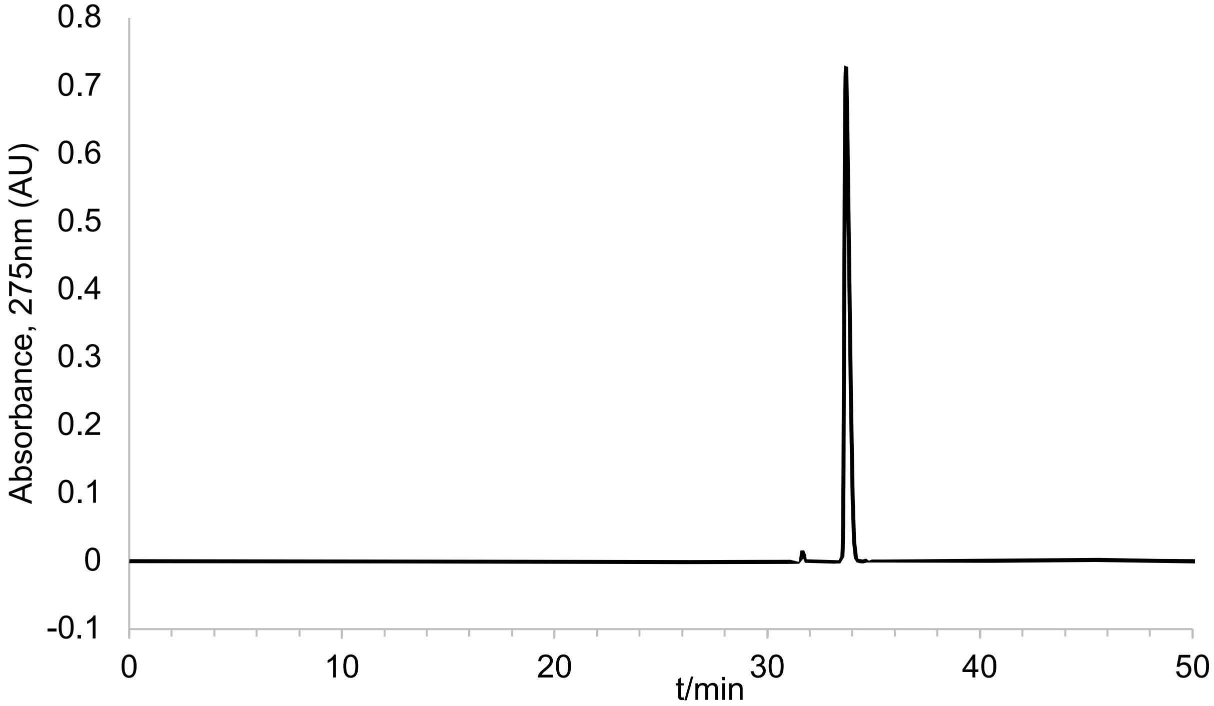


Analytical HPLC chromatogram of **4d**.

**4e.**

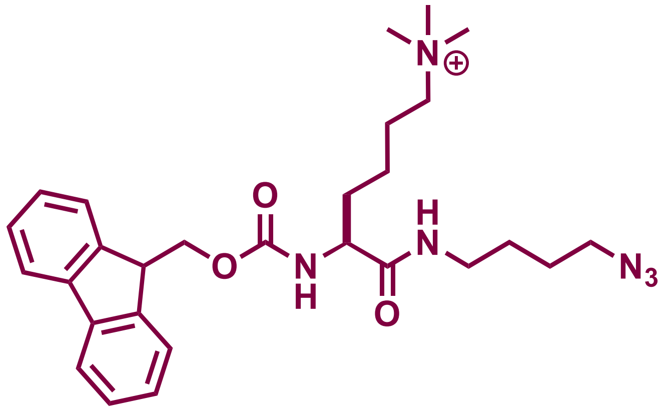


ESI-MS [M+H]+ m/z calcd. 507.3078 for $C_{28}H_{39}N_6O_3$, obsd. 507.3081. $^1$H NMR (600 MHz, $CDCl_3$) δ 1.52 (m, 1H), 1.64 (m, 2H), 1.84 (m, 1H), 2.02 (m, 1H), 3.27 (t, J= 6Hz, 2H), 3.29, 3.35 (brs, 9H), 3.42 (m, 1H), 3.79 (m, 1H), 4.19 (t, J= 6 Hz, 1H), 4.29 (m, 1H), 4.42 (m,1H), 5.92 (brs, 1H), , 7.30 (dt, J= 1, 6Hz, 2H), 7.38 (dt, J= 1, 6Hz, 2H), 7.60 (d, J= 4, 6Hz, 2H), 7.74 (d, J= 6Hz, 2H).

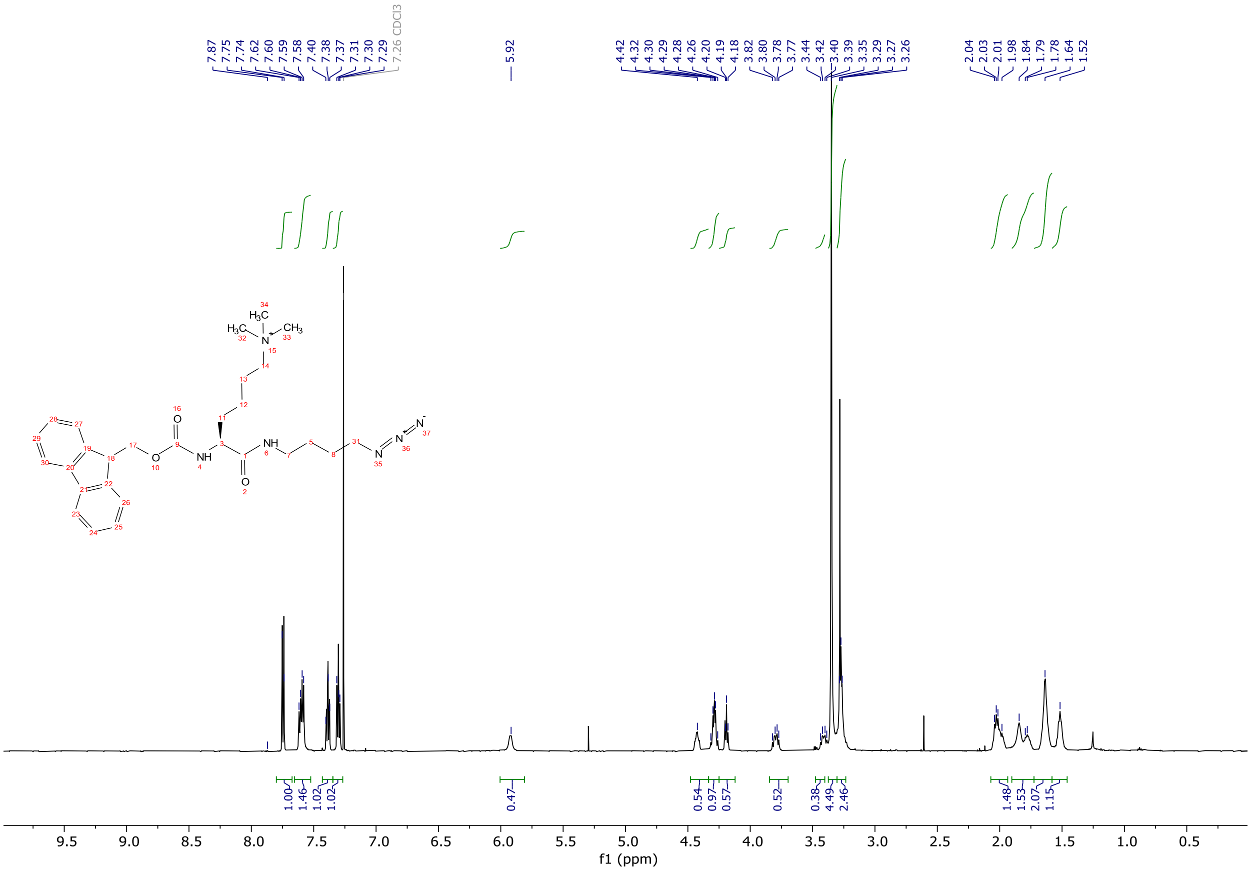

[1]H-NMR of **4e** in $CDCl_3$.

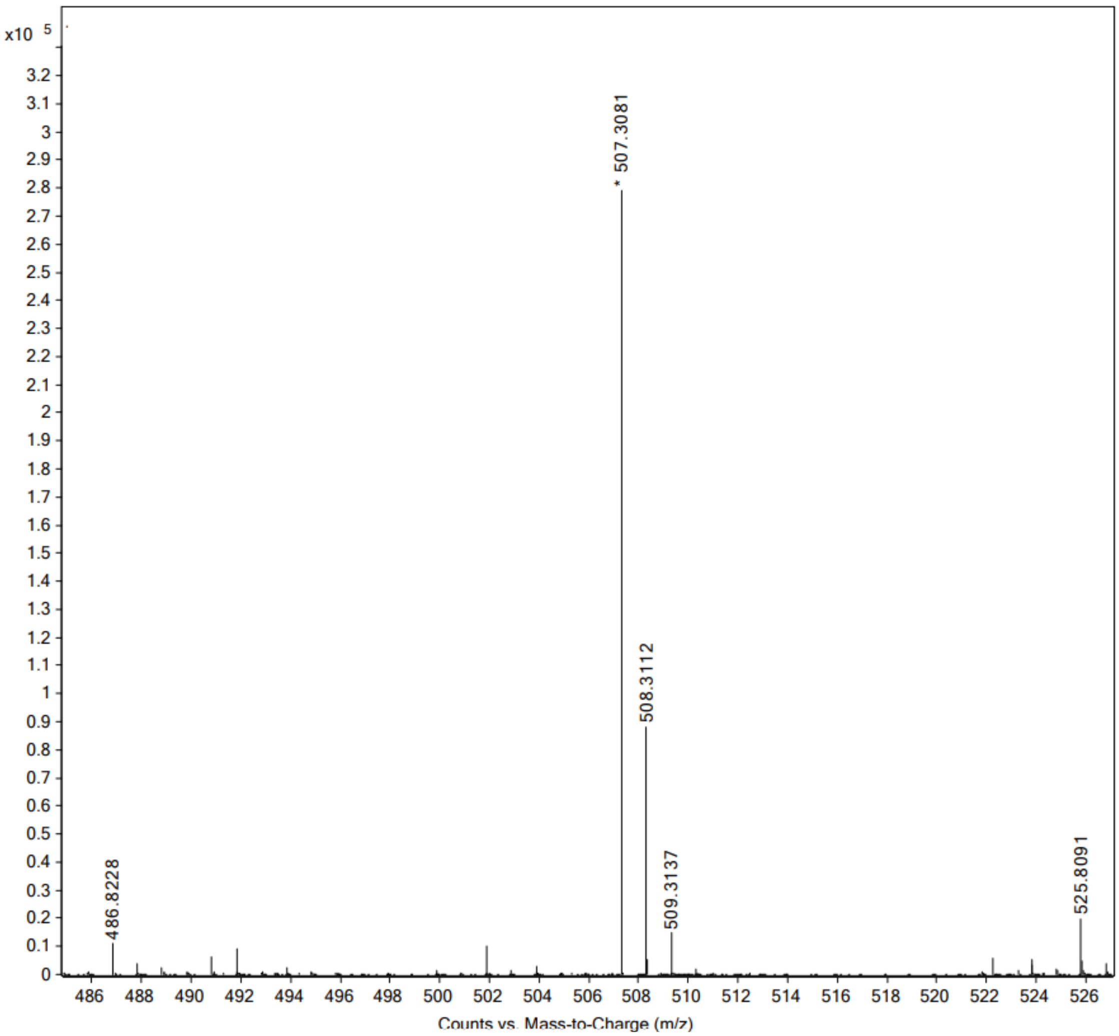


High resolution mass spectrum (ESI) of **4e**.

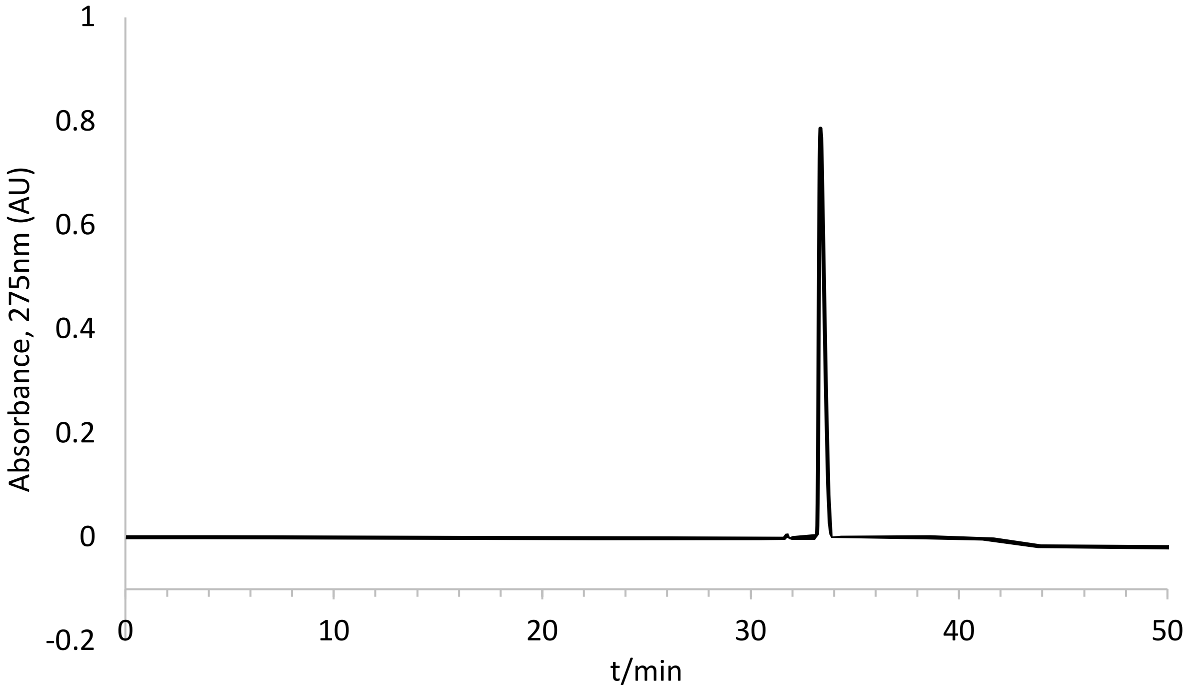


Analytical HPLC chromatogram of **4e**.

**Synthesis of 5a.**

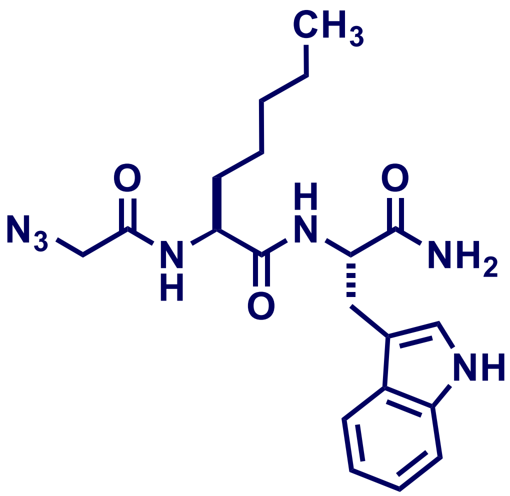

100 mg of Fmoc-aminomethyl)-phenoxy resin (Rink amide resin) of loading capacity 0.48 mmol/g was swollen in DCM after which the Fmoc protecting group was removed using 20% piperidine in DMF for 30 min. The resin was washed using alternating DCM and MeOH washes and finally rinsed with DMF. Fmoc-Trp (Boc)-OH was coupled to the resin using 4 equivalents of the amino acid and 4 equivalents each of Oxyma and DIC; the coupling reaction was done at room temperature with shaking for 2 h. The Fmoc group was similarly removed using 20% piperidine and following wash steps as described above, Fmoc-Nle-OH was similarly coupled to Trp(Boc) to make Fmoc-Lys(Boc)-Trp(Boc)-Rink amide resin. The Fmoc group was deprotected similarly and coupling of 2-azidoacetic acid was done using 4eq of 2-azidoacetic and 4eq of each of Oxyma and DIC. The peptide was cleaved off the resin using 95% TFA: 2.5% TIPS: 2.5% $H_2O$ and purified by preparative RP-HPLC on a on a Phenomenex Luna 10 µm C8(2) 100 Å (250 x 21.2 mm) using a 30 to 100% linear gradient of MeCN in $H_2O$ each containing 0.1% TFA at 10 mL/min. ESI-MS $[M+H]^+$ m/z calcd. 400.2092 for $C_{19}H_{26}N_7O_3$, obsd. 400.2095. $^1$H NMR (600 MHz, $CD_3CN$) δ $^1$H NMR (600 MHz, $cd_3cn$) δ 0.83 (t, J= 6Hz, 3H), 1.12-127 (m, 5H) 1.49 (m, 1H), 1.61 (m, 1H), 3.08 (dd, J= 2 and 6 Hz, 1H), 3.26 (dd, J= 2 and 6 Hz, 1H), 3.77 (ABd, J= 6Hz, 2H), 4.13 (q, J= 6 Hz,1H), 4.56 (m, 1H), 5.80 (brs, 1H), 6.42 (brs, 1H), 6.90 (brt, J= 6Hz, 1H), 7.05 (t, J= 6Hz, 1H), 7.10 (d, J=1 Hz, 1H), 7.13 (t, J= 6Hz, 1H), 7.38 (d, J=6 Hz, 1H), 7.60 (d, J=6 Hz, 1H), 9.18 (s, 1H). $^{13}$C NMR (151 MHz,

cd$_3$cn) δ 1.04, 1.11, 1.18, 1.32, 1.49, 14.14, 22.91, 28.02, 28.05, 28.20, 31.87, 31.91, 52.36, 52.39, 54.54, 54.62, 54.91, 54.94, 54.96, 55.01, 55.04, 63.85, 110.71, 112.43, 112.48, 118.83, 119.38, 119.92, 122.44, 124.77, 124.93, 128.35, 128.38, 137.26, 137.42, 169.90, 169.98, 173.10, 173.11, 173.18, 173.19, 175.68.

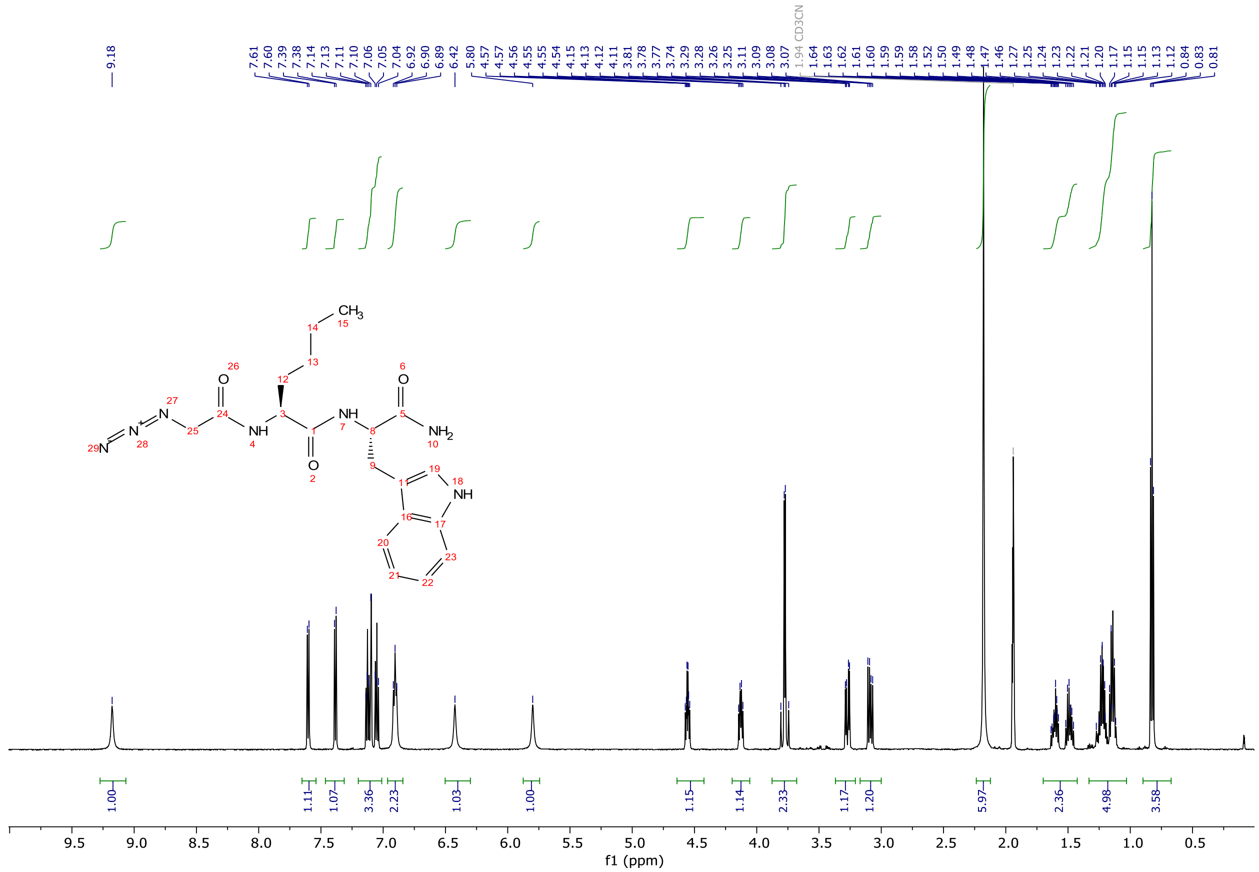

[1]H-NMR of **5a** in $CD_3CN$.

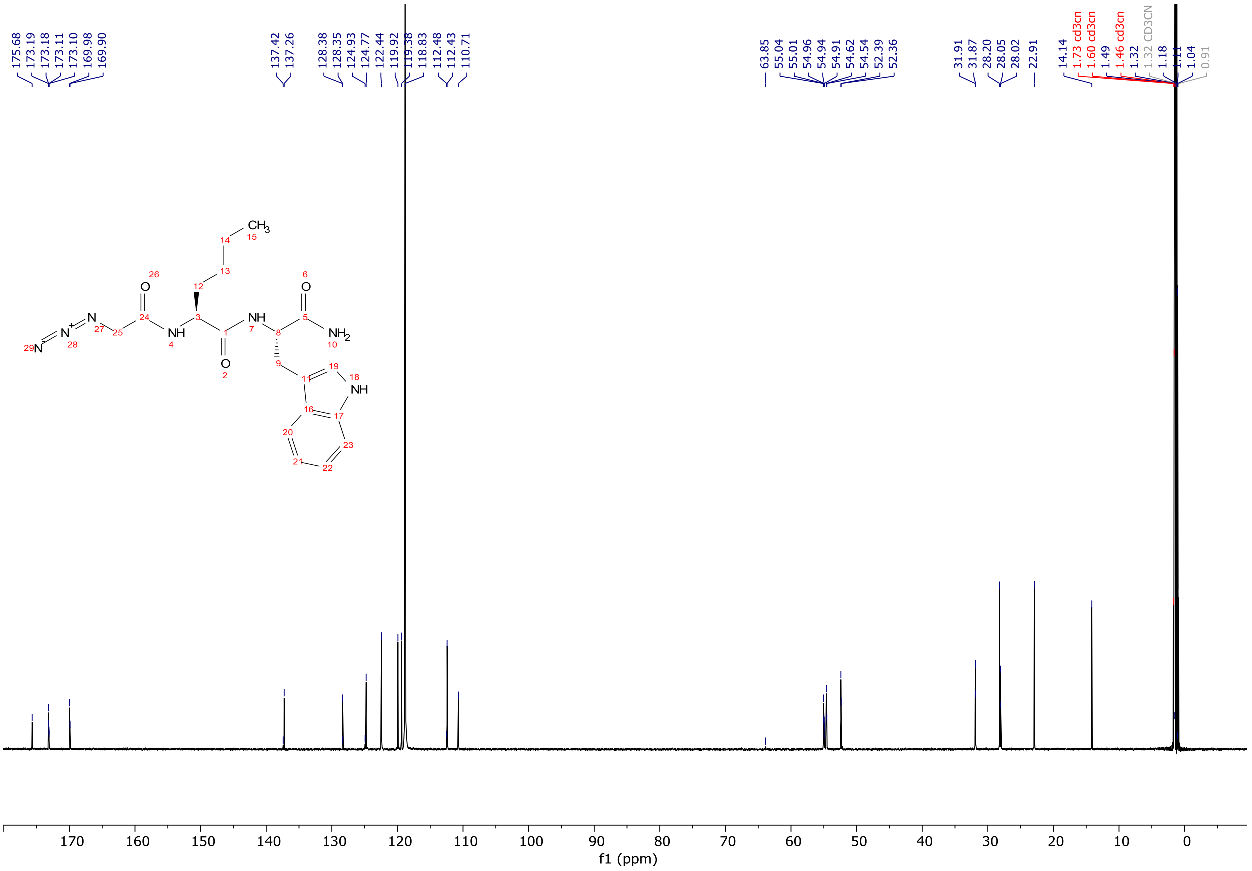


$^{13}$C-NMR of **5a** in $CD_3CN$.

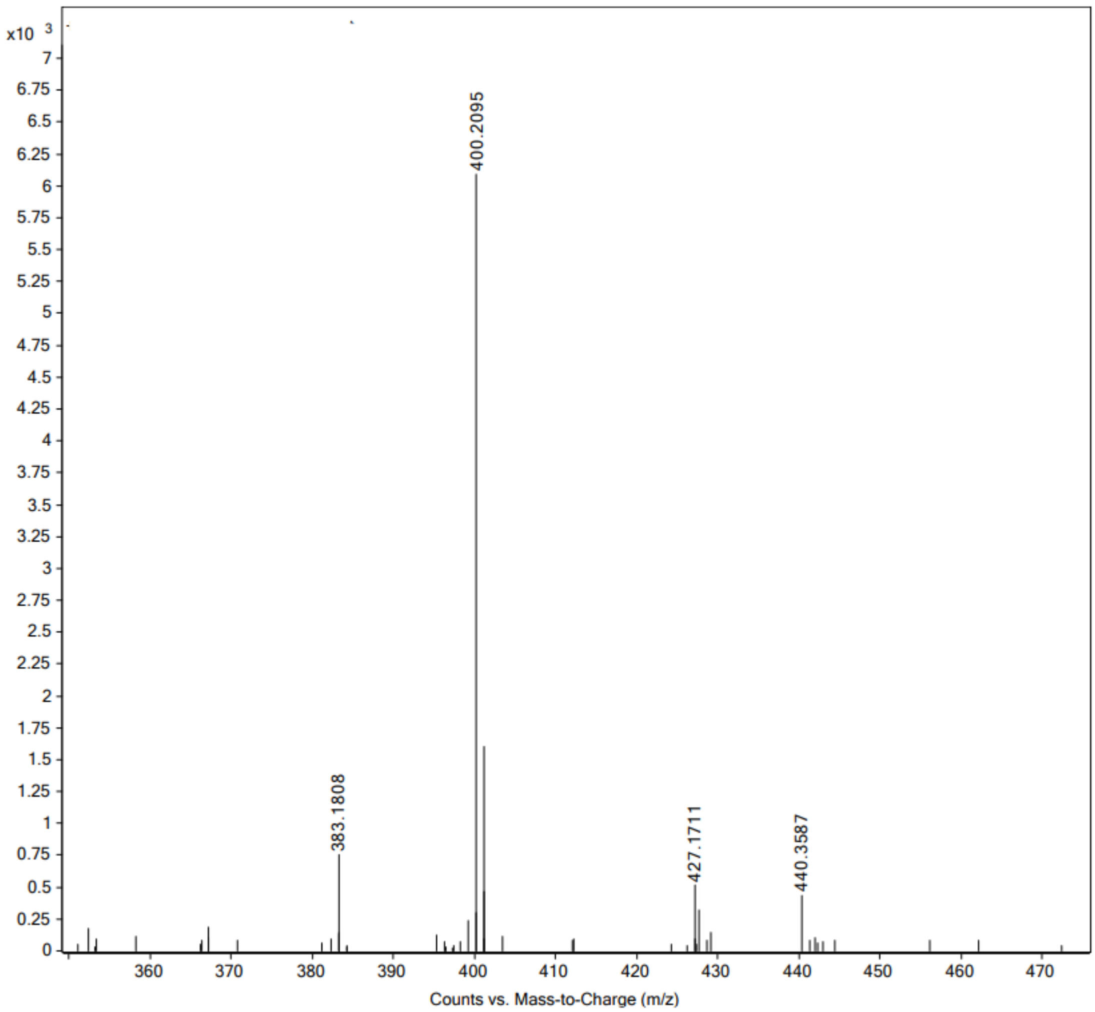


High resolution mass spectrum (ESI) of **5a**

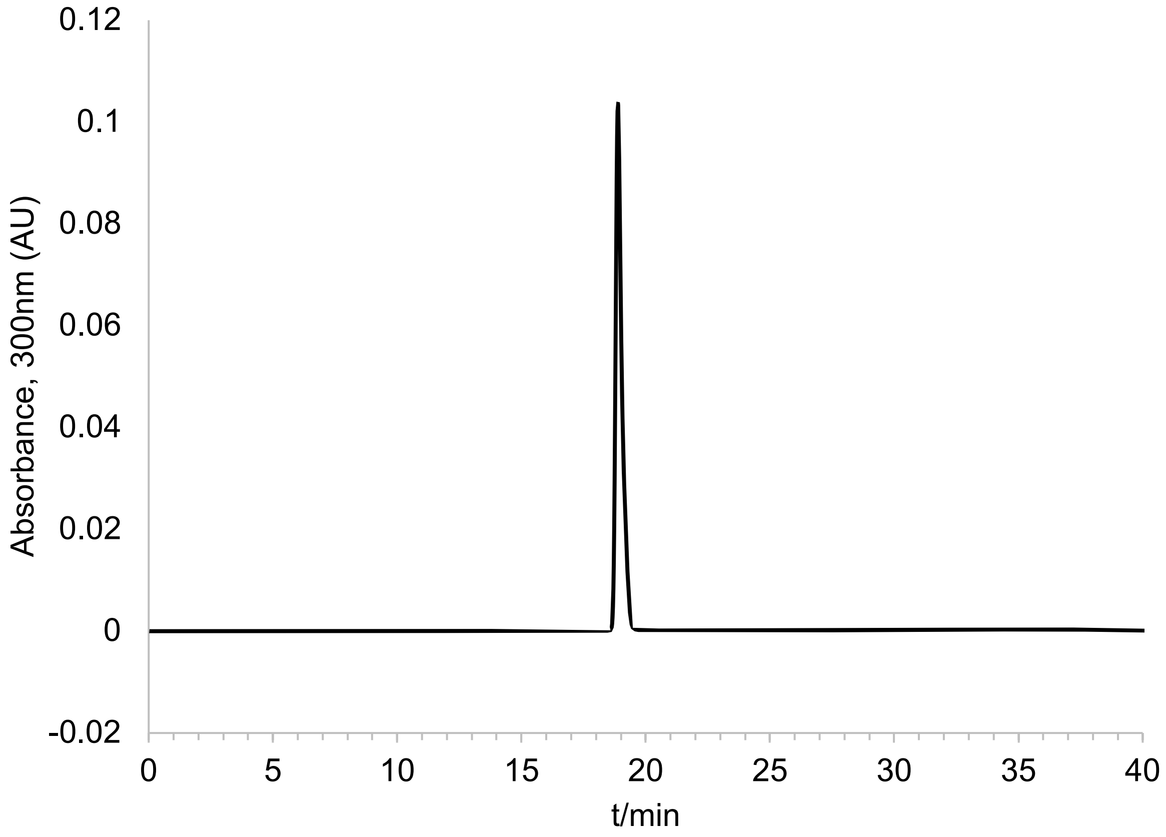


Analytical HPLC chromatogram of **5a**.

**Synthesis of 5b, 5c, 5d, 5e,** 2-azidoacetyl-Lys $(Me)_x$-Trp-$NH_2$ where x = 0, 1, 2, 3.

100 mg of Fmoc-(aminomethyl)-phenoxy resin (Rink amide resin) of loading capacity 0.48 mmol/g was swollen in DCM after which the Fmoc protecting group was removed using 20% piperidine in DMF for 30 min. The resin was washed using alternating DCM and MeOH washes and finally rinsed with DMF. The deprotected amine on Rink amide resin was coupled to Fmoc-Trp (Boc)-OH using 4 equivalents of the amino acid and 4 equivalents of each of Oxyma and DIC; the coupling reaction was done at room temperature with shaking for 2 h. The Fmoc group was similarly removed using 20% piperidine and following wash steps as described above, either Fmoc-Lys(Boc)-OH or Fmoc-Lys(Boc,Me)-OH or Fmoc-Lys$(Me)_2$-OH.HCl or Fmoc-Lys$(Me)_3$-$OH^+Cl^-$ was coupled to the deprotected amine using 4 equivalents of each of the amino acid, 4 eq. of Oxyma and 4 eq. DIC in DMF; the coupling reaction was done at room temperature with shaking for 2 h. The Fmoc group was deprotected similarly and coupling of 2-azidoacetic acid was done using 4 eq of 2-azidoacetic and 4 eq of each of Oxyma and DIC. The peptide was cleaved off the resin using 95% TFA: 2.5% TIPS: 2.5% $H_2O$ and purified by preparative RP-HPLC on a Phenomenex Luna 10 µm C8(2) 100 Å (250 x 21.2 mm) using a 30 to 100% linear gradient of MeCN in $H_2O$ each containing 0.1% TFA at 10 mL/min.

**5b.**

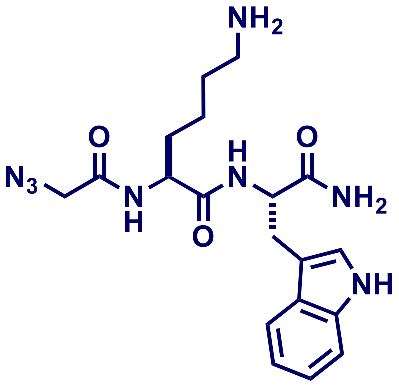


**5b** ESI-MS $[M+H]^+$ m/z calcd. 415.2201 for $C_{19}H_{27}N_8O_3$, obsd. 415.2200. $^1$H NMR (600 MHz, $CD_3CN$) δ 1.14-1.23 (m, 1H), 1.49-1.60 (m, 4H), 2.81 (t, J= 6Hz, 1H), , 3.08 (dd, J= 2 and 6 Hz, 1H), 3.27 (dd, J= 2 and 6Hz, 1H), , 3.81 (ABd, J= 6Hz, 2H),, 4.21 (q, J= 6Hz, 1H), 4.59 (m, 1H), 6.22 (brs, 1H), 6.77 (brs, 1H), 7.04 (t, J= 6Hz, 1H),, 7.11 (t, J= 6Hz, 1H), 7.38 (d, J=6 Hz, 1H), 7.59 (d, J=6 Hz, 1H), 9.51 (s, 1H).

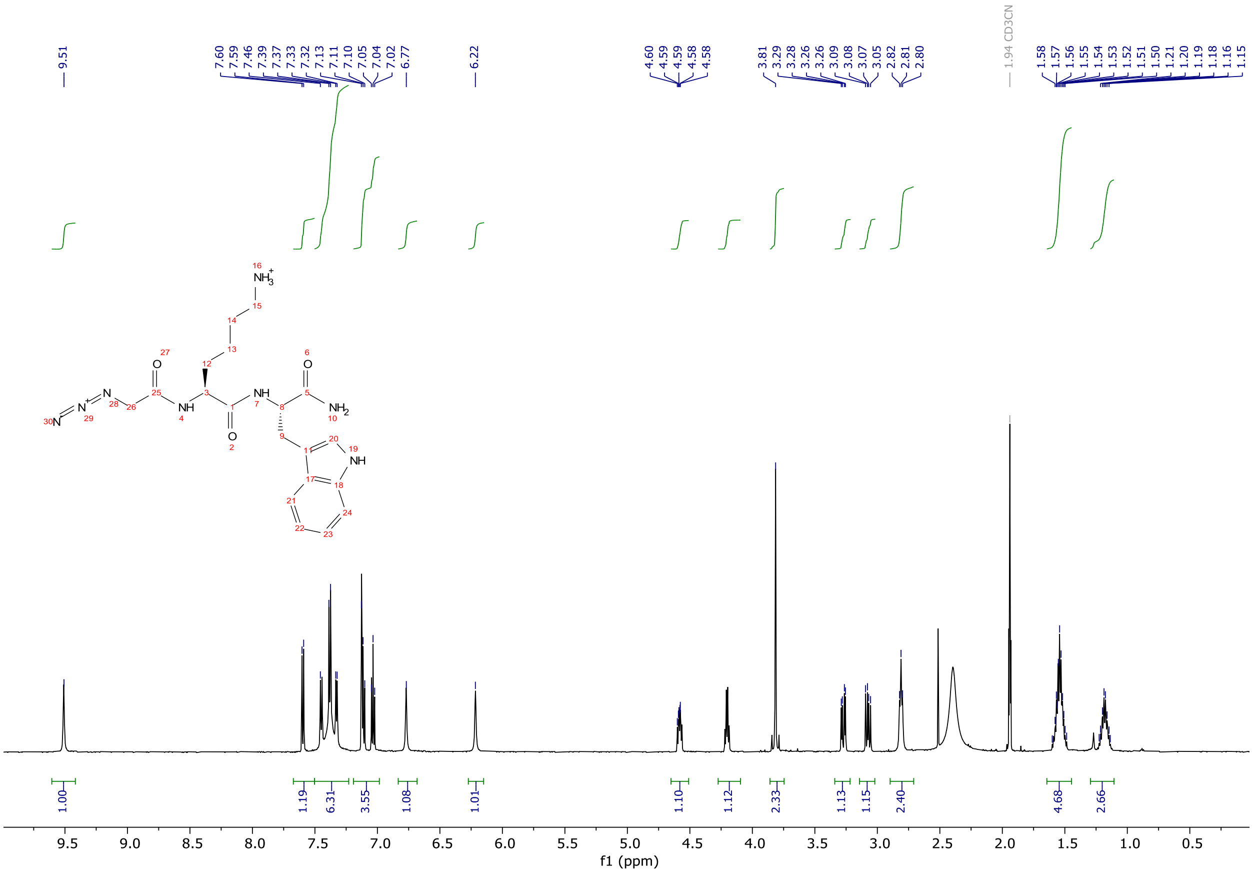

[1]H-NMR of **5b** in $CD_3CN$.

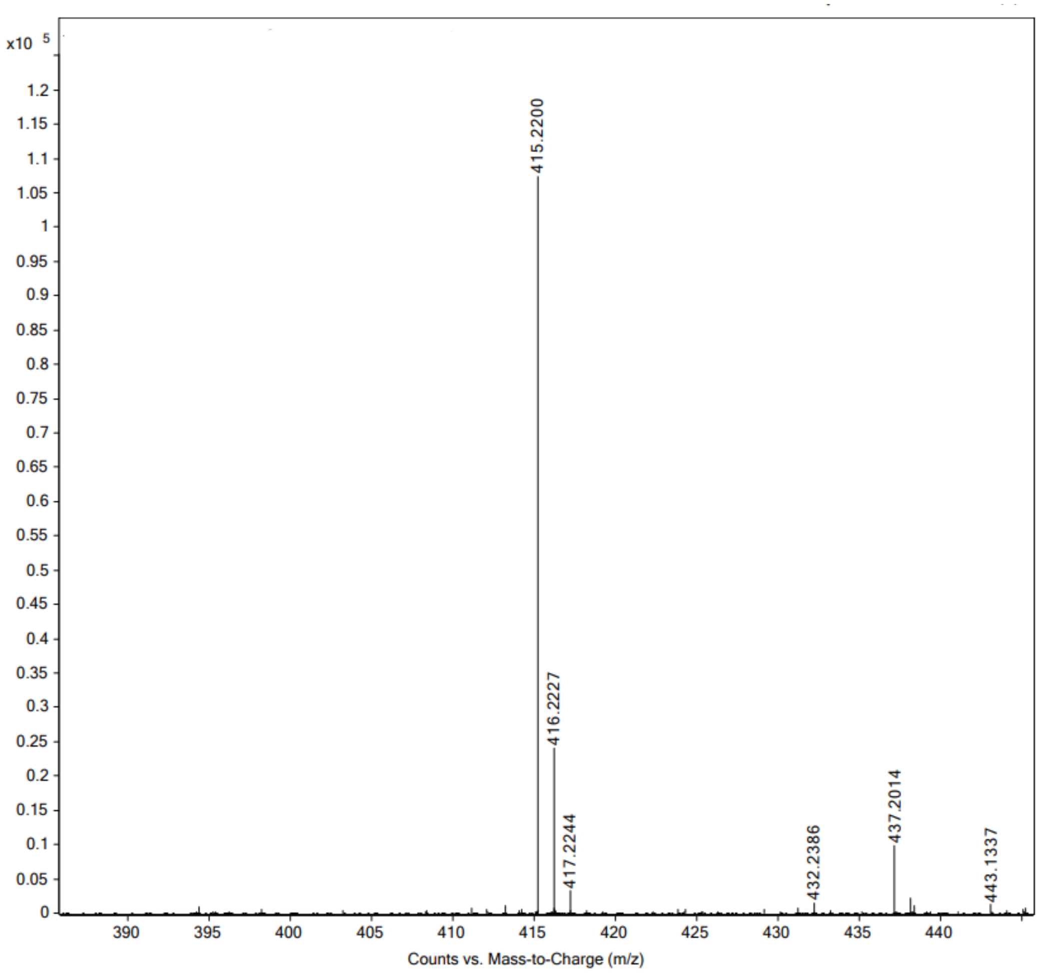


High resolution mass spectrum (ESI) of **5b**.

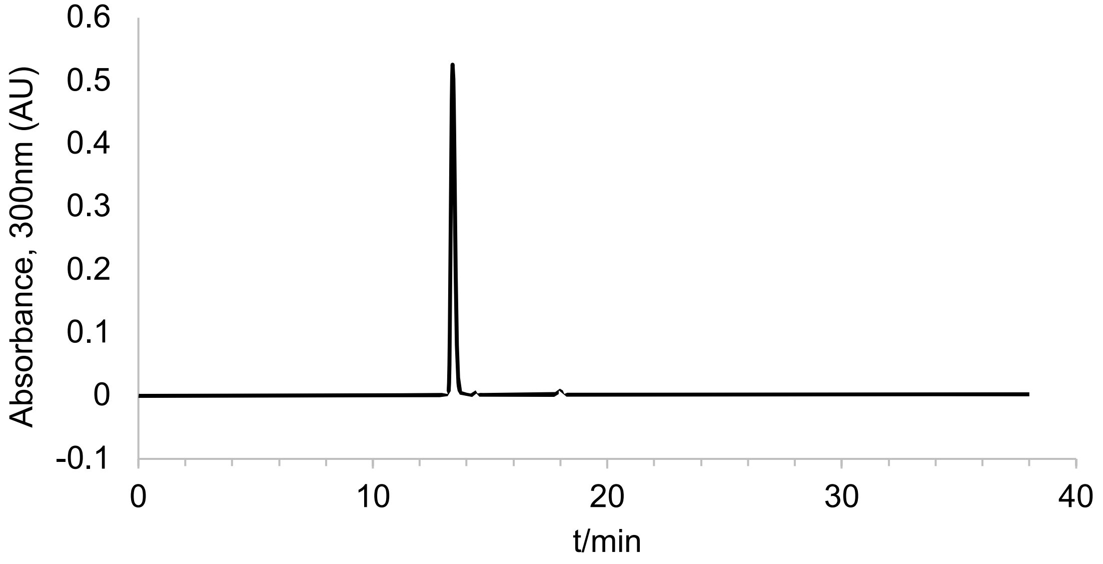


Analytical HPLC chromatogram of **5b**.

**5c.**

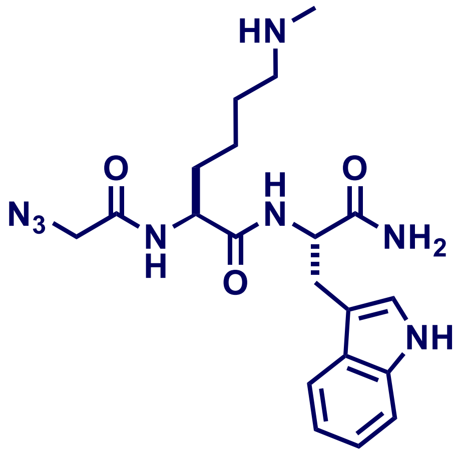


**5c** ESI-MS $[M+H]^+$ m/z calcd. 429.2357 for $C_{20}H_{29}N_8O_3$, obsd. 429.2360. $^1H$ NMR (600 MHz, $CD_3CN$) δ 1.14-1.23 (m, 1H), 1.49-1.60 (m, 4H), 2.81 (t, J= 6Hz, 1H), , 3.08 (dd, J= 2 and 6 Hz, 1H), 3.27 (dd, J= 2 and 6Hz, 1H), 3.81 (ABd, J= 6Hz, 2H),, 4.21 (q, J= 6Hz, 1H), 4.59 (m, 1H), 6.22 (brs, 1H), 6.67 (brs, 1H), 7.05 (t, J= 6Hz, 1H), 7.13 (t, J= 6Hz, 1H), 7.39 (d, J=6 Hz, 1H), 7.60 (d, J=6 Hz, 1H), 7.91 (brs, 1H), 8.10 (brs, 1H), 9.44 (s, 1H). Another minor isomer is visible in NMR. $^{13}C$ NMR (151 MHz, $CD_3CN$) δ 1.04, 1.18, 1.32, 22.49, 25.81, 28.18, 31.72, 33.82, 49.75, 52.34, 54.40, 54.69, 111.20, 112.41, 118.32, 119.45, 119.93, 122.49, 124.77, 137.45, 169.20, 172.31, 183.49.

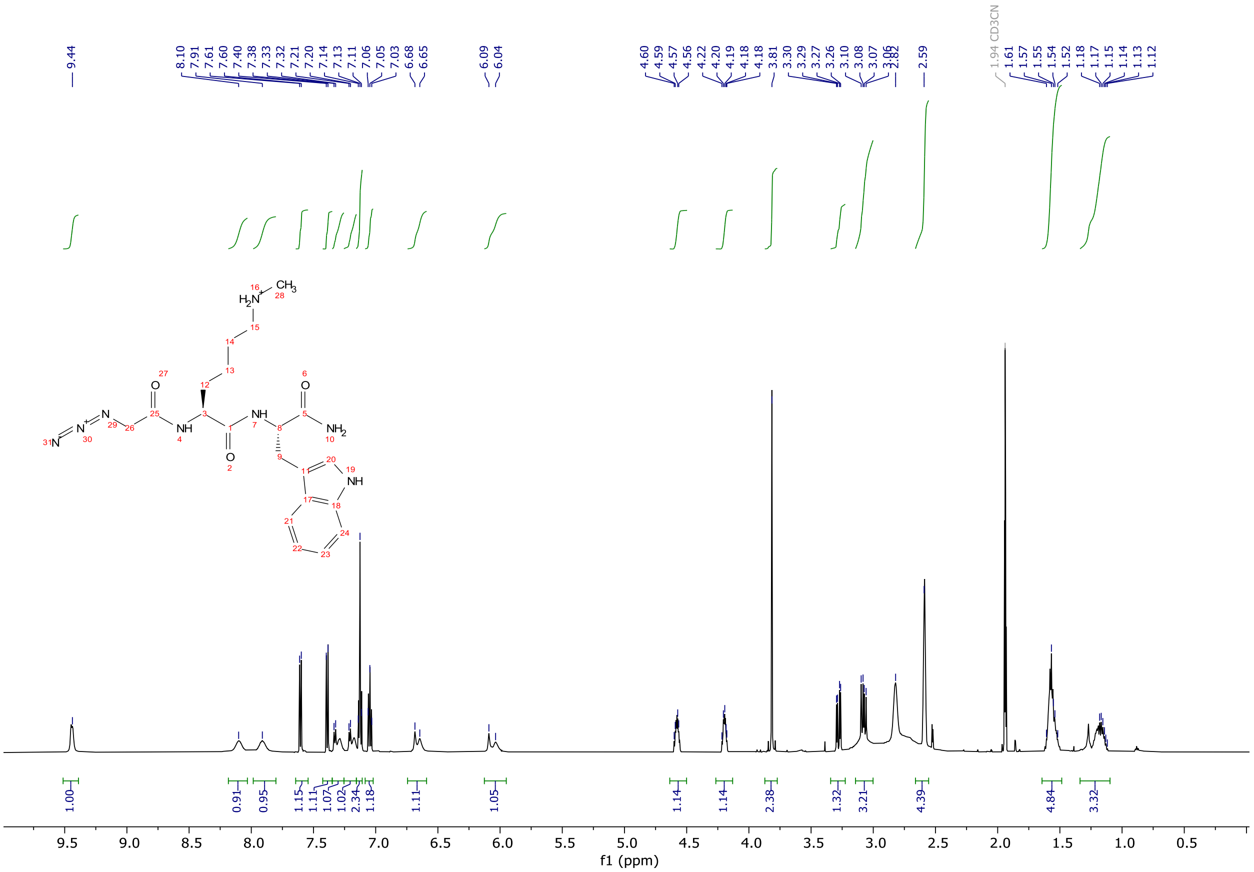

[1]H-NMR of **5c** in $CD_3CN$.

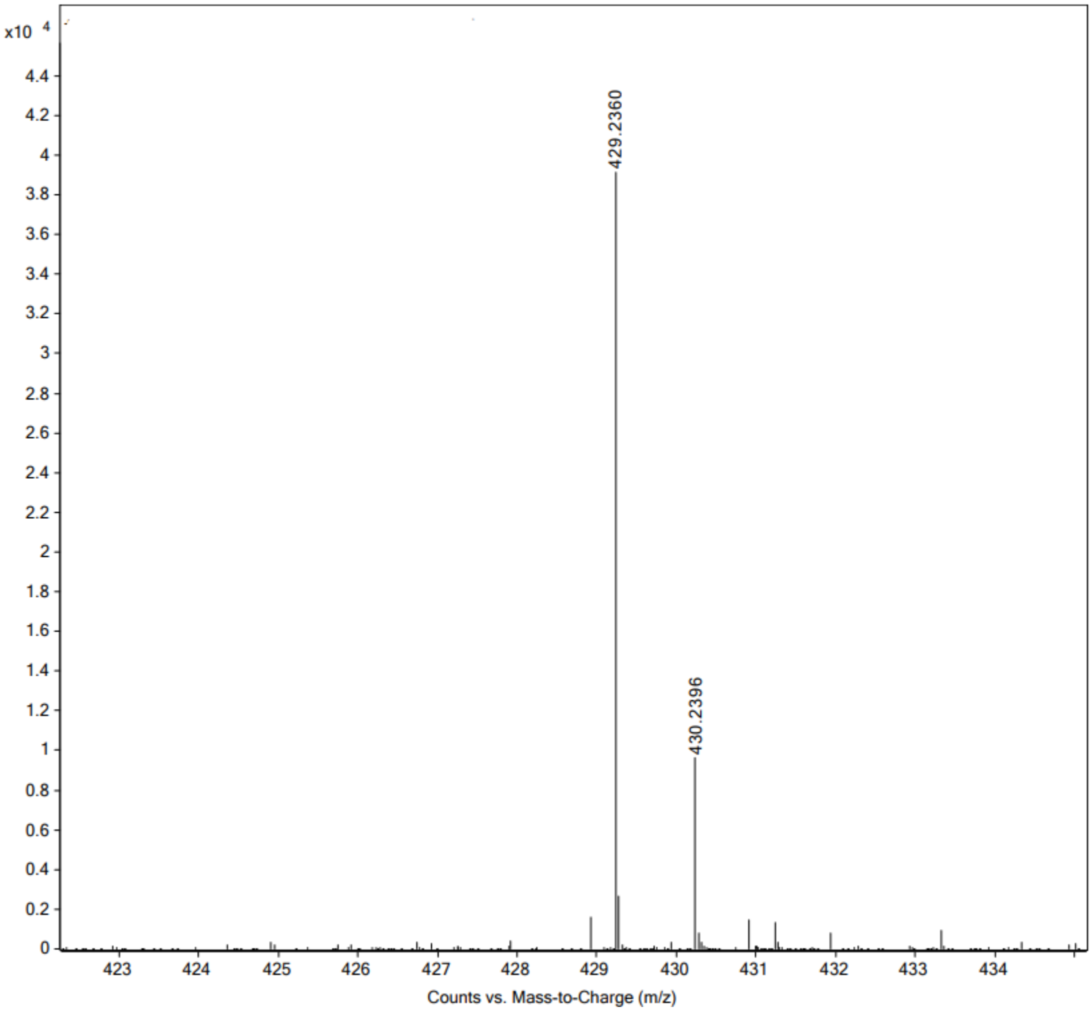


High resolution mass spectrum (ESI) of **5c**.

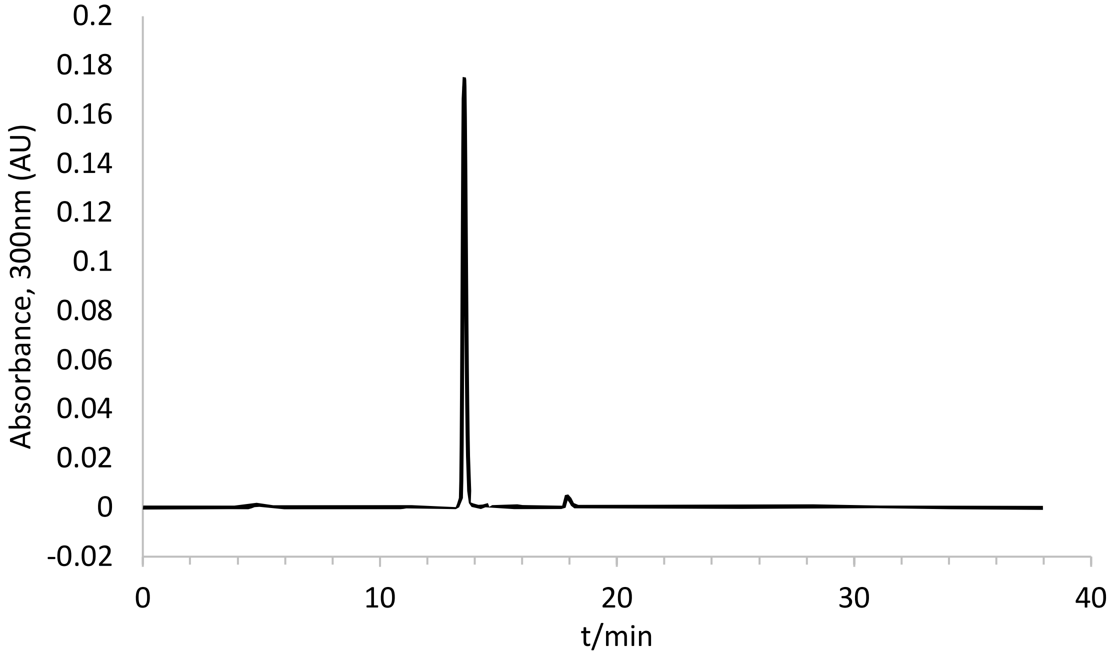


Analytical HPLC chromatogram of **5c**.

**5d.**

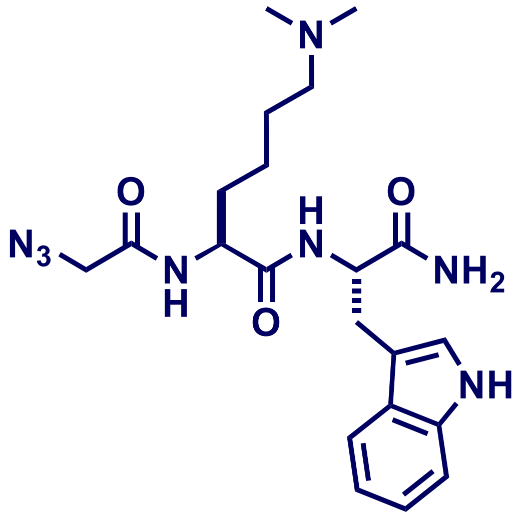


**5d** ESI-MS $[M+H]^+$ m/z calcd. 443.2514 for $C_{21}H_{31}N_8O_3$, obsd. 443.2516. $^1H$ NMR (600 MHz, $CD_3CN$) δ 1.08-1.18 (m, 2H), 1.50-1.63 (m, 4H), 2.72 (brs, 6H), 2.86 (q, J= 6Hz, 1H), 3.08 (dd, J= 2 and 6 Hz, 1H), 3.29 (dd, J= 2 and 6Hz, 1H), 3.81 (ABd, J= 6Hz, 2H), 4.18 (q, J= 6Hz, 1H), 4.60 (m, 1H), 6.00 (brs, 1H), 6.61 (brs, 1H), 7.05 (t, J= 6Hz, 1H), 7.10-7.14 (m, 4H), 7.39 (d, J=6 Hz, 1H), 7.60 (d, J=6 Hz, 1H), 9.39 (s, 2H). $^{13}C$ NMR (151 MHz, $CD_3CN$) δ 1.05, 1.60, 1.74, 22.60, 24.42, 28.20, 31.67, 43.46, 43.63, 52.37, 54.52, 54.61, 58.18, 111.25, 112.41, 118.34, 119.49, 119.96, 122.52, 124.74, 128.48, 137.46, 169.26, 172.12.

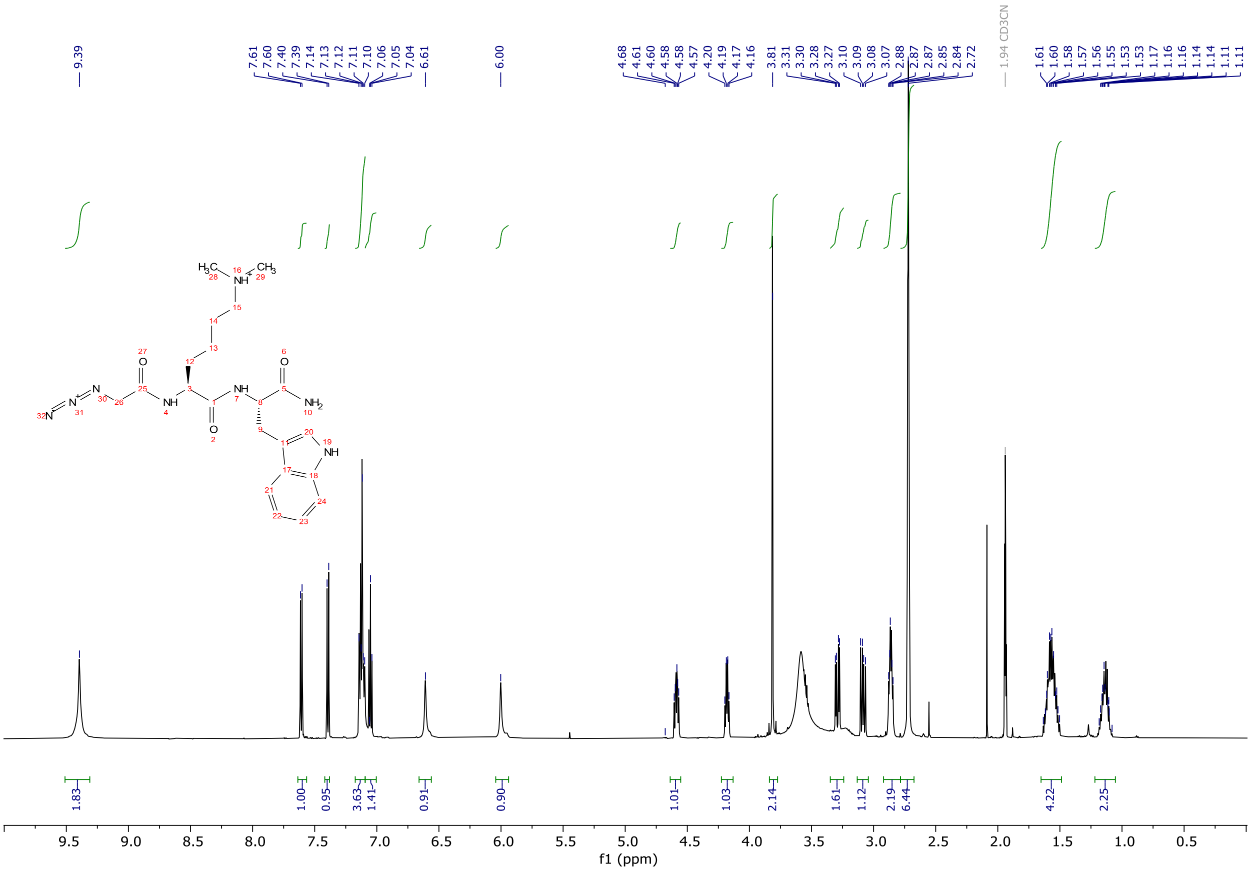

[1]H-NMR of **5d** in $CD_3CN$.

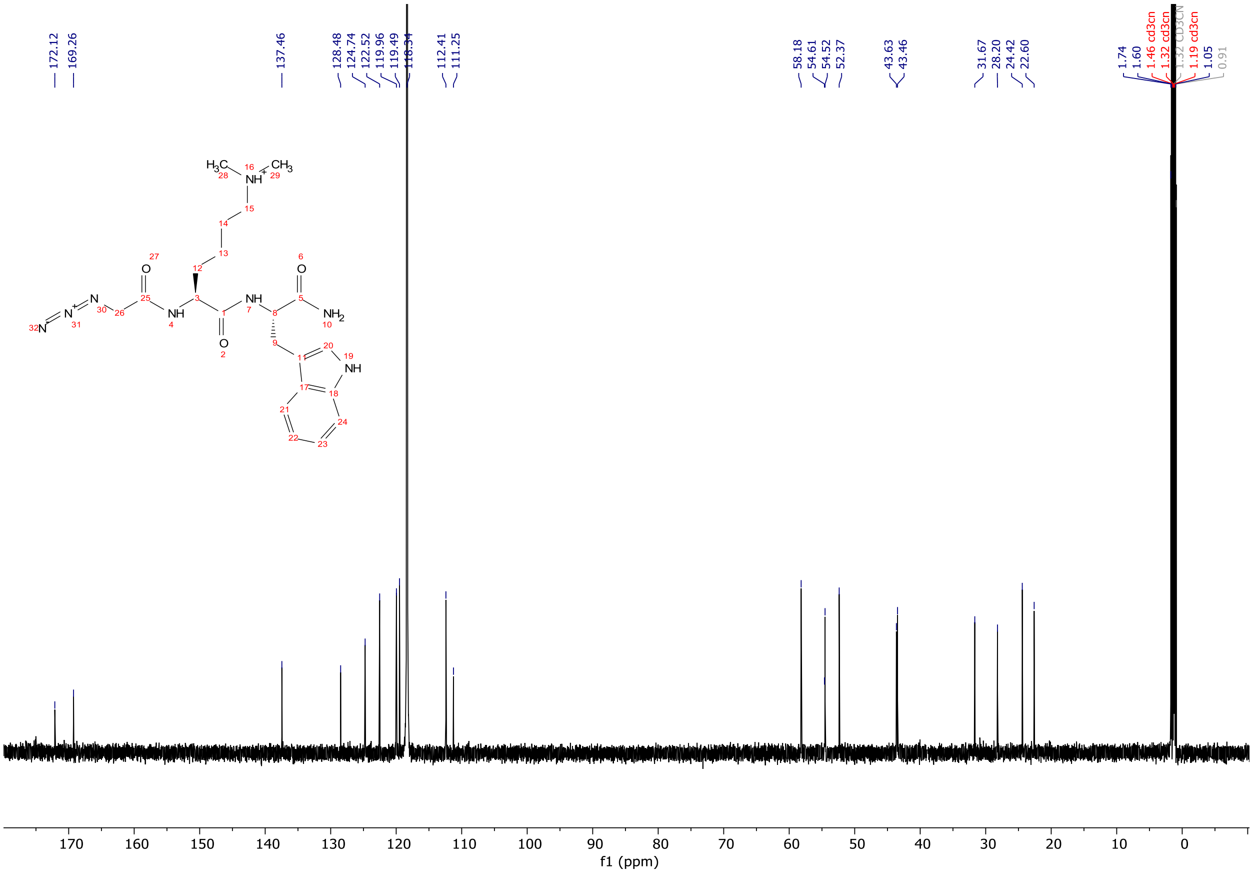

$^{13}$C-NMR of **5d** in $CD_3CN$.

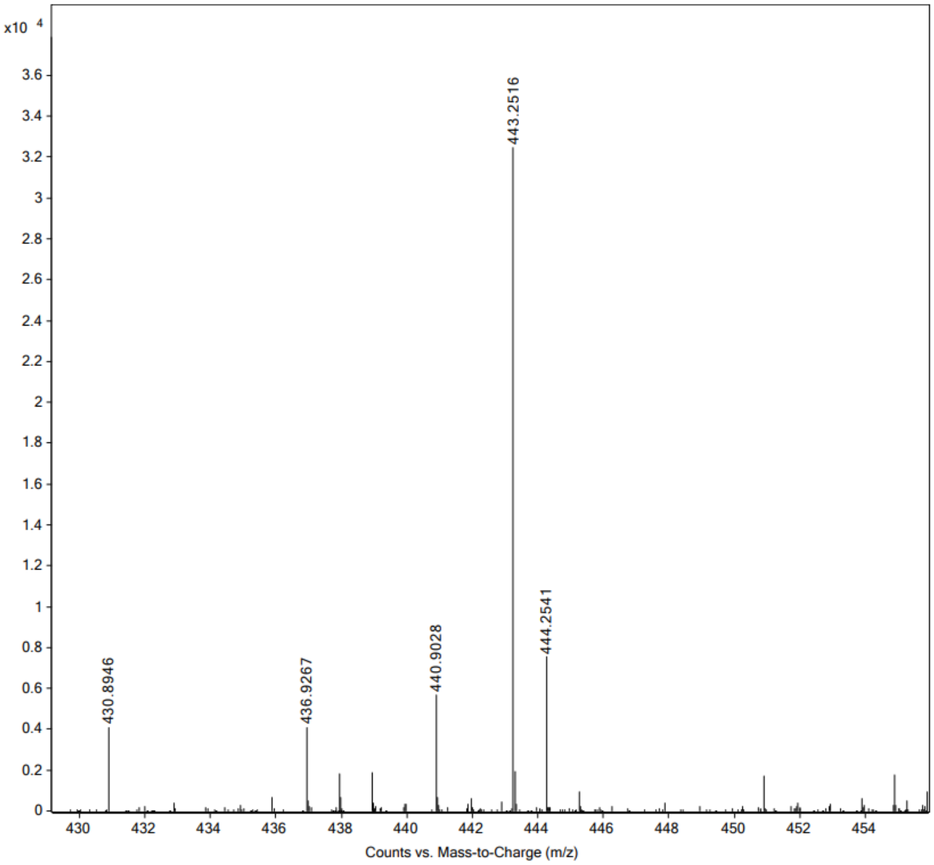


High resolution mass spectrum (ESI) of **5d**.

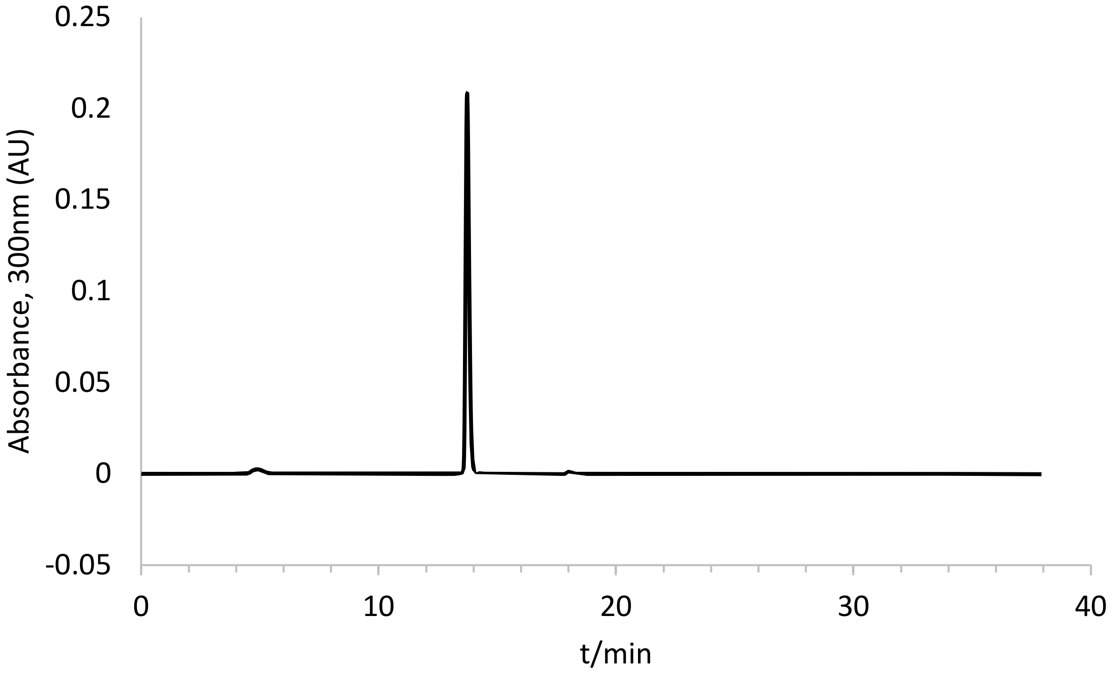


Analytical HPLC chromatogram of **5d**.

**5e.**

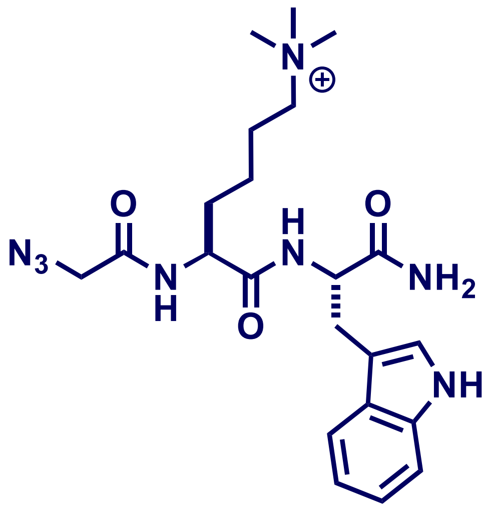


**5e** ESI-MS $[M+H]^+$ m/z calcd. 457.2679 for $C_{22}H_{33}N_8O_3$, obsd. 457.2680. $^1H$ NMR (600 MHz, $CD_3CN$) δ 1.10-1.18 (m, 2H), 1.53-1.68 (m, 4H), 2.95 (brs, 9H), 3.07 (m, 1H), 3.10 (dd, J= 2 and 6 Hz, 1H), 3.28 (dd, J= 2 and 6Hz, 1H), 3.82 (s, 2H), 4.21 (q, J= 6Hz, 1H), 4.57 (m, 1H), 5.87 (brs, 1H), 6.62 (brs, 1H), 7.05 (t, J= 6Hz, 1H), 7.11-7.14 (m, 2H), 7.27 (brs, 1H) 7.39 (d, J=6 Hz, 1H), 7.60 (d, J=6 Hz, 1H), 9.40 (s, 1H). $^{13}C$ NMR (151 MHz, $CD_3CN$) δ 22.66, 22.95, 28.26, 31.73, 52.35, 53.79, 53.82, 54.48, 54.73, 67.20, 111.38, 112.40, 118.33, 119.52, 119.93, 122.46, 124.83, 128.54, 137.44, 169.22, 174.72, 183.46.

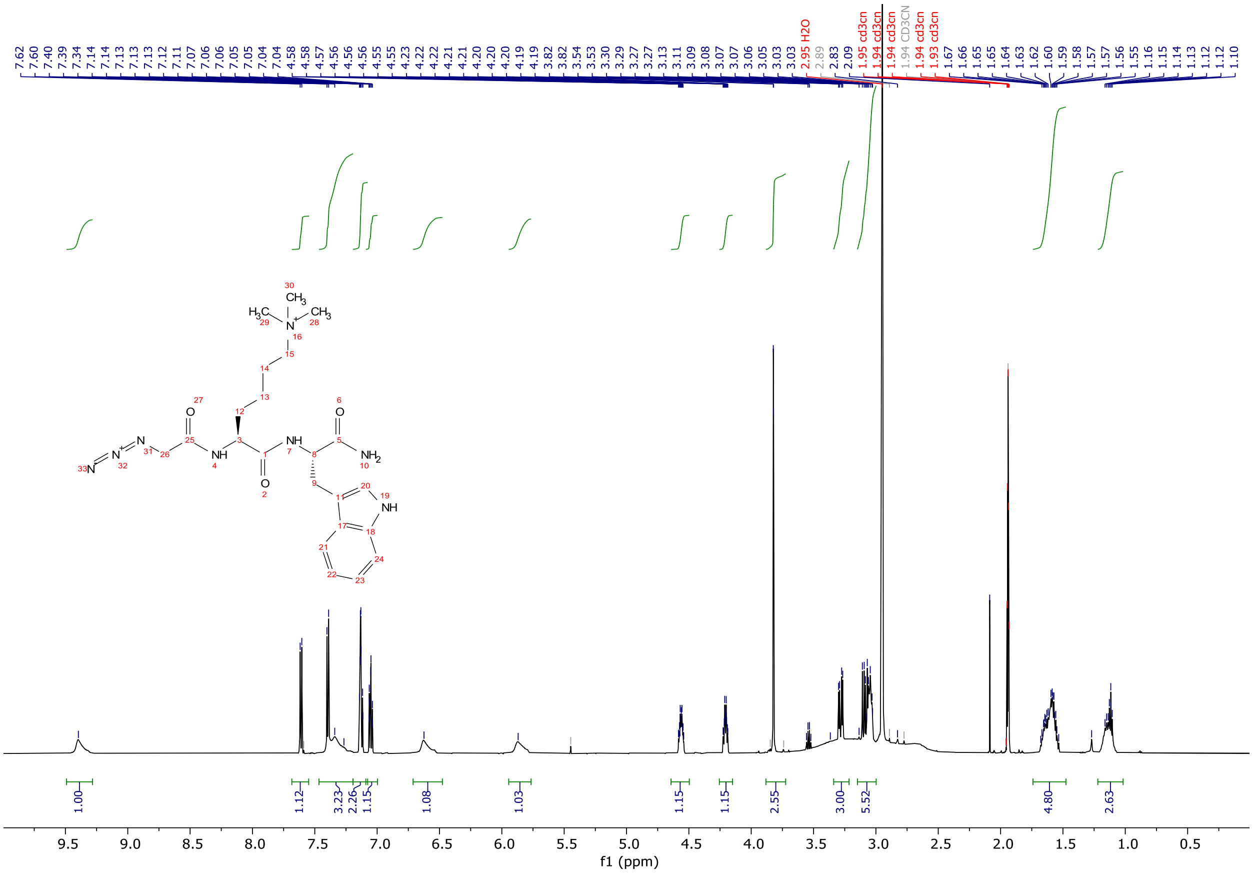

$^1$H-NMR of **5e** in $CD_3CN$.

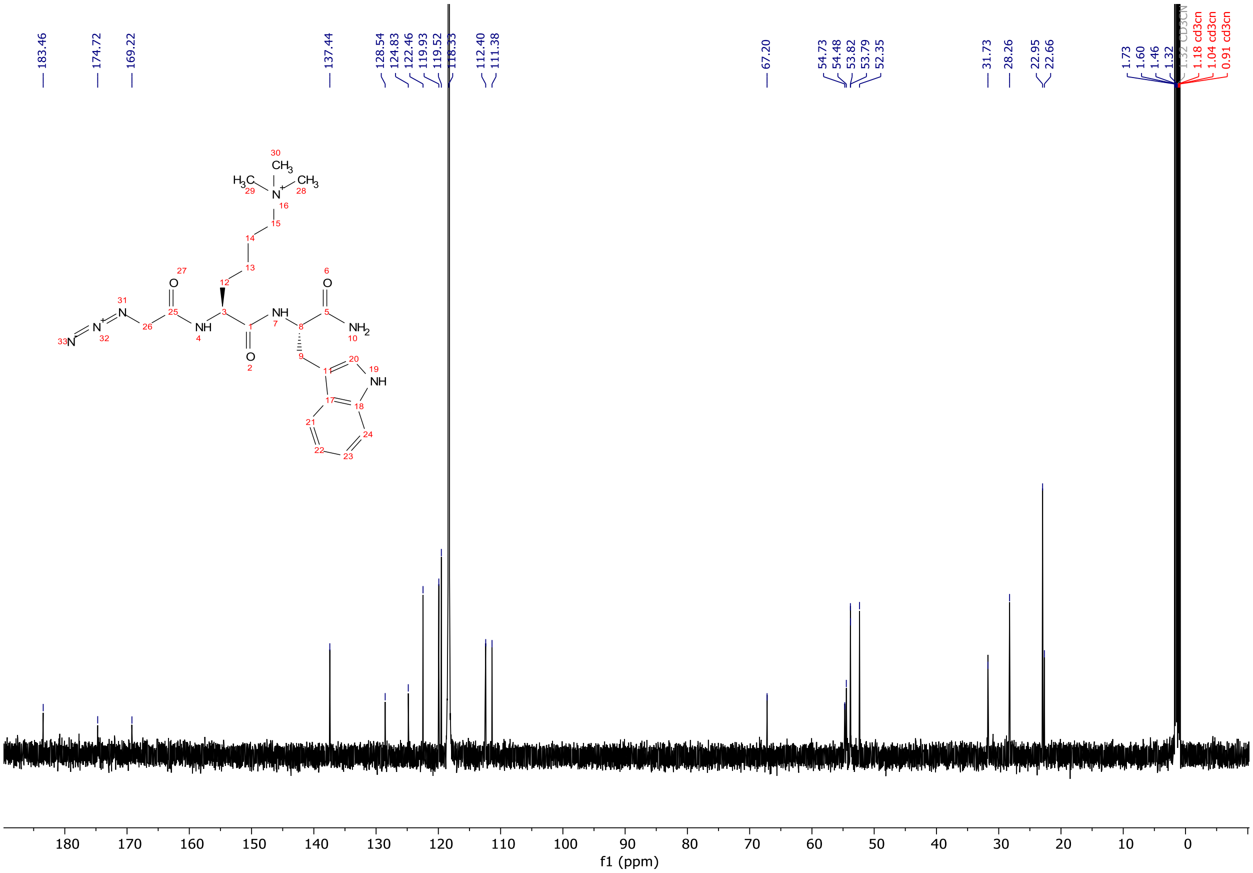


$^{13}$C-NMR of **5e** in $CD_3CN$.

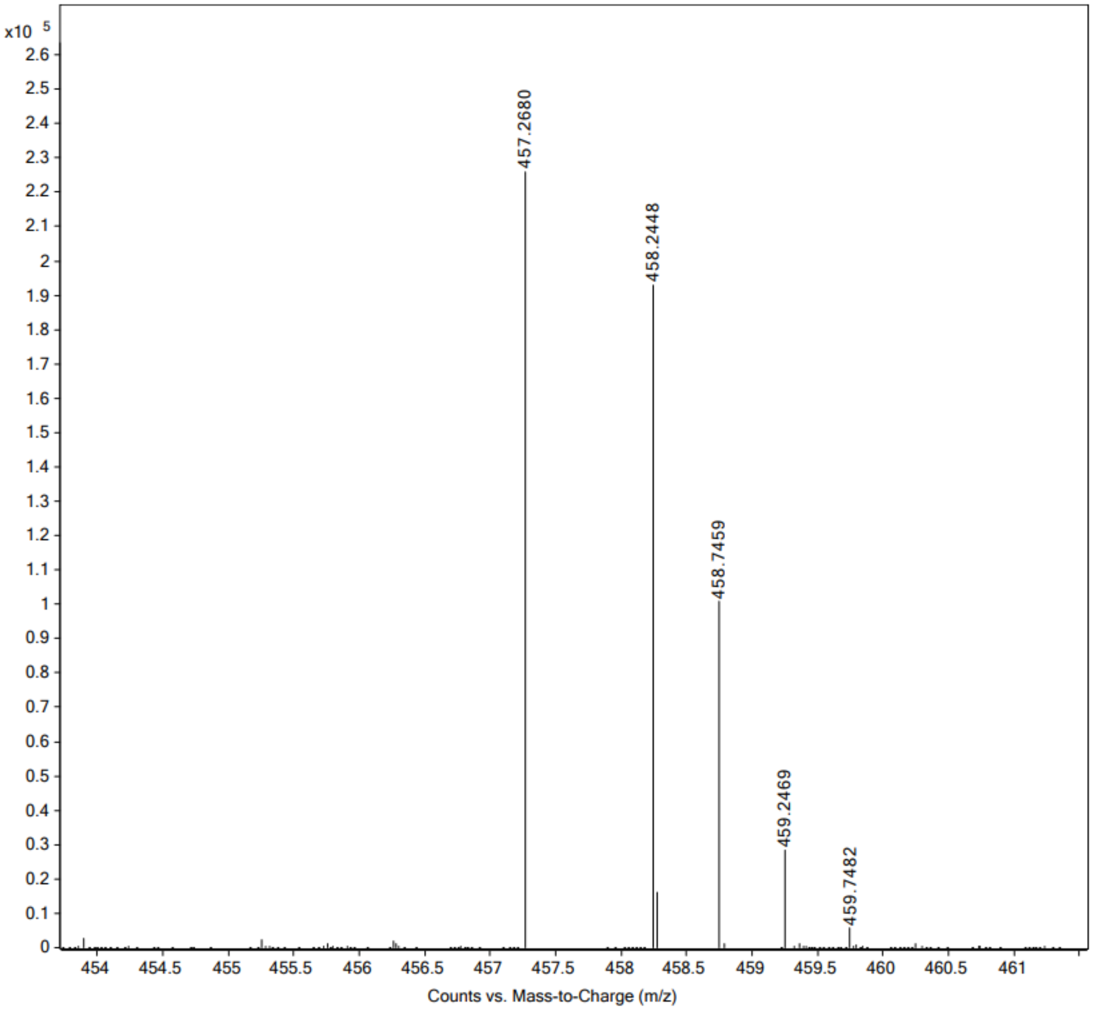


High resolution mass spectrum (ESI) of **5e**.

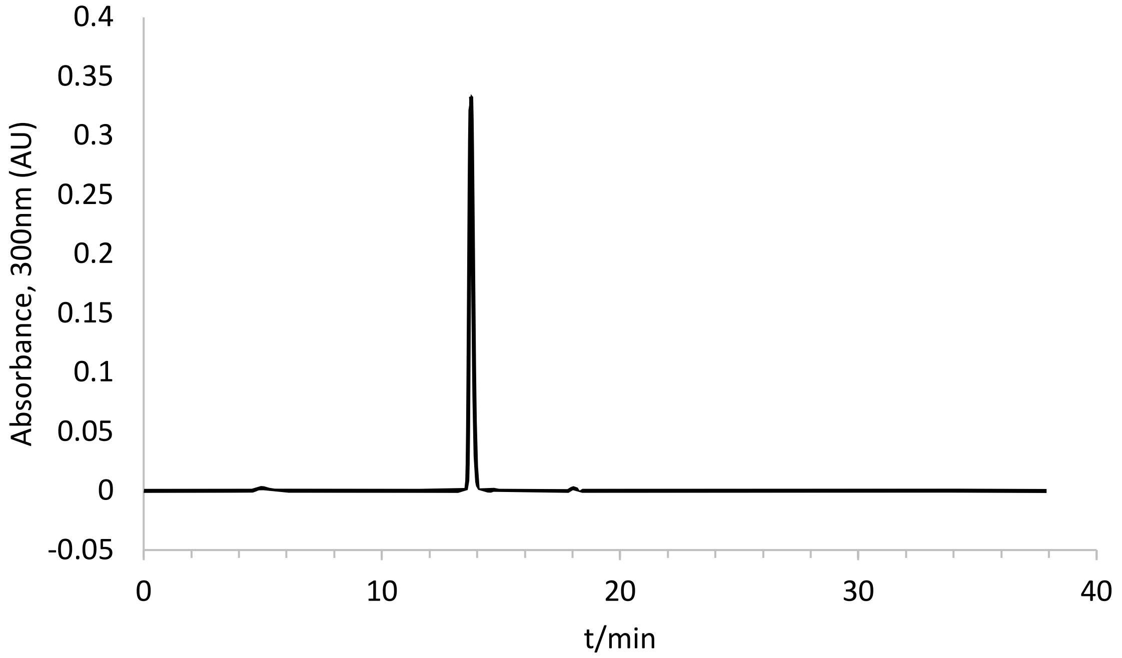


Analytical HPLC chromatogram of **5e**.

**Synthesis of 6a.**

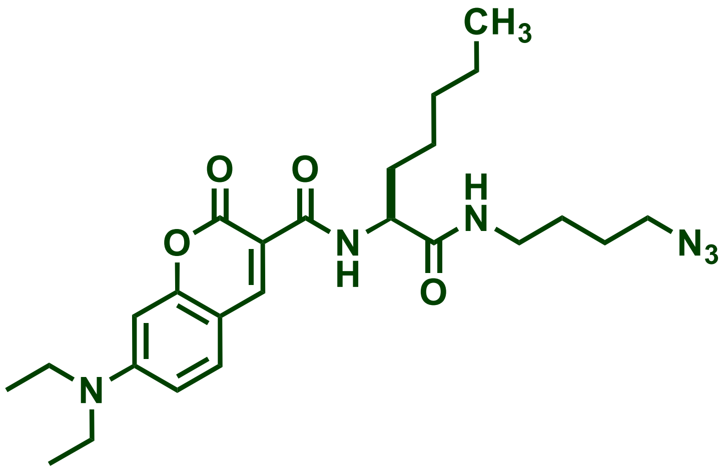


100 mg of chlorotrityl chloride resin (CTC resin) of loading capacity 1.5 mmol/g was swollen in dry DCM after which 1 eq Fmoc-Nle-OH (Nle = norleucine) was loaded on the resin as a solution of 5 mL of dry DCM mixed with 4 eq of dry DIPEA. The resin was shaken for 1 h at room temperature and washed 3X with a cocktail of DCM/MeOH/DIPEA (17:2:1) and then finally with DCM. The Fmoc group was removed using 20% piperidine and following wash steps with DCM and MeOH, 7-(diethylamino) coumarin-3-carboxylic acid was coupled to the norleucine using 4 equivalents of the coumarin and 4 equivalents each of Oxyma and DIC; the coupling reaction was done at room temperature with shaking for 2 h to make coumarin-Nle on trityl chloride resin. The resin was washed using alternating DCM and MeOH and finally rinsed with DCM. This coumarin-Nle conjugate was cleaved off the resin using 1% TFA in dry DCM. The cleavage cocktail was blown off and the resulting oil pellet was washed with ether 3X and dried. 4-azidobutylamine was coupled to the coumarin-Nle conjugate, in solution, using 2 equivalents of azidobutylamine, 2 eq. of HBTU and 4 eq. DIEA in DMF. The coupling reaction was done at room temperature with agitation for 2 h to make Coumarin-Nle-butyl azide. Excess DMF solvent was removed by rotary evaporation as a DMF-toluene azeotrope until dry powder was obtained. The powder was resuspended in nanopure water to make a white suspension. The suspension was centrifuged at 4000 x g for 10 min to separate the coupling reagents (aqueous layer) from the product (pellet). The pellet was resuspended

in 20% HPLC grade acetonitrile in water (0.1% TFA) and injected into preparative RP-HPLC for purification on a Phenomenex Luna 10 µm C8(2) 100 Å (250 x 21.2 mm) using a 20 to 100% linear gradient of MeCN in $H_2O$ each containing 0.1% TFA at 10 mL/min. ESI-MS $[M+H]^+$ m/z calcd. 471.2714 for $C_{24}H_{35}N_6O_4$, obsd. 471.2718. $^1$H NMR (600 MHz, $CDCl_3$) δ 0.89 (t, J= 6.0 Hz, 3H)1.18 (t, J= 6 Hz, 6H), 1.30-1.36 (m, 2H), 1.46-1.59 (m, 4H), 1.66-1.71 (m, 1H), 1.78-1.85 (m, 1H), 3.13-3.22 (m, 2H)  3.29 (t, J= 6 Hz, CH), 3.46 (q, J=  6 Hz, 1H, $CH_2$) , 4.40 (dq, J= 6, 12 Hz, 1H, CH), 6.55 (d, J= 2 Hz, 1H), 6.76 (dd, J= 2 and 12 Hz, 1H), 7.51 (d, J= 12 Hz, 1H), 8.60 (s, 1H), 9.03 (d, J= 12 Hz, 1H). $^{13}$C NMR (151 MHz, $CDCl_3$) δ 11.67, 13.23, 22.13, 25.83, 26.42, 27.49, 32.31, 32.34, 38.11, 38.23, 44.67, 50.78, 53.55, 53.58, 96.10, 108.05, 109.84, 110.19, 117.30, 131.34, 147.73, 152.92, 157.79, 162.51, 162.53, 171.62.

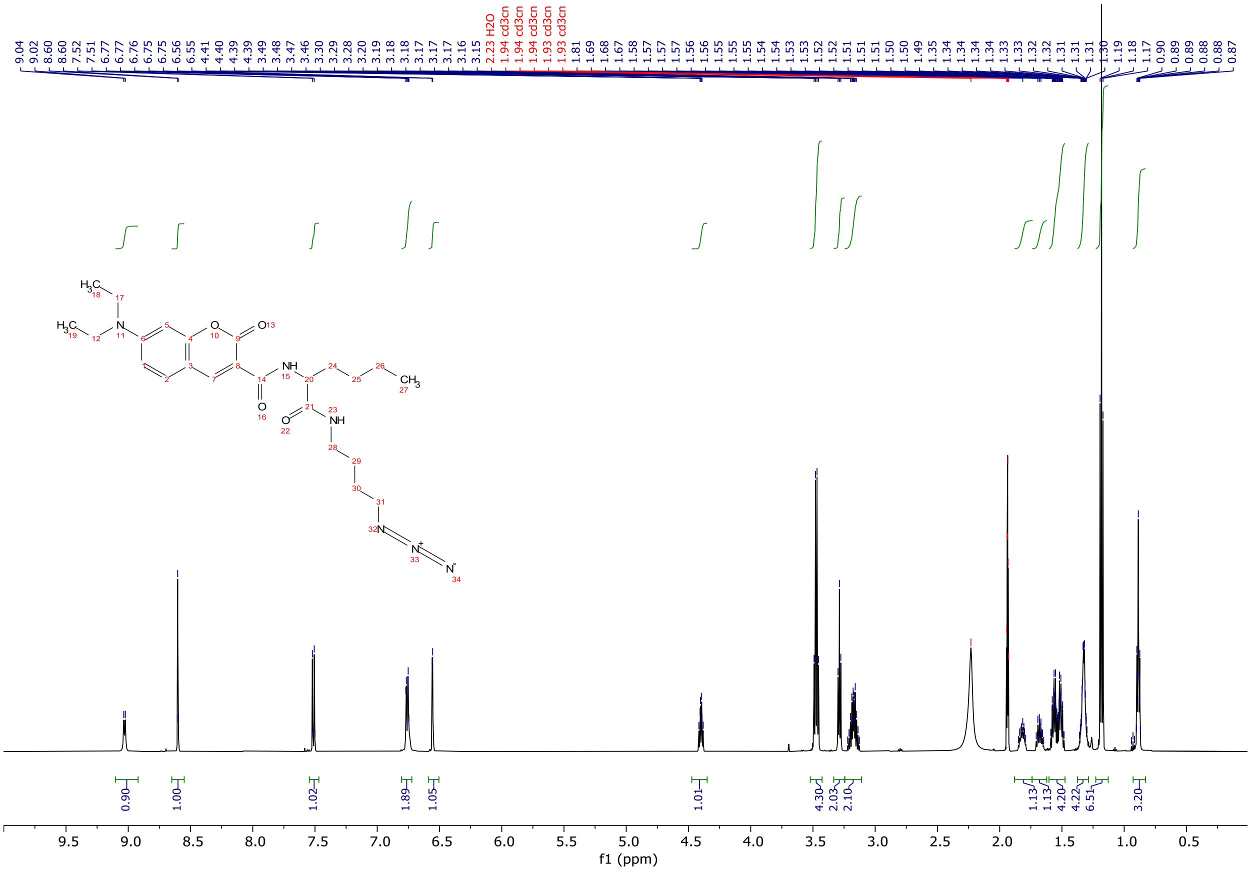

$^{1}$H-NMR of **6a** in $CDCl_3$.

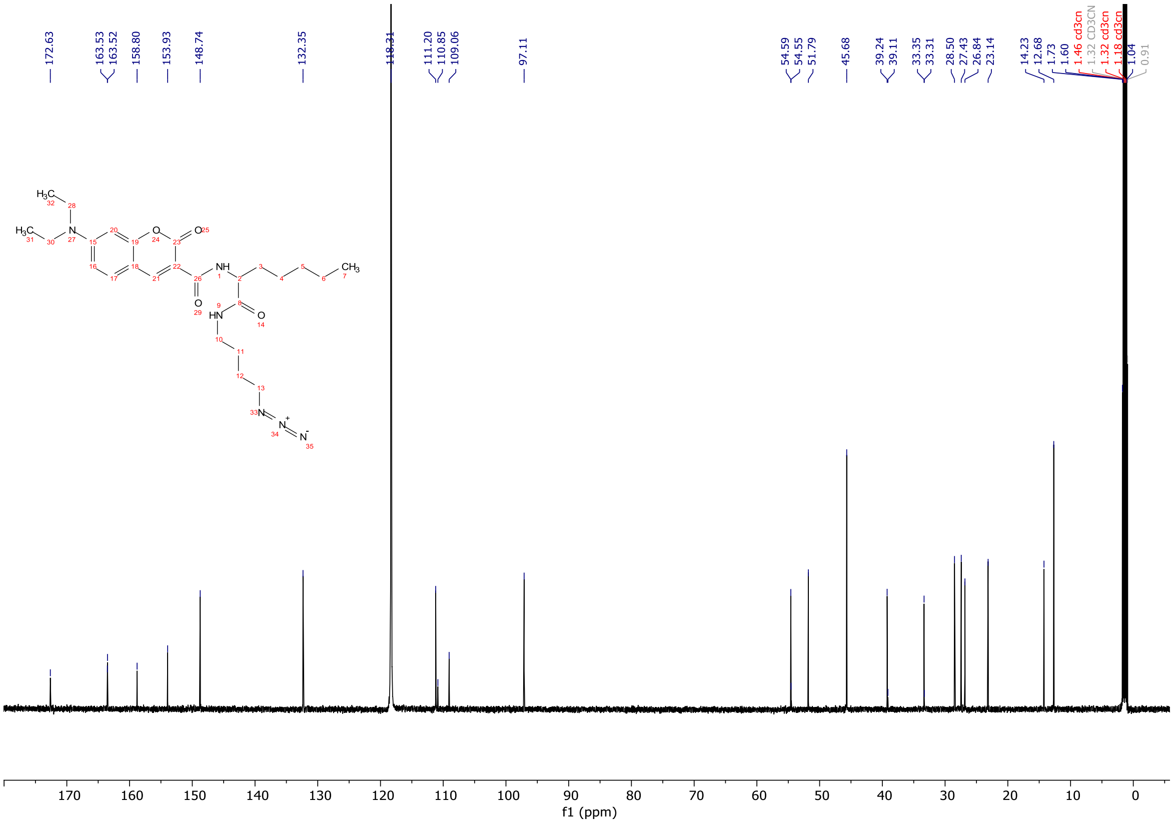


$^{13}$C-NMR of **6a** in $CDCl_3$.

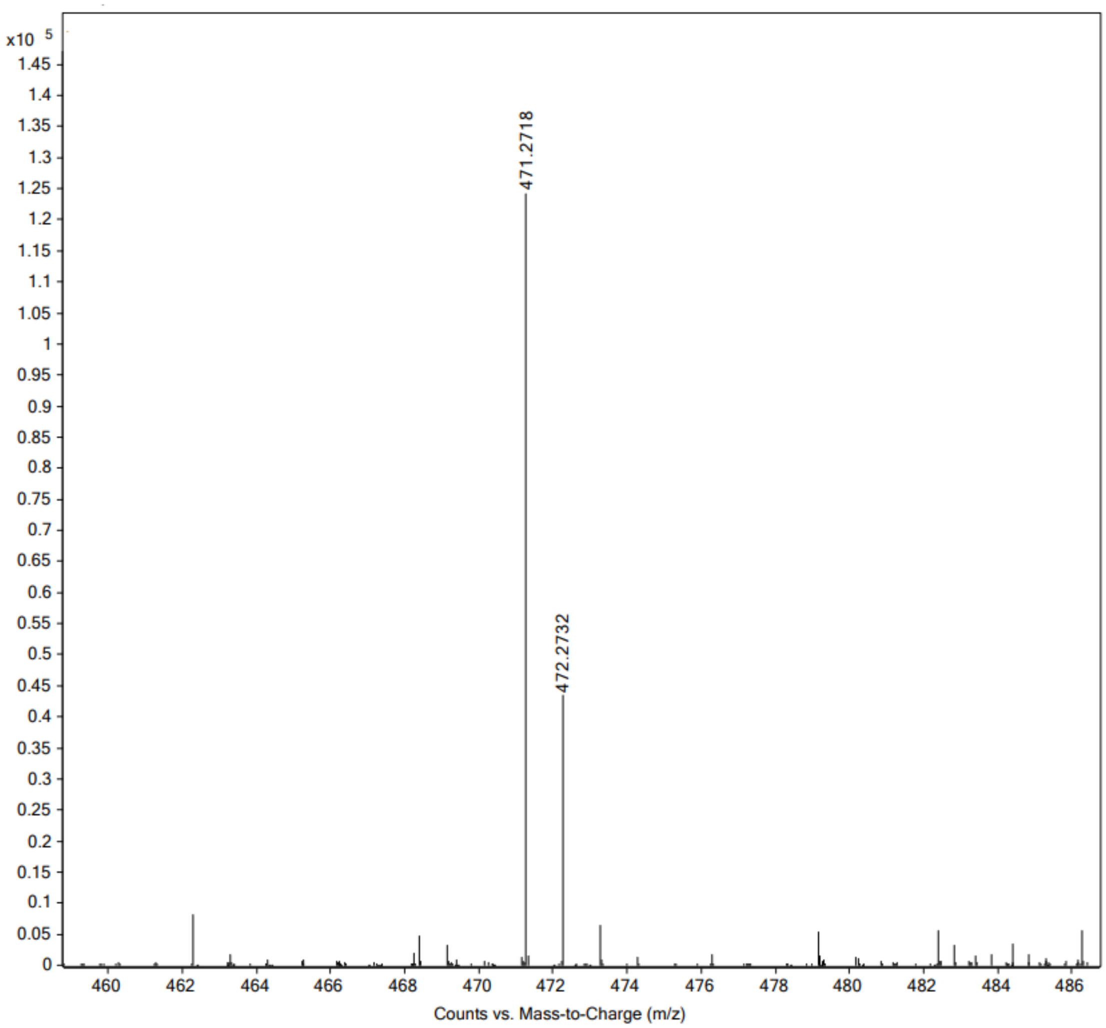


High resolution mass spectrum (ESI) of **6a**

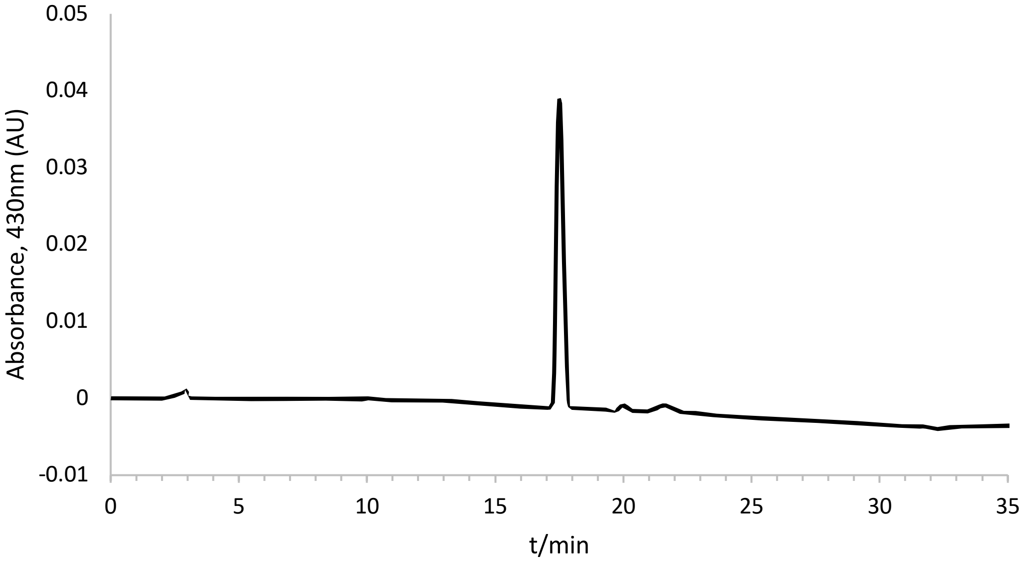


Analytical HPLC chromatogram of **6a**.

**Synthesis of 6b**.

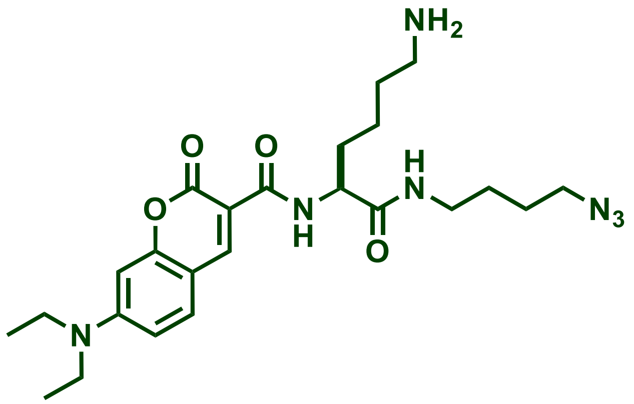


0.2 mol of a-Fmoc-e-$^{t}$Boc-protected lysine (Me) amino acid was mixed with 1-amino-4-butyl azide (0.22 mmol) in $CH_2Cl_2$ (2.0 mL) at room temperature. To this solution was added EDC (0.25 mmol) and the mixture was stirred at RT followed by LC-MS analysis upon completion of the reaction. The mixture was further diluted with $CH_2Cl_2$ (20 mL) and washed with water, dil. $NaHCO_3$ and brine. The $CH_2Cl_2$ layer was filtered through a $Na_2SO_4$ plug on a funnel and concentrated under reduced pressure to give a solid residue which was carried forward to the next reaction without purification or characterization. The left-over residue was dissolved in methanol (2 mL) and to it, was added LiOH (5M, 0.5 mL), the mixture was stirred at RT overnight and then, the next day, the reaction mixture was analyzed by mass spectrometry to confirm that the starting material was completely consumed to afford the desired Fmoc-deprotected amine. The mixture was carefully acidified with dil. HCl, concentrated under reduced pressure, followed by freezing and lyophilization to yield a solid residue which was used as is for the next reaction. The amine residue was dissolved in $CH_2Cl_2$ and then coumarin (0.18 mmol) mixed with EDC (0.2 mmol) was added to the mixture followed by stirring at RT; the progress of the reaction was followed by LC-MS. Upon formation of the product (after 4-6hrs), the reaction mixture was further diluted with $CH_2Cl_2$ (20 mL) and washed with water, dil. $NaHCO_3$ and brine. The $CH_2Cl_2$ layer was filtered through a $Na_2SO_4$ plug on a funnel and concentrated under reduced pressure to give a solid residue which was Boc-deprotected upon treatment with conc. HCl (0.5 mL) at RT overnight. The next day, the reaction mixture was diluted with acetonitrile (5 mL) and carefully neutralized with $Na_2SO_4$. The solid was removed by filtration through a 0.2 mm filter, and the remaining solution was injected into preparative RP-HPLC for purification on a Phenomenex Luna 10 μm C8(2) 100 Å (250 x 21.2 mm) using a 20 to 100% linear gradient of MeCN in $H_2O$ each containing 0.1% TFA at 10 mL/min. ESI-MS $[M+H]^+$ m/z calcd. 486.2823 for $C_{25}H_{36}N_7O_4$, obsd. 486.2824. $^1$H-NMR (600 MHz, $CD_3CN$) d 1.18 (t, J= 6 Hz, 6H), 1.41-1.58 (m, 4H), 1.67-1.75 (m, 1H), 1.48-1.90 (m, 1H), 2.93-3.02 (m, 2H) 3.19 (q, J= 6Hz, 2H) 3.29 (t, J= 6 Hz, CH), 3.46 (q, J= 6 Hz, 1H, $CH_2$) , 4.45 (d, J= 6 Hz, 1H), 6.50 (d, J= 2 Hz, 1H), 6.74 (dd, J= 2 and 12 Hz, 1H), 6.80 (brs, 1H) 7.06 (brt, J= 12 Hz, 1H), 8.55 (s, 1H, ArH), 9.15 (d, J= 6 Hz, 1H).

$^{13}$C NMR (151 MHz, $CD_3CN$) δ 11.69, 21.95, 25.73, 26.16, 31.97, 38.34, 39.73, 44.74, 50.77, 108.01, 109.18, 110.35, 131.41, 147.91, 153.06, 157.76, 162.51, 171.88.

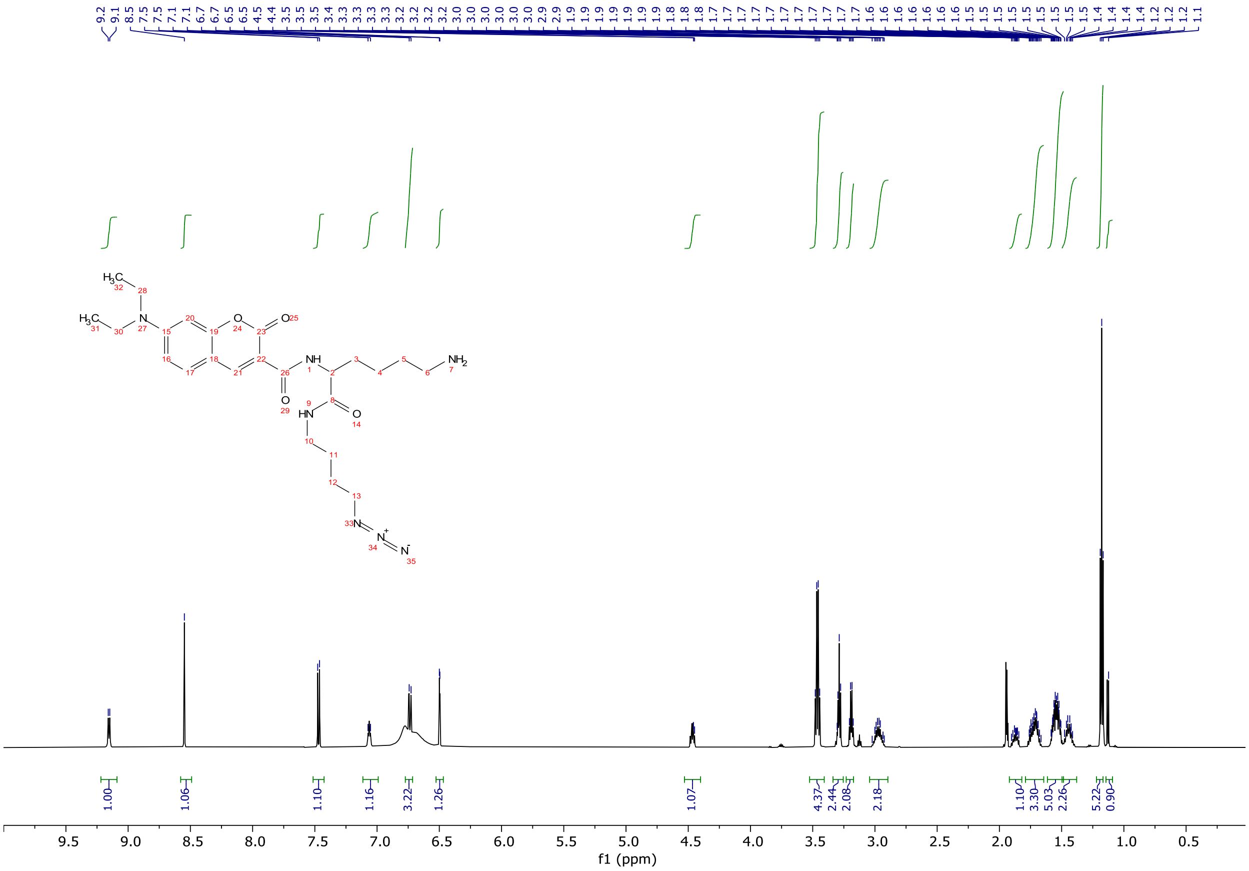

[1]H-NMR of **6b** in $CD_3CN$.

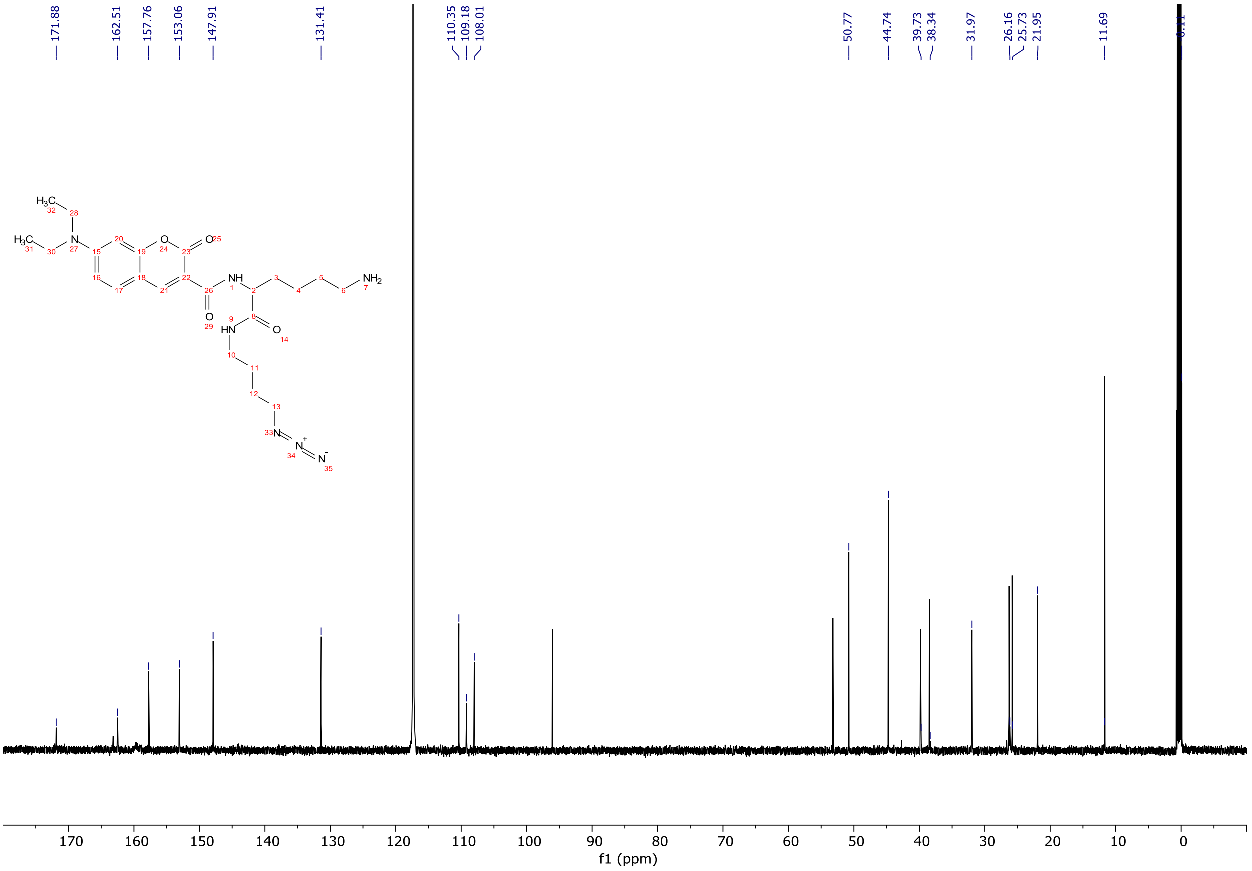


$^{13}$C-NMR of **6b** in $CDCl_3$.

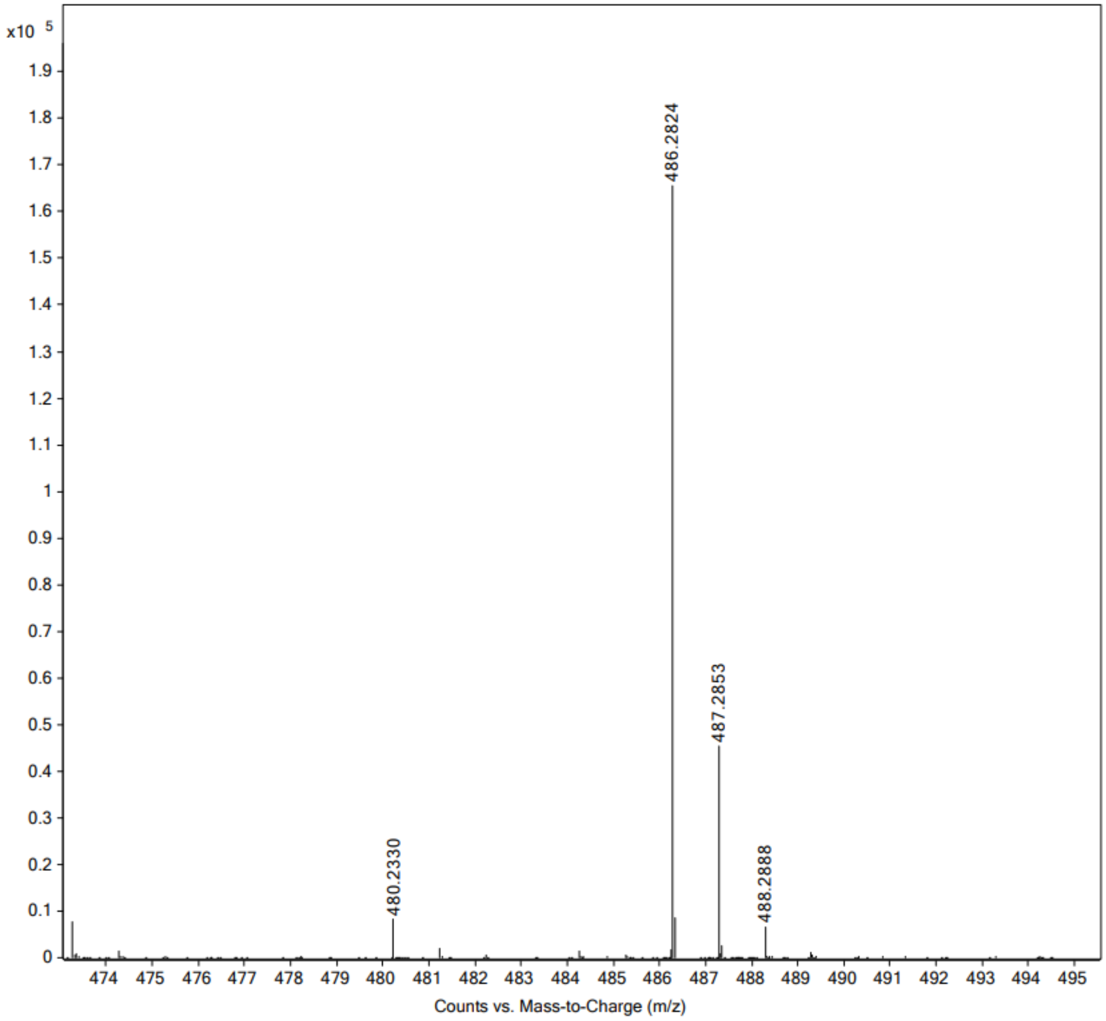


High resolution mass spectrum (ESI) of **6b**.

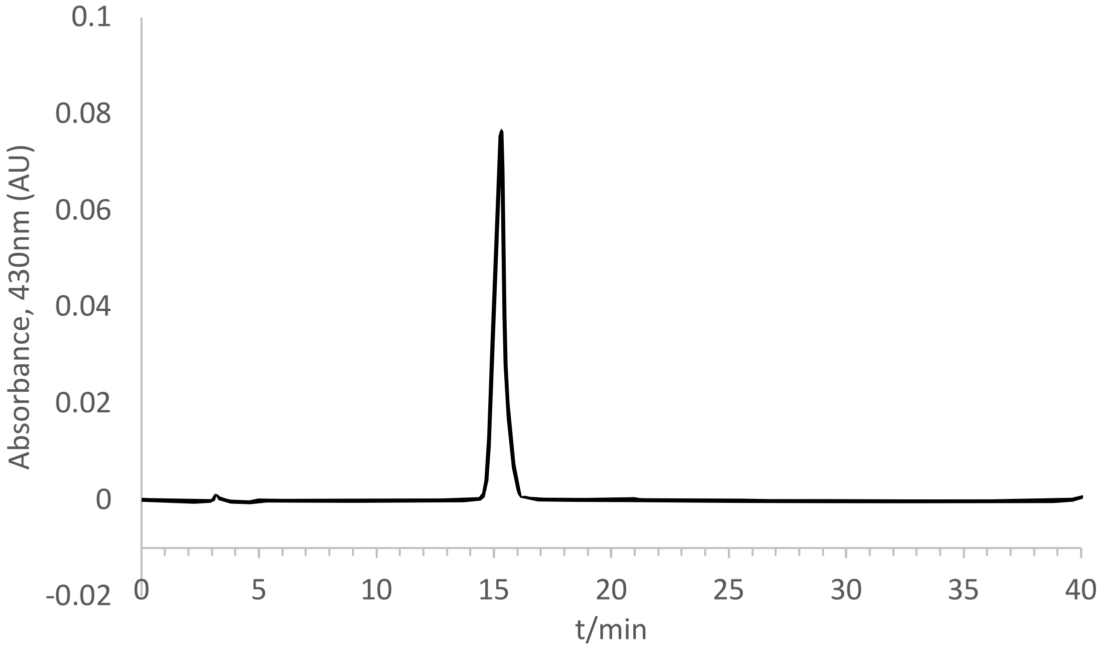


Analytical HPLC chromatogram of **6b**.

**Synthesis of 6c.**

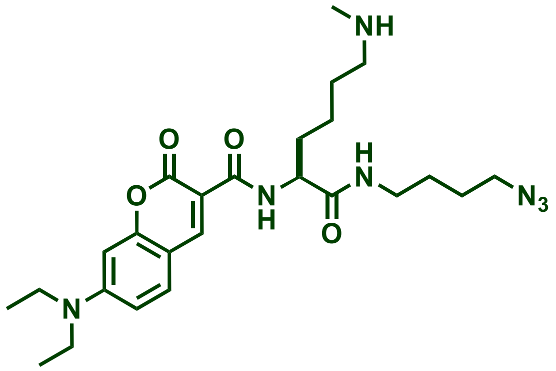


0.2 mmol of a-Fmoc-protected lysine (Me)$_2$ amino acid (96.4mg) was mixed with 1-amino-4-butyl azide (29 mg, 0.25mmol) in $CH_2Cl_2$ (2.0 mL) at room temperature. To this solution was added EDC (0.25 mmol) and the mixture was stirred at RT followed by LC-MS analysis upon completion of the reaction. The mixture was further diluted with $CH_2Cl_2$ (20 mL) and washed with water, dil. $NaHCO_3$ and brine. The $CH_2Cl_2$ layer was filtered through a $Na_2SO_4$ plug on a funnel and concentrated under reduced pressure to give a solid residue which was carried forward to the next reaction without purification or characterization. The left-over residue was dissolved in methanol (2 mL) and to it was added LiOH (5 M, 0.5 mL), the mixture was stirred at RT overnight and then, the next day, the reaction mixture was analyzed by mass spectrometry to confirm that the starting material was completely consumed to afford the desired Fmoc-deprotected amine. The mixture was carefully neutralized with dil. HCl (0.1 M), concentrated under reduced pressure, followed by freezing and lyophilization to yield a solid residue which was used as is for the next reaction. The amine residue was dissolved in $CH_2Cl_2$ and then coumarin (0.2 mmol) mixed with EDC (0.25mmol) was added to the mixture followed by stirring at RT; the progress of the reaction was followed by LC-MS. Upon formation of the product after 6hrs, the reaction mixture was further diluted with $CH_2Cl_2$ (20 mL) and washed with water, dil. $NaHCO_3$ and brine. The $CH_2Cl_2$ layer was filtered through a $Na_2SO_4$ plug on a funnel and concentrated under reduced pressure to give a residue. This residue was dissolved in 200 μL of MeCN and treated concentrated HCl (0.5mL), the mixture was stirred overnight, the LCMS analysis indicated deprotection of Boc group and formation of desired product. The mixture was diluted with 3.0 mL MeCN and concentrated under reduced pressure. Left over residue diluted with $CHCl_3$ (3 mL), carefully neutralized by adding solid $Na_2CO_3$, and solution was filtered through bed of celite, concentrated and purified as follows: The residue was dissolved in 1:1 (MeCN:$H_2O$, ~10mL). The solution was filtered through a 0.2 m filter, and filtrate was injected into preparative RP-HPLC for purification on a Phenomenex Luna 10 µm C8(2) 100 Å (250 x 21.2 mm) using a 20 to 100% linear gradient of MeCN in $H_2O$ each containing 0.1% TFA at 10 mL/min. Homogenous fractions showing absorbance at 220 and 410nm were collected, concentrated under reduced pressure using rotary evaporator to remove as much MeCN as possible, remaining solution was frozen at -80 °C overnight. Next day the frozen solution was lyophilized to obtained dried light orange solid. ESI-MS $[M+H]^+$ m/z 500.2980 calcd. 500.2980 for $C_{25}H_{38}N_7O_4$ obsd. 500.2981. $^1$H-NMR (600 MHz, $CD_3CN$) d 1.38 (t, J= 6 Hz, 6H), 1.41-1.58 (m, 4H), 1.67-1.75 (m, 1H), 1.48-1.90 (m, 1H), 2.94 (brt, 3H), 3.19-3.24 (m, 2H) 3.41 (q, J= 6Hz, 2H) 3.52 (t, J= 6 Hz, CH), 3.68 (q, J= 6 Hz, 4H, $CH_2$) , 4.68 (q, J= 6 Hz, 1H), 6.72 (s,1H), 6.95 (d, J= 6 Hz, 1H) 7.32 (brt, J= 12 Hz, 1H), 7.42 (brs, 1H), 7.69 (d, J= 6 Hz, 1H), 8.13 (brs, 1H), 8.76 (s, 1H), 9.38 (d, J= 6 Hz, 1H). $^{13}$C NMR (151 MHz, $CD_3CN$) δ 12.40, 22.81, 25.69, 26.55, 26.99, 32.63, 33.75, 39.28, 45.52, 49.91, 51.50, 54.01, 96.86, 108.78, 109.86, 111.14, 115.59, 132.17, 148.64, 153.79, 158.50, 163.23, 163.97, 172.75.

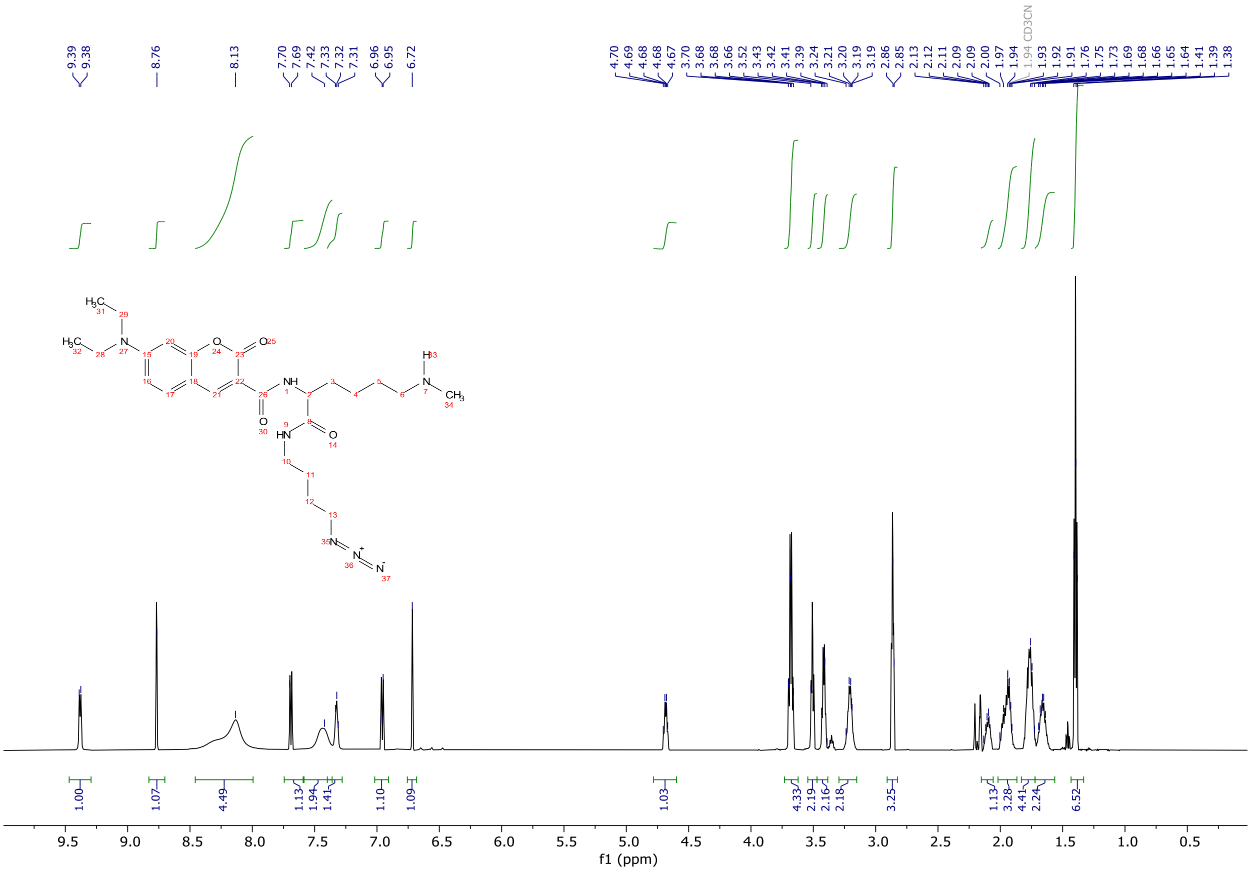


[1]H-NMR of **6c** in $CD_3CN$.

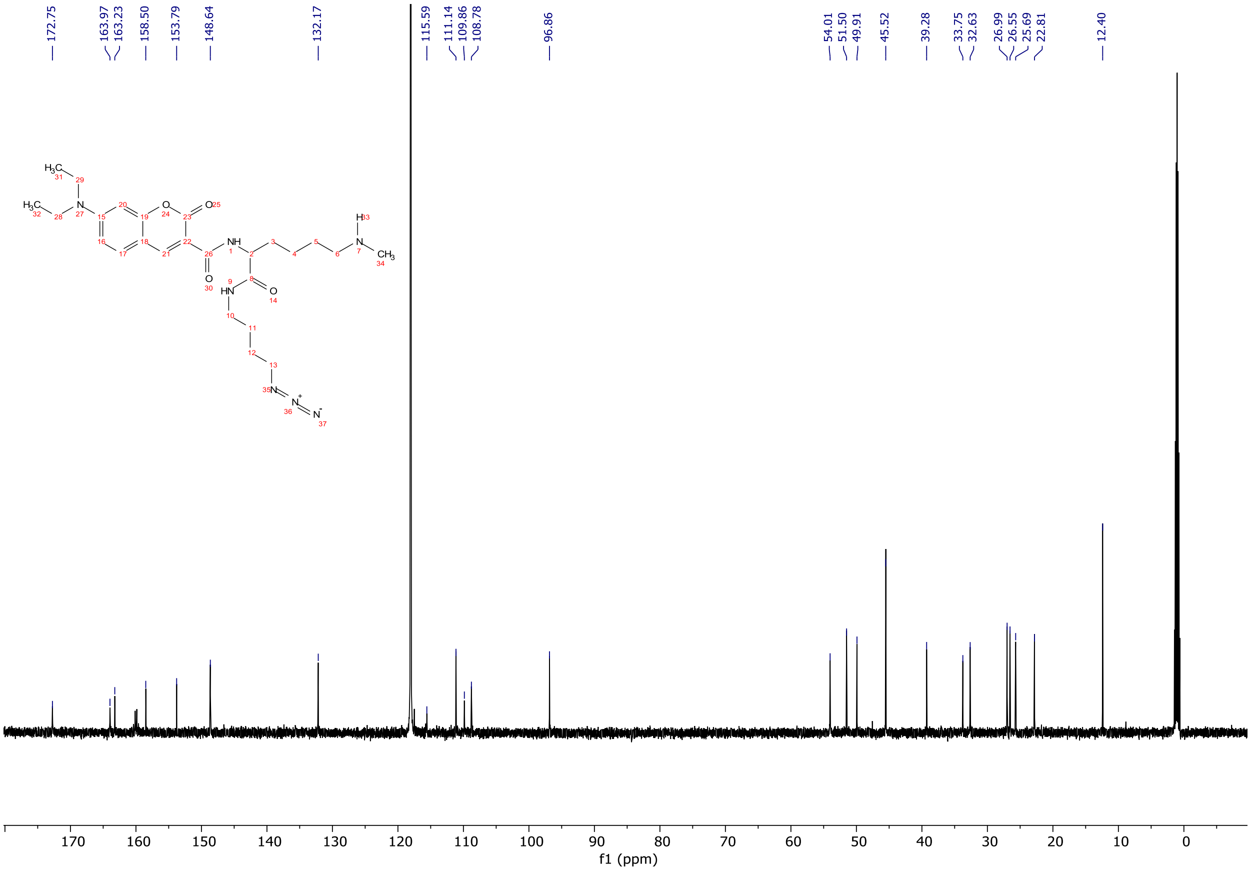


$^{13}$C-NMR of **6c** in $CD_3CN$.

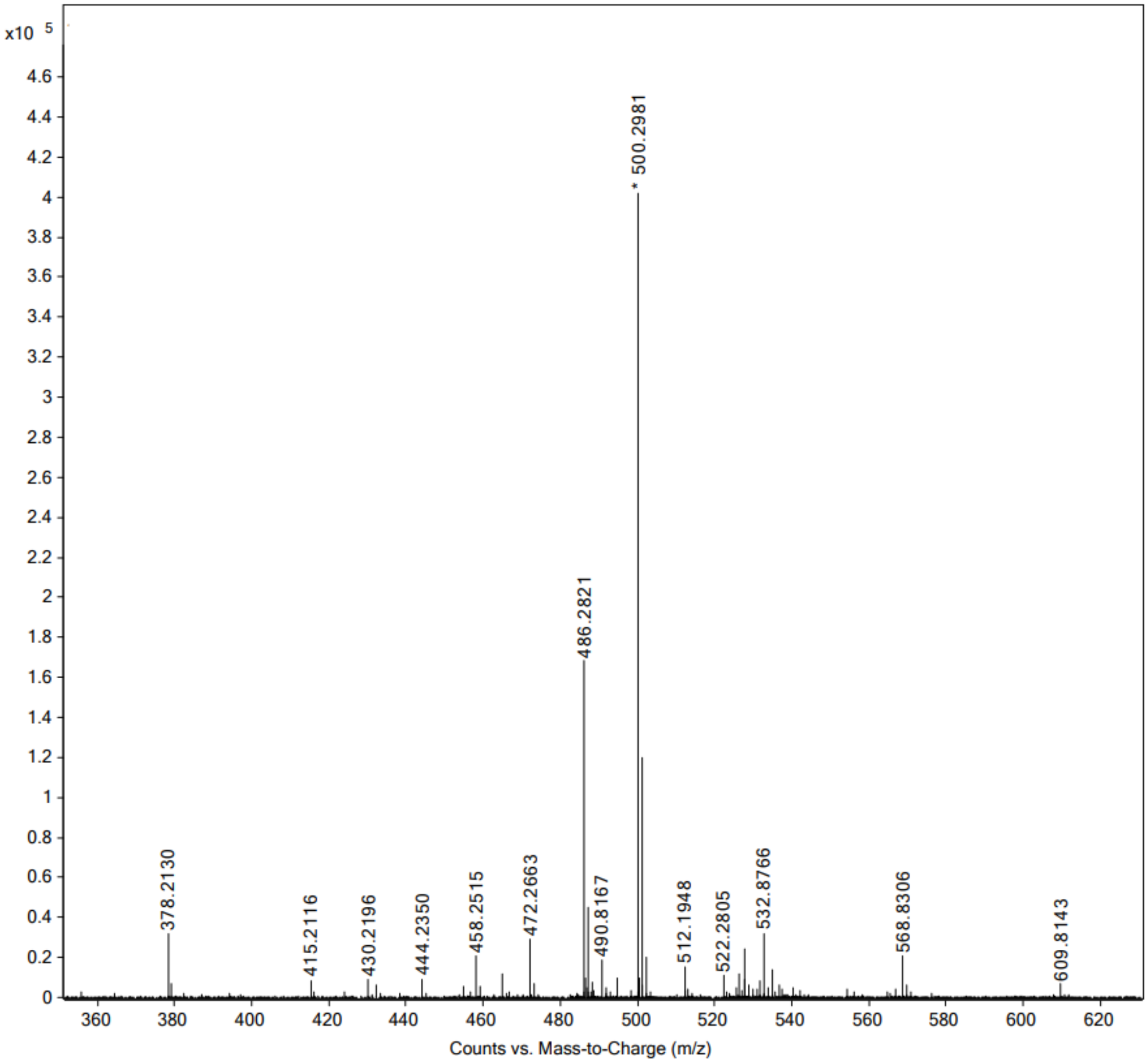


High resolution mass spectrum (ESI) of **6c**.

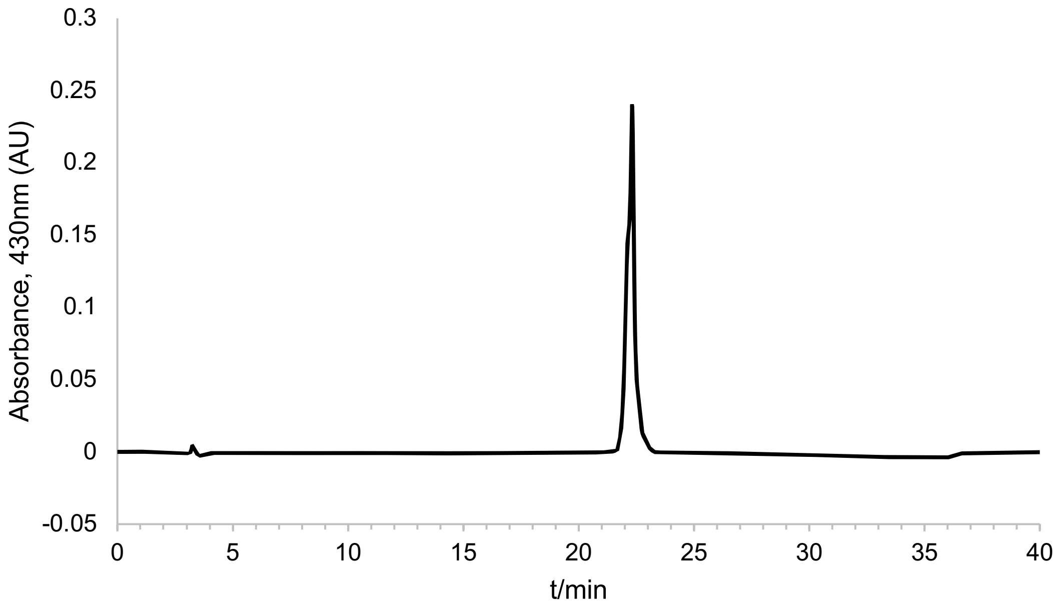


Analytical HPLC chromatogram of **6c**.

**Synthesis of 6d.**

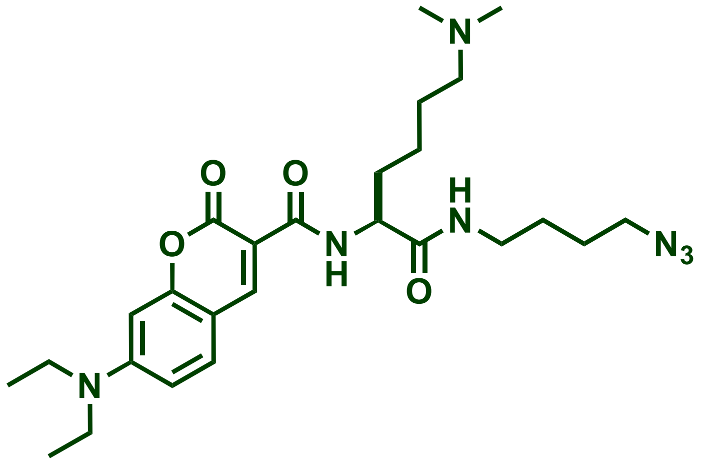


0.2 mmol of a-Fmoc-protected lysine $(Me)_2$ amino acid (79.2mg) was mixed with 1-amino-4-butyl azide (29 mg, 0.25 mmol) in $CH_2Cl_2$ (2.0 mL) at room temperature. To this solution was added EDC (0.25 mmol) and the mixture was stirred at RT followed by LC-MS analysis upon completion of the reaction. The mixture was further diluted with $CH_2Cl_2$ (20 mL) and washed with water, dil. $NaHCO_3$ and brine. The $CH_2Cl_2$ layer was filtered through a $Na_2SO_4$ plug on a funnel and concentrated under reduced pressure to give a solid residue which was carried forward to the next reaction without purification or characterization. The left-over residue was dissolved in methanol (2 mL) and to it was added LiOH (5 M, 0.5 mL), the mixture was stirred at RT overnight and then, the next day, the reaction mixture was analyzed by mass spectrometry to confirm that the starting material was completely consumed to afford the desired Fmoc-deprotected amine. The mixture was carefully acidified with dil. HCl, concentrated under reduced pressure, followed by freezing and lyophilization to yield a solid residue which was used as is for the next reaction. The amine residue was dissolved in $CH_2Cl_2$ and then coumarin (0.18 mmol) mixed with EDC (0.2 mmol) was added to the mixture followed by stirring at RT; the progress of the reaction was followed by LC-MS. Upon formation of the product after 6hrs, the reaction mixture was further diluted with $CH_2Cl_2$ (20 mL) and washed with water, dil. $NaHCO_3$ and brine. The $CH_2Cl_2$ layer was filtered through a $Na_2SO_4$ plug on a funnel and concentrated under reduced pressure to give a residue. The residue was dissolved in 1:1 (MeCN:$H_2O$, 8mL). The solution was filtered through a 0.2 m filter, and filtrate was injected into preparative RP-HPLC for purification on a Phenomenex Luna 10 μm C8(2) 100 Å (250 x 21.2 mm) using a 20 to 100% linear gradient of MeCN in $H_2O$ each containing 0.1% TFA at 10 mL/min. Homogenous fractions showing absorbance at 220 and 410nm were collected, concentrated under reduced pressure using rotary evaporator to remove as much MeCN as possible, remaining solution was frozen at -80 $^oC$ overnight. Next day the frozen solution was lyophilized to obtained dried light orange solid. ESI-HRMS $[M+H]^+$ m/z calcd. 514.3136 for $C_{26}H_{40}N_7O_4$, obsd. 514.3137. $^1H$-NMR (600 MHz, $CD_3CN$) 1.19 (t, J= 6 Hz, 6H), 1.36-1.41 (m, 2H), 1.51-1.58 (m, 4H), 1.70-1.77 (m, 3H), 1.85-1.90 (m, 1H), 2.78 (brs, 6H), 2.99-3.06 (m, 2H) 3.19 (q, J= 6Hz, 2H) 3.29 (t, J= 6 Hz, 2H), 3.47 (q, J= 6 Hz, 4H, $CH_2$), 4.47 (d, J= 6 Hz, 1H), 6.53 (d, J= 2 Hz, 1H), 6.75 (dd, J= 2 and 12 Hz, 1H), 6.99 (brs, 1H), 7.50 (d, J= 6, 1H), 8.58 (s, 1H, ArH), 8.70 (brs, 1H), 9.11 (d, J= 6 Hz, 1H). $^{13}C$ NMR (151 MHz, $CD_3CN$) δ 12.69, 22.64, 23.08, 24.57, 26.84, 27.35, 32.95, 39.42, 43.69, 43.76, 45.73, 51.79, 54.14, 58.54, 97.10, 109.04, 110.43, 110.46, 111.32, 118.33, 132.42, 148.87, 154.04, 158.81, 160.36, 160.61, 163.52, 163.92, 172.62.

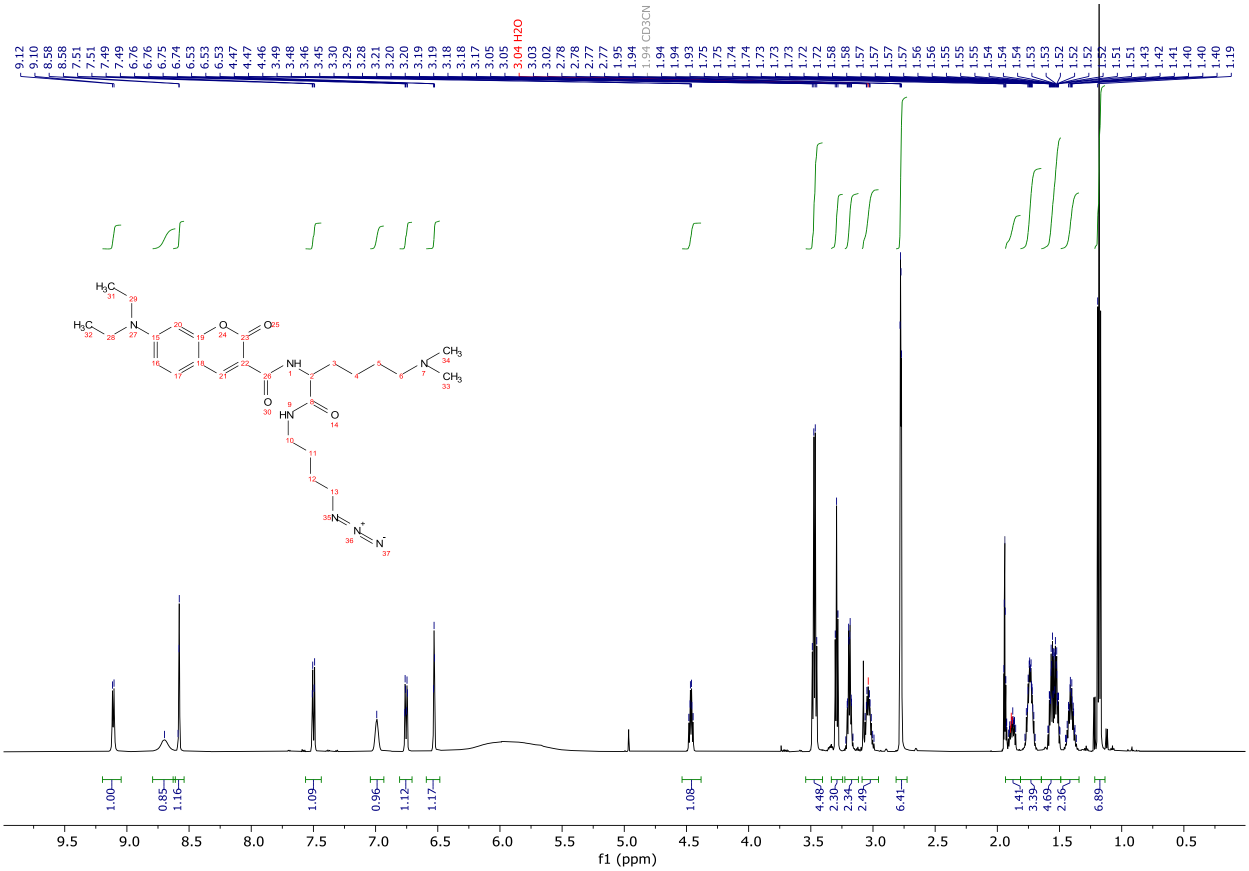

$^1$H-NMR of **6d** in $CD_3CN$.

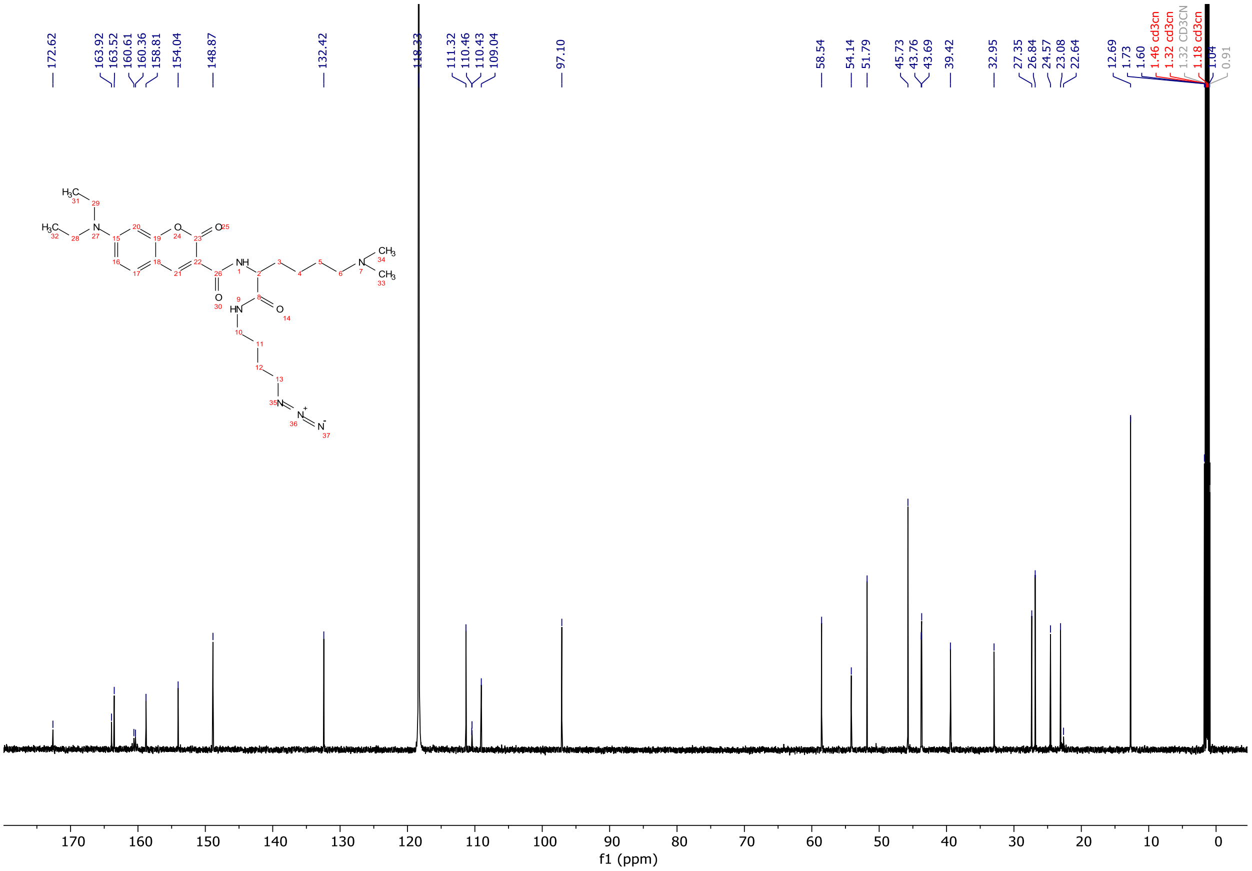

$^{13}$C-NMR of **6d** in $CD_3CN$.

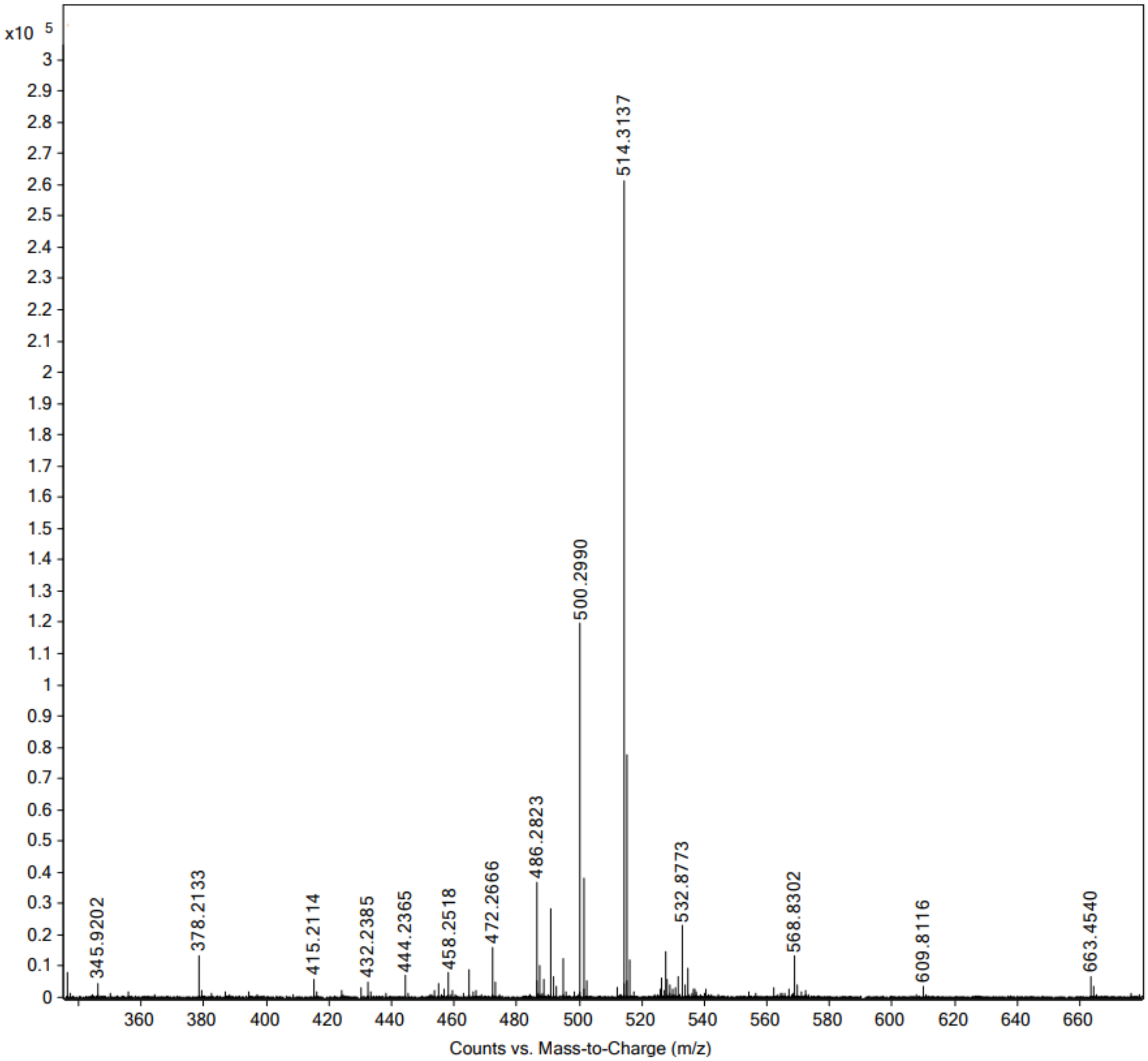


High resolution mass spectrum (ESI) of **6d**.

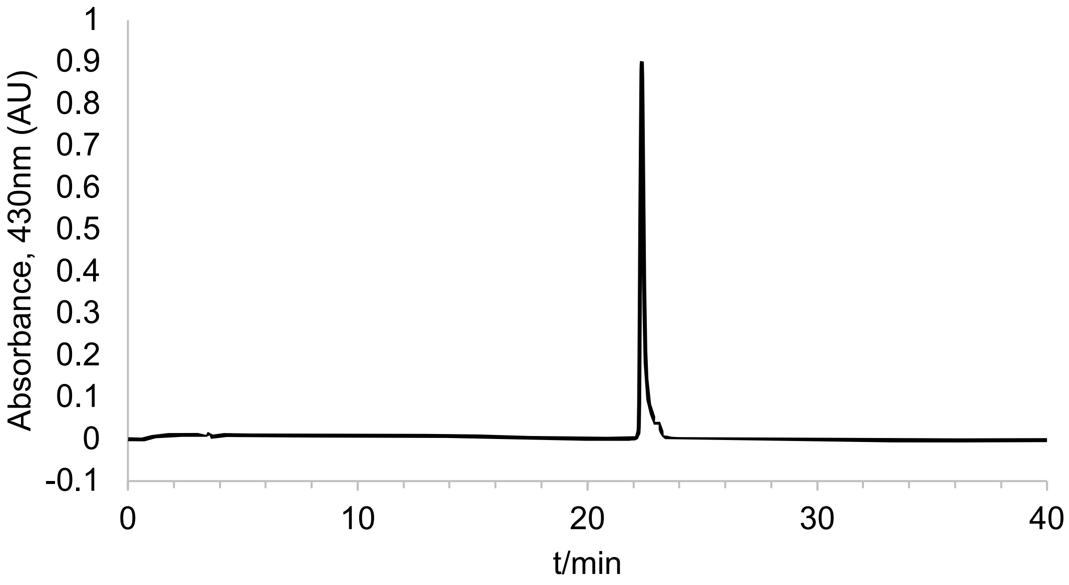


Analytical HPLC chromatogram of **6d**.

**Synthesis of 6e**.

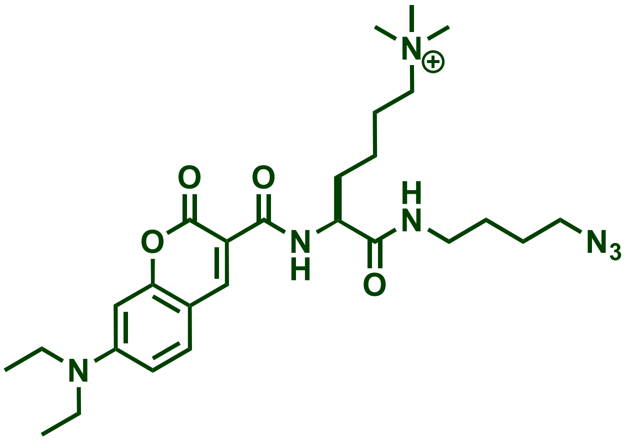

The solution of compound **6d** (25.5 mg, 0.05 mmol) was dissolved in MeCN (1 mL) and solid $NaHCO_3$ (20.0 mg) was added to it and sealed with rubber septum. To this mixture methyl iodide (50 μL) was added via syringe and the mixture was stirred overnight at RT. Next the LCMS analysis indicated starting material was consumed to form the desired quaternary ammonium product. The mixture was concentrated under reduced pressure to give a residue, which was dissolved in minimum amount of $CHCl_3$ (~100μL) and loaded on preparative TLC plate (1000 m, Silica Gel, Analtech), the plate was run with $CHCl_3$:MeOH (95:5)The lowest fluorescent band observed by UV light was removed and eluted with ($CHCl_3$:MeOH; 9:1, 0.5mL x 3). The combined eluent was concentrated under reduced pressure to yield orange oily residue, which analyzed by analytical HPLC and characterized by $^1H$, $^{13}C$-NMR and HRMS. ESI-HRMS $[M]^+$ m/z calcd. 528.3293 for $C_{27}H_{42}N_7O_4$, obsd. 528.3293. $^1H$-NMR (600 MHz, $CD_3CN$) 1.18 (t, J= 6 Hz, 6H), 1.38-1.42 (m, 2H), 1.55-1.61 (m, 4H), 1.76-1.83 (m, 3H), 2.38 (m, 2H), 3.10 (brs, 9H), 3.16-3.23 (m, 2H) 3.19 (q, J= 6Hz, 2H), 3.32 (t, J= 6 Hz, 2H), 3.47 (q, J= 6 Hz, 4H, $CH_2$), 4.61 (m, 1H), 6.54 (brs, 1H), 6.77 (dd, J= 2 and 12 Hz, 1H), 7.40 (brs, 1H), 7.52 (d, J= 6, 1H), 8.08 (s, 0.5H), 8.61 (s, 1H), 9.11 (d, J= 6 Hz, 1H). $^{13}C$ NMR (151 MHz, $CD_3CN$) δ 10.43, 11.72, 13.33, 21.91, 22.22, 22.66, 23.72, 25.92, 26.35, 28.69, 30.28, 32.20, 38.28, 38.77, 44.71, 50.81, 53.01, 66.24, 67.37, 96.05, 108.02, 110.29, 129.38, 131.40, 147.91, 152.98, 157.77, 162.49, 162.75, 171.49.

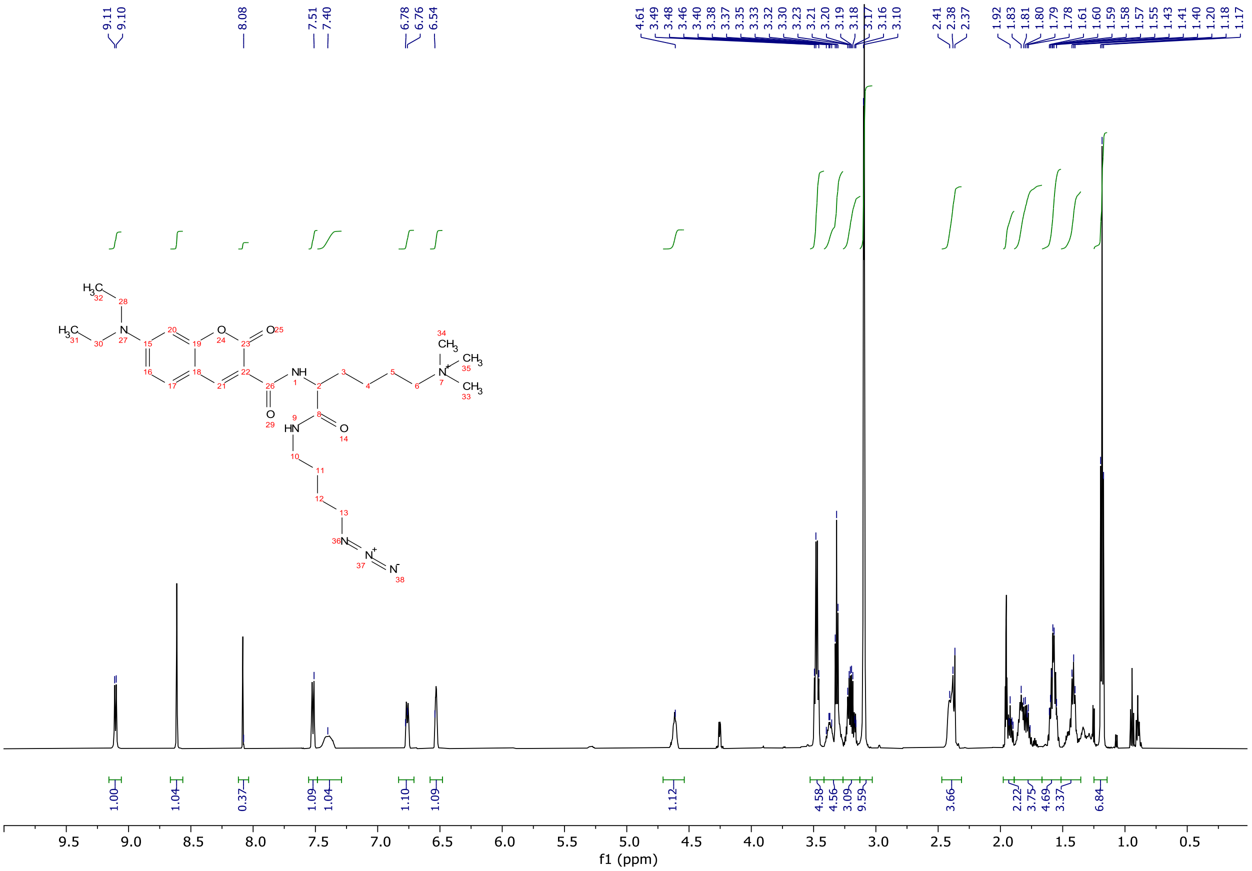

$^1$H-NMR of **6e** in $CD_3CN$.

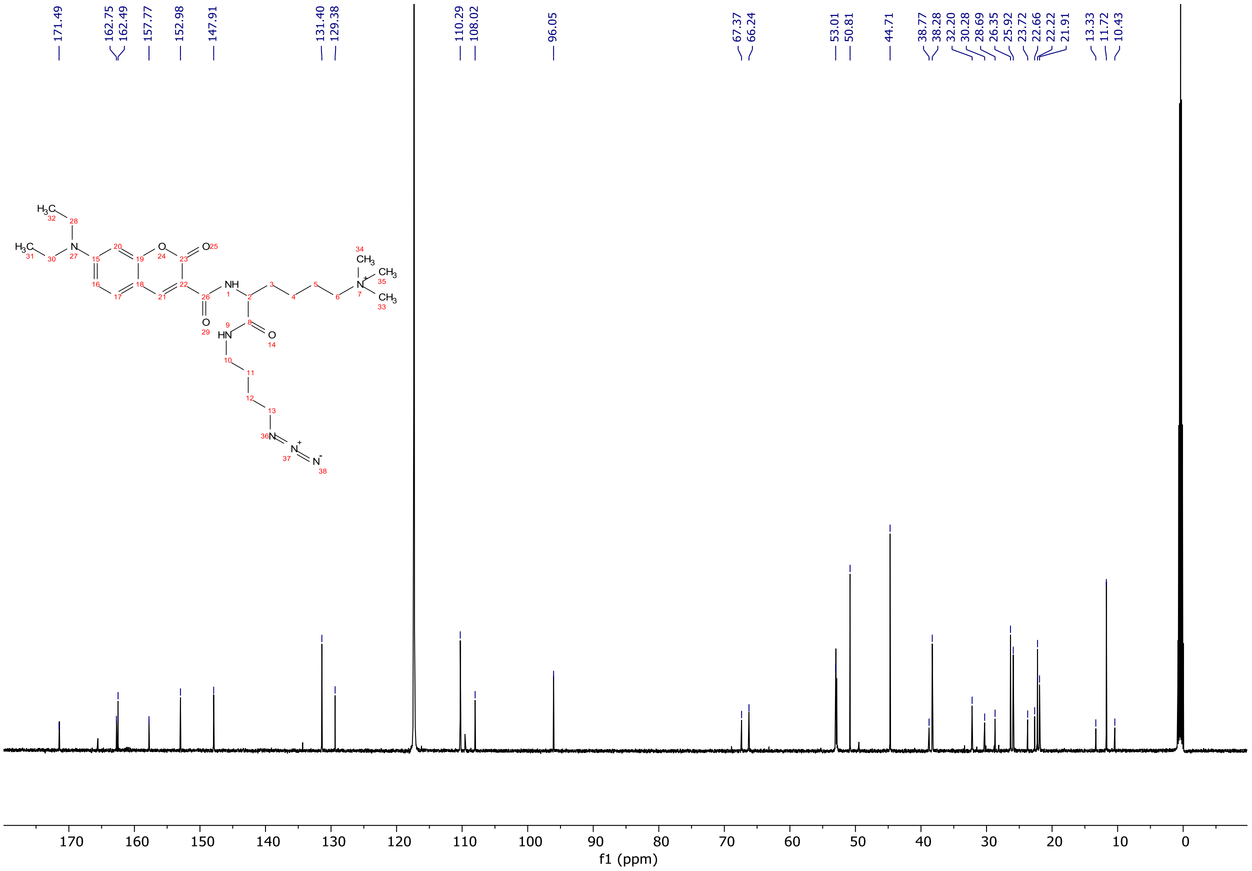


$^{13}$C-NMR of **6e** in $CD_3CN$.

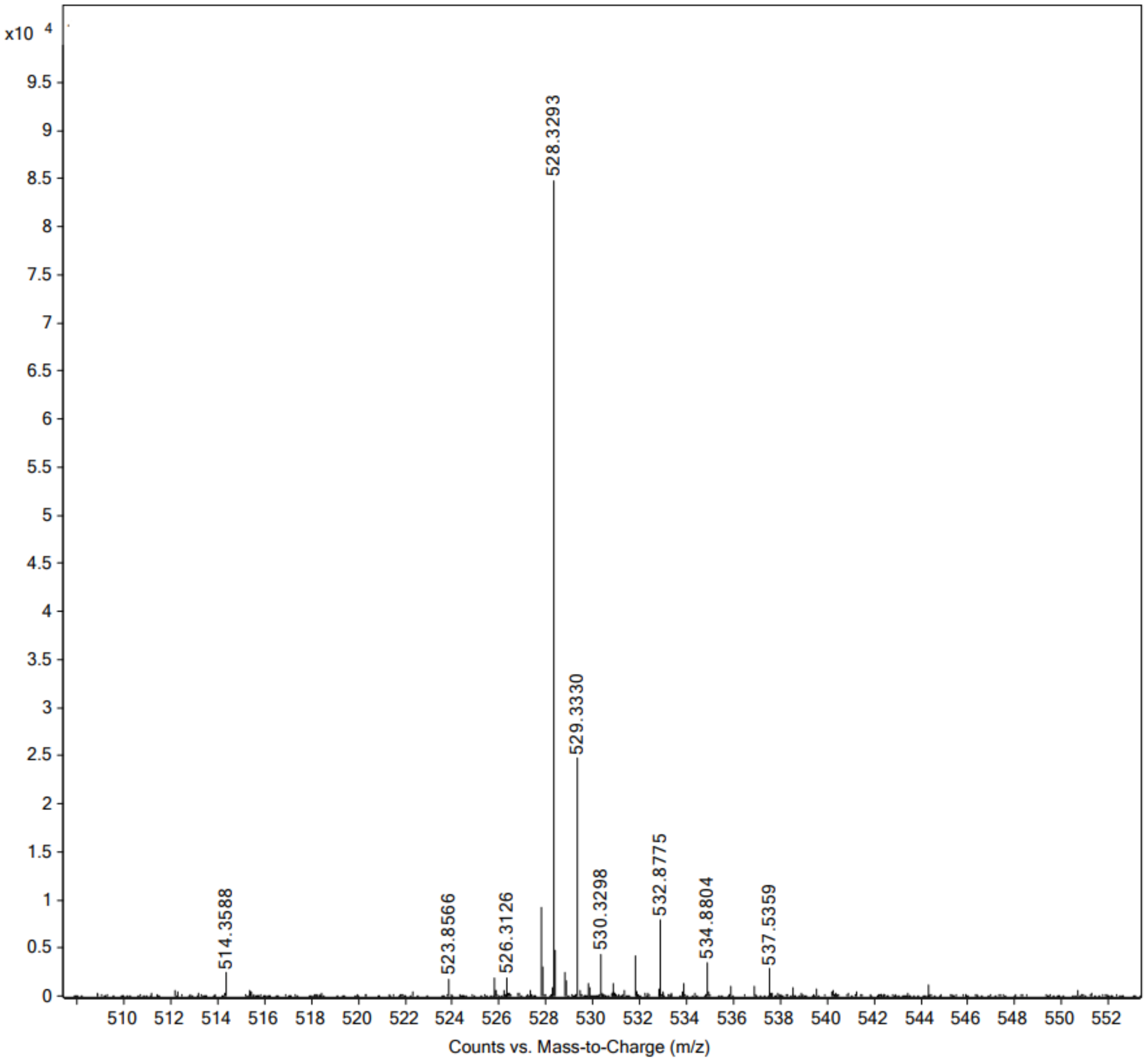


High resolution mass spectrum (ESI) of **6e**.

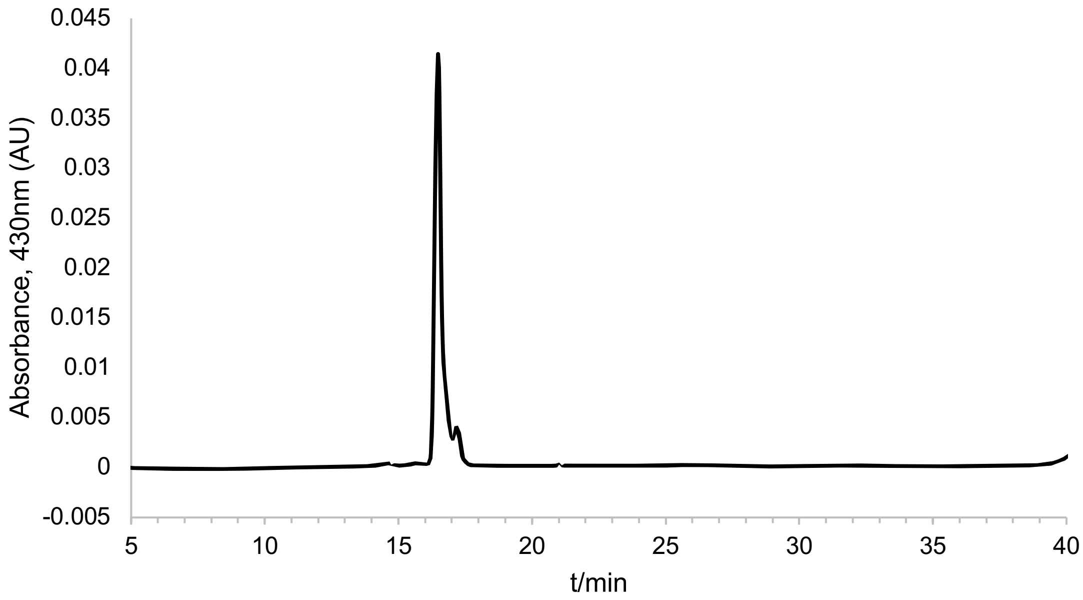


Analytical HPLC chromatogram of **6e**.

**1152 Library of azido-tagged molecules (Dong/Sharpless Library)**

**Plate 1**

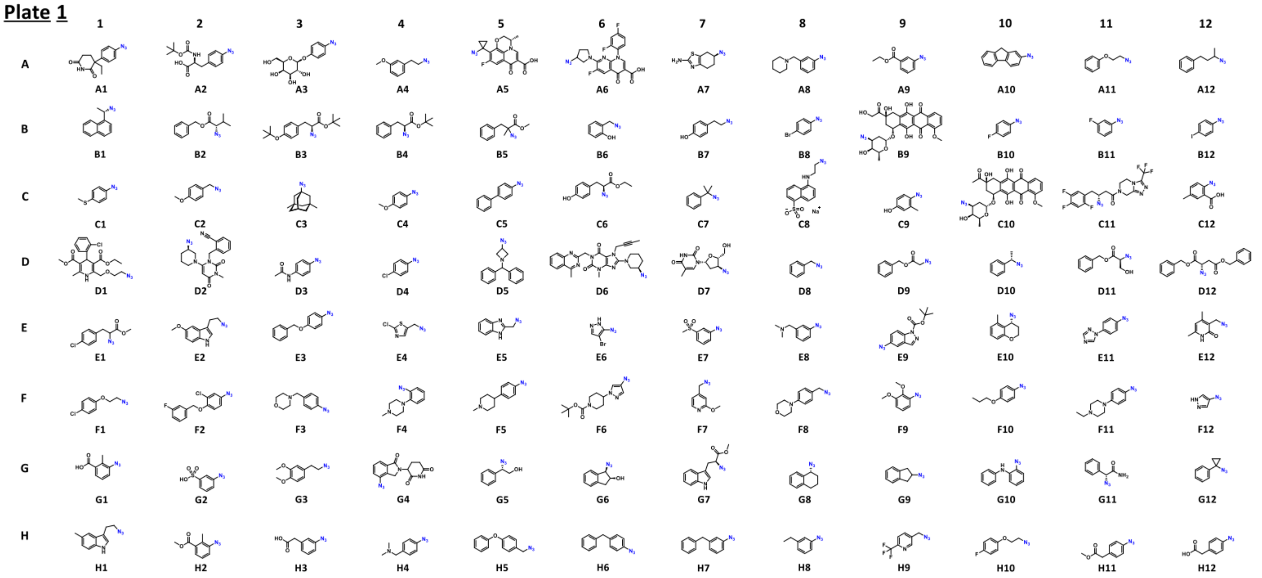


**Plate 2**

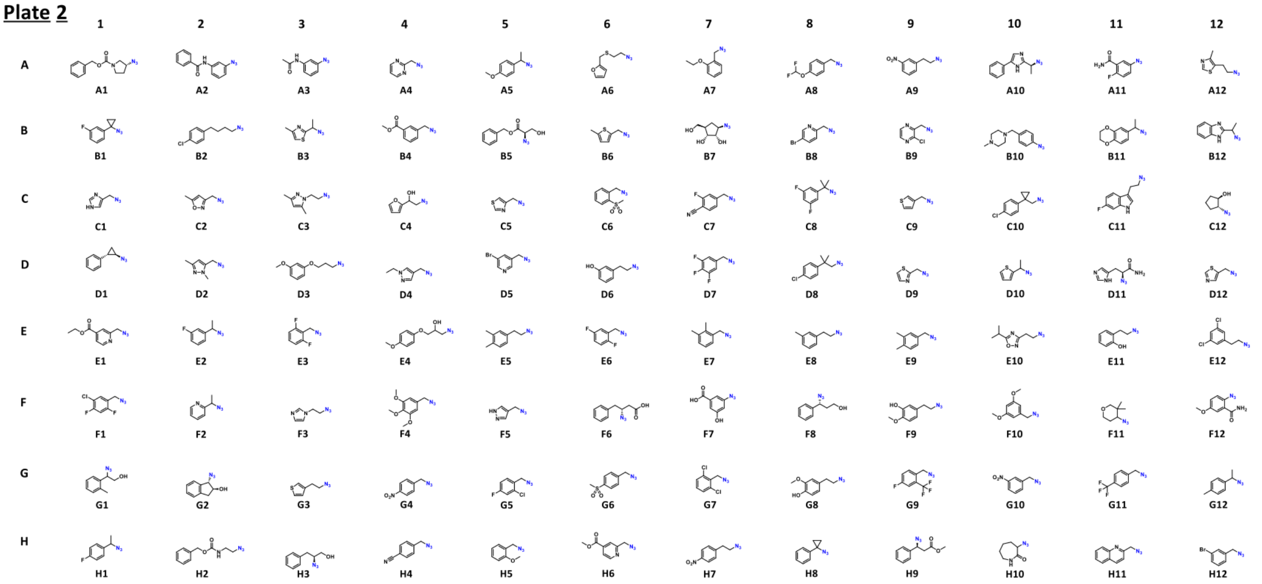

**Plate 3**

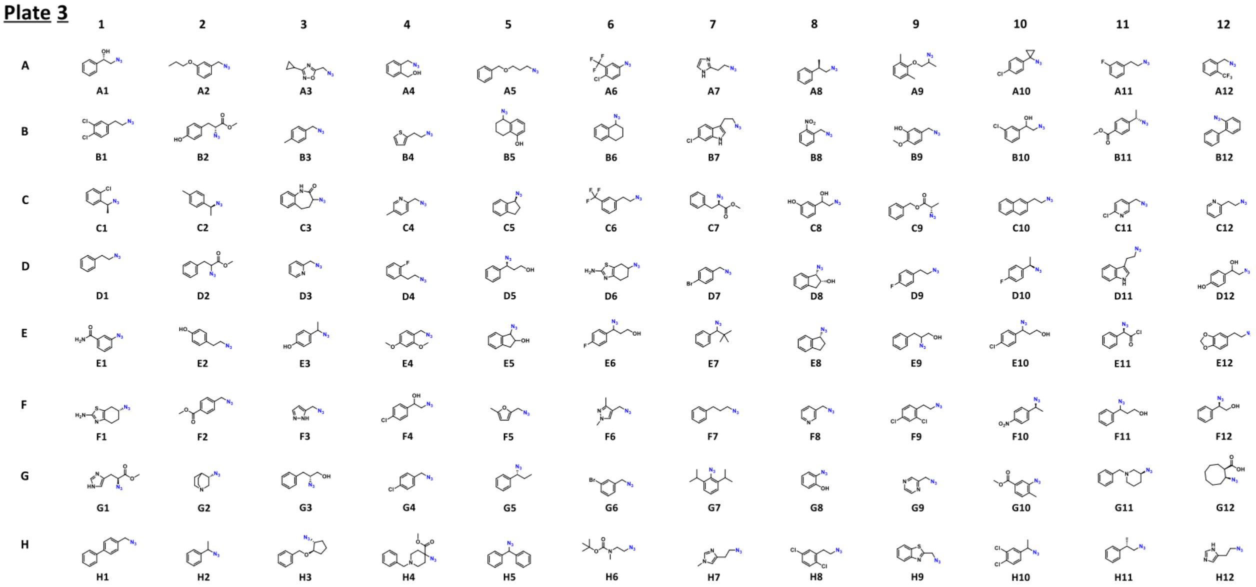


**Plate 4**

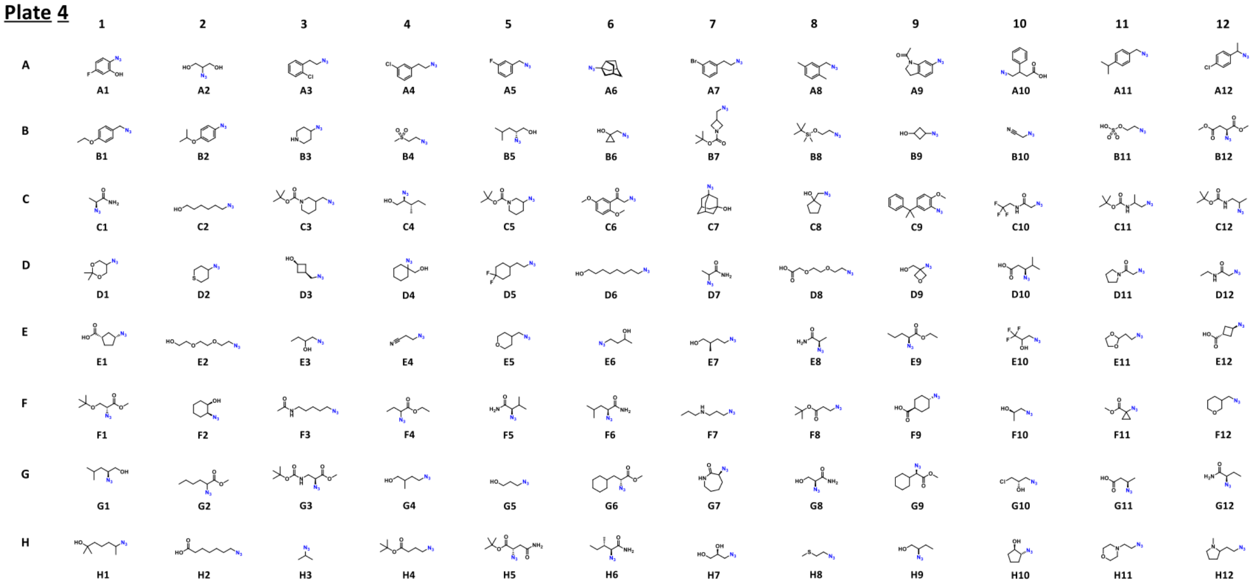


#### 404 Library of azido-tagged molecules (Enamine/Sigma)

1 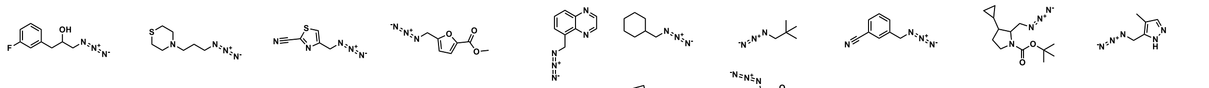

11 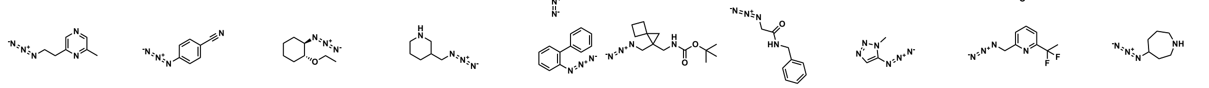

21 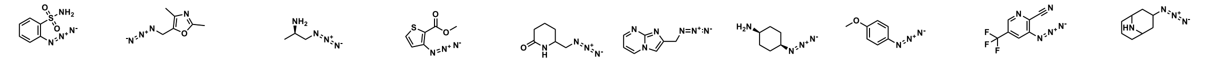

31 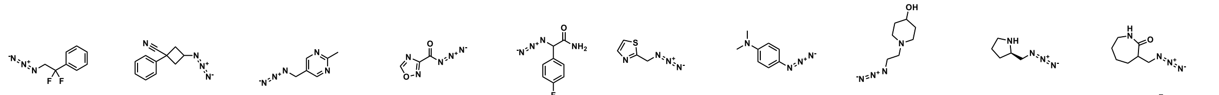

41 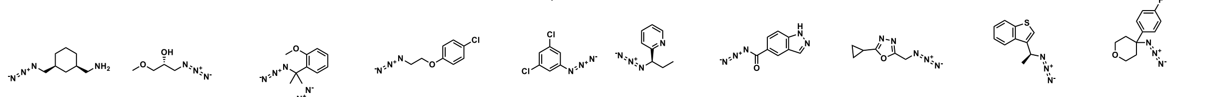

51 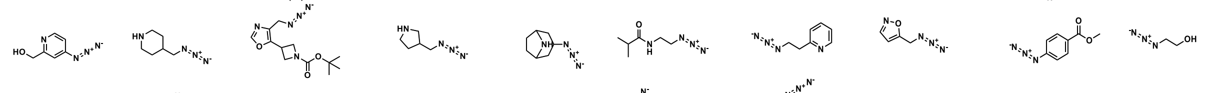

61 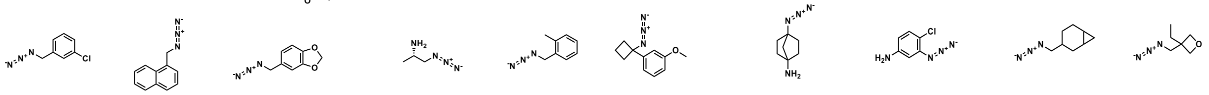

71 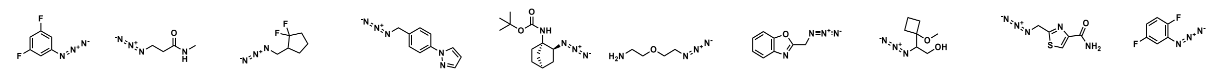

81 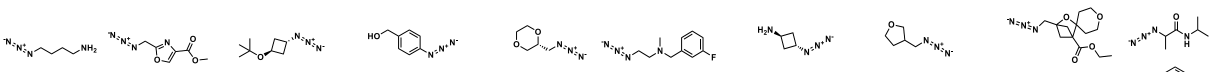

91 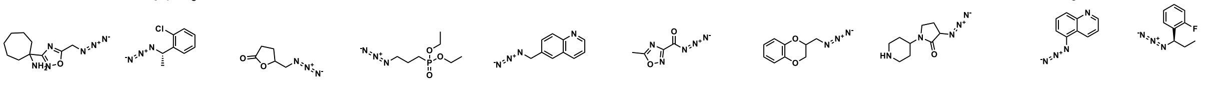

101 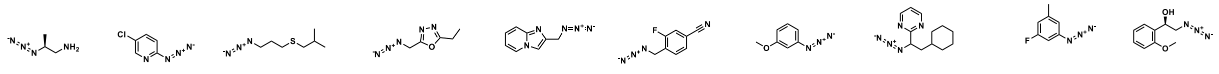

111 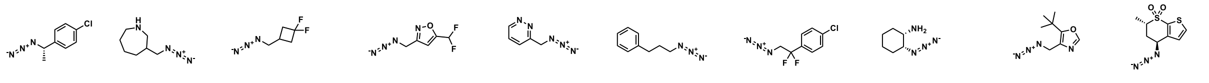

121 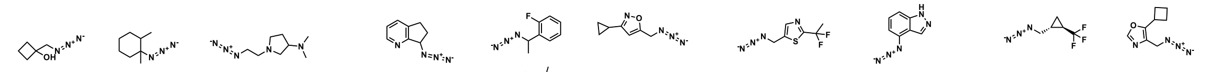

131 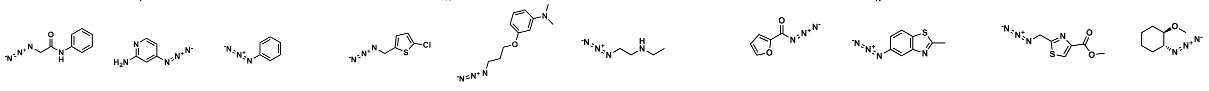

141 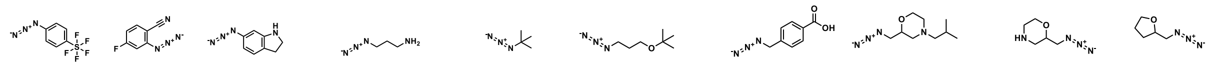

151 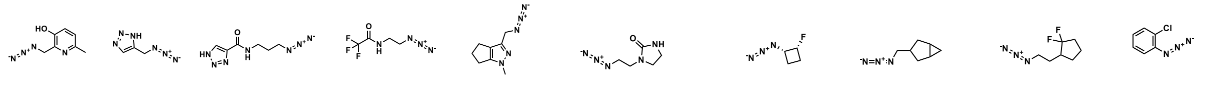

161 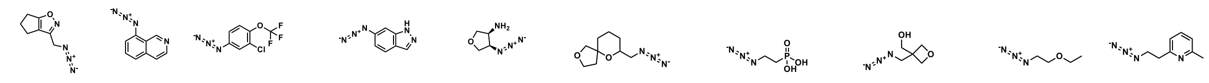

171

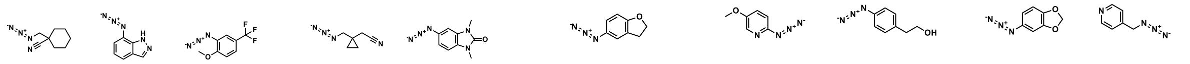

181

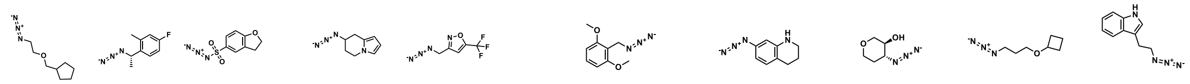

191

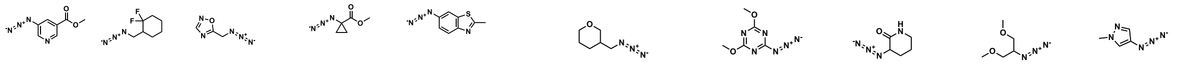

201

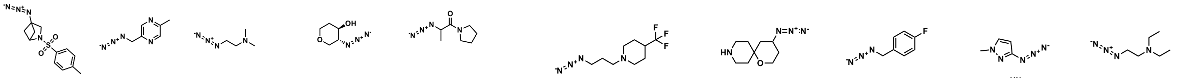

211

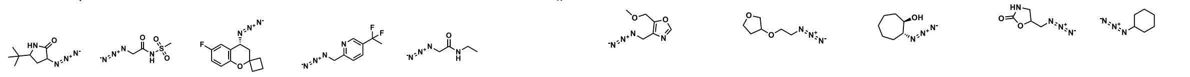

221

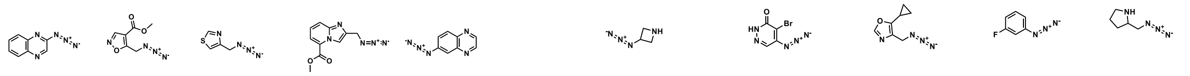

231

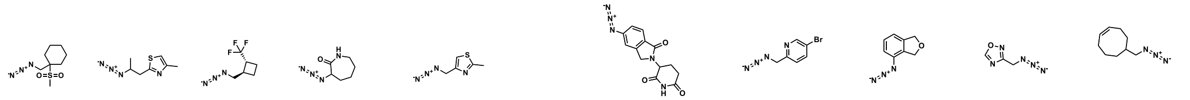

241

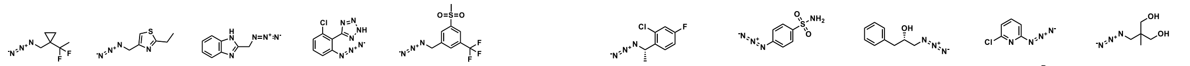

251

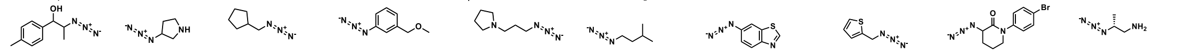

261

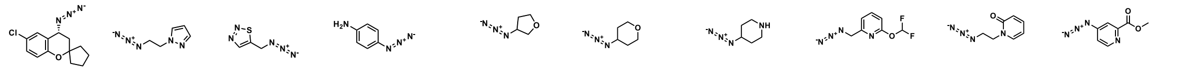

271

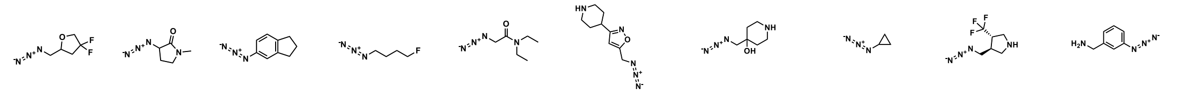

281

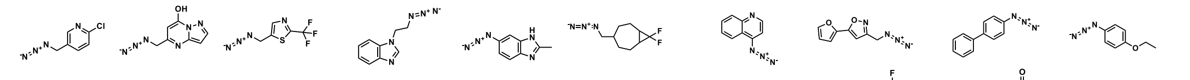

291

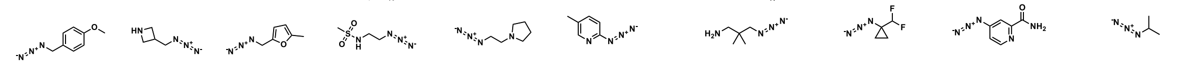

301

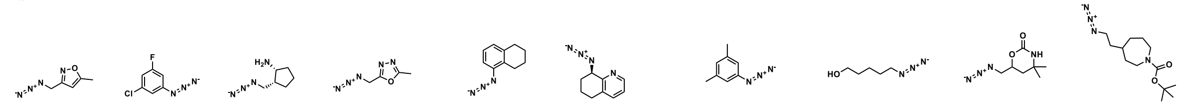

311

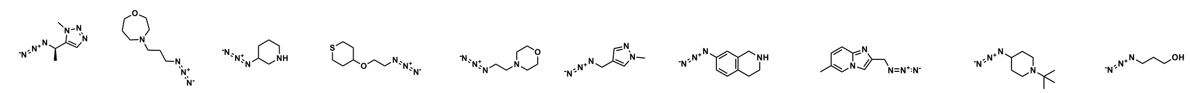

321

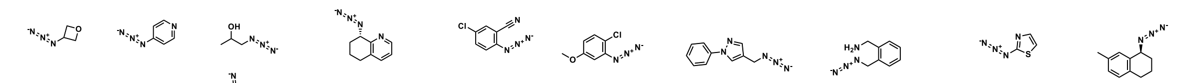

331

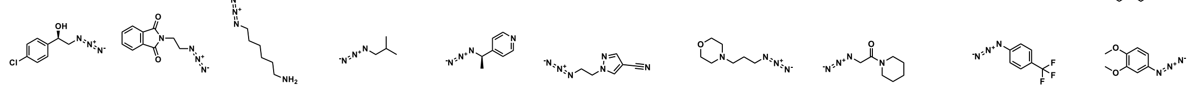

341

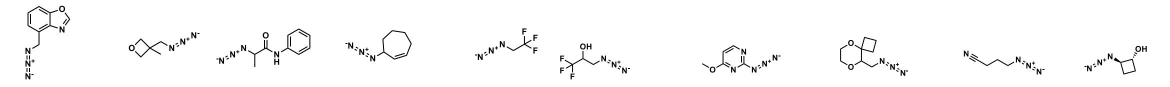

351

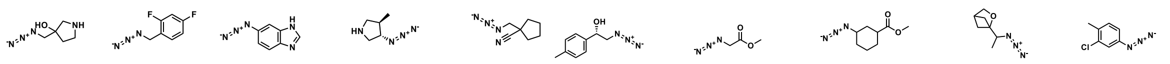

361

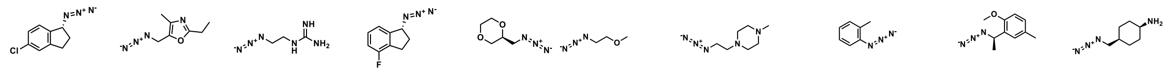

371

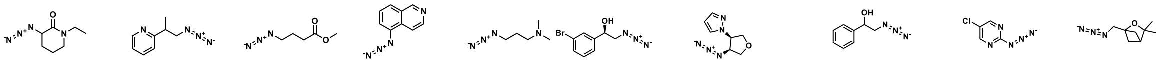

381

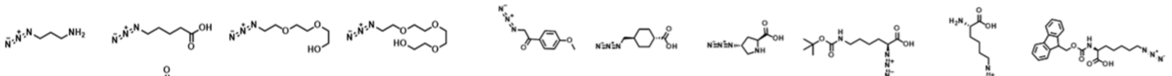

391

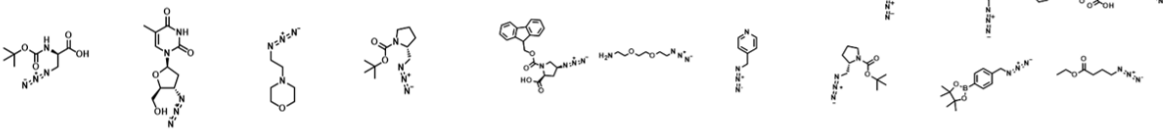

401

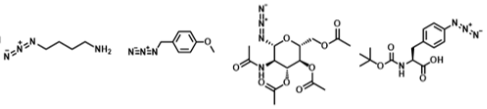

#### References


(1) Murrey HE, Judkins JC am Ende CW, Ballard TE, Fang Y, Riccardi K, Di L, Guilmette RE, Schwartz JW, Fox JM, Johnson DS. Systematic Evaluation of Bioorthogonal Reactions in Live Cells with Clickable HaloTag Ligands: Implications for Intracellular Imaging. *J. Am. Chem. Soc* **2015**, 137, 35, 11461–11475 DOI: https://doi.org/10.1021/jacs.5b06847

(2) Silverman SM, Moses JE, Sharpless B. Reengineering Antibiotics to Combat Bacterial Resistance: Click Chemistry [1,2,3]-Triazole Vancomycin Dimers with Potent Activity against MRSA and VRE. *A European Journal*, 2017, 23, 79-83.

(3) Phetsang W, Pelingon R, Butler MS, Pitt ME, Kaeslin G, Cooper MA, Blaskovich, MAT. Fluorescent Trimethoprim Conjugate Probes To Assess Drug Accumulation in Wild Type and Mutant Escherichia coli. *ACS Infect. Dis.* 2016; 2, 10, 688–701

(4) Paulen A, Gasser V, Hoegy F, Perraud Q, Pesset B, Schalk IJ, Mislin GLA. Synthesis and antibiotic activity of oxazolidinone–catechol conjugates against Pseudomonas aeruginosa*. Org. Biomol. Chem.,* 2015,**13**, 11401-11401

(5) Im WB, Choi SH, Park JY, Choi SH, Finn J, Yoon SH. Discovery of torezolid as a novel 5-hydroxymethyl-oxazolidinone antibacterial agent. *European Journal of Medicinal Chemistry*. 2011; 46 (4) 1027 – 1039.

(6) Zhao K, Ke W, Yin W, Li J, Qiang M, Ge Z. Facile Preparation and Radiotherapy Application of an Amphiphilic Block Copolymer Radiosensitizer. *ACS Macro Lett.* 2017 May 16;6(5):556-560.

(7) Meng G, Guo T, Ma T, Zhang J, Shen Y, Sharpless KB, Dong J. Modular click chemistry libraries for functional screens using a diazotizing reagent. *Nature*. 2019 Oct;574(7776):86-89.
